# VIRP1 bromodomain shapes nuclear condensate formation and has a positive effect on PSTVd accumulation

**DOI:** 10.64898/2026.06.10.730826

**Authors:** Bardani Eirini, Ostendorp Steffen, Andronis Christos, Asch Franziska, Ostendorp Anna, Katsarou Konstantina, Kehr Julia, Kalantidis Kriton

## Abstract

- Viroids are small, non-coding RNAs that rely on host factors for replication, intracellular trafficking and systemic movement. VIRP1, a Bromodomain and Extra-terminal domain (BET) protein, has previously been implicated in Potato spindle tuber viroid (PSTVd) infection, yet its precise role and mode of action remain unresolved.
- In this work, we highlight VIRP1 as the only Solanaceae BET protein containing a proline-rich domain, which overlaps with the PSTVd-binding site. VIRP1-deficient plants exhibit delayed flowering and increased ABA sensitivity, with differentially expressed genes enriched in stress-related pathways.
- VIRP1 forms condensates *in planta* and *in vitro*, consistent with phase-separation behaviour. Condensate morphology is altered by PSTVd RNA, deletion of the intrinsically disordered CTD and bromodomain mutations.
- VIRP1 is particularly important for the early establishment of PSTVd infection, while nuclear localization and bromodomain integrity are required for efficient viroid accumulation. By contrast, the disordered CTD region is dispensable for complementation of PSTVd accumulation.
- Our results support a model in which VIRP1 acts as a host nuclear factor that links stress-related functions, nuclear condensate formation and early viroid infection.

## INTRODUCTION

Viroids are the smallest known pathogens, consisting of 246-434-nt-long, circular, single-stranded RNA, able to cause important plant diseases. They are classified in two families, *Pospiviroidae* and *Avsunviroidae*, based on structural features and the replication site (Flores et al., 2015; Hammond, 2017; Kalantidis et al., 2025). Infection involves entry into cells and organelles, replication and systemic spread. Since viroids do not encode proteins (Katsarou, Adkar-Purushothama, et al., 2022), they rely on host factors for cell-to-cell transport, replication, and long-distance movement through the phloem (Flores et al., 2015; Navarro et al., 2021). Intramolecular base-pairing in the viroid RNA generates characteristic secondary structures that serve as host factor-binding sites or may protect viroids from host RNases (Flores et al., 2015; Navarro et al., 2021). Potato spindle tuber viroid (PSTVd, *Pospiviroid fusituberis*) is the type species of the nuclear-replicating family *Pospiviroidae*. Its 359-nt RNA forms a rod-like secondary structure typical for pospiviroids and contains a signal that directs it to the nucleus (Woo et al., 1999; Zhao et al., 2001), where it replicates via an asymmetric rolling-circle mechanism (Branch & Robertson, 1984). Its replication starts with transcription of the monomeric, circular, positive-strand RNA ((+)-RNA) by RNA polymerase II (Pol II) into multimeric, linear strands of negative polarity ((–)-RNA). These strands serve as replication intermediates for the production of multimeric, linear (+)-strands, which are then cleaved and circularized. While the (–)-strands reside in the nucleoplasm, (+)-RNAs are localized into the nucleolus (Flores et al., 2009; Kalantidis et al., 2025; Qi & Ding, 2003). Pol II is redirected by PSTVd to transcribe RNA instead of DNA, through a yet undefined mechanism, likely involving TFIIIA-7ZF (Dissanayaka Mudiyanselage et al., 2018, 2022; Y. Wang et al., 2016).

Our previous work identified Viroid-binding protein 1 (VIRP1) as a host factor that strongly interacts with (+)-PSTVd RNA (Maniataki et al., 2003; Martínez De Alba et al., 2003) and is crucial for PSTVd infectivity, as *VIRP1*-suppressed (*VIRP1*i) tobacco plants were found resistant to PSTVd mechanical inoculation (Kalantidis et al., 2007). VIRP1 contains a number of conserved regions, including a N-terminal P-loop (aa 21-28), a coiled-coil (aa 105-128), a proline-rich domain (aa 322-404) and a C-terminal serine-rich domain (aa 566-595) (Martínez De Alba et al., 2003). Its most striking feature, however, is the simultaneous presence of a bromodomain (BRD, aa 185-295) and a N-extra-terminal domain (N-ET, aa 399-480), classifying VIRP1 into the Bromodomain and N-extra-terminal domain (BET) protein family (Bardani et al., 2023; Filippakopoulos et al., 2012). The bromodomain is the “reader” of lysine acetylation mainly in histones (Dhalluin et al., 1999), whereas the N-ET domain recruits factors involved in transcriptional regulation (Rahman et al., 2011). PSTVd (+)-RNA binds VIRP1 through an atypical RNA-binding site (aa 313-403) (Martínez De Alba et al., 2003), which overlaps with the proline-rich domain. Despite its long-standing association with viroid infection (Kalantidis et al., 2007), the endogenous biological role of VIRP1 and the mechanistic basis of its pro-viroid activity remain unclear. BET proteins in animals function as chromatin readers and transcriptional regulators, linked to various cancer types, inflammation, and viral infections (C.-Y. Wang & Filippakopoulos, 2015). The human BET BRD4 is a central regulator of transcription elongation (Gaucher et al., 2012; Jang et al., 2005; Yang et al., 2005) that directly interacts with Pol II and phosphorylates its C-terminal domain (CTD) (Devaiah et al., 2012; Weissman et al., 2021). BRD4 also forms phase-separated condensates and is intrinsically disordered, with its CTD driving this property (Sabari et al., 2018). BRD4 condensate formation is considered to promote compartmentalization and concentration of biomolecules, aiding transcription (Kotekar et al., 2023). Additionally, BRD4 controls several long non-coding RNAs (lncRNAs), which can affect transcription (Pastori et al., 2015; Pistoni et al., 2021). Interestingly, human BET proteins have been reported to interact with viral proteins to mediate their retention on mitotic chromosomes (Ottinger et al., 2006; Schweiger et al., 2006; Wu et al., 2006; J. You et al., 2006).

In plants, BET proteins remain comparatively underexplored. *Arabidopsis thaliana* (*A. thaliana*) encodes 12 BETs, named as GENERAL TRANSCRIPTION FACTOR GROUP E (GTE) 1-12, which form 3 phylogenetic clusters: GTE2-7, GTE8-12 and GTE1/GTE6 (Airoldi et al., 2010). GTE1 is involved in seed germination (Duque & Chua, 2003), while GTE6 regulates mature leaf shape (Chua et al., 2005). GTE4 participates in mitotic cell cycle retention (Airoldi et al., 2010) and plant immunity (Zhou et al., 2022). GTE9, GTE10 and GTE11 mediate abscisic acid (ABA) responses (M. J. Kim et al., 2009; Misra et al., 2018). Recently, the closely related GTE2 and GTE7 were shown to monitor Pol II activity upon heat stress (Zheng et al., 2025) and regulate sulfur homeostasis in Arabidopsis (Luo et al., 2025). Regarding Solanaceae, there is currently no information about BET proteins and their endogenous functions (Bardani et al., 2023). Understanding VIRP1’s endogenous role will not only shed light to the functions of BET proteins in plants, but could further uncover putative host mechanisms exploited by viroids to fulfill their infection cycle.

Several roles have been proposed for VIRP1 in the viroid infection cycle. Given its nuclear localization, it was suggested that VIRP1 may facilitate PSTVd nuclear import (Kalantidis et al., 2007; Martínez De Alba et al., 2003). This hypothesis was supported by demonstrating that VIRP1 enhances nuclear localization of Q-satRNA of Cucumber mosaic virus (Family *Bromoviridae*, species *Cucumovirus CMV*) (Chaturvedi et al., 2014). As satellite RNAs exhibit similarities to viroids (Bruening, 1991), it was hypothesized that VIRP1 mediates pospiviroid nuclear import. Application of exogenous, purified VIRP1 mildly enhanced nuclear localization of Citrus Exocortis viroid (CEVd, *Pospiviroid exocortiscitri*) in a heterologous system using onion cells (Seo et al., 2021). Further experiments in Arabidopsis suggested a model where PSTVd forms a complex with GTE7 and IMPα-4 for nuclear import (Ma et al., 2022). However, whether VIRP1’s main or only role is to facilitate viroid nuclear entry is unclear yet. As VIRP1 contains a bromodomain, it could facilitate not only viroid transport into the nucleus but also subnuclear trafficking and association with chromatin and chromatin-related host factors. The work presented here addresses three questions: what endogenous functions VIRP1 fulfils in *N. benthamiana*; how VIRP1 domain architecture contributes to its (sub)nuclear localization and which VIRP1 features are required to support PSTVd infection. Our results indicate that VIRP1 is a stress-related BET protein that forms phase-separated subnuclear condensates and promotes early viroid infection through a bromodomain-dependent nuclear function.

## MATERIALS AND METHODS

### Plants

We used wild-type (WT) and *VIRP1*-suppressed (*VIRP1*i) *N. benthamiana* plants (Kalantidis et al., 2007) and *Columbia* (*Col-0) A. thaliana* plants. *N. benthamiana* plants constitutively expressing a human codon-optimized *Streptococcus pyogenes Cas9* were used for CRISPR/Cas9 mutagenesis, kindly provided by José Antonio Darós in IBMCP (Valencia, Spain) (Uranga et al., 2021). *N. benthamiana* plants were grown in greenhouse conditions. *Col-0 A. thaliana* plants were grown in a growth chamber at 22 °C, with 60% humidity, 16h light/8h dark, light intensity ∼120 µmol m^− 2^ s^− 1^. Images were captured using a Nikon D5100 camera (Nikon, Tokyo, Japan).

### Plasmids

The *pLB-PVX::sgRNA* construct was generated by amplifying a *VIRP1*-targeting sgRNA sequence and cloning into MluI-linearized pLB-PVX (Uranga et al., 2021). For all clonings, we used the *Solanum lycopersicum* (cultivar Rentita) *VIRP1* cDNA (GenBank: AJ249595.1). To clone *VIRP1* into pEarleyGate104 and pEarleyGate201, the sequence was first cloned into pENTR4-dual to generate pENTR4-VIRP1. This construct was also used for Gateway cloning into the destination vector pB2GW7 for stable *A. thaliana* transformation (*35S::VIRP1*). For the amplification of C-terminally truncated fragments, specific reverse primers were used. To clone VIRP1 into pEarleyGate103, it was first cloned into pENTR4-dual. NES-tagged VIRP1 (LALKLAGLDI) (Wen et al., 1995) was introduced into pENTR4-dual and then recombined it into pEarleyGate104. To clone *HA:VIRP1* (WT, NLS mutants, BRD mutants) into PVX-GW, the sequence was first amplified from *pEG201-VIRP1* and then cloned into pENTR4-dual. Respectively, to clone *HA:NES:VIRP1* into PVX-GW, the sequence was first amplified from *pEG104-NES:VIRP1* and then cloned into pENTR4-dual. All inserts were then cloned into the respective gateway vectors via LR clonase II (Thermo Fisher Scientific) according to the manufacturer’s instructions. All primers used are listed in **Table S1**. To introduce VIRP1 mutations, overlapping primers were used **(Table S1)**, designed using QuickChange Primer Design (agilent.com/store/primerDesignProgram.jsp) online software. After amplification (Kapa HiFi, Roche) and DpnI (New England Biolabs) digestion, T4 ligase (New England Biolabs) was used for circularization of the linear fragment. To clone tomato VIRP1 into pET16b vector for protein purification (His-tag), conventional cloning was performed using NdeI and XhoI restriction enzymes. To clone VIRP1 into pET28a-eYFP-SSG-MCS vector for protein purification, conventional cloning was performed using EcoRI and XhoI restriction enzymes **(Table S1)**. To generate the FIBRILLARIN:RFP construct, the sequence was first amplified from *N. benthamiana* cDNA and cloned into pENTR4-dual **(Table S1)**. The construct was used for gateway cloning into the destination vector pGWB454. The vectors PVX-GW, PVX:GFP and pSoup (Lacorte et al., 2010) were kindly provided by Dr. Christiano Lacorte (EMBRAPA). The construct PSTVd^NB^-AJ634596, expressing an infectious PSTVd dimer, was kindly provided by Dr. De Alba and Dr. Flores (Institute for Cellular and Molecular Plant Biology-IBMCP). For the synthesis of a PSTVd-detecting DIG-labeled RNA probe, the plasmid pHA106 was used (Tabler et al., 1992).

### Plant transformation

For the transformation of *A. thaliana* plants, we followed the method described in (Narusaka et al., 2010). Briefly, *Col-0 A. thaliana* plants were grown until the emergence of flowers and were dipped in GV3101 *A. tumefasciens* suspension (saturated culture) in 5% sucrose and 0.02% Silwet L-77. Plants were then placed in the dark for 24h. Seeds were harvested and transformed plants were screened using 120 μg/mL Basta (glufosinate ammonium, Bayer).

### Selection of single insert transformants

For the selection of transgenic *A. thaliana* plants carrying one insert of the *35S::VIRP1* transgene, Basta-resistant T_2_ seeds were planted in pots containing 1:1 humus: vermiculite and placed for 2-3 days at 4 °C in the dark. After transferring to a growth chamber (see Plants), seedlings were sprayed with 120 μg/mL Basta 2 and 3 weeks later. T_2_ plants with a 3:1 ratio of resistant: sensitive progeny were selected.

### Generation of *virp1 N. benthamiana mutants*

*To* generate CRISPR/Cas9 *N. benthamiana* mutants, the method described by Uranga et al (Uranga et al., 2021) was used, with modifications (Bardani et al., 2025). The sgRNA targeting the two *VIRP1* paralogs was designed using Benchling CRISPR Guide RNA Design Tool (benchling.com/crispr). *Cas9*-expressing *N. benthamiana* plants were inoculated with GV3101 agrobacteria carrying *pLB-PVX::sgVIRP1-tFT* (2 leaves per plant at the 4-6 leaf stage). Two weeks later, symptomatic leaves were isolated and used for plant regeneration. PVX-infected regenerated plants (F_0_) were screened through PCR sequencing and ICE analysis and mutants were selected for further analysis. Healthy F_1_ plants were screened through PCR sequencing and mutation analysis using Inference of CRISPR Edits (ICE, Synthego, ice.editco.bio/) and homozygous mutants were selected. Plant regeneration was performed based on Horsch et al. (Horsch et al., 1985) and has been described elsewhere (Bardani et al., 2025).

### Genomic DNA (gDNA) extraction

The CTAB protocol was followed, with modifications (Murray & Thompson, 1980). Briefly, 30 mg of leaf tissue were frozen and ground using liquid nitrogen and leaf tissue powder was diluted in 300 μl of CTAB buffer (2% CTAB, 100 mM Tris-HCl pH 8, 20 mM EDTA, 1.4 M NaCl, 1% PVP). After a 20 min incubation at 65 °C, an equal volume of chloroform was added. After centrifugation and isolation of the supernatant, DNA was precipitated with 0.8 volumes of isopropanol, washed with 70% ethanol and resuspended in 50 μl of distilled water.

### Sequencing of PCR fragments for mutant genotyping

DNA fragments were amplified by PCR using Taq polymerase (Minotech, Greece) and primers flanking the sgRNA target area **(Table S1)**. PCR fragments were sequenced by Genewiz, Azenta Life Sciences (Germany).

### Phenotypic *measurements*

Six weeks old *N. benthamiana* plants were used for the estimation of plant height and dry weight. Plant height was estimated as the vertical distance from the soil surface at the base of the stem to the highest point of the main shoot. Measurements were taken using a ruler and recorded in centimeters. Measurements were performed on n=10 plants. For dry weight estimation, whole plants were dried in a ventilated oven at 70 °C until constant weight was achieved (48-72h). Weight was measured using an analytical balance and recorded in grams. Measurements were performed on n=10 plants. Flowering time was estimated as the number of days from sowing until the appearance of the first visible flower bud on the main shoot. Three biological replicates were used with a total plant number of n=23 (WT), n=24 (*virp1*) and n=26 (*VIRP1i*).

### ABA treatment

To estimate ABA-mediated inhibition of germination in *N. benthamiana* seeds, *virp1, VIRP1i* and WT seeds were sterilized using 10% bleach and placed on Petri dishes containing MS medium (Murashige and Skoog, 4.4 g/L MS, 5.5 g/L plant agar, pH 5.6) with 0, 2.5 or 5 μM ABA (Duchefa Biochemie). For RT-qPCR, WT seedlings were vertically grown on MS for 2 weeks and then transferred to 0 or 10 μM ABA plates for 3 hours. Whole seedlings were isolated and a pool of 3 plants was considered as a biological replicate. *Col-0* and *35S::VIRP1 A. thaliana* seeds were first vernalized at 4 °C for 48h, sterilized and placed on ½ MS with 0, 0.5 or 1 μM ABA in a growth chamber (see Plants). Photos were taken 2 and 3 weeks after placement.

### Agroinfiltration

C58C1 or GV3101 *Agrobacterium tumefaciens* strains carrying the desired constructs were grown in LB medium with the appropriate antibiotics for selection. Agroinfiltration was performed in young *N. benthamiana* plants at the age of 4-6 leaves, as previously described (Katsarou, Kryovrysanaki, et al., 2022). An optical density at 600 nm (OD_600_) of 0.1 was used for PSTVd and PVX infections (see Plasmids). For YFP-and HA-constructs, an OD of 0.5 was used.

### Mechanical inoculation

*N. benthamiana* plants at the 4-leaf stage were inoculated with infectious sup from 4 week-PSTVd-agroinfected *N. benthamiana* plants, using carborundum (Prolabo, VWR). The infectious sup was homogenized in 50 mM NaPO_4_ pH 6.8 in a 1:4 ratio. For each plant two leaves were inoculated, with 50 μl each.

### RNA extraction and Northern blot

RNA extraction from young leaves and Northern blot analysis have been described previously (Katsarou, Kryovrysanaki, et al., 2022). Briefly, 4 μg of total RNA were separated on a denaturing agarose gel (1.4% agarose, 0.7% formaldehyde, 1x MOPS, 7 μg EtBr per 100 ml gel), in denaturing buffer (0.7% formaldehyde, 1x MOPS). After equilibration in 2x SSC, transfer was performed onto a 0.2 μm Nylon membrane (Whatman, GE Healthcare, USA) in 10x SSC via capillary transfer for 16-20 h. Crosslinking was achieved through UV radiation (Stratalinker, Stratagene, USA) with 1200 μJ cm^-2^ total energy. A PSTVd RNA(-) DIG-labelled probe (DIG RNA labelling mix, Roche) was generated as described previously (Katsarou, Kryovrysanaki, et al., 2022) by T7 *in vitro* transcription from a HindIII-linearized pHa106 plasmid (Tabler et al., 1992) and hybridization was performed at 65 °C in hybridization buffer (5x SSC, 1% SDS; 1× Denhardt’s; 25011mg/ml tRNA; 50% formamide). CDP-star (Roche) was used for the detection according to manufacturer’s instructions. Band intensity was quantified with the software Azure Sapphire and values were normalized using ribosomal 25S rRNA as a control.

### Reverse transcription and quantitative PCR

For RT-qPCR, tissue was isolated from young leaves, shoots, flower buds or roots of 6-week-old plants. 5 μg of total RNA were treated with DNaseI (Roche). After purification with phenol/chloroform, RNA was reverse-transcribed to cDNA using Minotech RT and random hexamers (Invitrogen), according to the manufacturer instructions. A 1:5 dilution of cDNA was used for gene amplification with KAPA SYBR® FAST qPCR Kit (KapaBiosystems) on the CFX Connect^™^ Real-Time PCR detection System (Bio-Rad). Annealing and extension were carried out in one step at either 60-62 °C for 20s, depending on the primers. Samples were processed in triplicates and efficiency of each primer set was calculated using plasmid standard curves in the range of 90-110%. L23, PP2A and FBOX were used as reference genes (Liu et al., 2012), validated in our samples and conditions using BestKeeper algorithm (Andersen et al., 2004). The results were normalized to the value of WT plants, which was arbitrarily fixed at 1. Expression analysis was performed using the Pfaffl method (Pfaffl et al., 2004). All primers are listed in **Table S1**.

### Protein extraction and western blot

Plant tissue from agroinfiltrated leaves was isolated 3 days post infiltration (dpi) for transient expression analysis. Plant tissue from systemic leaves was isolated 7 dpi for virus-mediated expression analysis. Total protein was extracted from liquid nitrogen-ground young leaves, adding 4:1 protein extraction buffer (50 mM Na_2_HPO_4_, 1 mM EDTA, 0.1% SDS, 0.1% Triton X-100, 10 mM DTT, 0.7 μl/ml β-ME). Proteins were analyzed on an 8% SDS polyacrylamide gel and transferred on ProtraN membrane (Whatman). For detection of HA-tagged and YFP-tagged proteins, we used anti-HA (1:2000, C29F4, Cell signaling) and anti-YFP (1:10000, AS11 1775, Agrisera), respectively, according to manufacturer’s instructions. Sapphire biomolecular imager (Azure Biosystems) and Azure Sapphire capture software were used for detection. Ponceau staining was used as a loading control.

### RNA-protein immunoprecipitation

Plant tissue from leaves co-agroinfiltrated with HA:VIRP1 and PSTVd RNA was isolated 3 dpi and UV-crosslinked (3 × 375 mJ/cm^2^). 200mg were used for extraction (150mM NaCl, 100mM Tris pH 8.0, 1mM EDTA, 10% glycerol, 0.2% NP-40, 1mM DTT, 1:100 protease inhibitors (Sigma) and 1:100 RNase inhibitor (Takara) and incubated with anti-HA (1:500, C29F4, Cell signaling) for 2h, then with magnetic beads for 1h (Pure Proteome, Merck). Washes were made with the same buffer. Proteinase K (Invitrogen) was used to release the bound RNA, followed by RNA extraction (see above). After cDNA synthesis (see above), the presence of immunoprecipitated PSTVd was detected with PCR **(Table S1)**.

### Protein expression and purification

6xHis-eYFP-tagged and 6xHis-tagged VIRP1 proteins were expressed in *E*.*coli* BL21 Rosetta2 (DE3) cells (Merck) in 400 mL autoinduction medium (Studier, 2005). The cultures were harvested by centrifugation at 5000xg and lysis was carried out in 60 mL of lysis buffer (50 mM Tris-HCl pH 7.5, 300 mM NaCl, 10 mM imidazole, 1 mM AEBSF, 1 mM DTT, 1x cOmplete protease inhibitor tablet) supplemented with 1 mg/mL lysozyme, 0.1 mg/mL DNase I and 0.1 mg/mL RNAse A. After sonication of the resuspension on ice (Branson Sonifier 250, 8 times 30 s, duty cycle 50%, output 6), the lysate was centrifuged at 35000xg at 12°C. Purification was achieved in two consecutive steps using nickel affinity chromatography followed by size exclusion chromatography. The centrifuged lysate was loaded onto a 5 mL HisTrap FF column (Cytiva) connected to an ÄKTA prime plus FPLC system run at 1 mL/min. The loaded column was washed with lysis buffer containing 1 M NaCl and proteins were eluted using an imidazole gradient from 0 to 1 M in buffer B (50 mM Tris-HCl pH 7.5, 300 mM NaCl, 1 M imidazole) over 15 column volumes. Fractions containing desired proteins were pooled and dialyzed overnight in 2 L of dialysis buffer (50 mM Tris-HCl pH 7.5, 300 mM NaCl, 1 mM DTT) in a SpectraPor dialysis membrane (MWCO: 10000 Da). Thrombin (20 u, Cytiva) was added to remove the 6xHis tag. On the next day, dialyzed proteins were concentrated to a final volume of 2 ml using an Amicon Ultra centrifugal filter device (MWCO: 10000 Da, Millipore) and subjected to size exclusion chromatography using a HiLoad 16/600 Superdex 200 pg column (Cytiva) equilibrated in gel filtration buffer (25 mM Tris-HCl pH 7.5, 200 mM NaCl, 1 mM DTT). The quality of all purified proteins was further accessed by MALDI-TOF mass spectrometric measurements using a Bruker Ultraflex extreme MALDI-TOF/TOF mass spectrometer and by UV/Vis spectrometry using a NanoDrop One photospectrometer.

### Phase separation assay

To induce protein condensation, purified proteins were diluted to a final concentration of 20 μM with 1x condensation buffer (50 mM Tris-HCl pH 7.5, 150 mM NaCl). Subsequently, 10% (w/v) PEG3350 was added as a crowding agent 1:1 with VIRP1 and incubated for 2 min prior to the measurements. 5 μL were transferred onto a glass slide (Labsolute, Th. Geyer GmbH, Germany) and carefully covered with a siliconised circular cover slide (22 mm diameter, Jena Bioscience, Germany). Condensate formation was observed by fluorescence microscopy using a BZ-X800 fluorescence microscope (Keyence, Germany), with a 100× oil immersion objective equipped with an eYFP filter system (Chroma, USA). All microscopic analyses were performed at an exposure time of 17 ms as a single z-plane without any black balance.

### Microscale Thermophoresis (MST)

PSTVd (+) and (-) RNA was *in vitro* transcribed from a HindIII-linearized pHa106 plasmid (Tabler et al., 1992) by T7 and SP6 RNA polymerase (Ribomax, Promega), respectively, using Cy5-labeled UTP. Titration series were prepared with the unlabeled protein using MST buffer (25 mM Tris-HCl pH 8.0, 150 mM NaCl, 1 mM DTT, 0.1% Tween-20 and 0.1 mg/ml BSA) and a starting concentration of 50-75 µM. Labeled RNA was added 1:1 with a final concentration of 20 nM for labeled RNA in each reaction mixture. Everything was mixed well by pipetting and centrifuged down in a microcentrifuge. Protein and RNA were incubated for around two minutes. Monolith™ Series standard capillaries (Nanotemper technologies, Germany) were used to load 16 samples and placed in the sample slide of the Monolith NT.115 MicroScale Thermophoresis device (Nanotemper technologies). Binding affinity measurements were performed at 40% LED and medium MST-power for the first measurement of each individual RNA. Each measurement was done at least three times. For each repetition a new titration series was prepared.

### Microscopy

For scanning electron microscopy, leaf samples from 6 weeks old plants were isolated and prepared as described previously (Bardani et al., 2025). Observations were made using a scanning electron microscope (JEOL, model JSM-IT700HR, Tokyo, Japan) at an operating voltage of 80 kV.

For fluorescent microscopy, the optical microscope Eclipse E800 (Nikon) was used, equipped with a UV lamp and the suitable filter. Photos were taken using ProgRed CF (Jenoptik) and software ProgResCapture Pro V2.8.8.

For confocal microscopy, images were captured with a Leica SP8 laser scanning confocal microscope. Leaf samples were imaged with a 40x oil objective. Processing was performed with FiJi (imagej.net/software/fiji/). YFP was excited using a 514-nm argon laser and emission was detected at 520-550 nm. DAPI was excited using a 405-nm argon laser and emission was detected at 461 nm.

### RNA sequencing

Two different RNA sequencing experiments were conducted, the first using *VIRP1i* and WT *N. benthamiana* plants and the second using *virp1* homozygous mutants (*virp1-1*) and *Cas9*-expressing plants. In each experiment, 3 plants per genotype were analyzed. Young leaves were isolated at the stage of 6 leaves and RNA was extracted and purified with phenol-chloroform.

### Bioinformatic analysis

Library construction was performed using the Lexogen QuantSeq 3’ mRNA-Seq Library Prep Kit FWD for Illumina, generating single-end reads of 75 bp average size. The quality of the raw reads before and after the QC steps was assessed with FastQC (Andrews S., n.d.). Quality and adapter trimming was performed with BBDuk (n.d.) using the settings recommended by the manufacturer, i.e. k=13 ktrim=r useshortkmers=t mink=5 qtrim=r trimq=10 minlength=20. The draft genome v1.0.1 of *Nicotiana benthamiana* (Bombarely et al., 2012) obtained from Sol Genomics Network (solgenomics.net) was indexed using hisat2-build. Reads obtained after trimming were aligned to the indexed Niben101 genome using hisat2 (D. Kim et al., 2019) with --score-min L,0,-0.5. Aligned reads were counted with htseq-count (Anders et al., 2015). Differential expression analysis of RNA-seq data was carried out using the edgeR workflow via the R package SARTools (Varet et al., 2016). Annotation and functional enrichment analysis of Differentially Expressed genes was performed either via the agriGOv2 (Tian et al., 2017) server or using the clusterProfiler (Yu et al., 2012) Bioconductor R package.

### Additional software

Figures were assembled using Photoshop CS5 (Adobe Systems Incorporated, California, USA). FiJi (imagej.net/software/fiji/) was used for image measurements. GraphPad Prism6 (GraphPad Software, San Diego, California, USA) was used for graph representations and statistical analysis. MEGA11 was used for sequence alignment and phylogenetic analysis (Neighbour Joining, bootstrap =1000). Shiny GO was used for gene ontology analysis (bioinformatics.sdstate.edu/go/). Fuzdrop was used for phase separation prediction (fuzdrop.bio.unipd.it/predictor) (Hatos et al., 2022). Alphafold was used for 3D protein structure prediction (alphafold.ebi.ac.uk/). NLStradamus (moseslab.csb.utoronto.ca/NLStradamus/) (Nguyen Ba et al., 2009) and cNLS Mapper (nls-mapper.iab.keio.ac.jp) (Kosugi et al., 2009) were used for NLS prediction. For the prediction of bromodomain active center structure, Swiss-PdbViewer 4.1 was used.

## RESULTS

### VIRP1 is a distinctive Solanaceae BET protein

VIRP1, carrying a bromodomain and a N-extra-terminal domain, belongs to the BET protein family; however, there is no information on other BET proteins in Solanaceae and how they are related to VIRP1. Therefore, we aimed to investigate the total BET family members of tomato and *N. benthamiana* in terms of length, domain organization and phylogenetic relationships. Using the Sol Genomics database, we identified 10 BET proteins in tomato and 21 in the allotetraploid *N. benthamiana*, showing homology to VIRP1 **(Fig. 1a-b)**. All 10 tomato BETs contained a bromodomain and a N-extra-terminal domain, as well as additional domains, e.g. coiled-coil domain or low-complexity regions rich in specific amino acid residues **(Fig. 1a)**. We noticed a similar domain architecture in the 21 BETs of *N. benthamiana*, with two of them lacking the NET domain (Niben101Scf08142g00004.1 and Niben101Scf13540g04027.1) **(Fig. 1b)**. Phylogenetic analysis of BET proteins in tomato, *N. benthamiana* and *A. thaliana* revealed three distinct clusters of BET proteins (I-III). The first cluster (I) contained homologs of *A. thaliana* GTE8-12; the second cluster (II) contained homologs of GTE1 and GTE6, whereas the third cluster (III) contained orthologs of GTE2, GTE3, GTE4, GTE5 and GTE7 **(Fig. 1c)**. VIRP1 grouped with the third cluster (III) and showed the closest homology to GTE7 and GTE2 **(Fig. 1c)**, with tomato VIRP1 having an amino acid similarity of 44.4% with GTE7 and 42.2% with GTE2, respectively **(Table S2a). Table S2b** lists the 10 tomato BET proteins, along with their chromosome locations, protein lengths, molecular weights, and isoelectric points. The closest homologs of each gene in *N. benthamiana* and *A. thaliana* are also indicated. Noteworthy, the proline-rich domain is unique to tomato VIRP1 (Solyc01g106280.3.1) and to the 2 *N. benthamiana* paralogs (Niben101Scf02392g00001.1 and Niben101Scf18637g02010.1). This low-complexity region, residing between the bromodomain and the NET domain **(Fig. 1a-b)**, overlaps with the RNA-binding site, with which PSTVd RNA interacts (Martínez De Alba et al., 2003). We conclude that tomato and *N. benthamiana* VIRP1 proteins belong to protein families with several members that show similarities in domain arrangement and biochemical properties. However, VIRP1 is the only BET protein carrying a proline-rich domain coinciding with the PSTVd-interacting site.

**Fig. 1.**
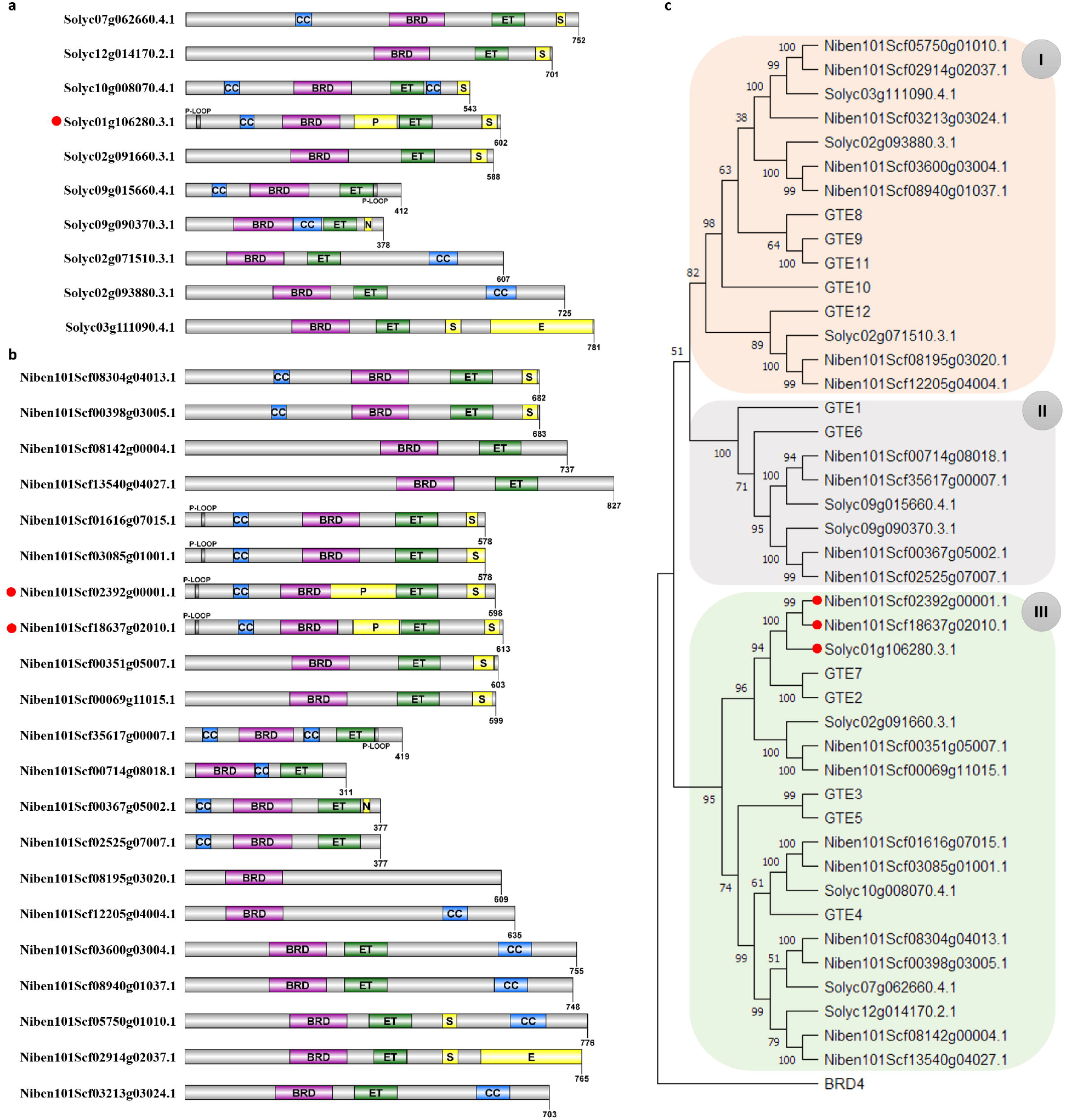
Domain architecture and phylogenetic analysis of BET proteins in tomato and *N. benthamiana*. **a**,**b**. Domain architecture of BET protein family in tomato (a) and *N. benthamiana* (b), respectively; CC: coiled-coil; BRD: bromodomain; ET: extra-terminal domain; S: serine-rich domain; P: proline-rich domain; N: asparagine-rich domain; E: glutamine-rich domain. **c**. Phylogenetic tree of tomato and *N. benthamiana* BET proteins, compared with the Arabidopsis BET homologs; colored boxes indicate the three subgroups of BET proteins; human BRD4 was used as a root; analysis was performed using the Neighbor-Joining algorithm (bootstrap=1000); red dots indicate VIRP1 homologs.

### VIRP1 contributes to endogenous stress-related gene regulation

Our previously described *VIRP1*i plants have been valuable tools to demonstrate the crucial role of VIRP1 in viroid infectivity (Kalantidis et al., 2007). However, these plants may still express low levels of *VIRP1*, which might suffice for at least some physiological function. Additionally, *VIRP1*i plants complicate complementation experiments, as any exogenous VIRP1 construct will be targeted by the transgenic hairpin. Here, we aimed to generate and evaluate *virp1 N. benthamiana* mutants using a CRISPR/Cas9 approach based on the delivery of the sgRNA through a PVX-based vector (Bardani et al., 2025; Uranga et al., 2021); the method is briefly described in **Fig. S1a**. As an allotetraploid, *N. benthamiana* carries two copies of the *VIRP1* gene, therefore a sgRNA was designed to target both paralogs at the 5’ end of the coding sequence **(Fig. 2a)**. In **Fig. S1b** the percentage of edits in PVX-infected regenerated plants (F_0_) is shown, while in **Fig. S1c** the percentage of knockout mutations in healthy F_1_ plants is indicated. In **Fig. 2b**, the homozygous *virp1* mutant (*virp1-1*) is represented, containing 2 types of mutations: a deletion of 17-nt and an insertion of 1-nt. This mutant was used for all subsequent experiments.

**Fig. 2.**
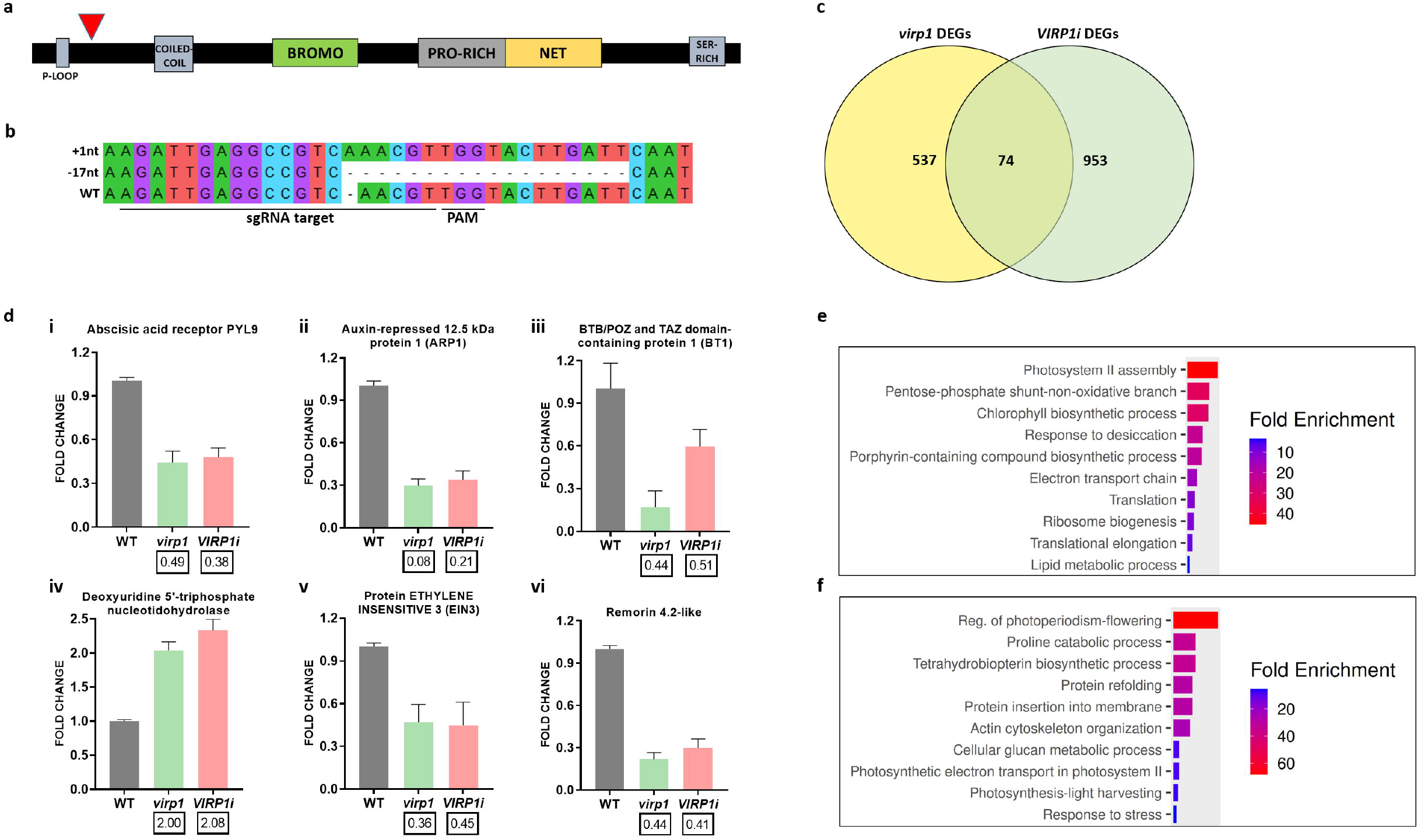
Generation of *virp1* mutants and RNA sequencing analysis of *virp1* and *VIRP1*i *N. benthamiana* lines. **a**. Mapping of *VIRP1-*targeting sgRNA on *N. benthamiana* VIRP1 protein (red arrow). **b**. Editing analysis of homozygous *virp1-1* F_1_ mutant; the sgRNA target sequence and the PAM are indicated; +1: insertion of 1-nt; -17: deletion of 17-nt. **c**. Venn diagram showing the common differentially expressed genes (DEGs) between *virp1* mutants and *VIRP1*-suppressed plants. **d**. qPCR analysis of selected common DEGs; the respective fold change found in each RNA sequencing analysis is indicated in a box below each bar. **e**. GO term enrichment of *virp1 N. benthamiana* DEGs. **f**. GO term enrichment of *VIRP1i N. benthamiana DEGs. Shiny*GO 0.85 was used for GO enrichment analysis (FDR cutoff 0.05).

As BET proteins are considered transcriptional activators, we asked whether VIRP1 regulates gene expression in *N. benthamiana*. To adress this, we performed high-throughput RNA sequencing in *virp1* and *VIRP1*i plants to detect differentially expressed genes (DEGs) **(Tables S3-S5)**. We found 611 DEGs in *virp1* mutants and 1027 DEGs in *VIRP1i* plants, setting log_2_fold change ≥0.5 and padj <0.05 **(Fig. 2c)**. When we compared the two datasets, we detected a relatively low number of 74 common genes, which are presented in **Table 1**. Among the 74 common genes, several were related to chloroplast functions and photosynthesis, such as *Cytochrome b6, 30S ribosomal proteins S7/S14, Ycf6* and *Ribulose bisphosphate carboxylase large chain* **(Table 1)**. Many common genes were linked to stress and defense responses, including dormancy/auxin associated family proteins, involved in (a)biotic stress responses (Rae et al., 2014; Roy et al., 2020), and bifunctional nucleases, involved in defense response and ABA-derived callose deposition (M. K. You et al., 2010) **(Table 1)**. Furthermore, *ABA receptor PYL9* and *Magnesium chelatase subunit H*, both involved in ABA responses (Shen et al., 2006; X. Zhang et al., 2013), were found differentially expressed in both datasets **(Table 1)**. To evaluate our RNA sequencing analyses, we performed qPCR on selected common genes: *ABA receptor PYL9* **(Fig. 2d-i)**, *auxin-repressed 12*.*5 kDa protein* **(Fig. 2d-ii)**, *BTB/POZ and TAZ domain-containing protein 1* **(Fig. 2d-iii)**, *Deoxyuridine 5’-triphosphate nucleotidohydrolase* **(Fig. 2d-iv)**, *Protein ETHYLENE INSENSITIVE 3* **(Fig. 2d-v)** and *Remorin 4*.*2-like* **(Fig. 2d-vi)** showed similar expression patterns to those found in our sequencing analyses **(Table 1)**.

**Table 1.**
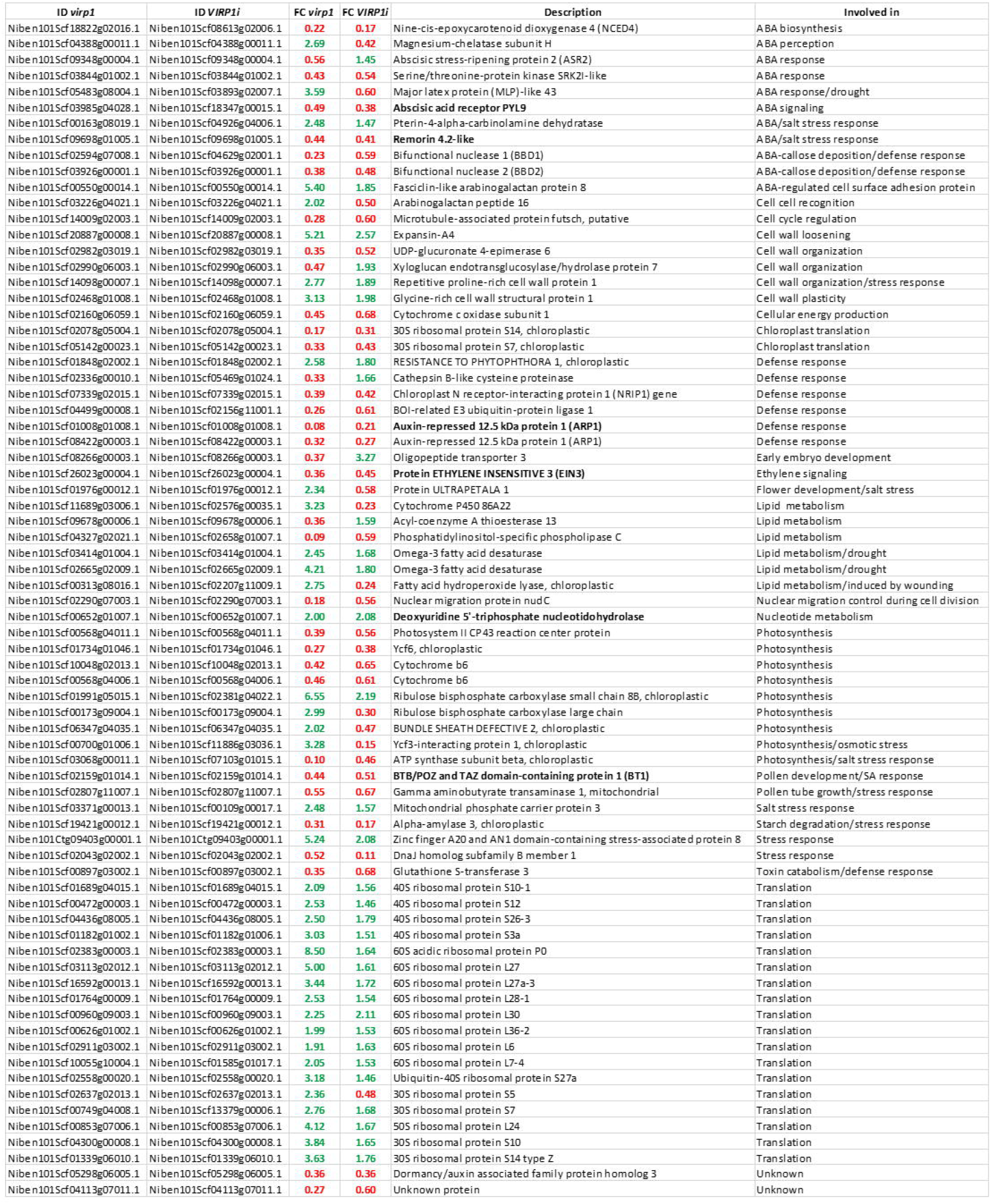
Common DEGs in *virp1* mutants and *VIRP1i* plants. Green color represents upregulation and red color represents downregulation. With bold are indicated the 6 genes selected for qPCR verification. FC: fold change. The descrption of each gene was adjusted using BLAST NCBI. For the role of each gene (last column) Uniprot database was used.

Following Gene Ontology (GO) analysis, we observed the following shared functional groups of DEGs between *virp1* mutants and *VIRP1i* plants: i) *Photosynthesis and energy-related genes*: Several DEGs are involved either in photosynthesis and chloroplast functions (Rubisco large and small units, PSII components, chloroplast ribosomal proteins) or in energy-related processes (sugar metabolism, lipid metabolism, ATP metabolism, electron transport chain, proton transport) **(Fig. 2e-f, Tables S4-S5)**. This highlights that VIRP1, partially or fully suppressed, impacts photosynthetic and energy-related pathways, pinpointing a role in metabolic regulation. ii) *Abiotic stress-related genes*: A significant number of genes in both *virp1* and *VIRP1i* plants are linked to abiotic stress response, including ABA-related genes, dormancy/auxin-associated proteins, remorins, heat-shock proteins and cytochrome P450 family enzymes **(Fig. 2e-f, Tables S4-S5)**. The following result is represented by “response to desiccation” in *virp1* mutants and by “response to stress” and “proline catabolic process” in *VIRP1i* plants **(Fig. 2e-f)**, with the latter closely related to osmotic/drought stress responses (Ghosh et al., 2022; Wei et al., 2022; Q. Zhang & Bartels, 2018). Some enriched gene groups were more prominent in *virp1* mutants. These groups include i) *chromatin-related proteins*, such as histone variants (H2A, H2B, H3.2, H3.3, H4), topoisomerases and chromatin-remodelers, ii) *transcription-related proteins*, such as transcription factors and RNA polymerase subunits and iii) *genes involved in translation*, such as 60S/40S ribosomal subunits and translation elongation factors **(Fig. 2e, Table S4)**. Finally, several gene groups were found uniquely enriched in *VIRP1i* plants, mainly linked to i) *photoperiodism/flowering regulation*, ii) *cytoskeleton organization*, iii) *cell wall metabolism* and iv) *signal transduction* and *transport* **(Fig. 2f, Table S5)**.

### Loss of VIRP1 affects flowering and ABA responses

Our transcriptome analyses revealed an altered gene expression profile in *virp1* mutants and *VIRP1i* plants, indicating alterations in stress response, photosynthesis and metabolic pathways. *virp1* mutants and *VIRP1i* plants did not present prominent phenotypic alterations compared to WT plants (data not shown); however, we noticed a subtle but statistically significant reduction in plant height and dry weight in both *VIRP1i* and *virp1* plants **(Fig. 3a-b)**, which aligns with chloroplast-and photosynthesis-related DEGs in both analyses **(Fig. 2e-f, Tables S4-S5)**. We conclude that *VIRP1* downregulation has a subtle but negative impact on plant growth. We further analyzed flowering time in our plant lines and discovered a slight delay in flowering of both *virp1* mutants and *VIRP1i* plants **(Fig. 3c)**. The above observation aligns with the downregulation of flowering-related genes in *VIRP1i* plants **(Fig. 2f, Table S5)**. This enrichment was not observed in the transcriptome analysis of *virp1* mutants; however, a strong downregulation (log_2_FC=-2.215) of *FLOWERING LOCUS T* was detected **(Table S4)**. As VIRP1 is most abundantly expressed in flowers compared to other tissues **(Fig. S2a)**, the above findings suggest a role of VIRP1 in flowering regulation. After observing the above subtle but consistent phenotypic alterations, we asked wether they could be accompanied by underlying defects in leaf epidermal development. Scanning electron microscopy (SEM) was therefore used to examine the leaf surface in detail. Both *virp1* mutants and *VIRP1i* plants exhibited a reduced stomata density with no changes in stomata morphology **(Fig. 3d-e, Fig. S2b)**.

**Fig. 3.**
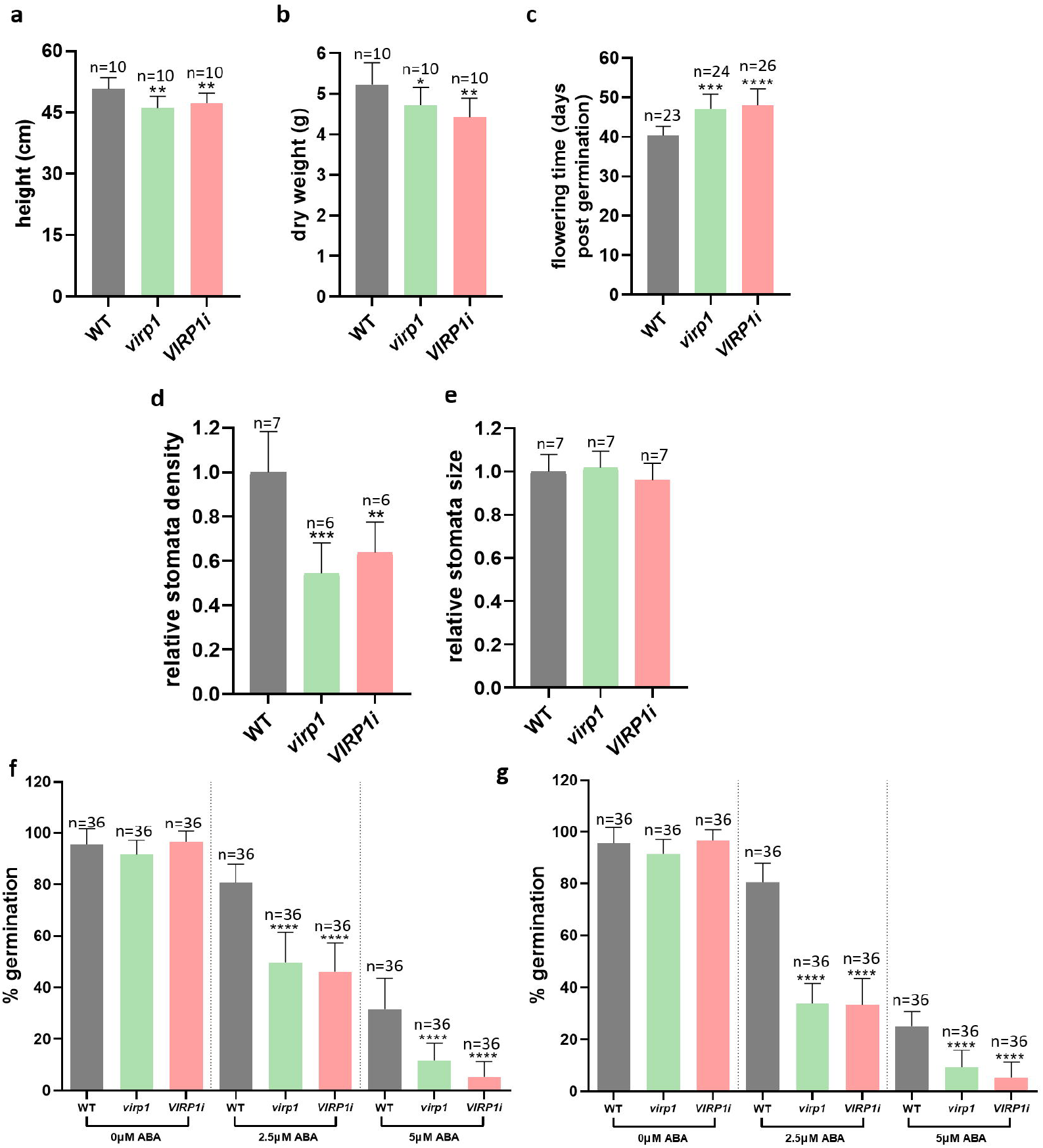
Phenotypic analysis of *virp1* and *VIRP1i N. benthamiana* lines. **a-c**. Analysis of plant height (a), dry weight (b) and flowering time (c) in WT, *virp1 and VIRP1-s*uppressed plants. **d**,**e**. Measurements of stomata density (d) and stomata size (e) in WT, *virp1* and *VIRP1*-suppressed plants. **f**,**g**. Analysis of *virp1* and *VIRP1*-suppressed plants germination under ABA treatment 2 weeks (f) and 3 weeks (g) after incubation with 0, 2.5 and 5μM ABA. Statistical analysis was performed in all cases with unpaired Student’s t-test with *p<0.05, **p<0.01 and ***p<0.001.

RNA sequencing analysis revealed a link of VIRP1 to abiotic stress responses, with several common DEGs related to ABA signaling **(Table 1)**. Based on this finding, we further aimed to investigate VIRP1 involvement in ABA responses. *VIRP1* expression was not affected by ABA in *N. benthamiana* plants **(Fig. S2c)**. However, when we placed *virp1* and *VIRP1i* seeds to ABA-containing medium, we observed a significantly increased sensitivity of both *virp1* mutants and *VIRP1i* plants to ABA-mediated inhibition of germination compared to WT plants **(Fig. 3f-g, Fig S2d)**. In support of this observation, *VIRP1*-overexpressing (*VIRP1* OE) *Col-0 A. thaliana* seeds showed resistance to ABA-mediated inhibition of germination compared to WT plants **(Fig. S2e-g)**. The above results suggest that VIRP1 acts as a negative regulator of ABA responses.

### VIRP1 is required for early PSTVd establishment

Our previous study demonstrated a crucial role of VIRP1 in PSTVd infectivity, as *VIRP1i N. benthamiana* plants were found resistant to PSTVd mechanical inoculation 4 weeks post infection (wpi) (Kalantidis et al., 2007). To further analyze how viroid infectivity is affected by VIRP1, we compared PSTVd levels in *virp1* mutants and *VIRP1*i plants using either mechanical inoculation or agroinfiltration. As shown in **Fig. 4a**, both *virp1* and *VIRP1*i plants are resistant to PSTVd mechanical inoculation at 4wpi. However, infection with PSTVd through agroinfiltration revealed considerable viroid titers in both *virp1* and *VIRP1*i plants at 3wpi, approximately 50% compared to WT plants **(Fig. 4b-c)**. To investigate the above in more detail, we assessed a time-course of PSTVd infectivity through mechanical infection and agroinfiltration. Our results showed that, through mechanical inoculation, low levels of PSTVd appeared at 5wpi in both *virp1* and *VIRP1*i plants, and remained low compared to WT up to 7wpi **(Fig. 4d)**. Using agroinfiltration, PSTVd levels in *virp1* and *VIRP1*i plants appeared lower than WT up to 3wpi; however, at later time-points they approached the viroid levels of WT plants **(Fig. 4e)**. The above results indicate that VIRP1 is mainly critical for early viroid establishment and that, once PSTVd is nuclear and systemic, VIRP1 is dispensable.

**Fig. 4.**
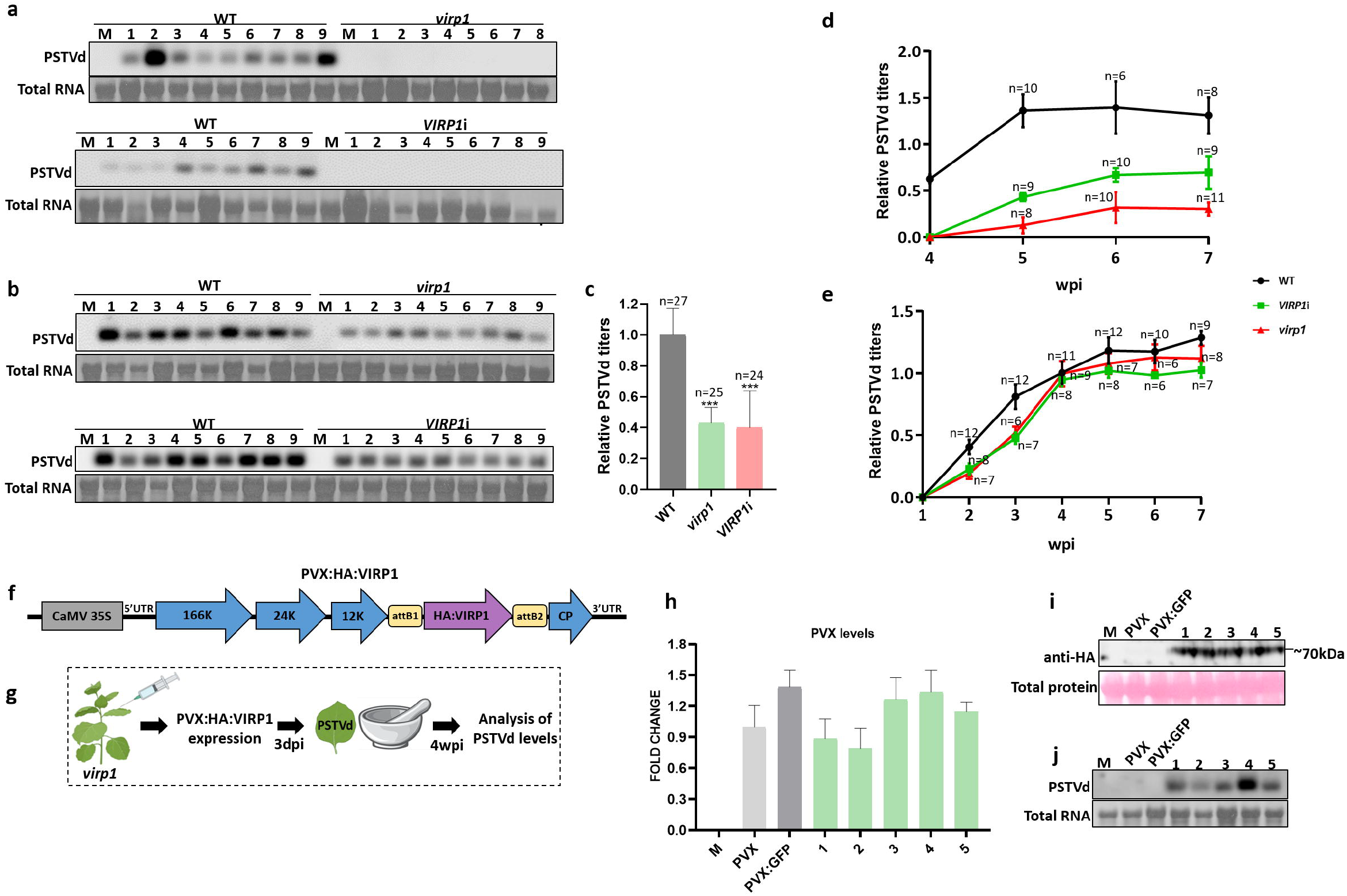
Analysis of PSTVd infectivity and complementation of *virp1* mutants. **a**. PSTVd levels in *virp1* mutants and *VIRP1*-suppressed plants 4 weeks after mechanical inoculation. **b**,**c**. PSTVd levels in *virp1* mutants and *VIRP1*-suppressed plants 3 weeks after agroinfiltration. **d**. Time-course of PSTVd levels after mechanical inoculation of *virp1* mutants and *VIRP1*-suppressed plants. **e**. Time-course of PSTVd levels after agroinoculation of *virp1* mutants and *VIRP1*-suppressed plants. **f**. PVX-GW construct used for VIRP1 complementation. **g**. Method followed for VIRP1 complementation: *virp1* mutants are inoculated with PVX-vector expressing HA-tagged VIRP1; 3 days later, inoculated plants are mechanically infected with PSTVd; PSTVd levels are monitored 4 weeks later. **h**. Analysis of PVX levels in selected plants with qPCR. **i**. VIRP1 protein levels in selected plants after PVX:HA:VIRP1 expression. **j**. PSTVd levels in selected plants 4 weeks after mechanical inoculation. PVX: *virp1* mutant inoculated with PVX empty vector; PVX:GFP: *virp1* mutant inoculated with the PVX vector overexpressing GFP; 1-5: selected *virp1* plants inoculated with PVX:HA:VIRP1. Statistical analysis was performed with unpaired Student’s t-test with *p<0.05, **p<0.01 and ***p<0.001; M: mock.

### Virus-mediated *VIRP1* expression restores PSTVd infectivity in *virp1 N. benthamiana* mutants

In order to understand how VIRP1 affects PSTVd infectivity, we aimed to establish a complementation system that would enable us to test the effects of VIRP1 mutation/truncation on viroid titers. To achieve this, we implemented a Gateway-compatible PVX-based erexpression vector (Lacorte et al., 2010), in which we cloned an HA-tagged tomato *VIRP1* **(Fig. 4f)**. We briefly describe the steps followed for complementation in **Fig. 4g**: *virp1* mutants are inoculated with PVX:HA:VIRP1 through agroinfiltration; 3 days later, plants are mechanically infected with PSTVd and viroid titers are analyzed 4 wpi. We employed mechanical inoculation, as *virp1* mutants are completely resistant to PSTVd up to 4wpi using the above method **(Fig. 4d)**. As controls, we used mock plants, plants infected with PVX empty vector, as well as plants infected with PVX:GFP **(Fig. 4h-j)**. First, PVX infection was estimated using RT-qPCR. We found comparable PVX levels in PVX-, PVX:GFP- and PVX:HA:VIRP1-inoculated samples **(Fig. 4h)**. Furthermore, we observed comparable HA:VIRP1 protein levels in PVX:HA:VIRP1-inoculated samples through Western blot analysis **(Fig. 4i)**. When we analyzed PSTVd titers in *virp1* mutants 4 weeks after PVX:HA:VIRP1 inoculation, we detected robust viroid levels **(Fig. 4j)**. Using the above method, we were able to restore PSTVd infectivity in *virp1* mutants, by successfully complementing VIRP1 function **(Fig. 4j)**.

### VIRP1 forms phase-separated nuclear condensates, which are affected by PSTVd RNA

To examine how different VIRP1 domains may play a role in PSTVd infectivity using the complementation system described above, we first aimed to study in detail WT VIRP1 localization. It is already known that VIRP1 localizes in the nucleus of healthy and PSTVd-infected cells (Kalantidis et al., 2007). In order to investigate VIRP1 localization in more detail, we fused tomato *VIRP1* to either a N-terminal YFP or a C-terminal GFP. Observation of transformed nuclei using confocal and fluorescent microscopy revealed that, in both fusions, VIRP1 forms distinct droplet-like puncta inside the cell nucleus. These puncta vary in number and size **(Fig. 5a)** and appear not to overlap with the nucleolus, as shown by co-expression of YFP:VIRP1 with FIBRILLARIN:RFP **(Fig. S3a)**. Since PSTVd RNA binds VIRP1, we asked whether it can affect VIRP1 localization. When we observed YFP:VIRP1 in PSTVd-infected leaves (3 wpi), we found that the nuclear puncta were larger, on average ∼2.5 fold compared to healthy plants **(Fig. 5a, Fig. S3b-c)**. Protein levels were not affected however, as YFP:VIRP1 levels were found similar in healthy and PSTVd-infected plants **(Fig. 5b)**.

**Fig. 5.**
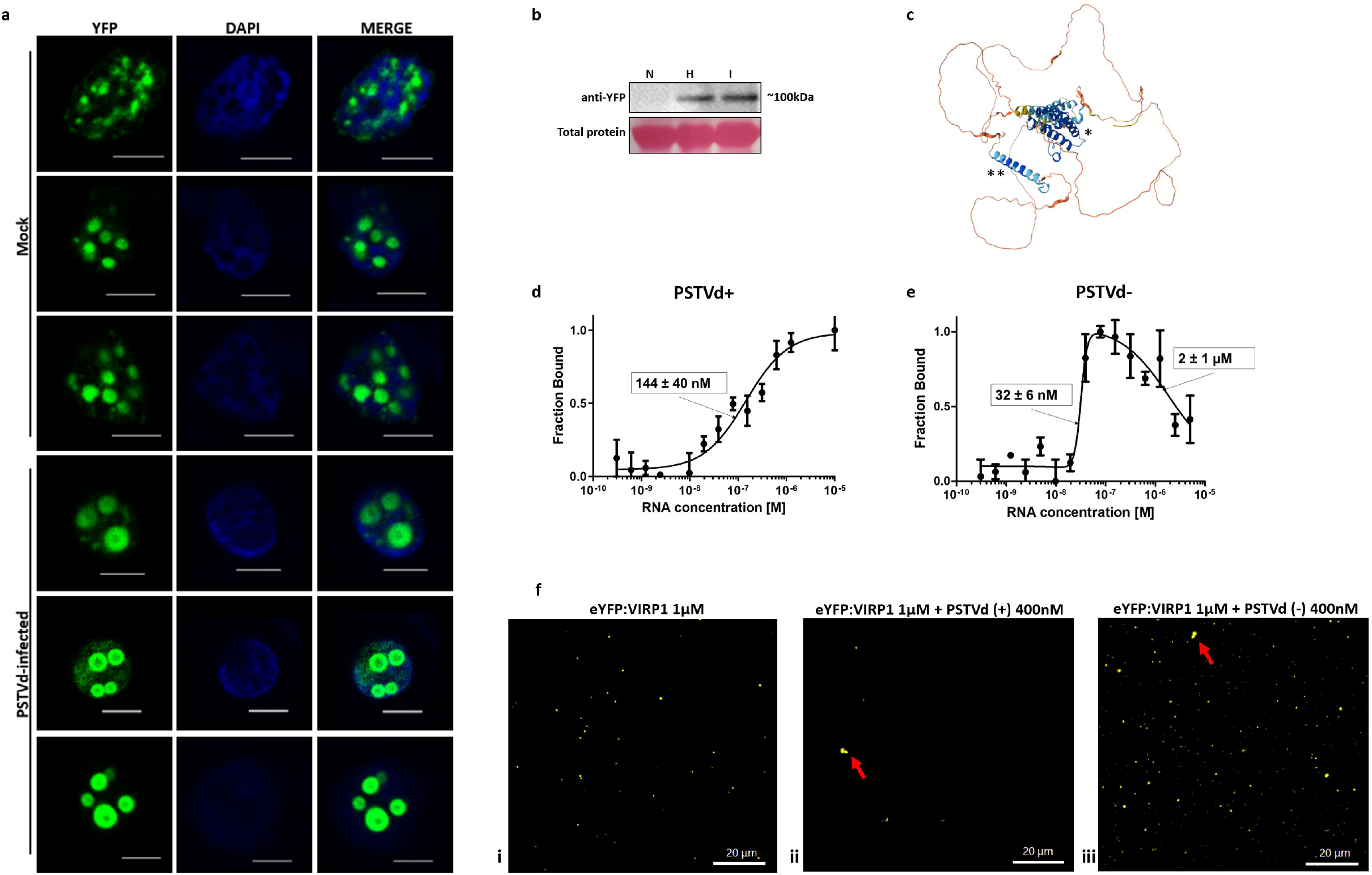
VIRP1 forms phase-separated nuclear condensates, which are affected by PSTVd RNA. **a**. Confocal microscopy of YFP:VIRP1 in healthy and PSTVd-infected *N. benthamiana* leaves (3wpi); scale bar=5μm. **b**. Protein levels of YFP:VIRP1 in healthy (H) and PSTVd-infected (I) leaves; N: negative control. **c**. Prediction of VIRP1 3D structure in Alphafold; the bromodomain (*) and NET domain (**) are indicated. **d**,**e**. Microscale thermophoresis showing binding of PSTVd (+) RNA (d) and PSTVd (-) RNA (e) to VIRP1. **f**. *In vitro* codensation assay of eYFP-tagged VIRP1 in the presence of PSTVd RNA; in panels ii and iii (+) and (-) PSTVd RNA was added respectively. Larger VIRP1 condensates in ii and iii are indicated with red arrows.

Further analyses suggested that the observed VIRP1 nuclear puncta resembled phase-separated condensates. Alphafold revealed that VIRP1 is a highly disordered protein, with the largest part of the predicted 3D structure being unstructured, setting aside the highly structured bromodomain and NET domain **(Fig. 5c)**. As disordered domains are often linked to liquid-liquid phase separation (LLPS) (Elbaum-Garfinkle et al., 2015; Molliex et al., 2015; Nott et al., 2015), we next used the FuzDrop prediction software to explore the probability of VIRP1 promoting LLPS. Indeed, VIRP1 is predicted to promote LLPS with a high probability of 0.9995 **(Fig. S3d)**. Furthermore, disordered domains are indicated as droplet-promoting regions across the VIRP1 sequence (1-92aa, 122-193aa, 290-418aa, 481-602aa) **(Fig. S3d)**. To test if indeed VIRP1 is a phase-separated protein, we purified His-tagged and eYFP-tagged VIRP1 **(Fig. S3e)** and performed *in vitro* assays. LLPS could be induced in 10% PEG 4000 and low salt concentration (<150 mM) at neutral pH. Phase-separated condensates began to appear at VIRP1 concentrations starting from 1 μM, whereas free eYFP remained disperse **(Fig. S3f)**. To verify the binding of *in vitro*-purified VIRP1 to PSTVd, (+) and (-) PSTVd RNA was used for *in vitro* Microscale thermophoresis (MST) assays. Indeed, PSTVd RNA of both polarities showed a strong affinity for eYFP-VIRP1 **(Fig. 5d-e)**. In contrast, Cy5-labeled miR164, used as a negative control, did not show any binding **(Fig. S3g)**. Since binding was confirmed, PSTVd RNA was added to the assays to examine VIRP1 condensate formation. When 400 nM of PSTVd RNA of (+) or (-) polarity was added to eYFP:VIRP1, larger condensates were observed sporadically **(Fig. 5f)**. When we added Cy5-labeled miR164 in a 1:1 ratio, we observed a recruitment of RNA into eYFP-VIRP1 condensates, however there was no effect on condensate appearance **(Fig. S3h)**. We conclude that VIRP1 forms phase-separated condensates, the size of which increases in the presence of PSTVd RNA.

### VIRP1 nuclear localization and bromodomain integrity are substantial for viroid infection, whereas the CTD is dispensable

After demonstrating VIRP1 phase separation in the nucleus, we aimed to investigate how modifications across VIRP1 sequence may disrupt this characteristic subnuclear behavior, which could also indicate functional changes that may affect PSTVd infectivity. We selected to modify a number of VIRP1 sites that may potentially be crucial for VIRP1 localization and function. As VIRP1 has been proposed to facilitate viroid nuclear entry, we first asked to what extent VIRP1 nuclear localization is important for PSTVd replication. Using prediction tools, we identified 2 putative nuclear localization signals (NLSs) in VIRP1 amino acid sequence with high probability **(Fig. 6a)**. However, confocal microscopy revealed exclusively nuclear signal in the respective NLS single and double mutants, with no significant changes in condensate formation **(Fig. 6b)**. We speculated that VIRP1 either contains multiple or atypical NLSs, or its nuclear localization is mediated by other mechanisms. To produce a cytoplasmic version of the protein, we N-terminally fused VIRP1 to a nuclear exit signal (NES) **(Fig. S4a)**. Confocal microscopy revealed cytoplasmic signal of VIRP1, however an important fraction still remained nuclear **(Fig. 6c)**. Protein levels were found similar in YFP:VIRP1 and YFP:NES:VIRP1 **(Fig. 6d)**. Using the previously described PVX-based complementation method **(Fig. 4f-j)**, we expressed HA:NES:VIRP1 **(Fig. 6e)** in *virp1* mutants. After verifying PVX and protein expression levels **(Fig. 6f-g)**, we observed a statistically significant reduction in PSTVd titers **(Fig. 6h-i)**. We therefore conclude that reduction of VIRP1 nuclear accumulation negatively affects viroid infectivity.

**Fig. 6.**
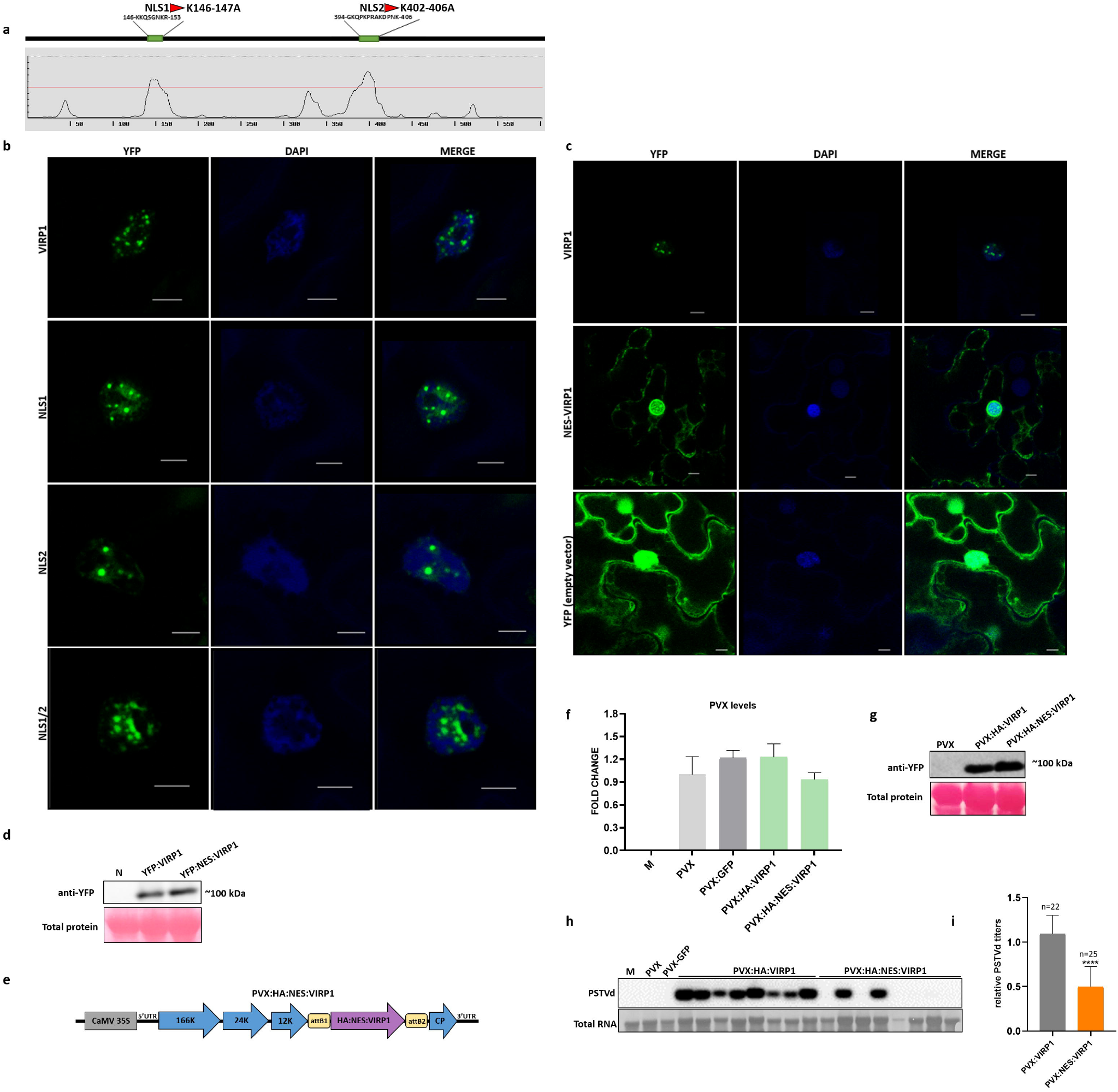
Effect of NES:VIRP1 fusion on VIRP1 localization and PSTVd infectivity. **a**. Prediction of putative NLSs on VIRP1 sequence (cutoff=0.5). The sequence of each putative NLS is indicated, as well as the mutations generated (NLS1: K146-147A; NLS2: K402-406A). **b**. Confocal microscopy of YFP-tagged NLS single and double mutants; scale bar=5μm. **c**. Confocal microscopy of YFP:NES:VIRP1; scale bar=5μm. **d**. Protein levels of YFP-tagged VIRP1 and NES:VIRP1. **e**. PVX construct carrying HA:NES:VIRP1, used for the complementation experiment. **f**. PVX levels of infected plants used for complementation experiment (n=3). **g**. Protein levels of PVX:HA:VIRP1 and PVX:HA:VIRP1 from complementation experiment. **h**. Indicative northern blot of complementation experiment showing PSTVd levels after PVX:HA:VIRP1 and PVX:HA:VIRP1 overexpression in *virp1* mutants. **i**. Quantification of PSTVd levels from 3 independent complementation experiments. Statistical analysis was performed with unpaired Student’s t-test with *p<0.05, **p<0.01 and ***p<0.001; M: mock; N: negative control.

As VIRP1 contains a large disordered CTD (residues 481-602), predicted to induce droplet formation **(Fig. S3d)**, we asked whether its deletion may affect VIRP1 subnuclear localization, by disrupting the normal appearance of the condensates. We designed a number of truncations, leading to the following VIRP1 fragments: 1-550aa, 1-500aa, 1-482aa, 1-458aa **(Fig. 7a)**. We first asked whether these fragments, when expressed in *N. benthamiana* leaves, produced similar protein levels as the intact VIRP1 protein. We detected all truncated versions of VIRP1 at higher protein levels compared to the intact protein, which was detected at relatively low levels **(Fig. 7b-c)**. When we N-terminally fused the above fragments to YFP, we did not observe any alterations in the localization of truncations 1-550aa and 1-500aa compared to the intact protein **(Fig. 7d)**. However, in the fragments 1-482aa and 1-458aa we observed condensates smaller in size and more dispersed compared to the intact protein **(Fig. 7d)**. In all cases, localization was exclusively nuclear **(Fig. 7d)**. We concluded that deletion of the disordered CTD of VIRP1 can affect its subnuclear localization, leading to a more dispersed distribution of the protein in the nucleus compared to the intact protein. Part of the CTD may also be responsible for the protein levels, possibly affecting protein stability or turnover **(Fig. 7b-c)**.

**Fig. 7.**
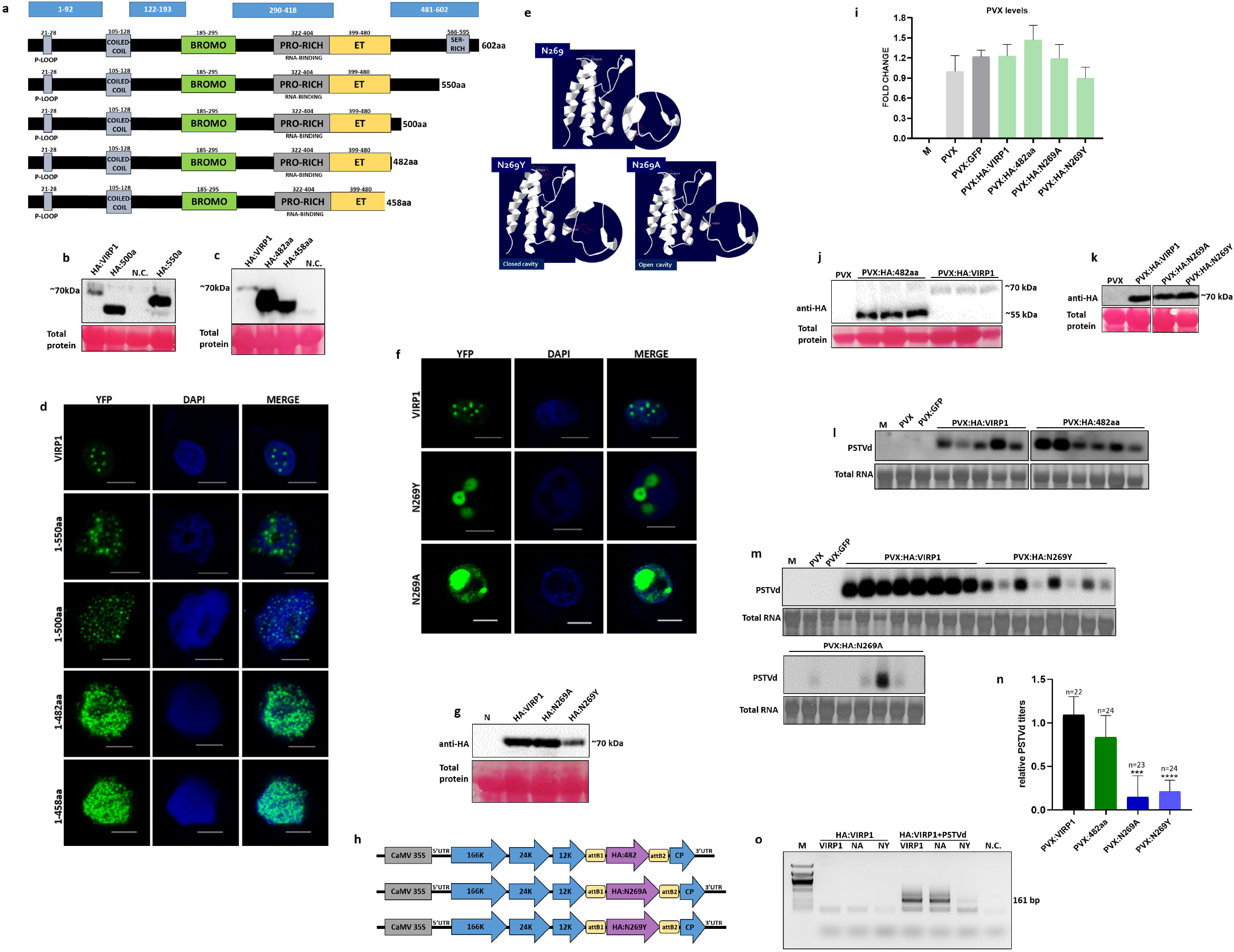
Effect of VIRP1 C-terminal truncation and bromodomain mutations on PSTVd infectivity. **a**. C-terminally truncated versions of VIRP1; in blue boxes above, the disordered, droplet-promoting regions are depicted (according to FuzDrop). **b**. Confocal microscopy of YFP:VIRP1 and C-terminally truncated versions; scale bar=5μm. **c**,**d**. Protein levels of HA-tagged VIRP1 and C-terminal truncations. **e**. *In silico* prediction of the effect of N269 mutation to either Y or A on the site of acetyl-lysine recognition in VIRP1 bromodomain (Swiss-PdbViewer 4.1). **f**. Confocal microscopy of YFP-fused VIRP1 and bromodomain mutants; scale bar=5μm. **g**. Protein levels of HA-tagged VIRP1 and bromodomain mutants. **h**. Constructs of PVX:HA:482aa, PVX:HA:N269A and PVX:HA:N269Y. **i**. PVX levels of infected plants used for complementation experiment (n=3). **j-k**. Protein levels of HA-tagged VIRP1, truncation 1-482aa and bromodomain mutants from complementation experiment. **l**,**m**. Indicative northern blots from complementation experiments showing PSTVd levels after overexpression of 1-482aa truncation (L) and bromodomain mutants (M) in *virp1* plants. **n**. Quantification of PSTVd levels from 3 independent complementation experiments. Statistical analysis was performed with unpaired Student’s t-test with *p<0.05, **p<0.01 and ***p<0.001; M: mock; N: negative control. **o**. PSTVd-binding of VIRP1 bromodomain mutants; after HA:VIRP1 immunoprecipitation, RNA was extracted and cDNA was synthesized; PCR using PSTVd-specific primers verifies PSTVd binding on VIRP1 and bromodomain mutants (NA, NY); M: marker (lamda/PstI).

To examine whether the VIRP1 bromodomain may be important for the protein’s subnuclear localization, we chose to mutate a conserved aminoacid of the bromodomain, the asparagine residue in position 269 (N269) **(Fig. S4b)**, which is crucial for acetyl-lysine recognition in human BET proteins (Filippakopoulos et al., 2012; Floyd et al., 2013; Umehara et al., 2010). We designed 2 different mutations, N269Y and N269A, for which *in silico* analysis predicted alterations in the site of acetyl-lysine recognition, leading to a rather “closed” or “open” cavity that may affect recognition **(Fig. 7e)**. Confocal microscopy revealed a change in subnuclear localization of N269Y and N269A, showing larger and fewer condensates compared to WT VIRP1 **(Fig. 7f)**. Protein levels of HA-tagged bromodomain mutants appeared similar to WT VIRP1 **(Fig. 7g)**. We concluded that mutation of the conserved N269 residue of the VIRP1 bromodomain disrupts VIRP1 subnuclear localization, leading to altered condensate formation, without affecting protein levels.

To test whether changes in VIRP1 subnuclear behavior may have an effect on viroid infectivity, we employed the previously described method **(Fig. 4f-j)** to complement with VIRP1 versions that exhibited significant differences in subnuclear behavior **(Fig. 7h)**. More specifically, we chose the fragment 1-482aa, which is the longest fragment displaying dispersed nuclear condensates **(Fig. 7d)**, as well as the two bromodomain mutants, N269A and N269Y, displaying fewer and larger nuclear condensates **(Fig. 7g)**. After verifying PVX infection and protein overexpression **(Fig. 7i-k)**, we assessed viroid levels in inoculated plants. HA:1-482 complementation revealed no statistically significant difference in viroid levels compared to intact VIRP1 **(Fig. 7l,n)**. However, complementation with HA:N269A or HA:N269Y, resulted to a significant reduction in viroid levels compared to WT VIRP1 expression **(Fig. 7m-n)**. Immunoprecipitation of HA:VIRP1 with PSTVd RNA showed that PSTVd RNA binding is not abolished in the bromodomain mutants **(Fig. 7o)**. Our results indicate that an intact VIRP1 bromodomain is crucial for PSTVd accumulation, whereas VIRP1 CTD is dispensable.

## DISCUSSION

Through this study, we show that VIRP1 belongs to the BET protein family, being more closely related to GTE7 and GTE2 of *A. thaliana* **(Fig. 1c)**. Most BET members share a bromodomain and a N-ET domain, as well as conserved domains and regions, including a coiled-coil and a serine-rich C-terminal domain **(Fig. 1a-b)**. Among tomato and *N. benthamiana* BET members, a proline-rich domain is present exclusively in VIRP1 **(Fig. 1a-b)**. This domain, residing between the bromodomain and the N-ET domain (aa 322-404) (Martínez De Alba et al., 2003), overlaps with the PSTVd RNA-binding site (Martínez De Alba et al., 2003) and is absent from all other BET members in tomato and *N. benthamiana*; it is however present in GTE2, GTE7 and GTE5 BETs of *A. thaliana* (Bardani et al., 2023), with GTE7 being a PSTVd-binding protein (Ma et al., 2022). We propose that this low-complexity proline-rich domain is a specialized, adaptive interface that mediates viroid interaction, rather than a core BET functional component. To that end, there is possibly no redundancy among BET members regarding the positive role of VIRP1 in PSTVd replication.

Through transcriptome analysis, we show that VIRP1 affects the expression of genes linked to abiotic stress responses – mainly ABA signaling and drought/osmotic stress – and chloroplast metabolism **(Table 1, Tables S4-S5, Fig. 2e-f)**. This leads to subtle reductions in height and dry weight **(Fig. 3a-b)**, a detectable delay in flowering time **(Fig. 3c)**, reduced stomata density **(Fig. 3 d-e, Fig. S2b)** and increased sensitivity to ABA-mediated inhibition of germination, positioning VIRP1 as a negative regulator of ABA responses **(Fig. 3f-g, Fig. S2d-g)**. In support of these findings, several BET members in *A. thaliana* (GTE1, GTE9-11), as well as non-BET bromodomain-containing proteins (BRD1, BRD2, BRD13) are involved in ABA signaling (Bardani et al., 2023). Furthermore, a number of bromodomain-containing proteins are involved in the regulation of flowering and circadian rhythm, such as BRAHMA (BRM) (Bardani et al., 2023). Since these functions are intertwined, with ABA response regulating flowering (Martignago et al., 2020), these findings suggest that VIRP1, through regulation of gene expression, controls ABA signaling, fine-tuning photosynthesis, energy metabolism, and circadian rhythm. Additionally, the effect both on ABA-mediated inhibition of germination and stomatal density suggests that VIRP1 may be involved in a broader ABA-dependent regulatory network that includes stomata development.

By comparing our RNA sequencing analyses, we found a number of enriched gene groups unique for *virp1* mutants, mainly related to chromatin, transcription and translation **(Fig. 2e, Table S4)**. An interpretation of the above could be that complete loss of VIRP1 function drives fundamental gene expression and chromatin remodeling changes, possibly reflecting VIRP1’s direct role in gene regulation and chromatin organization. Some gene groups were found exclusively enriched in *VIRP1i* plants, mainly linked to photoperiodism, metabolism and signal transduction **(Fig. 2f, Table S5)**. Overall, transcriptomic differences observed between *virp1* and *VIRP1i* lines could be a result of *VIRP1* knockout versus knockdown. Alternatively, they could be the direct or indirect effects of RNAi-triggered phenomena or position effects due to transgene insertion (Neumeier & Meister, 2021; Senthil-Kumar & Mysore, 2011).

By performing time-course experiments, we showed that PSTVd can systemically infect VIRP1-defective *N. benthamiana* plants, although with lower efficiency **(Fig. 4d-e)**. Mechanical inoculation introduces PSTVd into a limited number of epidermal and mesophyll cells. Furthermore, PSTVd RNA is not protected from cytoplasmic RNA decay and has to achieve nuclear entry for replication. This leads to a strong and persistent restriction of PSTVd accumulation. The delayed and low-level infection in this case reflects possible escape events where PSTVd exploits alternative, low-efficiency pathways to achieve replication. On the contrary, agroinfiltration leads to completely different infection dynamics; continuous PSTVd transcription, simultaneous infection of multiple cells, and direct introduction into the nucleus of infected cells bypasses the early establishment bottlenecks observed during mechanical inoculation experiments, leading to partial early PSTVd restriction, followed by full recovery at later time-points **(Fig. 4e)**. Together, the above results suggest that VIRP1 enhances early infection establishment, possibly by facilitating nuclear targeting and/or trafficking in early infected cells. Once PSTVd has entered the initial cell nuclei and has gone systemic, VIRP1 is no longer necessary.

Additionally, the similar results obtained in *VIRP1i* and *virp1* lines indicate that VIRP1 involvement in PSTVd replication is dosage-independent beyond a certain threshold, and that any residual VIRP1 levels in *VIRP1i* plants are below functional significance. Using a PVX-based expression vector, we complemented tomato *VIRP1* into *virp1* mutants, and restoried PSTVd infectivity **(Fig. 4f-j)**. This is the first time, to our knowledge, that VIRP1 complementation has been achieved. Furthermore, using this method, we have established a convenient and time-saving complementation system that can be used to investigate in detail the role of different VIRP1 regions in PSTVd accumulation.

Using confocal microscopy, we observed that VIRP1 forms macromolecular nuclear assemblies **(Fig. 5a)**. *In vitro* condensation assays suggested that these assemblies are the result of LLPS **(Fig. 5f, Fig. S3f)**. In the presence of PSTVd RNA, the condensates appeared significantly larger in size when VIRP1 was transiently expressed in PSTVd-infected plants **(Fig. 5a, Fig. S3b-c)**. Supporting this observation, *in vitro* condensation assay in the presence of PSTVd RNA led to the formation of sporadic aggregates **(Fig. 5f)**. This could be a result of the direct binding of PSTVd RNA to VIRP1 **(Fig. 5d-e)**, which may have further functional implications, something that is yet to be discovered. It is to note that, as our LLPS designation was based on PEG-crowding *in vitro*, formal biophysical characterization remains for future work. Deletion of the disordered VIRP1 CTD (truncations 1-482aa and 1-458aa) led to smaller and more abundant condensates dispersed throughout the nucleus **(Fig. 7d)**. In all truncations, protein levels were found to be higher compared to intact VIRP1, suggesting that the disordered CTD might affect protein stability or turnover **(Fig. 7b-c)**. When we mutated the conserved asparagine N269 bromodomain residue **(Fig. 7e)**, we observed larger and fewer condensates inside the cell nuclei **(Fig. 7f)**. This observation aligns well with studies in human BRD4, where bromodomain mutations in conserved aminoacids were shown to abolish acetyl-lysine recognition and chromatin association (Jung et al., 2014), while disruption of chromatin binding resulted in altered condensate formation (Strom et al., 2024). On the contrary, mutations of positively charged lysine residues in putative NLSs across VIRP1 sequence did not lead to any major changes in VIRP1 localization **(Fig. 6a-b)**. We proposed that either VIRP1 is directed to the nucleus by multiple NLSs, or that its nuclear entry is mediated by atypical NLSs or other mechanisms. By N-terminally fusing a NES to VIRP1 **(Fig. S4a)**, we generated a partially nuclear-partially cytoplasmic VIRP1 version **(Fig. 6c)**.

Complementation analysis revealed, that reducing nuclear accumulation of VIRP1 significantly compromises PSTVd infection **(Fig. 6h-i)**, indicating that a predominantly nuclear VIRP1 is required, as suggested by previous studies (Ma et al., 2022; Seo et al., 2021), while the C-terminal disordered domain is dispensable **(Fig. 7l,n)**. The most compelling mechanistic result, however, is the functional importance of the bromodomain. Mutations in the conserved bromodomain N269 residue strongly impaired viroid infection **(Fig. 7m-n)**, without abolishing PSTVd binding **(Fig. 7o)**. The fact that the deletion of the disordered CTD affects condensate formation but not viroid accumulation further supports that the pro-viroid activity of VIRP1 may be bromodomain-dependent and not condensate formation-dependent. Here, it is worth noting that while complementation relies on viral overexpression and thus may not fully reflect endogenous dosage, VIRP1 variants were expressed at comparable levels, allowing direct comparison of domain/mutant effects on PSTVd accumulation. Recently, GTE7 and GTE2, the closest Arabidopsis VIRP1 homologs, were shown to bind acetylated histones, whereas this binding was demonstrated to facilitate the association of the closely related GTE4 with chromatin (Qian et al., 2024). We therefore speculate that the positive role of VIRP1 in viroid replication is not restricted to viroid nuclear import, but is also linked to VIRP1’s association with chromatin via its bromodomain. A plausible hypothesis is that VIRP1 association with open chromatin brings the viroid in proximity to Pol II, constituting VIRP1 a proper vehicle for viroid trafficking to specific subnuclear sites. In support of this speculation, PSTVd localization to histones was shown in an earlier study, however the importance of this finding was not clear (Wolff et al., 1985). Alternatively, VIRP1 may act as a linker that attaches PSTVd to mitotic chromosomes during cell division; this could facilitate nuclear retention of PSTVd and stabilization in dividing cells, as has been demonstrated for BRD4, through binding to the E2 protein of Papillomaviruses (Li et al., 2014; McBride et al., 2021). As nuclear-replicating viroids do not encode any proteins, they might have evolved to directly bind chromatin-attaching proteins for mitotic chromosome retaining and partitioning to daughter nuclei. Such a mechanism would be particularly critical during early stages of infection, when viroid copies are limited and loss during cell division would have a strong negative impact on the establishment of infection. Therefore, we propose that VIRP1 is required for early viroid establishment rather than for sustained replication and that this role may be mechanistically linked to VIRP1’s bromodomain-mediated chromatin association. Several aspects that would favor the above hypotheses remain open. Direct evidence that VIRP1 bromodomain mutants lose chromatin association is not yet provided, and the precise relation of VIRP1 condensates to known nuclear bodies or to viroid replication sites remains to be established. These limitations do not weaken the central conclusions, but they point to the next mechanistic questions raised by the study. Overall, the data support a model in which VIRP1 is a stress-related BET protein that has been co-opted by PSTVd to facilitate early infection. Its unique proline-rich viroid-binding region, combined with a bromodomain-dependent nuclear function and condensate-forming capacity, places VIRP1 at the intersection of RNA recognition, chromatin-linked organisation and host susceptibility to viroid infection.

## Supporting information

Sopplemental fig 2

## Acknowledgements

The authors would like to thank Myrsini Froudaki for the generation of *VIRP1*-overexpressing *A. thaliana* lines and Paraskevi Kallemi for the FIBRILLARIN:RFP construct. We also thank Maro Iliopoulou, Villy Michalopoulou, Mireia Uranga and Konstantina Arkoulaki for technical assistance in the project, as well as Stephanos Papadakis for helping with SEM and Stelios Mavridis for assistance at the greenhouse.

We acknowledge funding to KJ from the European Research Council (ERC) under the European Union’s Horizon 2020 research and innovation programme (GA No. 810131) and from the DFG (project number 433194101, Research Unit 5116).

## Author contributions

BE, OS, KKo and KKr designed the research; BE, OS, AF, OA and KKo performed the research; AC performed the data analysis; KKr and KJ secured funding; BE wrote the manuscript; OS, KKo, KJ and KKr revised the manuscript.

## Figures and Tables legends

**Fig. S1. Screening steps for the generation of *virp1 N. benthamiana mutants*. a**. Method followed for the generation of *virp1* mutants: *Cas9*-expressing plants are inoculated with a PVX-vector carrying the desired sgRNA; infected tissue (2wpi) is isolated and used for plant regeneration through tissue culture; PVX-infected regenerated plants (F_0_) are screened for edits using ICE; PVX-free progeny of selected plants (F_1_) are analyzed using ICE. **b**. Editing analysis of PVX-infected regenerated plants (F_0_). **c**. Analysis of knockout mutations in PVX-free edited progeny (F_1_). Red arrows indicate selected plants; for editing analysis, the ICE online tool (Synthego) was used.

**Fig. S2. Comparing *virp1* mutants and *VIRP1i N. benthamiana* plants. a**. Expression levels of *VIRP1* in different tissues of N. *benthamiana*. L: leaves; F: flowers; S: shoot; R: root. **b**. Photos of leaf surface of WT, *virp1* and *VIRP1i* plants obtained using scanning electron microscopy (SEM); in the upper panels, white asterisks indicate positions of stomata; photos of x300 magnitude where used for the quantification of stomata density (upper panels); photos of x3000 magnitude where used for the quantification of stomata size (lower panels). **c**. Nb*VIRP1* expression levels after 10μM ABA treatment. **d**. Germination of WT, *virp1* and *VIRP1*-suppressed *N. benthamiana* plants in MS plates supplied with 0μM, 2.5μM and 5μM ABA 3 weeks after placement. **e**. Germination of *VIRP1*-overexpressing *A. thaliana* plants under ABA treatment 3 weeks after placement. **f**,**g**. Germination analysis of *Col-0* and *VIRP1*-overexpressing *A. thaliana* plants under ABA treatment 2 weeks (f) and 3 weeks (g) after placement under 0, 0.5 and 1μM ABA. Statistical analysis was performed with unpaired Student’s t-test with *p<0.05, **p<0.01 and ***p<0.001.

**Fig. S3. VIRP1 localization and phase separation. a**. Confocal microscopy of YFP:VIRP1 and FIBRILLARIN:RFP; scale bar=5μm. **b**. Fluorescent microscopy of YFP:VIRP1 (i) and VIRP1:GFP (ii) in healthy leaves and YFP:VIRP1 in PSTVd-infected leaves (iii); scale bar=5μm. **c**. Analysis of aggregate size in healthy and infected cells; fluorescent microscopy photos were used for the analysis. **d**. Prediction of VIRP1 probability of phase separation using FuzDrop. **e**. Purification of recombinantly expressed eYFP-tagged VIRP1 from *E*.*coli*. Final purity of VIRP1 was approximately 90% after evaluation on a 10 % SDS gel. Identity, modifications and degradations were checked via In-Gel trypsin digestion with a subsequent MALDi-TOF mass spectrometric analysis. Analyzed samples are marked with an asterisk. Fraction 17 showed highest purity and was concentrated and snap frozen until further used. Note that minor impurities are DnaK from *E*.*coli* and VIRP1 with an aberrant migration at approximately 125 kDa. **f**. *In vitro* condensation assay of His-and eYFP-tagged VIRP1. Purified proteins were set to a final concentration of 1 µM and PEG 4000 to a final concentration of 10 % was added. **g**. MST measurements of eYFP:VIRP1 towards miR164. **h**. Incorporation of unspecific RNA into eYFP:VIRP1 condensates. Cy5-labeled RNA oligomer miR164 was added to eYFP-tagged VIRP1 during in vitro codnesation assays in a 1:1 ratio. Colocalization of Cy5 and eYFP signal highlight a recruitment of RNAs into formed condesates.

**Fig. S4. NES and bromodomain sequence alignment. a**. NES sequence fused to VIRP1 for cytoplasmic localization. **b**. Alignment of partial bromodomain sequences from tomato and *N. benthamiana* VIRP1, GTE Arabidopsis homologs, as well as the second bromodomain of human BETs (BD2).

**Table S1. Primers used in this study**.

**Table S2. BET homologs in tomato (*S. lycopersicum*) and *N. benthamiana*. a**. Aminoacid similarity between SlVIRP1, the two paralogs of NbVIRP1 (a and b) and GTE7/GTE2. **b**. BET proteins in tomato. The chromosome on which each gene is located, protein length, molecular weight and isoelectric point are indicated, as well as the closest homologs in *N. benthamiana* and *A. thaliana* (based on aa sequence similarity).

**Table S3. Small RNA sequencing libraries**

**Table S4. Transcriptome analysis of *virp1 N. benthamiana* mutants**.

**Table S5. Transcriptome analysis of *VIRP1i N. benthamiana* plants**.

## References

Airoldi, C. A., Rovere, F. D., Falasca, G., Marino, G., Kooiker, M., Altamura, M. M., Citterio, S., & Kater, M. M. (2010). The Arabidopsis BET Bromodomain Factor GTE4 Is Involved in Maintenance of the Mitotic Cell Cycle during Plant Development. Plant Physiology, 152(3), 1320–1334. 10.1104/pp.109.150631

Anders, S., Pyl, P. T., & Huber, W. (2015). HTSeq—A Python framework to work with high-throughput sequencing data. Bioinformatics, 31(2), 166–169. 10.1093/bioinformatics/btu638

Andersen, C. L., Jensen, J. L., & Ørntoft, T. F. (2004). Normalization of Real-Time Quantitative Reverse Transcription-PCR Data: A Model-Based Variance Estimation Approach to Identify Genes Suited for Normalization, Applied to Bladder and Colon Cancer Data Sets. Cancer Research, 64(15), 5245–5250. 10.1158/0008-5472.CAN-04-0496

Andrews S. (n.d.). FastQC: a quality control tool for high throughput sequence data. [Computer software]. Retrieved https://www.bioinformatics.babraham.ac.uk/projects/fastqc/

Bardani, E., Kallemi, P., Tselika, M., Katsarou, K., & Kalantidis, K. (2023). Spotlight on Plant Bromodomain Proteins. Biology, 12(8), 1076. 10.3390/biology12081076

Bardani, E., Katsarou, K., Mitta, E., Andronis, C., Štefková, M., Wassenegger, M., & Kalantidis, K. (2025). Broadening the Nicotiana benthamiana research toolbox through the generation of dicer-like mutants using CRISPR/Cas9 approaches. Plant Science, 356, 112490. 10.1016/j.plantsci.2025.112490

Bombarely, A., Rosli, H. G., Vrebalov, J., Moffett, P., Mueller, L. A., & Martin, G. B. (2012). A Draft Genome Sequence of Nicotiana benthamiana to Enhance Molecular Plant-Microbe Biology Research. Molecular Plant-Microbe Interactions®, 25(12), 1523–1530. 10.1094/MPMI-06-12-0148-TA

Branch, A. D., & Robertson, H. D. (1984). A Replication Cycle for Viroids and Other Small Infectious RNA’s. Science, 223(4635), 450–455. 10.1126/science.6197756

Bruening, G. (1991). Current Review Replication of a Plant Virus Satellite RNA: Evidence Favors Transcription of Circular Templates of Both Polarities. Molecular Plant-Microbe Interactions, 4(3), 219. 10.1094/MPMI-4-219

Bushnell B. (n.d.). BBMap [Computer software]. Retrieved BBMap - Bushnell B. - sourceforge.net/projects/bbmap/

Chaturvedi, S., Kalantidis, K., & Rao, A. L. N. (2014). A Bromodomain-Containing Host Protein Mediates the Nuclear Importation of a Satellite RNA of Cucumber Mosaic Virus. Journal of Virology, 88(4), 1890–1896. 10.1128/JVI.03082-13

Chua, Y. L., Channelière, S., Mott, E., & Gray, J. C. (2005). The bromodomain protein GTE6 controls leaf development in Arabidopsis by histone acetylation at ASYMMETRIC LEAVES1. Genes & Development, 19(18), Article 18. 10.1101/gad.352005

Devaiah, B. N., Lewis, B. A., Cherman, N., Hewitt, M. C., Albrecht, B. K., Robey, P. G., Ozato, K., Sims, R. J., & Singer, D. S. (2012). BRD4 is an atypical kinase that phosphorylates Serine2 of the RNA Polymerase II carboxy-terminal domain. Proceedings of the National Academy of Sciences, 109(18), 6927–6932. 10.1073/pnas.1120422109

Dhalluin, C., Carlson, J. E., Zeng, L., He, C., Aggarwal, A. K., & Zhou, M.-M. (1999). Structure and ligand of a histone acetyltransferase bromodomain. 399.

Dissanayaka Mudiyanselage, S. D., Ma, J., Pechan, T., Pechanova, O., Liu, B., & Wang, Y. (2022). A remodeled RNA polymerase II complex catalyzing viroid RNA-templated transcription. PLOS Pathogens, 18(9), e1010850. 10.1371/journal.ppat.1010850

Dissanayaka Mudiyanselage, S. D., Qu, J., Tian, N., Jiang, J., & Wang, Y. (2018). Potato Spindle Tuber Viroid RNA-Templated Transcription: Factors and Regulation. Viruses, 10(9), 503. 10.3390/v10090503

Duque, P., & Chua, N.-H. (2003). IMB1, a bromodomain protein induced during seed imbibition, regulates ABA- and phyA-mediated responses of germination in Arabidopsis. The Plant Journal, 35(6), 787–799. 10.1046/j.1365-313X.2003.01848.x

Elbaum-Garfinkle, S., Kim, Y., Szczepaniak, K., Chen, C. C.-H., Eckmann, C. R., Myong, S., & Brangwynne, C. P. (2015). The disordered P granule protein LAF-1 drives phase separation into droplets with tunable viscosity and dynamics. Proceedings of the National Academy of Sciences, 112(23), 7189–7194. 10.1073/pnas.1504822112

Filippakopoulos, P., Picaud, S., Mangos, M., Keates, T., Lambert, J.-P., Barsyte-Lovejoy, D., Felletar, I., Volkmer, R., Müller, S., Pawson, T., Gingras, A.-C., Arrowsmith, C. H., & Knapp, S. (2012). Histone Recognition and Large-Scale Structural Analysis of the Human Bromodomain Family. Cell, 149(1), 214–231. 10.1016/j.cell.2012.02.013

Flores, R., Gas, M.-E., Molina-Serrano, D., Nohales, M.-Á., Carbonell, A., Gago, S., De La Peña, M., & Daròs, J.-A. (2009). Viroid Replication: Rolling-Circles, Enzymes and Ribozymes. Viruses, 1(2), 317–334. 10.3390/v1020317

Flores, R., Minoia, S., Carbonell, A., Gisel, A., Delgado, S., López-Carrasco, A., Navarro, B., & Di Serio, F. (2015). Viroids, the simplest RNA replicons: How they manipulate their hosts for being propagated and how their hosts react for containing the infection. Virus Research, 209, 136–145. 10.1016/j.virusres.2015.02.027

Floyd, S. R., Pacold, M. E., Huang, Q., Clarke, S. M., Lam, F. C., Cannell, I. G., Bryson, B. D., Rameseder, J., Lee, M. J., Blake, E. J., Fydrych, A., Ho, R., Greenberger, B. A., Chen, G. C., Maffa, A., Del Rosario, A. M., Root, D. E., Carpenter, A. E., Hahn, W. C., … Yaffe, M. B. (2013). The bromodomain protein Brd4 insulates chromatin from DNA damage signalling. Nature, 498(7453), 246–250. 10.1038/nature12147

Gaucher, J., Boussouar, F., Montellier, E., Curtet, S., Buchou, T., Bertrand, S., Hery, P., Jounier, S., Depaux, A., Vitte, A.-L., Guardiola, P., Pernet, K., Debernardi, A., Lopez, F., Holota, H., Imbert, J., Wolgemuth, D. J., Gérard, M., Rousseaux, S., & Khochbin, S. (2012). Bromodomain-dependent stage-specific male genome programming by Brdt: Brdt: a master regulator of spermatogenesis. The EMBO Journal, 31(19), 3809–3820. 10.1038/emboj.2012.233

Ghosh, U. K., Islam, M. N., Siddiqui, M. N., Cao, X., & Khan, M. A. R. (2022). Proline, a multifaceted signalling molecule in plant responses to abiotic stress: Understanding the physiological mechanisms. Plant Biology, 24(2), 227–239. 10.1111/plb.13363

Hammond, R. W. (2017). Economic Significance of Viroids in Vegetable and Field Crops. In Viroids and Satellites (pp. 5–13). Elsevier. 10.1016/B978-0-12-801498-1.00001-2

Hatos, A., Tosatto, S. C. E., Vendruscolo, M., & Fuxreiter, M. (2022). FuzDrop on AlphaFold: Visualizing the sequence-dependent propensity of liquid–liquid phase separation and aggregation of proteins. Nucleic Acids Research, 50(W1), W337–W344. 10.1093/nar/gkac386

Horsch, R. B., Fry, J. E., Hoffmann, N. L., Wallroth, M., Eichholtz, D., Rogers, S. G., & Fraley, R. T. (1985). A Simple and General Method for Transferring Genes into Plants. Science, 227(4691), 1229–1231. 10.1126/science.227.4691.1229

Jang, M. K., Mochizuki, K., Zhou, M., Jeong, H.-S., Brady, J. N., & Ozato, K. (2005). The Bromodomain Protein Brd4 Is a Positive Regulatory Component of P-TEFb and Stimulates RNA Polymerase II-Dependent Transcription. Molecular Cell, 19(4), Article 4. 10.1016/j.molcel.2005.06.027

Jung, M., Philpott, M., Müller, S., Schulze, J., Badock, V., Eberspächer, U., Moosmayer, D., Bader, B., Schmees, N., Fernández-Montalván, A., & Haendler, B. (2014). Affinity Map of Bromodomain Protein 4 (BRD4) Interactions with the Histone H4 Tail and the Small Molecule Inhibitor JQ1. Journal of Biological Chemistry, 289(13), 9304–9319. 10.1074/jbc.M113.523019

Kalantidis, K., Denti, M. A., Tzortzakaki, S., Marinou, E., Tabler, M., & Tsagris, M. (2007). Virp1 Is a Host Protein with a Major Role in Potato Spindle Tuber Viroid Infection in Nicotiana Plants. Journal of Virology, 81(23), 12872–12880. 10.1128/JVI.00974-07

Kalantidis, K., Tselika, M., Kallemi, P., Bardani, E., Kryovrysanaki, N., & Katsarou, K. (2025). Derailing the host machinery to achieve replication: How viroid and viroid-like RNAs successfully copy their genomes in hostile territory. RNA Biology, 22(1), 1–19. 10.1080/15476286.2025.2538269

Katsarou, K., Adkar-Purushothama, C. R., Tassios, E., Samiotaki, M., Andronis, C., Lisón, P., Nikolaou, C., Perreault, J.-P., & Kalantidis, K. (2022). Revisiting the Non-Coding Nature of Pospiviroids. Cells, 11(2), 265. 10.3390/cells11020265

Katsarou, K., Kryovrysanaki, N., & Kalantidis, K. (2022). Detection of Viroid RNA and vd-siRNA in N. benthamiana Plants: Northern Blot Analyses for Viroid and vd-siRNAs. In A. L. N. Rao, I. Lavagi-Craddock, & G. Vidalakis (Eds.), Viroids (Vol. 2316, pp. 287–312). Springer US. 10.1007/978-1-0716-1464-8_24

Kim, D., Paggi, J. M., Park, C., Bennett, C., & Salzberg, S. L. (2019). Graph-based genome alignment and genotyping with HISAT2 and HISAT-genotype. Nature Biotechnology, 37(8), 907–915. 10.1038/s41587-019-0201-4

Kim, M. J., Shin, R., & Schachtman, D. P. (2009). A Nuclear Factor Regulates Abscisic Acid Responses in Arabidopsis. Plant Physiology, 151(3), Article 3. 10.1104/pp.109.144766

Kosugi, S., Hasebe, M., Tomita, M., & Yanagawa, H. (2009). Systematic identification of cell cycle-dependent yeast nucleocytoplasmic shuttling proteins by prediction of composite motifs. Proceedings of the National Academy of Sciences, 106(25), 10171–10176. 10.1073/pnas.0900604106

Kotekar, A., Singh, A. K., & Devaiah, B. N. (2023). BRD4 and MYC: Power couple in transcription and disease. The FEBS Journal, 290(20), 4820–4842. 10.1111/febs.16580

Lacorte, C., Ribeiro, S. G., Lohuis, D., Goldbach, R., & Prins, M. (2010). Potato virus X and Tobacco mosaic virus-based vectors compatible with the GatewayTM cloning system. Journal of Virological Methods, 164(1–2), 7–13. 10.1016/j.jviromet.2009.11.005

Li, J., Li, Q., Diaz, J., & You, J. (2014). Brd4-Mediated Nuclear Retention of the Papillomavirus E2 Protein Contributes to Its Stabilization in Host Cells. Viruses, 6(1), 319–335. 10.3390/v6010319

Liu, D., Shi, L., Han, C., Yu, J., Li, D., & Zhang, Y. (2012). Validation of Reference Genes for Gene Expression Studies in Virus-Infected Nicotiana benthamiana Using Quantitative Real-Time PCR. PLoS ONE, 7(9), e46451. 10.1371/journal.pone.0046451

Luo, R., Zhuo, C., Liang, M., Liu, H., Gu, X., Zhao, F., & Huang, X. (2025). The GTE7 - IES2B - GTE2 complex epigenetically regulates sulfur homeostasis in Arabidopsis thaliana. New Phytologist, 248(1), 211–230. 10.1111/nph.70398

Ma, J., Dissanayaka Mudiyanselage, S. D., Park, W. J., Wang, M., Takeda, R., Liu, B., & Wang, Y. (2022). A nuclear import pathway exploited by pathogenic noncoding RNAs. The Plant Cell, 34(10), Article 10. 10.1093/plcell/koac210

Maniataki, E., Tabler, M., & Tsagris, M. (2003). Viroid RNA systemic spread may depend on the interaction of a 71-nucleotide bulged hairpin with the host protein VirP1. RNA, 9(3), 346–354. 10.1261/rna.2162203

Martignago, D., Siemiatkowska, B., Lombardi, A., & Conti, L. (2020). Abscisic Acid and Flowering Regulation: Many Targets, Different Places. International Journal of Molecular Sciences, 21(24), 9700. 10.3390/ijms21249700

Martínez De Alba, A. E., Sägesser, R., Tabler, M., & Tsagris, M. (2003). A Bromodomain-Containing Protein from Tomato Specifically Binds Potato Spindle Tuber Viroid RNA In Vitro and In Vivo. Journal of Virology, 77(17), Article 17. 10.1128/JVI.77.17.9685-9694.2003

McBride, A. A., Warburton, A., & Khurana, S. (2021). Multiple Roles of Brd4 in the Infectious Cycle of Human Papillomaviruses. Frontiers in Molecular Biosciences, 8, 725794. 10.3389/fmolb.2021.725794

Misra, A., McKnight, T. D., & Mandadi, K. K. (2018). Bromodomain proteins GTE9 and GTE11 are essential for specific BT2-mediated sugar and ABA responses in Arabidopsis thaliana. Plant Molecular Biology, 96(4–5), 393–402. 10.1007/s11103-018-0704-2

Molliex, A., Temirov, J., Lee, J., Coughlin, M., Kanagaraj, A. P., Kim, H. J., Mittag, T., & Taylor, J. P. (2015). Phase Separation by Low Complexity Domains Promotes Stress Granule Assembly and Drives Pathological Fibrillization. Cell, 163(1), 123–133. 10.1016/j.cell.2015.09.015

Murray, M. G., & Thompson, W. F. (1980). Rapid isolation of high molecular weight plant DNA. Nucleic Acids Research, 8(19), 4321–4326. 10.1093/nar/8.19.4321

Narusaka, M., Shiraishi, T., Iwabuchi, M., & Narusaka, Y. (2010). The floral inoculating protocol: A simplified Arabidopsis thaliana transformation method modified from floral dipping. Plant Biotechnology, 27(4), 349–351. 10.5511/plantbiotechnology.27.349

Navarro, B., Flores, R., & Di Serio, F. (2021). Advances in Viroid-Host Interactions. Annual Review of Virology, 8(1), 305–325. 10.1146/annurev-virology-091919-092331

Neumeier, J., & Meister, G. (2021). siRNA Specificity: RNAi Mechanisms and Strategies to Reduce Off-Target Effects. Frontiers in Plant Science, 11, 526455. 10.3389/fpls.2020.526455

Nguyen Ba, A. N., Pogoutse, A., Provart, N., & Moses, A. M. (2009). NLStradamus: A simple Hidden Markov Model for nuclear localization signal prediction. BMC Bioinformatics, 10(1), 202. 10.1186/1471-2105-10-202

Nott, T. J., Petsalaki, E., Farber, P., Jervis, D., Fussner, E., Plochowietz, A., Craggs, T. D., Bazett-Jones, D. P., Pawson, T., Forman-Kay, J. D., & Baldwin, A. J. (2015). Phase Transition of a Disordered Nuage Protein Generates Environmentally Responsive Membraneless Organelles. Molecular Cell, 57(5), 936–947. 10.1016/j.molcel.2015.01.013

Ottinger, M., Christalla, T., Nathan, K., Brinkmann, M. M., Viejo-Borbolla, A., & Schulz, T. F. (2006). Kaposi’s Sarcoma-Associated Herpesvirus LANA-1 Interacts with the Short Variant of BRD4 and Releases Cells from a BRD4- and BRD2/RING3-Induced G 1 Cell CycleArrest. Journal of Virology, 80(21), Article 21. 10.1128/JVI.00804-06

Pastori, C., Kapranov, P., Penas, C., Peschansky, V., Volmar, C.-H., Sarkaria, J. N., Bregy, A., Komotar, R., St. Laurent, G., Ayad, N. G., & Wahlestedt, C. (2015). The Bromodomain protein BRD4 controls HOTAIR, a long noncoding RNA essential for glioblastoma proliferation. Proceedings of the National Academy of Sciences, 112(27), 8326–8331. 10.1073/pnas.1424220112

Pfaffl, M. W., Tichopad, A., Prgomet, C., & Neuvians, T. P. (2004). Determination of stable housekeeping genes, differentially regulated target genes and sample integrity: BestKeeper – Excel-based tool using pair-wise correlations. Biotechnology Letters, 26(6), 509–515. 10.1023/B:BILE.0000019559.84305.47

Pistoni, M., Rossi, T., Donati, B., Torricelli, F., Polano, M., & Ciarrocchi, A. (2021). Long Noncoding RNA NEAT1 Acts as a Molecular Switch for BRD4 Transcriptional Activity and Mediates Repression of BRD4/WDR5 Target Genes. Molecular Cancer Research, 19(5), 799–811. 10.1158/1541-7786.MCR-20-0324

Qi, Y., & Ding, B. (2003). Differential Subnuclear Localization of RNA Strands of Opposite Polarity Derived from an Autonomously Replicating Viroid[W]. The Plant Cell, 15(11), 2566–2577. 10.1105/tpc.016576

Qi, Y., Peélissier, T., Itaya, A., Hunt, E., Wassenegger, M., & Ding, B. (2004). Direct Role of a Viroid RNA Motif in Mediating Directional RNA Trafficking across a Specific Cellular Boundary[W]. The Plant Cell, 16(7), 1741–1752. 10.1105/tpc.021980

Qian, F., Zhao, Q.-Q., Zhou, J.-X., Yuan, D.-Y., Liu, Z.-Z., Su, Y.-N., Li, L., Chen, S., & He, X.-J. (2024). The GTE4–EML chromatin reader complex concurrently recognizes histone acetylation and H3K4 trimethylation in Arabidopsis. The Plant Cell, 37(1), koae330. 10.1093/plcell/koae330

Rae, G. M., Uversky, V. N., David, K., & Wood, M. (2014). DRM1 and DRM2 expression regulation: Potential role of splice variants in response to stress and environmental factors in Arabidopsis. Molecular Genetics and Genomics, 289(3), 317–332. 10.1007/s00438-013-0804-2

Rahman, S., Sowa, M. E., Ottinger, M., Smith, J. A., Shi, Y., Harper, J. W., & Howley, P. M. (2011). The Brd4 Extraterminal Domain Confers Transcription Activation Independent of pTEFb by Recruiting Multiple Proteins, Including NSD3. Molecular and Cellular Biology, 31(13), 2641–2652. 10.1128/MCB.01341-10

Roy, S., Saxena, S., Sinha, A., & Nandi, A. K. (2020). DORMANCY/AUXIN ASSOCIATED FAMILY PROTEIN 2 of Arabidopsis thaliana is a negative regulator of local and systemic acquired resistance. Journal of Plant Research, 133(3), 409–417. 10.1007/s10265-020-01183-2

Sabari, B. R., Dall’Agnese, A., Boija, A., Klein, I. A., Coffey, E. L., Shrinivas, K., Abraham, B. J., Hannett, N. M., Zamudio, A. V., Manteiga, J. C., Li, C. H., Guo, Y. E., Day, D. S., Schuijers, J., Vasile, E., Malik, S., Hnisz, D., Lee, T. I., Cisse, I. I., … Young, R. A. (2018). Coactivator condensation at super-enhancers links phase separation and gene control. Science, 361(6400), eaar3958. 10.1126/science.aar3958

Schweiger, M.-R., You, J., & Howley, P. M. (2006). Bromodomain Protein 4 Mediates the Papillomavirus E2 Transcriptional Activation Function. Journal of Virology, 80(9), 4276–4285. 10.1128/JVI.80.9.4276-4285.2006

Senthil-Kumar, M., & Mysore, K. S. (2011). Caveat of RNAi in Plants: The Off-Target Effect. In H. Kodama & A. Komamine (Eds.), RNAi and Plant Gene Function Analysis (Vol. 744, pp. 13–25). Humana Press. 10.1007/978-1-61779-123-9_2

Seo, H., Kim, K., & Park, W. J. (2021). Effect of VIRP1 Protein on Nuclear Import of Citrus Exocortis Viroid (CEVd). Biomolecules, 11(1), Article 1. 10.3390/biom11010095

Shen, Y.-Y., Wang, X.-F., Wu, F.-Q., Du, S.-Y., Cao, Z., Shang, Y., Wang, X.-L., Peng, C.-C., Yu, X.-C., Zhu, S.-Y., Fan, R.-C., Xu, Y.-H., & Zhang, D.-P. (2006). The Mg-chelatase H subunit is an abscisic acid receptor. Nature, 443(7113), 823–826. 10.1038/nature05176

Strom, A. R., Eeftens, J. M., Polyachenko, Y., Weaver, C. J., Watanabe, H.-F., Bracha, D., Orlovsky, N. D., Jumper, C. C., Jacobs, W. M., & Brangwynne, C. P. (2024). Interplay of condensation and chromatin binding underlies BRD4 targeting. Molecular Biology of the Cell, 35(6), ar88. 10.1091/mbc.E24-01-0046

Studier, F. W. (2005). Protein production by auto-induction in high-density shaking cultures. Protein Expression and Purification, 41(1), 207–234. 10.1016/j.pep.2005.01.016

Tabler, M., Tzortzakaki, S., & Tsagris, M. (1992). Processing of linear longer-than-unit-length potato spindle tuber viroid RNAs into infectious monomeric circular molecules by a G-specific endoribonuclease. Virology, 190(2), 746–753. 10.1016/0042-6822(92)90912-9

Tian, T., Liu, Y., Yan, H., You, Q., Yi, X., Du, Z., Xu, W., & Su, Z. (2017). agriGO v2.0: A GO analysis toolkit for the agricultural community, 2017 update. Nucleic Acids Research, 45(W1), W122–W129. 10.1093/nar/gkx382

Umehara, T., Nakamura, Y., Jang, M. K., Nakano, K., Tanaka, A., Ozato, K., Padmanabhan, B., & Yokoyama, S. (2010). Structural Basis for Acetylated Histone H4 Recognition by the Human BRD2 Bromodomain. Journal of Biological Chemistry, 285(10), 7610–7618. 10.1074/jbc.M109.062422

Uranga, M., Aragonés, V., Selma, S., Vázquez-Vilar, M., Orzáez, D., & Daròs, J. (2021). Efficient Cas9 multiplex editing using unspaced sgRNA arrays engineering in a Potato virus X vector. The Plant Journal, 106(2), 555–565. 10.1111/tpj.15164

Varet, H., Brillet-Guéguen, L., Coppée, J.-Y., & Dillies, M.-A. (2016). SARTools: A DESeq2- and EdgeR-Based R Pipeline for Comprehensive Differential Analysis of RNA-Seq Data. PLOS ONE, 11(6), e0157022. 10.1371/journal.pone.0157022

Wang, C.-Y., & Filippakopoulos, P. (2015). Beating the odds: BETs in disease. Trends in Biochemical Sciences, 40(8), Article 8. 10.1016/j.tibs.2015.06.002

Wang, Y., Qu, J., Ji, S., Wallace, A. J., Wu, J., Li, Y., Gopalan, V., & Ding, B. (2016). A Land Plant-Specific Transcription Factor Directly Enhances Transcription of a Pathogenic Noncoding RNA Template by DNA-Dependent RNA Polymerase II. The Plant Cell, 28(5), 1094–1107. 10.1105/tpc.16.00100

Wei, T.-L., Wang, Z.-X., He, Y.-F., Xue, S., Zhang, S.-Q., Pei, M.-S., Liu, H.-N., Yu, Y.-H., & Guo, D.-L. (2022). Proline synthesis and catabolism-related genes synergistically regulate proline accumulation in response to abiotic stresses in grapevines. Scientia Horticulturae, 305, 111373. 10.1016/j.scienta.2022.111373

Weissman, J. D., Singh, A. K., Devaiah, B. N., Schuck, P., LaRue, R. C., & Singer, D. S. (2021). The intrinsic kinase activity of BRD4 spans its BD2-B-BID domains. Journal of Biological Chemistry, 297(5), 101326. 10.1016/j.jbc.2021.101326

Wen, W., Meinkotht, J. L., Tsien, R. Y., & Taylor, S. S. (1995). Identification of a signal for rapid export of proteins from the nucleus. Cell, 82(3), 463–473. 10.1016/0092-8674(95)90435-2

Wolff, P., Gilz, R., Schumacher, J., & Riesner, D. (1985). Complexes of viroids with histones and other proteins. Nucleic Acids Research, 13(2), 355–367. 10.1093/nar/13.2.355

Woo, Y., Itaya, A., Owens, R. A., Tang, L., Hammond, R. W., Chou, H., Lai, M. M. C., & Ding, B. (1999). Characterization of nuclear import of potato spindle tuber viroid RNA in permeabilized protoplasts. The Plant Journal, 17(6), 627–635. 10.1046/j.1365-313X.1999.00412.x

Wu, S.-Y., Lee, A.-Y., Hou, S. Y., Kemper, J. K., Erdjument-Bromage, H., Tempst, P., & Chiang, C.-M. (2006). Brd4 links chromatin targeting to HPV transcriptional silencing. Genes & Development, 20(17), 2383–2396. 10.1101/gad.1448206

Yang, Z., Yik, J. H. N., Chen, R., He, N., Jang, M. K., Ozato, K., & Zhou, Q. (2005). Recruitment of P-TEFb for Stimulation of Transcriptional Elongation by the Bromodomain Protein Brd4. Molecular Cell, 19(4), 535–545. 10.1016/j.molcel.2005.06.029

You, J., Srinivasan, V., Denis, G. V., Harrington, W. J., Ballestas, M. E., Kaye, K. M., & Howley, P. M. (2006). Kaposi’s Sarcoma-Associated Herpesvirus Latency-Associated Nuclear Antigen Interacts with Bromodomain Protein Brd4 on Host Mitotic Chromosomes. Journal of Virology, 80(18), Article 18. 10.1128/JVI.00502-06

You, M. K., Shin, H. Y., Kim, Y. J., Ok, S. H., Cho, S. K., Jeung, J. U., Yoo, S. D., Kim, J. K., & Shin, J. S. (2010). Novel Bifunctional Nucleases, OmBBD and AtBBD1, Are Involved in Abscisic Acid-Mediated Callose Deposition in Arabidopsis. Plant Physiology, 152(2), 1015–1029. 10.1104/pp.109.147645

Yu, G., Wang, L.-G., Han, Y., & He, Q.-Y. (2012). clusterProfiler: An R Package for Comparing Biological Themes Among Gene Clusters. OMICS: A Journal of Integrative Biology, 16(5), 284–287. 10.1089/omi.2011.0118

Zhang, Q., & Bartels, D. (2018). Molecular responses to dehydration and desiccation in desiccation-tolerant angiosperm plants. Journal of Experimental Botany, 69(13), 3211–3222. 10.1093/jxb/erx489

Zhang, X., Jiang, L., Wang, G., Yu, L., Zhang, Q., Xin, Q., Wu, W., Gong, Z., & Chen, Z. (2013). Structural Insights into the Abscisic Acid Stereospecificity by the ABA Receptors PYR/PYL/RCAR. PLoS ONE, 8(7), e67477. 10.1371/journal.pone.0067477

Zhao, Y., Owens, R. A., & Hammond, R. W. (2001). Use of a vector based on Potato virus X in a whole plant assay to demonstrate nuclear targeting of Potato spindle tuber viroid. Journal of General Virology, 82(6), 1491–1497. 10.1099/0022-1317-82-6-1491

Zheng, X., Zuo, Z., Yao, P., Li, X., Zhang, Q., & Chen, X. (2025). Bromodomain-containing proteins interact with a non-canonical RNA polymerase II kinase to maintain gene expression upon heat stress. Nature Plants, 11(7), 1416–1428. 10.1038/s41477-025-02044-3

Zhou, Q., Sun, Y., Zhao, X., Yu, Y., Cheng, W., Lu, L., Chu, Z., & Chen, X. (2022). Bromodomain-containing factor GTE4 regulates Arabidopsis immune response. BMC Biology, 20(1), Article 1. 10.1186/s12915-022-01454-5

