## Supplementary figures and images for "VIRP1 bromodomain shapes nuclear condensate formation and has a positive effect on PSTVd accumulation"

### Sopplemental fig 2

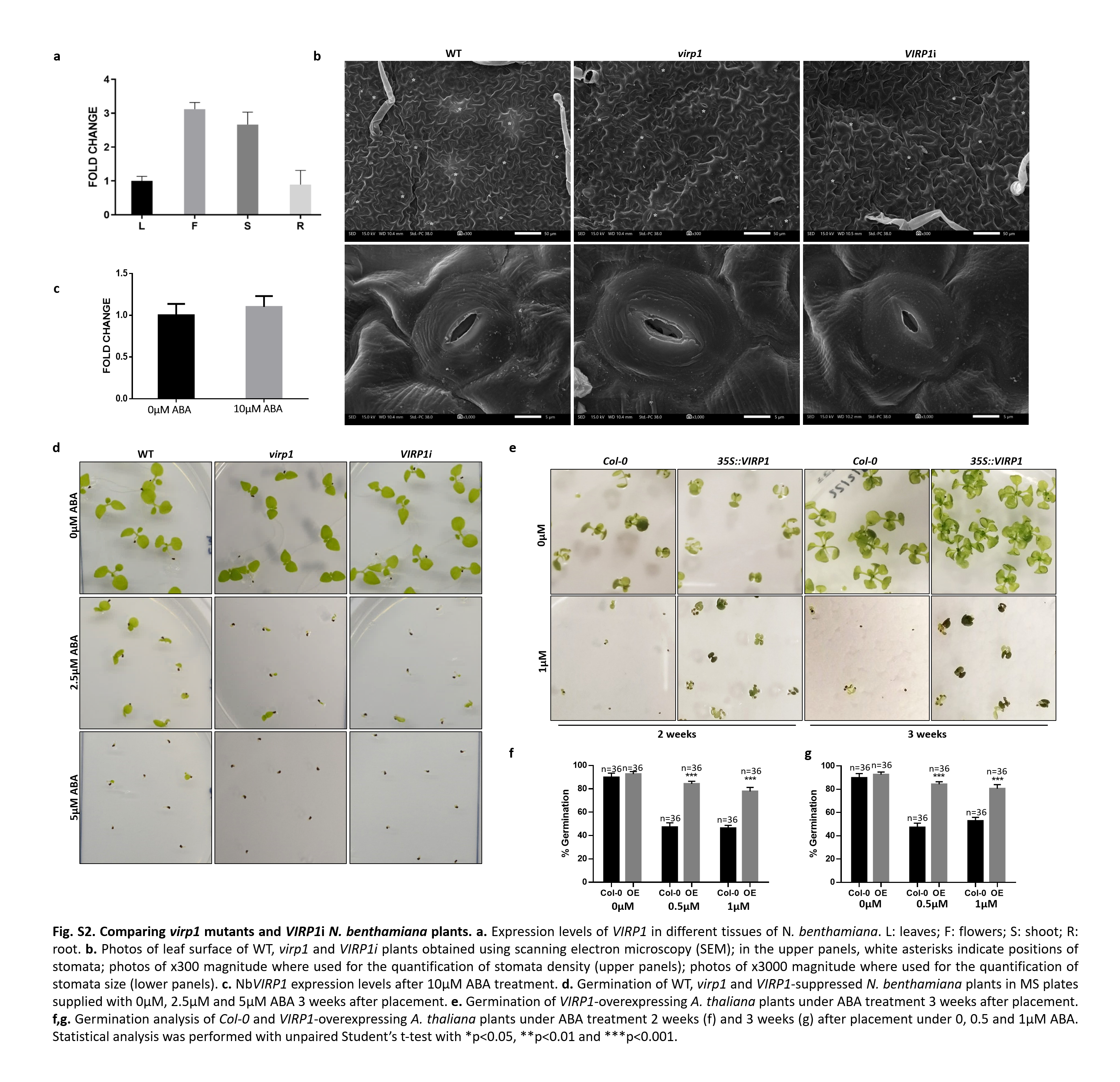

### Supplemental fig 1

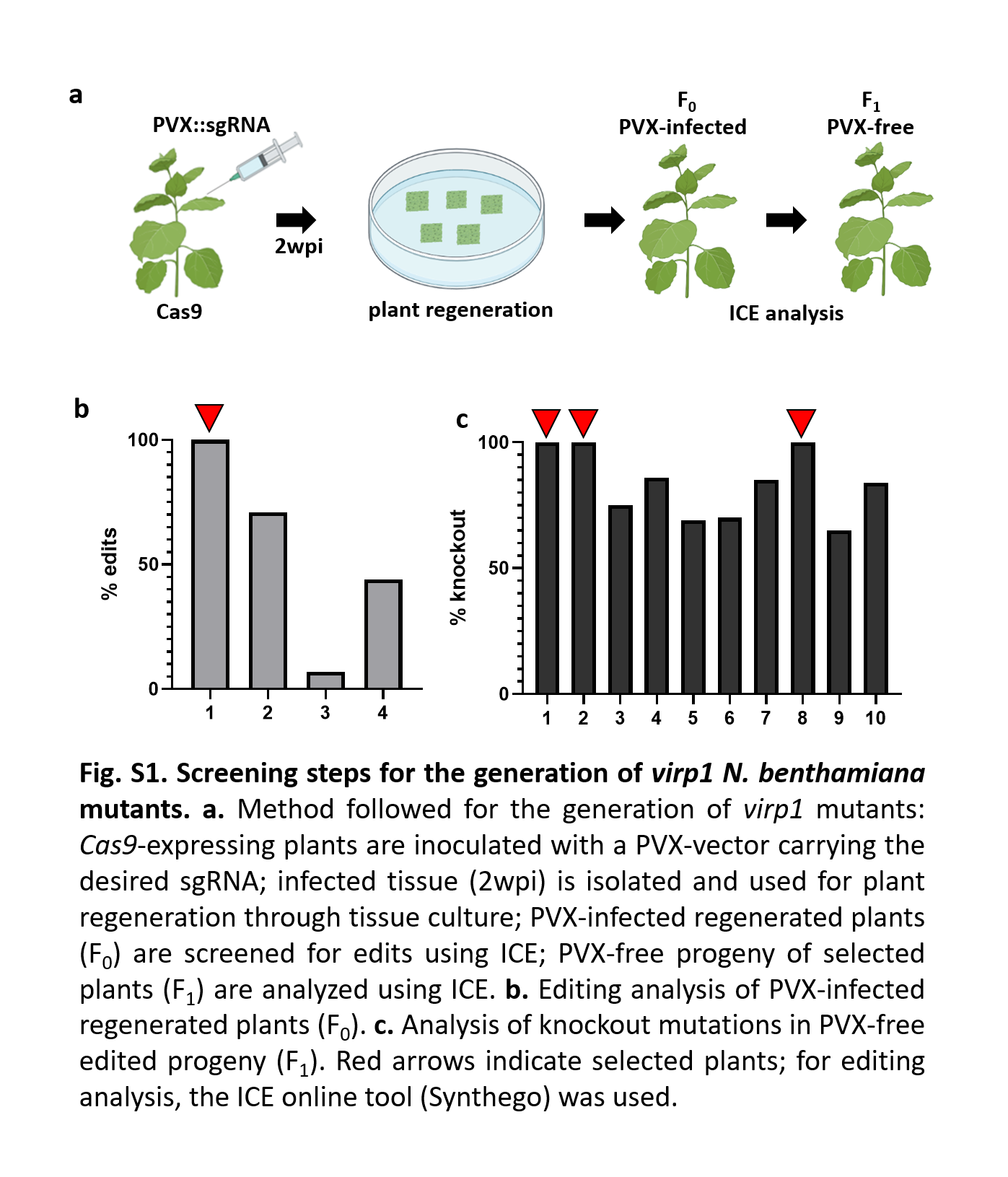

### Supplemental Fig 3

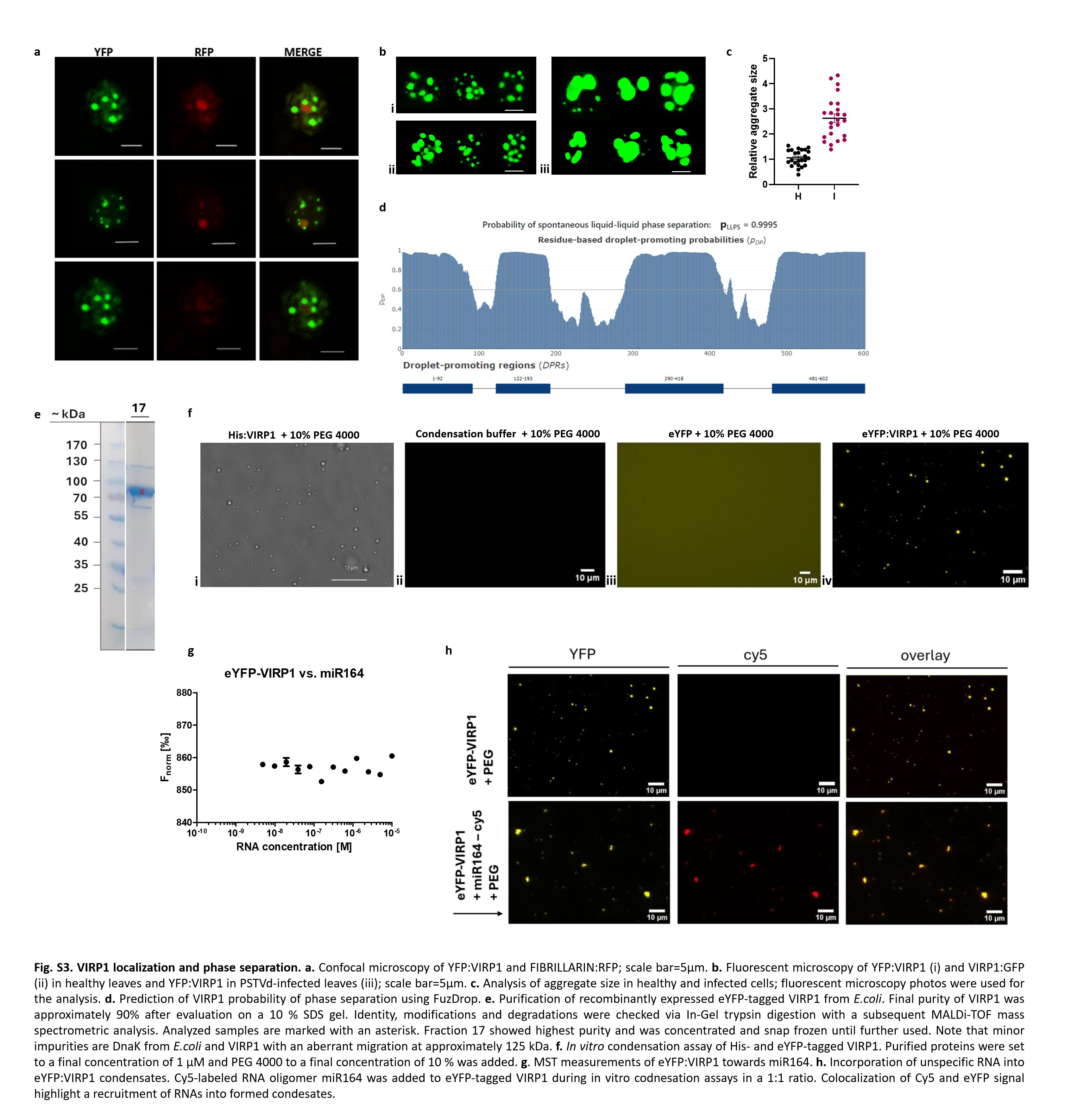

### Supplemental Figure 4

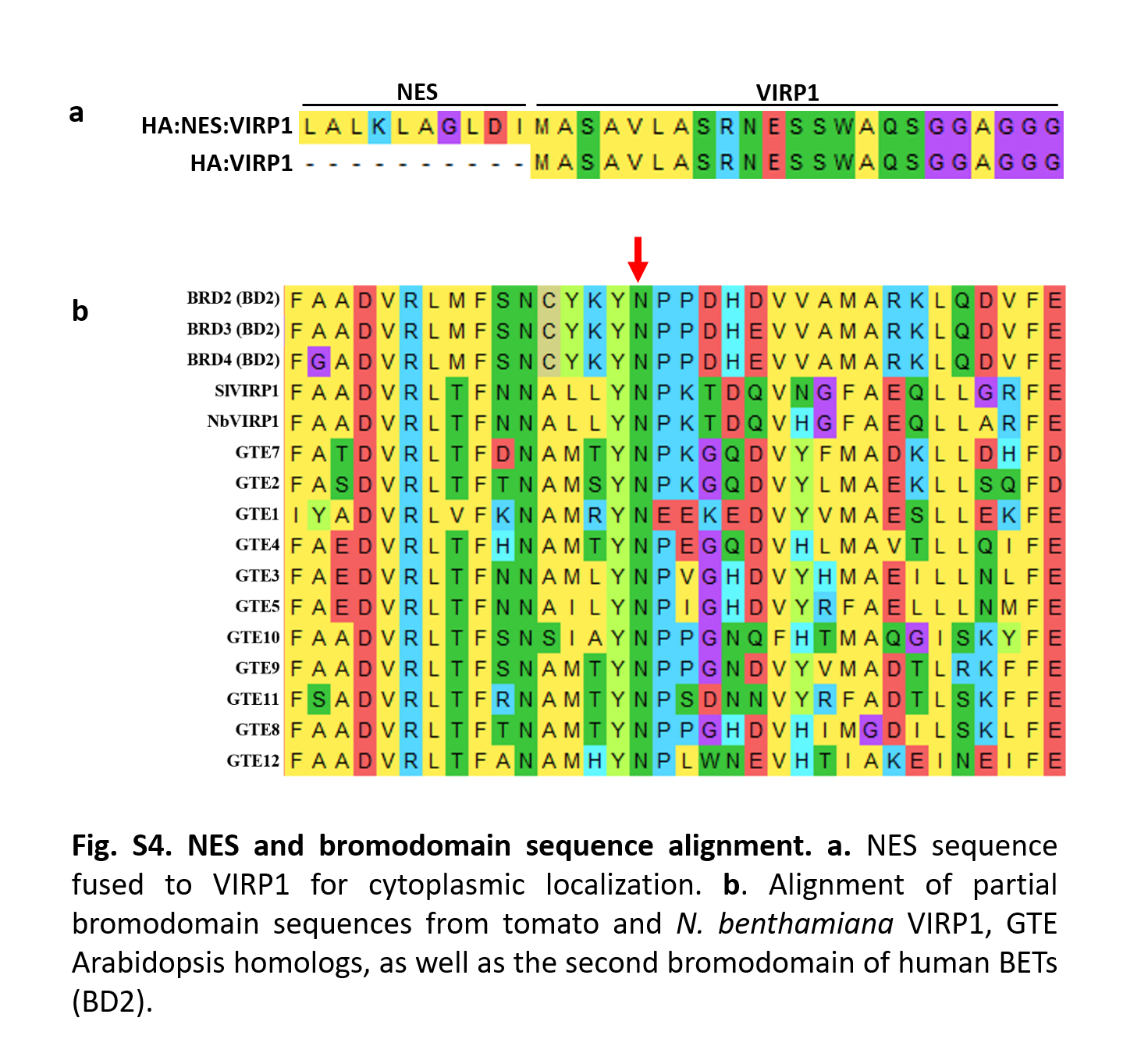

### Supplemental Table 1

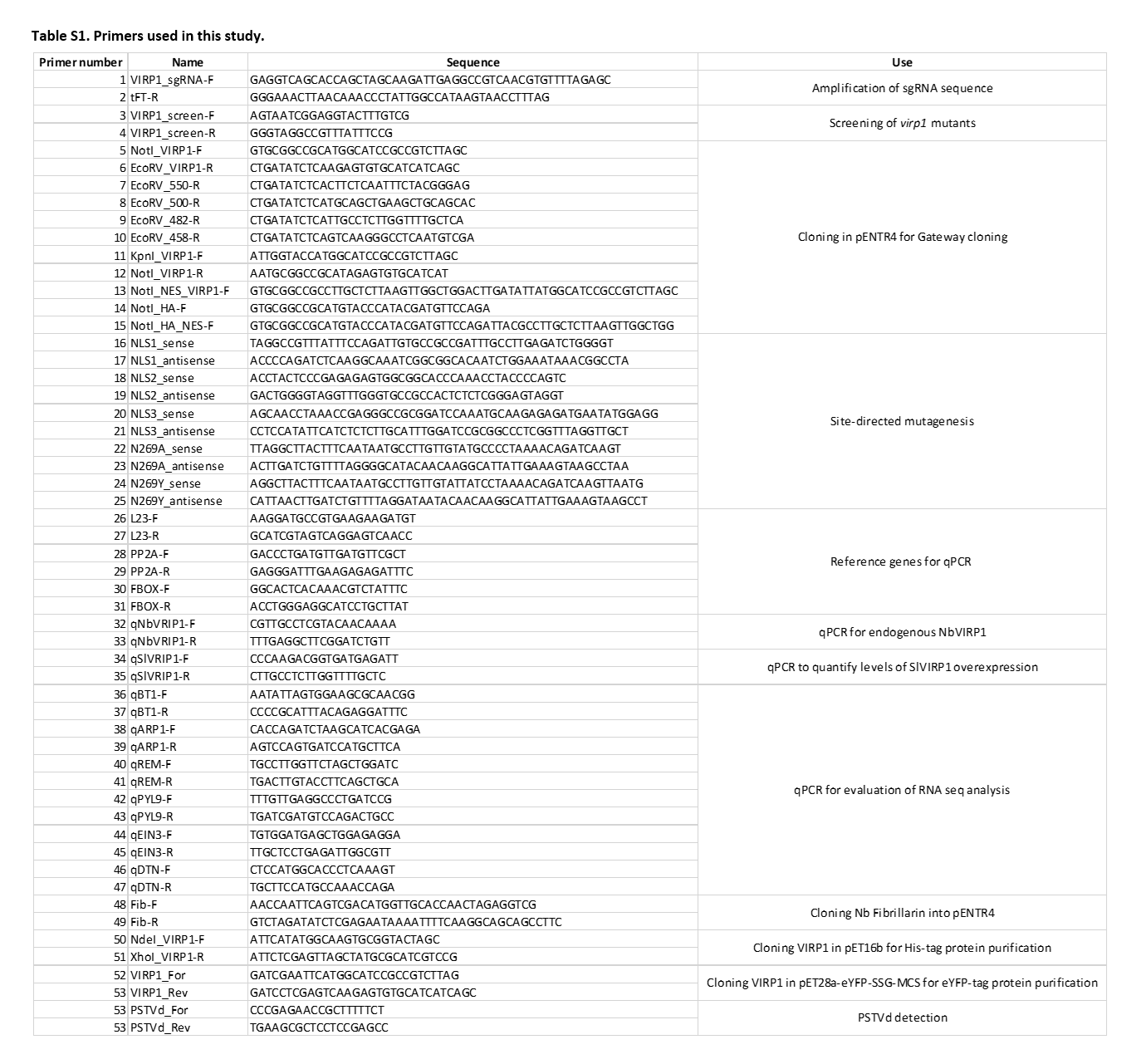

### Supplemental Table 2

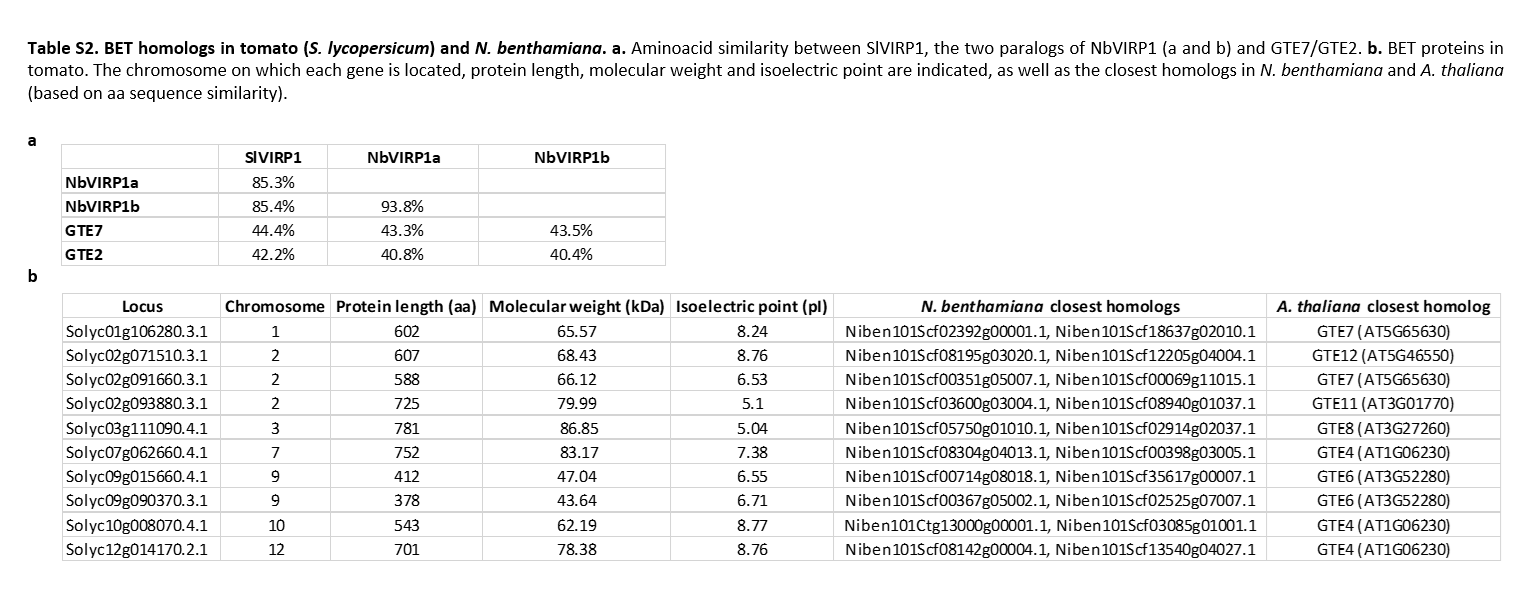

### Supplemental Table 3

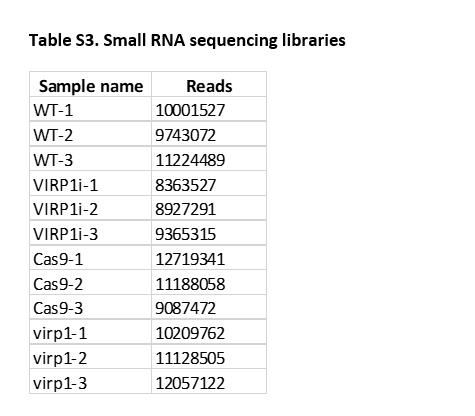
