## Supplemental Table 4 for "VIRP1 bromodomain shapes nuclear condensate formation and has a positive effect on PSTVd accumulation"

**Table S4. Transcriptome analysis of *virp1i N. benthamiana* mutants.**

a. **Upregulated**

| **ID** | **WT av. reads** | ***VIRP1i* av. reads** | **log2FC** | **padj** | **Blast-Hit-Accession** | **Human-Readable-Description** | **Interpro-ID (Description)** | **Gene-Ontology-ID (Name)** |
| --- | --- | --- | --- | --- | --- | --- | --- | --- |
| Niben101Scf13259g00024.1 | 13 | 0 | -6.686 | 4.49867E-05 | ref\|WP_032467012.1\| | RNA methyltransferase [Streptococcus pyogenes] | NA | NA |
| Niben101Scf07349g00005.1 | 10 | 0 | -6.248 | 0.000294263 | sp\|P55748\|CBP22_HORVU | Serine carboxypeptidase II-2 | IPR001563 (Peptidase S10, serine carboxypeptidase), IPR029058 (Alpha/Beta hydrolase fold) | GO:0004185 (serine-type carboxypeptidase activity), GO:0006508 (proteolysis) |
| Niben101Scf00271g09020.1 | 16 | 0 | -5.044 | 3.67047E-06 | NA | Unknown protein | NA | NA |
| Niben101Scf02059g02014.1 | 14 | 0 | -4.723 | 0.001842639 | NA | Unknown protein | NA | NA |
| Niben101Scf11512g00003.1 | 23 | 1 | -3.837 | 0.000965359 | NA | Unknown protein | NA | NA |
| Niben101Scf01009g01017.1 | 15 | 1 | -3.638 | 8.53572E-07 | emb\|CDX93386.1\| | BnaC04g45100D [Brassica napus] | NA | NA |
| Niben101Scf14883g03003.1 | 10 | 1 | -3.531 | 0.000792199 | sp\|Q8CX33\|GLMS_SHEON | Glutamine--fructose-6-phosphate aminotransferase [isomerizing] | IPR005855 (Glucosamine-fructose-6-phosphate aminotransferase, isomerising) | GO:0004360 (glutamine-fructose-6-phosphate transaminase (isomerizing) activity), GO:0005975 (carbohydrate metabolic process), GO:0016051 (carbohydrate biosynthetic process), GO:0030246 (carbohydrate binding) |
| Niben101Scf10489g00006.1 | 13 | 1 | -3.369 | 0.001589278 | AT3G17690.1 | cyclic nucleotide gated channel 19 LENGTH=729 | IPR014710 (RmlC-like jelly roll fold) | NA |
| Niben101Scf06958g00007.1 | 12 | 1 | -3.248 | 0.000497694 | AT1G27150.1 | Tetratricopeptide repeat (TPR)-like superfamily protein LENGTH=468 | IPR011990 (Tetratricopeptide-like helical domain) | GO:0005515 (protein binding) |
| Niben101Scf02043g02002.1 | 11 | 1 | -3.181 | 0.005547978 | sp\|B9JZ89\|DNAJ_AGRVS | Chaperone protein DnaJ | IPR001623 (DnaJ domain), IPR002939 (Chaperone DnaJ, C-terminal) | GO:0006457 (protein folding), GO:0051082 (unfolded protein binding) |
| Niben101Scf03737g03008.1 | 11 | 1 | -3.158 | 0.000950653 | AT5G52580.2 | RabGAP/TBC domain-containing protein LENGTH=690 | IPR000195 (Rab-GTPase-TBC domain), IPR021935 (Domain of unknown function DUF3548) | GO:0005097 (Rab GTPase activator activity), GO:0032313 (regulation of Rab GTPase activity) |
| Niben101Scf03823g01006.1 | 11 | 1 | -3.117 | 0.001748364 | sp\|Q43111\|PME3_PHAVU | Pectinesterase 3 | IPR006501 (Pectinesterase inhibitor domain), IPR011050 (Pectin lyase fold/virulence factor) | GO:0004857 (enzyme inhibitor activity), GO:0005618 (cell wall), GO:0030599 (pectinesterase activity), GO:0042545 (cell wall modification) |
| Niben101Scf00457g06012.1 | 10 | 1 | -3.066 | 0.016245873 | sp\|Q6DRB1\|GLE1_DANRE | Nucleoporin GLE1 | IPR012476 (GLE1-like) | GO:0005643 (nuclear pore), GO:0016973 (poly(A)+ mRNA export from nucleus) |
| Niben101Scf01486g02004.1 | 33 | 4 | -3 | 4.56963E-07 | AT4G31170.1 | Protein kinase superfamily protein LENGTH=412 | IPR002912 (ACT domain), IPR011009 (Protein kinase-like domain) | GO:0004672 (protein kinase activity), GO:0004674 (protein serine/threonine kinase activity), GO:0005524 (ATP binding), GO:0006468 (protein phosphorylation), GO:0008152 (metabolic process), GO:0016597 (amino acid binding), GO:0016772 (transferase activity, transferring phosphorus-containing groups) |
| Niben101Scf03488g06005.1 | 15 | 2 | -2.954 | 0.000146022 | gb\|ACZ64220.1\| | RPM1 interacting protein 4 transcript 2 [Lactuca saligna] | IPR008700 (Pathogenic type III effector avirulence factor Avr cleavage site) | NA |
| Niben101Scf02497g01020.1 | 58 | 7 | -2.941 | 3.60323E-15 | sp\|A6QP05\|DHR12_BOVIN | Dehydrogenase/reductase SDR family member 12 | IPR002347 (Glucose/ribitol dehydrogenase) | GO:0008152 (metabolic process), GO:0016491 (oxidoreductase activity) |
| Niben101Scf01243g00004.1 | 14 | 2 | -2.868 | 0.002013212 | emb\|CDY14672.1\| | BnaC05g19290D [Brassica napus] | NA | NA |
| Niben101Scf04099g09005.1 | 12 | 1 | -2.856 | 0.009421686 | sp\|P74518\|LRTA_SYNY3 | Light-repressed protein A homolog | IPR003489 (Ribosomal protein S30Ae/sigma 54 modulation protein) | GO:0044238 (primary metabolic process) |
| Niben101Scf11952g00008.1 | 14 | 2 | -2.843 | 0.002838954 | tpg\|DAA47775.1\| | TPA: hypothetical protein ZEAMMB73_248377 [Zea mays] | NA | NA |
| Niben101Scf03638g02002.1 | 11 | 1 | -2.825 | 0.001605157 | sp\|Q9M9M4\|CSLD3_ARATH | Cellulose synthase-like protein D3 | IPR005150 (Cellulose synthase), IPR013083 (Zinc finger, RING/FYVE/PHD-type), IPR029044 (Nucleotide-diphospho-sugar transferases) | GO:0016020 (membrane), GO:0016760 (cellulose synthase (UDP-forming) activity), GO:0030244 (cellulose biosynthetic process) |
| Niben101Scf07623g01035.1 | 28 | 3 | -2.781 | 0.024422998 | sp\|Q3J5H6\|GLND_RHOS4 | Bifunctional uridylyltransferase/uridylyl-removing enzyme | NA | NA |
| Niben101Scf00753g03016.1 | 16 | 2 | -2.776 | 0.00182607 | sp\|Q9XF43\|KCS6_ARATH | 3-ketoacyl-CoA synthase 6 | IPR012392 (Very-long-chain 3-ketoacyl-CoA synthase), IPR016039 (Thiolase-like) | GO:0003824 (catalytic activity), GO:0006633 (fatty acid biosynthetic process), GO:0008152 (metabolic process), GO:0008610 (lipid biosynthetic process), GO:0016020 (membrane), GO:0016747 (transferase activity, transferring acyl groups other than amino-acyl groups) |
| Niben101Scf06172g04003.1 | 13 | 2 | -2.728 | 0.005837906 | sp\|P50258\|TBAD_PHYPO | Tubulin alpha-1A chain | IPR000217 (Tubulin), IPR023123 (Tubulin, C-terminal) | GO:0003924 (GTPase activity), GO:0005200 (structural constituent of cytoskeleton), GO:0005525 (GTP binding), GO:0005874 (microtubule), GO:0006184 (GTP catabolic process), GO:0007017 (microtubule-based process), GO:0043234 (protein complex), GO:0051258 (protein polymerization) |
| Niben101Scf01959g01001.1 | 15 | 2 | -2.713 | 0.000291688 | ref\|NP_001047761.1\| | product [Oryza sativa Japonica Group] | NA | NA |
| Niben101Scf01240g05002.1 | 10 | 1 | -2.699 | 0.018700104 | AT5G20110.1 | Dynein light chain type 1 family protein LENGTH=209 | IPR001372 (Dynein light chain, type 1/2) | GO:0005875 (microtubule associated complex), GO:0007017 (microtubule-based process) |
| Niben101Scf11886g03036.1 | 10 | 1 | -2.699 | 0.005864599 | emb\|CDY68259.1\| | BnaCnng58260D [Brassica napus] | NA | NA |
| Niben101Scf08160g02011.1 | 10 | 1 | -2.698 | 0.009730659 | NA | Unknown protein | NA | NA |
| Niben101Scf01063g10002.1 | 10 | 1 | -2.686 | 0.034295037 | sp\|Q8RWS8\|PP199_ARATH | Pentatricopeptide repeat-containing protein | IPR002885 (Pentatricopeptide repeat), IPR011990 (Tetratricopeptide-like helical domain), IPR018087 (Glycoside hydrolase, family 5, conserved site) | GO:0004553 (hydrolase activity, hydrolyzing O-glycosyl compounds), GO:0005515 (protein binding), GO:0005975 (carbohydrate metabolic process) |
| Niben101Scf00428g05020.1 | 10 | 1 | -2.683 | 0.012716326 | AT1G20960.1 | U5 small nuclear ribonucleoprotein helicase, putative LENGTH=2171 | IPR014001 (Helicase superfamily 1/2, ATP-binding domain), IPR027417 (P-loop containing nucleoside triphosphate hydrolase) | GO:0003676 (nucleic acid binding), GO:0005524 (ATP binding) |
| Niben101Scf06942g02033.1 | 10 | 1 | -2.668 | 0.028434722 | sp\|Q56YT0\|LAC3_ARATH | Laccase-3 | IPR017761 (Laccase) | GO:0005507 (copper ion binding), GO:0016491 (oxidoreductase activity), GO:0046274 (lignin catabolic process), GO:0048046 (apoplast), GO:0052716 (hydroquinone:oxygen oxidoreductase activity), GO:0055114 (oxidation-reduction process) |
| Niben101Scf06809g00012.1 | 10 | 1 | -2.658 | 0.036637245 | AT2G36770.1 | UDP-Glycosyltransferase superfamily protein LENGTH=496 | IPR002213 (UDP-glucuronosyl/UDP-glucosyltransferase) | GO:0008152 (metabolic process), GO:0016758 (transferase activity, transferring hexosyl groups) |
| Niben101Scf02026g04007.1 | 15 | 2 | -2.657 | 0.000753344 | AT2G25180.1 | response regulator 12 LENGTH=596 | IPR010402 (CCT domain), IPR011006 (CheY-like superfamily) | GO:0000160 (phosphorelay signal transduction system), GO:0005515 (protein binding) |
| Niben101Scf00419g00001.1 | 21 | 3 | -2.621 | 7.61532E-05 | ref\|WP_025069310.1\| | acetyl-CoA carboxylase [Bacteroides propionicifaciens] | IPR007934 (Alpha-L-arabinofuranosidase B, arabinose-binding domain), IPR012878 (Protein of unknown function DUF1680) | GO:0046373 (L-arabinose metabolic process), GO:0046556 (alpha-L-arabinofuranosidase activity) |
| Niben101Scf18879g04004.1 | 24 | 4 | -2.575 | 0.009730659 | sp\|B6EUA9\|PR40A_ARATH | Pre-mRNA-processing protein 40A | IPR001202 (WW domain), IPR002713 (FF domain), IPR027237 (Pre-mRNA-processing protein PRP40) | GO:0005515 (protein binding) |
| Niben101Scf19421g00012.1 | 16 | 2 | -2.564 | 0.01340991 | AT1G69830.1 | alpha-amylase-like 3 LENGTH=887 | IPR015902 (Glycoside hydrolase, family 13) | GO:0003824 (catalytic activity), GO:0004556 (alpha-amylase activity), GO:0005509 (calcium ion binding), GO:0005975 (carbohydrate metabolic process), GO:0043169 (cation binding) |
| Niben101Scf08613g02006.1 | 29 | 4 | -2.562 | 8.24307E-06 | sp\|O65572\|CCD1_ARATH | Carotenoid 9,10(9',10')-cleavage dioxygenase 1 | IPR004294 (Carotenoid oxygenase) | NA |
| Niben101Scf06310g00007.1 | 11 | 2 | -2.505 | 0.024270484 | AT3G10560.1 | Cytochrome P450 superfamily protein LENGTH=514 | IPR001128 (Cytochrome P450) | GO:0005506 (iron ion binding), GO:0016705 (oxidoreductase activity, acting on paired donors, with incorporation or reduction of molecular oxygen), GO:0020037 (heme binding), GO:0055114 (oxidation-reduction process) |
| Niben101Scf22728g00001.1 | 11 | 2 | -2.488 | 0.005667131 | sp\|Q9ZU11\|ATB15_ARATH | Homeobox-leucine zipper protein ATHB-15 | NA | NA |
| Niben101Scf39332g00017.1 | 11 | 2 | -2.484 | 0.003836471 | sp\|P47990\|XDH_CHICK | Xanthine dehydrogenase/oxidase | IPR000674 (Aldehyde oxidase/xanthine dehydrogenase, a/b hammerhead), IPR002888 ([2Fe-2S]-binding), IPR005107 (CO dehydrogenase flavoprotein, C-terminal), IPR008274 (Aldehyde oxidase/xanthine dehydrogenase, molybdopterin binding), IPR012675 (Beta-grasp domain), IPR016166 (FAD-binding, type 2) | GO:0003824 (catalytic activity), GO:0008762 (UDP-N-acetylmuramate dehydrogenase activity), GO:0009055 (electron carrier activity), GO:0016491 (oxidoreductase activity), GO:0016614 (oxidoreductase activity, acting on CH-OH group of donors), GO:0046872 (metal ion binding), GO:0050660 (flavin adenine dinucleotide binding), GO:0051536 (iron-sulfur cluster binding), GO:0051537 (2 iron, 2 sulfur cluster binding), GO:0055114 (oxidation-reduction process) |
| Niben101Scf16608g00001.1 | 25 | 4 | -2.482 | 3.8204E-06 | sp\|Q3C1H3\|ATPF_NICSY | ATP synthase subunit b, chloroplastic | IPR002146 (ATPase, F0 complex, subunit B/B', bacterial/chloroplast) | GO:0015078 (hydrogen ion transmembrane transporter activity), GO:0015986 (ATP synthesis coupled proton transport), GO:0045263 (proton-transporting ATP synthase complex, coupling factor F(o)) |
| Niben101Scf00479g01016.1 | 11 | 2 | -2.457 | 0.044200053 | sp\|Q10281\|GBLP_SCHPO | Guanine nucleotide-binding protein subunit beta-like protein | IPR015943 (WD40/YVTN repeat-like-containing domain) | GO:0005515 (protein binding) |
| Niben101Scf01240g12007.1 | 31 | 5 | -2.456 | 1.07773E-06 | sp\|Q07E28\|CTTB2_NEONE | Cortactin-binding protein 2 | IPR014710 (RmlC-like jelly roll fold), IPR020683 (Ankyrin repeat-containing domain), IPR021789 (Potassium channel, plant-type) | GO:0005515 (protein binding) |
| Niben101Scf04963g04005.1 | 11 | 2 | -2.441 | 0.01478479 | sp\|P0C1U4\|GUN9_ORYSJ | Endoglucanase 9 | IPR001701 (Glycoside hydrolase, family 9), IPR008928 (Six-hairpin glycosidase-like) | GO:0003824 (catalytic activity), GO:0004553 (hydrolase activity, hydrolyzing O-glycosyl compounds), GO:0005975 (carbohydrate metabolic process) |
| Niben101Scf00077g05011.1 | 17 | 3 | -2.431 | 0.002255897 | sp\|Q9CR16\|PPID_MOUSE | Peptidyl-prolyl cis-trans isomerase D | IPR001179 (Peptidyl-prolyl cis-trans isomerase, FKBP-type, domain), IPR011990 (Tetratricopeptide-like helical domain), IPR023566 (Peptidyl-prolyl cis-trans isomerase, FKBP-type) | GO:0005515 (protein binding), GO:0006457 (protein folding) |
| Niben101Scf00988g04026.1 | 12 | 2 | -2.419 | 0.006754545 | sp\|A2AJK6\|CHD7_MOUSE | Chromodomain-helicase-DNA-binding protein 7 | IPR027417 (P-loop containing nucleoside triphosphate hydrolase) | NA |
| Niben101Scf01073g05003.1 | 32 | 5 | -2.403 | 5.20905E-05 | sp\|P78958\|G3P1_SCHPO | Glyceraldehyde-3-phosphate dehydrogenase 1 | IPR020831 (Glyceraldehyde/Erythrose phosphate dehydrogenase family) | GO:0006006 (glucose metabolic process), GO:0016620 (oxidoreductase activity, acting on the aldehyde or oxo group of donors, NAD or NADP as acceptor), GO:0050661 (NADP binding), GO:0051287 (NAD binding), GO:0055114 (oxidation-reduction process) |
| Niben101Scf10620g00003.1 | 10 | 2 | -2.394 | 0.012579582 | sp\|Q9NPJ3\|ACO13_HUMAN | Acyl-coenzyme A thioesterase 13 | IPR029069 (HotDog domain) | NA |
| Niben101Scf02467g01003.1 | 14 | 2 | -2.385 | 0.005175203 | sp\|Q9FEW9\|OPR3_SOLLC | 12-oxophytodienoate reductase 3 | IPR013785 (Aldolase-type TIM barrel) | GO:0003824 (catalytic activity), GO:0010181 (FMN binding), GO:0016491 (oxidoreductase activity), GO:0055114 (oxidation-reduction process) |
| Niben101Scf02408g02009.1 | 20 | 3 | -2.384 | 0.000868946 | AT2G15620.1 | nitrite reductase 1 LENGTH=586 | IPR005117 (Nitrite/Sulfite reductase ferredoxin-like domain), IPR006067 (Nitrite/sulphite reductase 4Fe-4S domain) | GO:0016491 (oxidoreductase activity), GO:0020037 (heme binding), GO:0051536 (iron-sulfur cluster binding), GO:0055114 (oxidation-reduction process) |
| Niben101Scf00063g03008.1 | 12 | 2 | -2.383 | 0.023269348 | AT2G03350.1 | Protein of unknown function, DUF538 LENGTH=179 | IPR007493 (Protein of unknown function DUF538) | NA |
| Niben101Scf08149g01007.1 | 10 | 2 | -2.379 | 0.024709764 | sp\|Q03065\|RPSB_NOSS1 | RNA polymerase sigma-B factor | IPR014284 (RNA polymerase sigma-70 like domain) | GO:0003677 (DNA binding), GO:0003700 (sequence-specific DNA binding transcription factor activity), GO:0006352 (DNA-templated transcription, initiation), GO:0006355 (regulation of transcription, DNA-templated), GO:0016987 (sigma factor activity) |
| Niben101Scf04410g03007.1 | 10 | 2 | -2.358 | 0.014074117 | sp\|Q9ZPQ3\|GEK1_ARATH | D-aminoacyl-tRNA deacylase | IPR007508 (D-aminoacyl-tRNA deacylase) | GO:0016788 (hydrolase activity, acting on ester bonds), GO:0051499 (D-aminoacyl-tRNA deacylase activity) |
| Niben101Scf00262g00006.1 | 10 | 2 | -2.339 | 0.025719435 | sp\|A6MMB7\|PSBD_CHLSC | Photosystem II D2 protein | NA | NA |
| Niben101Scf05056g02015.1 | 76 | 13 | -2.337 | 2.84539E-09 | sp\|P40937\|RFC5_HUMAN | Replication factor C subunit 5 | IPR003959 (ATPase, AAA-type, core), IPR008921 (DNA polymerase III, clamp loader complex, gamma/delta/delta subunit, C-terminal), IPR027417 (P-loop containing nucleoside triphosphate hydrolase) | GO:0003677 (DNA binding), GO:0005524 (ATP binding), GO:0006260 (DNA replication) |
| Niben101Scf11383g01001.1 | 12 | 2 | -2.336 | 0.026524819 | ref\|XP_002512246.1\| | conserved hypothetical protein [Ricinus communis] gb\|EEF49698.1\| conserved hypothetical protein [Ricinus communis] | IPR010903 (Protein of unknown function DUF1517) | NA |
| Niben101Scf01037g01016.1 | 14 | 2 | -2.33 | 0.003873829 | sp\|Q0KL01\|UBX2B_MOUSE | UBX domain-containing protein 2B | IPR009060 (UBA-like), IPR012989 (SEP domain), IPR029071 (Ubiquitin-related domain) | GO:0005515 (protein binding) |
| Niben101Scf01834g02011.1 | 129 | 23 | -2.33 | 5.79868E-10 | AT1G49975.1 | INVOLVED IN: photosynthesis; LOCATED IN: photosystem I, chloroplast, thylakoid membrane; EXPRESSED IN: 20 plant structures; EXPRESSED DURING: 13 growth stages; | IPR008796 (Photosystem I PsaN, reaction centre subunit N) | GO:0005516 (calmodulin binding), GO:0009522 (photosystem I), GO:0015979 (photosynthesis), GO:0042651 (thylakoid membrane) |
| Niben101Scf02795g00008.1 | 227 | 41 | -2.327 | 0.00010679 | sp\|P09144\|PSAB_CHLRE | Photosystem I P700 chlorophyll a apoprotein A2 | IPR001280 (Photosystem I PsaA/PsaB) | GO:0009522 (photosystem I), GO:0009579 (thylakoid), GO:0015979 (photosynthesis), GO:0016021 (integral component of membrane) |
| Niben101Scf03141g09006.1 | 12 | 2 | -2.306 | 0.018130506 | AT1G66500.1 | Pre-mRNA cleavage complex II LENGTH=416 | IPR007087 (Zinc finger, C2H2) | GO:0046872 (metal ion binding) |
| Niben101Scf00691g04016.1 | 13 | 2 | -2.305 | 0.002420138 | AT5G58000.1 | Reticulon family protein LENGTH=487 | IPR003388 (Reticulon) | NA |
| Niben101Scf04629g02001.1 | 32 | 6 | -2.282 | 0.000239267 | sp\|Q93VH2\|BBD2_ARATH | Bifunctional nuclease 2 | IPR003729 (Bifunctional nuclease domain) | GO:0004518 (nuclease activity) |
| Niben101Scf05282g00001.1 | 15 | 3 | -2.271 | 0.010224147 | sp\|O04504\|PPR27_ARATH | Pentatricopeptide repeat-containing protein | IPR002885 (Pentatricopeptide repeat), IPR011990 (Tetratricopeptide-like helical domain) | GO:0005515 (protein binding) |
| Niben101Scf06633g02009.1 | 75 | 14 | -2.262 | 2.12552E-08 | sp\|Q9T076\|ENL2_ARATH | Early nodulin-like protein 2 | IPR008972 (Cupredoxin) | GO:0009055 (electron carrier activity) |
| Niben101Scf19043g00021.1 | 11 | 2 | -2.257 | 0.024270484 | sp\|Q8H038\|KATAM_ORYSJ | Xyloglucan galactosyltransferase KATAMARI1 homolog | IPR004263 (Exostosin-like) | NA |
| Niben101Scf01008g01008.1 | 1668 | 314 | -2.247 | 0.003223347 | AT2G33830.2 | Dormancy/auxin associated family protein LENGTH=108 | IPR008406 (Dormancyauxin associated) | NA |
| Niben101Scf02752g01011.1 | 16 | 3 | -2.245 | 0.012263231 | sp\|Q8VY89\|APC7_ARATH | Anaphase-promoting complex subunit 7 | IPR011990 (Tetratricopeptide-like helical domain) | GO:0005515 (protein binding) |
| Niben101Scf01001g06011.1 | 22 | 4 | -2.242 | 0.000602397 | sp\|O80340\|ERF78_ARATH | Ethylene-responsive transcription factor 4 | IPR016177 (DNA-binding domain) | GO:0003677 (DNA binding), GO:0003700 (sequence-specific DNA binding transcription factor activity), GO:0006355 (regulation of transcription, DNA-templated) |
| Niben101Scf24355g00006.1 | 11 | 2 | -2.225 | 0.047999884 | sp\|Q8KCD8\|DNAJ_CHLTE | Chaperone protein DnaJ | IPR001623 (DnaJ domain) | NA |
| Niben101Scf01942g06001.1 | 13 | 2 | -2.222 | 0.02840987 | gb\|KEH43633.1\| | late embryogenesis abundant protein, putative [Medicago truncatula] | NA | NA |
| Niben101Scf19421g00012.1 | 14 | 3 | -2.216 | 0.016468532 | AT1G69830.1 | alpha-amylase-like 3 LENGTH=887 | IPR015902 (Glycoside hydrolase, family 13) | GO:0003824 (catalytic activity), GO:0004556 (alpha-amylase activity), GO:0005509 (calcium ion binding), GO:0005975 (carbohydrate metabolic process), GO:0043169 (cation binding) |
| Niben101Scf07790g02006.1 | 14 | 3 | -2.214 | 0.007900009 | AT3G10560.1 | Cytochrome P450 superfamily protein LENGTH=514 | IPR001128 (Cytochrome P450) | GO:0005506 (iron ion binding), GO:0016705 (oxidoreductase activity, acting on paired donors, with incorporation or reduction of molecular oxygen), GO:0020037 (heme binding), GO:0055114 (oxidation-reduction process) |
| Niben101Scf08026g02001.1 | 11 | 2 | -2.206 | 0.036749931 | AT2G35330.1 | RING/U-box superfamily protein LENGTH=738 | IPR013083 (Zinc finger, RING/FYVE/PHD-type) | GO:0005515 (protein binding), GO:0008270 (zinc ion binding) |
| Niben101Scf05222g04018.1 | 14 | 3 | -2.192 | 0.007747848 | NA | Unknown protein | NA | NA |
| Niben101Scf03679g03009.1 | 26 | 5 | -2.19 | 0.000115249 | AT5G51100.1 | Fe superoxide dismutase 2 LENGTH=305 | IPR001189 (Manganese/iron superoxide dismutase) | GO:0004784 (superoxide dismutase activity), GO:0006801 (superoxide metabolic process), GO:0046872 (metal ion binding), GO:0055114 (oxidation-reduction process) |
| Niben101Scf04187g00028.1 | 14 | 3 | -2.184 | 0.007566311 | sp\|P19484\|TFEB_HUMAN | Transcription factor EB | IPR011598 (Myc-type, basic helix-loop-helix (bHLH) domain) | GO:0046983 (protein dimerization activity) |
| Niben101Scf04639g03005.1 | 17 | 3 | -2.168 | 0.021210268 | sp\|B4U7U4\|FTSH_HYDS0 | ATP-dependent zinc metalloprotease FtsH | IPR003959 (ATPase, AAA-type, core), IPR009010 (Aspartate decarboxylase-like domain), IPR027417 (P-loop containing nucleoside triphosphate hydrolase), IPR029067 (CDC48 domain 2-like) | GO:0005524 (ATP binding) |
| Niben101Scf02852g02007.1 | 13 | 3 | -2.146 | 0.02583416 | sp\|P12062\|CB26_PETSP | Chlorophyll a-b binding protein 37, chloroplastic | IPR022796 (Chlorophyll A-B binding protein), IPR023329 (Chlorophyll a/b binding protein domain) | GO:0009765 (photosynthesis, light harvesting), GO:0016020 (membrane) |
| Niben101Scf04328g00045.1 | 10 | 2 | -2.143 | 0.029308011 | sp\|Q5IBL2\|YCF3_PLALA | Photosystem I assembly protein Ycf3 | NA | NA |
| Niben101Scf01072g01008.1 | 15 | 3 | -2.142 | 0.00429705 | AT1G78290.2 | Protein kinase superfamily protein LENGTH=343 | IPR011009 (Protein kinase-like domain) | GO:0004672 (protein kinase activity), GO:0004674 (protein serine/threonine kinase activity), GO:0005524 (ATP binding), GO:0006468 (protein phosphorylation), GO:0016772 (transferase activity, transferring phosphorus-containing groups) |
| Niben101Scf03152g00017.1 | 28 | 6 | -2.131 | 0.000287557 | sp\|Q7YJY8\|PSBA_CALFG | Photosystem Q(B) protein | IPR000484 (Photosynthetic reaction centre, L/M) | GO:0009772 (photosynthetic electron transport in photosystem II), GO:0019684 (photosynthesis, light reaction), GO:0045156 (electron transporter, transferring electrons within the cyclic electron transport pathway of photosynthesis activity) |
| Niben101Scf02576g00035.1 | 13 | 3 | -2.121 | 0.005058165 | AT5G08250.1 | Cytochrome P450 superfamily protein LENGTH=488 | IPR001128 (Cytochrome P450) | GO:0005506 (iron ion binding), GO:0016705 (oxidoreductase activity, acting on paired donors, with incorporation or reduction of molecular oxygen), GO:0020037 (heme binding), GO:0055114 (oxidation-reduction process) |
| Niben101Scf15579g00004.1 | 13 | 3 | -2.12 | 0.017824509 | emb\|CDY11325.1\| | BnaA03g01610D [Brassica napus] | NA | NA |
| Niben101Scf06462g01001.1 | 13 | 3 | -2.111 | 0.027459737 | sp\|Q56WM6\|ROGFE_ARATH | Rop guanine nucleotide exchange factor 14 | IPR005512 (PRONE domain) | GO:0005089 (Rho guanyl-nucleotide exchange factor activity) |
| Niben101Scf05025g10008.1 | 12 | 2 | -2.098 | 0.026321513 | ref\|XP_002519322.1\| | DNA binding protein, putative [Ricinus communis] gb\|EEF43186.1\| DNA binding protein, putative [Ricinus communis] | IPR003656 (Zinc finger, BED-type), IPR007021 (Domain of unknown function DUF659), IPR008906 (HAT dimerisation domain, C-terminal), IPR012337 (Ribonuclease H-like domain) | GO:0003676 (nucleic acid binding), GO:0003677 (DNA binding), GO:0046983 (protein dimerization activity) |
| Niben101Scf10489g00006.1 | 23 | 5 | -2.097 | 0.000100023 | AT3G17690.1 | cyclic nucleotide gated channel 19 LENGTH=729 | IPR014710 (RmlC-like jelly roll fold) | NA |
| Niben101Scf00775g05002.1 | 35 | 7 | -2.092 | 0.000211593 | sp\|Q9C8K2\|AB12G_ARATH | ABC transporter G family member 12 | IPR027417 (P-loop containing nucleoside triphosphate hydrolase) | NA |
| Niben101Scf13411g00007.1 | 11 | 2 | -2.085 | 0.027880722 | sp\|Q8GYP6\|PPR49_ARATH | Pentatricopeptide repeat-containing protein | IPR002885 (Pentatricopeptide repeat), IPR011990 (Tetratricopeptide-like helical domain) | GO:0005515 (protein binding) |
| Niben101Scf07247g01002.1 | 11 | 2 | -2.08 | 0.014180489 | AT5G10270.1 | cyclin-dependent kinase C;1 LENGTH=505 | IPR011009 (Protein kinase-like domain) | GO:0004672 (protein kinase activity), GO:0004674 (protein serine/threonine kinase activity), GO:0005524 (ATP binding), GO:0006468 (protein phosphorylation), GO:0016772 (transferase activity, transferring phosphorus-containing groups) |
| Niben101Scf14662g00007.1 | 16 | 3 | -2.075 | 0.003262411 | sp\|O14647\|CHD2_HUMAN | Chromodomain-helicase-DNA-binding protein 2 | IPR000330 (SNF2-related), IPR001650 (Helicase, C-terminal), IPR016197 (Chromo domain-like), IPR025260 (Domain of unknown function DUF4208), IPR027417 (P-loop containing nucleoside triphosphate hydrolase) | GO:0003677 (DNA binding), GO:0005524 (ATP binding) |
| Niben101Scf10224g01005.1 | 18 | 4 | -2.072 | 0.008213345 | sp\|Q8LPL6\|AP2A1_ARATH | AP-2 complex subunit alpha-1 | IPR009028 (Coatomer/calthrin adaptor appendage, C-terminal subdomain), IPR013041 (Coatomer/clathrin adaptor appendage, Ig-like subdomain), IPR016024 (Armadillo-type fold), IPR017104 (Adaptor protein complex AP-2, alpha subunit) | GO:0005488 (binding), GO:0006886 (intracellular protein transport), GO:0008565 (protein transporter activity), GO:0015031 (protein transport), GO:0016020 (membrane), GO:0016192 (vesicle-mediated transport), GO:0030117 (membrane coat), GO:0030131 (clathrin adaptor complex) |
| Niben101Scf05749g01009.1 | 11 | 2 | -2.068 | 0.033106689 | NA | Unknown protein | NA | NA |
| Niben101Scf02207g11009.1 | 103 | 22 | -2.063 | 1.31958E-11 | emb\|CAC86897.1\| | allene oxide synthase [Medicago truncatula] gb\|KEH40077.1\| cytochrome P450 family fatty acid hydroperoxide lyase [Medicago truncatula] | IPR001128 (Cytochrome P450) | GO:0004497 (monooxygenase activity), GO:0005506 (iron ion binding), GO:0016705 (oxidoreductase activity, acting on paired donors, with incorporation or reduction of molecular oxygen), GO:0020037 (heme binding), GO:0055114 (oxidation-reduction process) |
| Niben101Scf01378g00004.1 | 16 | 3 | -2.058 | 0.028353374 | sp\|O48837\|LRKS2_ARATH | Receptor like protein kinase S.2 | IPR011009 (Protein kinase-like domain) | GO:0004672 (protein kinase activity), GO:0004674 (protein serine/threonine kinase activity), GO:0005524 (ATP binding), GO:0006468 (protein phosphorylation), GO:0016772 (transferase activity, transferring phosphorus-containing groups) |
| Niben101Scf03370g07004.1 | 36 | 8 | -2.05 | 4.63476E-07 | sp\|Q76FS2\|TBB8_ORYSJ | Tubulin beta-8 chain | IPR000217 (Tubulin) | GO:0005200 (structural constituent of cytoskeleton), GO:0005525 (GTP binding), GO:0005874 (microtubule), GO:0007017 (microtubule-based process) |
| Niben101Scf03227g00001.1 | 20 | 4 | -2.043 | 0.005053075 | sp\|Q01780\|EXOSX_HUMAN | Exosome component 10 | IPR002121 (HRDC domain), IPR012337 (Ribonuclease H-like domain) | GO:0000166 (nucleotide binding), GO:0003676 (nucleic acid binding), GO:0003824 (catalytic activity), GO:0005622 (intracellular), GO:0006139 (nucleobase-containing compound metabolic process), GO:0008408 (3'-5' exonuclease activity), GO:0044237 (cellular metabolic process) |
| Niben101Scf06123g00002.1 | 14 | 3 | -2.041 | 0.038039812 | NA | Unknown protein | NA | NA |
| Niben101Scf08290g00012.1 | 17 | 4 | -2.038 | 0.039180596 | NA | Unknown protein | NA | NA |
| Niben101Scf11670g02004.1 | 14 | 3 | -2.033 | 0.008076741 | AT5G24470.1 | pseudo-response regulator 5 LENGTH=667 | IPR010402 (CCT domain), IPR011006 (CheY-like superfamily) | GO:0000160 (phosphorelay signal transduction system), GO:0005515 (protein binding) |
| Niben101Scf05371g06005.1 | 13 | 3 | -2.029 | 0.033917134 | sp\|Q8BHN5\|RBM45_MOUSE | RNA-binding protein 45 | IPR012677 (Nucleotide-binding alpha-beta plait domain) | GO:0000166 (nucleotide binding), GO:0003676 (nucleic acid binding) |
| Niben101Scf04914g00017.1 | 23 | 5 | -2.026 | 0.000204411 | AT1G29690.1 | MAC/Perforin domain-containing protein LENGTH=561 | IPR020864 (Membrane attack complex component/perforin (MACPF) domain) | NA |
| Niben101Scf04161g00003.1 | 15 | 3 | -2.022 | 0.007900009 | sp\|Q9LIG2\|RLK6_ARATH | Receptor-like protein kinase | IPR011009 (Protein kinase-like domain) | GO:0004672 (protein kinase activity), GO:0005524 (ATP binding), GO:0006468 (protein phosphorylation), GO:0016772 (transferase activity, transferring phosphorus-containing groups) |
| Niben101Scf03824g00016.1 | 18 | 4 | -2.02 | 0.004491115 | AT5G43630.1 | zinc knuckle (CCHC-type) family protein LENGTH=831 | IPR001878 (Zinc finger, CCHC-type), IPR004343 (Plus-3) | GO:0003676 (nucleic acid binding), GO:0003677 (DNA binding), GO:0005634 (nucleus), GO:0006352 (DNA-templated transcription, initiation), GO:0008270 (zinc ion binding), GO:0016570 (histone modification) |
| Niben101Scf16378g01012.1 | 19 | 4 | -2.02 | 0.014991313 | sp\|P46510\|CAHX_FLABI | Carbonic anhydrase | IPR001765 (Carbonic anhydrase) | GO:0004089 (carbonate dehydratase activity), GO:0008270 (zinc ion binding), GO:0015976 (carbon utilization) |
| Niben101Scf00506g02003.1 | 41 | 9 | -2.011 | 0.000131821 | sp\|P15904\|POR_AVESA | Protochlorophyllide reductase | IPR002347 (Glucose/ribitol dehydrogenase) | GO:0008152 (metabolic process), GO:0016491 (oxidoreductase activity) |
| Niben101Scf08679g02001.1 | 21 | 5 | -1.991 | 0.000595162 | sp\|Q8VYI9\|NLTL5_ARATH | Non-specific lipid-transfer protein-like protein | IPR016140 (Bifunctional inhibitor/plant lipid transfer protein/seed storage helical domain) | NA |
| Niben101Scf00644g00011.1 | 28 | 6 | -1.976 | 0.000264437 | sp\|E2RQ15\|RAB25_CANFA | Ras-related protein Rab-25 | IPR001806 (Small GTPase superfamily), IPR002041 (Ran GTPase), IPR005225 (Small GTP-binding protein domain), IPR024156 (Small GTPase superfamily, ARF type), IPR027417 (P-loop containing nucleoside triphosphate hydrolase) | GO:0003924 (GTPase activity), GO:0005525 (GTP binding), GO:0005622 (intracellular), GO:0006184 (GTP catabolic process), GO:0006886 (intracellular protein transport), GO:0006913 (nucleocytoplasmic transport), GO:0007165 (signal transduction), GO:0007264 (small GTPase mediated signal transduction), GO:0015031 (protein transport), GO:0016020 (membrane) |
| Niben101Scf06687g03002.1 | 16 | 4 | -1.966 | 0.002671194 | sp\|Q9H7Z3\|NRDE2_HUMAN | Protein NRDE2 homolog | IPR011990 (Tetratricopeptide-like helical domain), IPR013633 (siRNA-mediated silencing protein NRDE-2) | GO:0005515 (protein binding) |
| Niben101Scf00698g02020.1 | 16 | 4 | -1.958 | 0.030098874 | sp\|P19484\|TFEB_HUMAN | Transcription factor EB | IPR011598 (Myc-type, basic helix-loop-helix (bHLH) domain) | GO:0046983 (protein dimerization activity) |
| Niben101Scf05009g04012.1 | 26 | 6 | -1.956 | 0.0002778 | AT5G02810.1 | pseudo-response regulator 7 LENGTH=727 | IPR010402 (CCT domain), IPR011006 (CheY-like superfamily) | GO:0000160 (phosphorelay signal transduction system), GO:0005515 (protein binding) |
| Niben101Scf03772g02010.1 | 32 | 7 | -1.947 | 0.005288031 | AT4G17370.1 | Oxidoreductase family protein LENGTH=368 | IPR004104 (Oxidoreductase, C-terminal), IPR016040 (NAD(P)-binding domain) | GO:0008152 (metabolic process), GO:0016491 (oxidoreductase activity), GO:0055114 (oxidation-reduction process) |
| Niben101Scf15666g00009.1 | 32 | 7 | -1.946 | 0.000851273 | sp\|P74518\|LRTA_SYNY3 | Light-repressed protein A homolog | IPR003489 (Ribosomal protein S30Ae/sigma 54 modulation protein) | GO:0044238 (primary metabolic process) |
| Niben101Scf01326g10015.1 | 10 | 2 | -1.942 | 0.03985301 | gb\|KEH40157.1\| | FACT complex subunit SSRP1 [Medicago truncatula] | IPR000969 (Structure-specific recognition protein), IPR009071 (High mobility group box domain), IPR011993 (Pleckstrin homology-like domain), IPR013719 (Domain of unknown function DUF1747), IPR024954 (SSRP1 domain) | GO:0003677 (DNA binding), GO:0005634 (nucleus) |
| Niben101Scf00739g04026.1 | 16 | 4 | -1.938 | 0.012079656 | sp\|Q86JD4\|SEC11_DICDI | Signal peptidase complex catalytic subunit sec11 | IPR001733 (Peptidase S26B, eukaryotic signal peptidase), IPR015927 (Peptidase S24/S26A/S26B/S26C), IPR028360 (Peptidase S24/S26, beta-ribbon domain) | GO:0006465 (signal peptide processing), GO:0008233 (peptidase activity), GO:0016020 (membrane) |
| Niben101Scf01371g07014.1 | 12 | 3 | -1.927 | 0.028591545 | sp\|Q504L7\|ZN574_RAT | Zinc finger protein 574 | IPR013087 (Zinc finger C2H2-type/integrase DNA-binding domain) | GO:0003676 (nucleic acid binding), GO:0046872 (metal ion binding) |
| Niben101Scf07493g03007.1 | 16 | 4 | -1.926 | 0.048815013 | sp\|P0CB38\|PAB4L_HUMAN | Polyadenylate-binding protein 4-like | IPR006515 (Polyadenylate binding protein, human types 1, 2, 3, 4), IPR012677 (Nucleotide-binding alpha-beta plait domain) | GO:0000166 (nucleotide binding), GO:0003676 (nucleic acid binding), GO:0003723 (RNA binding) |
| Niben101Scf00777g00004.1 | 20 | 5 | -1.924 | 0.023357478 | AT2G25625.1 | unknown protein; FUNCTIONS IN: molecular_function unknown; INVOLVED IN: biological_process unknown; LOCATED IN: chloroplast; EXPRESSED IN: 10 plant structures; EXPRESSED DURING: LP.06 six leaves visible, LP.04 four leaves visible, 4 anthesis, petal differentiation and expansion stage; Has 24 Blast hits to 24 proteins in 9 species: Archae - 0; Bacteria - 0; Metazoa - 0; Fungi - 0; Plants - 24; Viruses - 0; Other Eukaryotes - 0 (source: NCBI BLink). LENGTH=152 | NA | NA |
| Niben101Scf11774g00003.1 | 21 | 5 | -1.919 | 0.000812181 | sp\|Q6UWW0\|LCN15_HUMAN | Lipocalin-15 | IPR011038 (Calycin-like) | NA |
| Niben101Scf08422g00003.1 | 45 | 11 | -1.914 | 7.11338E-07 | AT2G33830.2 | Dormancy/auxin associated family protein LENGTH=108 | IPR008406 (Dormancyauxin associated) | NA |
| Niben101Scf15817g00002.1 | 20 | 5 | -1.909 | 0.010607815 | sp\|P48699\|RBL_DENCL | Ribulose bisphosphate carboxylase large chain | IPR000685 (Ribulose bisphosphate carboxylase, large subunit, C-terminal) | GO:0000287 (magnesium ion binding), GO:0015977 (carbon fixation), GO:0016984 (ribulose-bisphosphate carboxylase activity) |
| Niben101Scf05124g01020.1 | 35 | 8 | -1.905 | 0.000167076 | sp\|P42046\|PSAC_CUCSA | Photosystem I iron-sulfur center | IPR001133 (NADH-ubiquinone oxidoreductase chain 4L/K), IPR017491 (Photosystem I protein PsaC) | GO:0009055 (electron carrier activity), GO:0009522 (photosystem I), GO:0009773 (photosynthetic electron transport in photosystem I), GO:0015979 (photosynthesis), GO:0016651 (oxidoreductase activity, acting on NAD(P)H), GO:0042651 (thylakoid membrane), GO:0042773 (ATP synthesis coupled electron transport), GO:0051536 (iron-sulfur cluster binding), GO:0051539 (4 iron, 4 sulfur cluster binding), GO:0055114 (oxidation-reduction process) |
| Niben101Scf03006g05001.1 | 16 | 4 | -1.903 | 0.007848302 | sp\|Q9FLP0\|MA651_ARATH | 65-kDa microtubule-associated protein 1 | IPR007145 (Microtubule-associated protein, MAP65/Ase1/PRC1) | GO:0000226 (microtubule cytoskeleton organization), GO:0000910 (cytokinesis), GO:0008017 (microtubule binding) |
| Niben101Scf02156g02008.1 | 13 | 3 | -1.894 | 0.037260617 | sp\|Q96300\|14337_ARATH | 14-3-3-like protein GF14 nu | IPR000308 (14-3-3 protein), IPR023409 (14-3-3 protein, conserved site), IPR023410 (14-3-3 domain) | GO:0019904 (protein domain specific binding) |
| Niben101Scf04713g03003.1 | 24 | 6 | -1.894 | 0.01052324 | AT4G17900.1 | PLATZ transcription factor family protein LENGTH=227 | IPR006734 (Protein of unknown function DUF597) | NA |
| Niben101Scf06349g00010.1 | 13 | 3 | -1.882 | 0.028946445 | sp\|P46824\|KLC_DROME | Kinesin light chain | IPR011990 (Tetratricopeptide-like helical domain) | GO:0005515 (protein binding) |
| Niben101Scf08160g05013.1 | 11 | 3 | -1.877 | 0.034816571 | sp\|P52674\|CYSJ_THIRO | Sulfite reductase [NADPH] flavoprotein alpha-component | IPR001094 (Flavodoxin), IPR007197 (Radical SAM), IPR013785 (Aldolase-type TIM barrel), IPR013917 (tRNA wybutosine-synthesis), IPR029039 (Flavoprotein-like) | GO:0003824 (catalytic activity), GO:0005506 (iron ion binding), GO:0010181 (FMN binding), GO:0016491 (oxidoreductase activity), GO:0051536 (iron-sulfur cluster binding), GO:0055114 (oxidation-reduction process) |
| Niben101Scf03150g07003.1 | 22 | 5 | -1.873 | 0.002783316 | AT3G46340.1 | Leucine-rich repeat protein kinase family protein LENGTH=889 | IPR001611 (Leucine-rich repeat), IPR024788 (Malectin-like carbohydrate-binding domain) | GO:0005515 (protein binding) |
| Niben101Scf00260g00003.1 | 19 | 5 | -1.869 | 0.01342984 | sp\|Q9SYT0\|ANXD1_ARATH | Annexin D1 | IPR001464 (Annexin) | GO:0005509 (calcium ion binding), GO:0005544 (calcium-dependent phospholipid binding) |
| Niben101Scf11444g00013.1 | 48 | 12 | -1.867 | 5.28842E-08 | AT3G02750.3 | Protein phosphatase 2C family protein LENGTH=527 | IPR001932 (Protein phosphatase 2C (PP2C)-like domain), IPR015655 (Protein phosphatase 2C) | GO:0003824 (catalytic activity) |
| Niben101Scf05467g00022.1 | 17 | 4 | -1.866 | 0.010753468 | gb\|AES99821.2\| | RNA polymerase II transcription factor SIII (elongin) subunit A [Medicago truncatula] | IPR010684 (RNA polymerase II transcription factor SIII, subunit A) | GO:0005634 (nucleus), GO:0006355 (regulation of transcription, DNA-templated), GO:0016021 (integral component of membrane) |
| Niben101Scf06236g01024.1 | 35 | 9 | -1.856 | 0.001073967 | sp\|Q84W04\|CNIH4_ARATH | Protein cornichon homolog 4 | IPR003377 (Cornichon) | GO:0016020 (membrane), GO:0035556 (intracellular signal transduction) |
| Niben101Ctg14374g00002.1 | 16 | 4 | -1.85 | 0.006119011 | AT4G19670.1 | RING/U-box superfamily protein LENGTH=532 | IPR013083 (Zinc finger, RING/FYVE/PHD-type) | GO:0005515 (protein binding), GO:0008270 (zinc ion binding) |
| Niben101Scf03722g00008.1 | 14 | 3 | -1.849 | 0.018148753 | AT2G37420.1 | ATP binding microtubule motor family protein LENGTH=1039 | IPR001752 (Kinesin motor domain), IPR027417 (P-loop containing nucleoside triphosphate hydrolase), IPR027640 (Kinesin-like protein) | GO:0003777 (microtubule motor activity), GO:0005524 (ATP binding), GO:0005871 (kinesin complex), GO:0007018 (microtubule-based movement), GO:0008017 (microtubule binding) |
| Niben101Scf02280g02004.1 | 25 | 6 | -1.839 | 0.002461631 | AT3G08510.1 | phospholipase C 2 LENGTH=581 | IPR001192 (Phosphoinositide phospholipase C family), IPR011992 (EF-hand domain pair) | GO:0004435 (phosphatidylinositol phospholipase C activity), GO:0005515 (protein binding), GO:0006629 (lipid metabolic process), GO:0007165 (signal transduction), GO:0008081 (phosphoric diester hydrolase activity), GO:0035556 (intracellular signal transduction) |
| Niben101Scf00185g01001.1 | 112 | 28 | -1.831 | 2.58853E-10 | emb\|CBY05509.1\| | ASR4 protein [Solanum peruvianum] | IPR003496 (ABA/WDS induced protein) | GO:0006950 (response to stress) |
| Niben101Scf00383g04013.1 | 612 | 151 | -1.83 | 0.002555882 | NA | Unknown protein | IPR000167 (Dehydrin) | GO:0006950 (response to stress), GO:0009415 (response to water) |
| Niben101Scf04216g07016.1 | 16 | 4 | -1.815 | 0.021629424 | sp\|A9IM54\|RL21_BART1 | 50S ribosomal protein L21 | IPR028909 (Ribosomal protein L21-like) | GO:0003723 (RNA binding), GO:0003735 (structural constituent of ribosome), GO:0005622 (intracellular), GO:0005840 (ribosome), GO:0006412 (translation) |
| Niben101Scf13558g01008.1 | 35 | 9 | -1.811 | 3.50099E-05 | AT5G20940.1 | Glycosyl hydrolase family protein LENGTH=626 | IPR002772 (Glycoside hydrolase family 3 C-terminal domain), IPR026892 (Glycoside hydrolase family 3) | GO:0004553 (hydrolase activity, hydrolyzing O-glycosyl compounds), GO:0005975 (carbohydrate metabolic process) |
| Niben101Scf07347g00001.1 | 12 | 3 | -1.809 | 0.02071283 | sp\|Q5F5V1\|RS4_NEIG1 | 30S ribosomal protein S4 | IPR002942 (RNA-binding S4 domain), IPR022801 (Ribosomal protein S4/S9) | GO:0003723 (RNA binding) |
| Niben101Scf05154g01013.1 | 33 | 8 | -1.806 | 0.006027459 | emb\|CDX92841.1\| | BnaC07g40990D [Brassica napus] | NA | NA |
| Niben101Scf01460g04028.1 | 11 | 3 | -1.803 | 0.048202571 | sp\|Q9QZA2\|PDC6I_RAT | Programmed cell death 6-interacting protein | IPR004328 (BRO1 domain), IPR025304 (ALIX V-shaped domain) | GO:0005515 (protein binding) |
| Niben101Scf04700g08003.1 | 17 | 4 | -1.799 | 0.010699243 | AT4G31590.1 | Cellulose-synthase-like C5 LENGTH=692 | IPR029044 (Nucleotide-diphospho-sugar transferases) | NA |
| Niben101Scf02937g00003.1 | 17 | 4 | -1.798 | 0.014135465 | sp\|Q9STV0\|GWD2_ARATH | Alpha-glucan water dikinase 2 | IPR002192 (Pyruvate phosphate dikinase, PEP/pyruvate-binding), IPR013816 (ATP-grasp fold, subdomain 2) | GO:0003824 (catalytic activity), GO:0005524 (ATP binding), GO:0016301 (kinase activity), GO:0016310 (phosphorylation) |
| Niben101Scf01740g07011.1 | 23 | 6 | -1.785 | 0.002466463 | AT5G53550.1 | YELLOW STRIPE like 3 LENGTH=675 | IPR004813 (Oligopeptide transporter, OPT superfamily) | GO:0055085 (transmembrane transport) |
| Niben101Scf04094g02006.1 | 13 | 3 | -1.778 | 0.019792908 | AT5G64740.1 | cellulose synthase 6 LENGTH=1084 | IPR005150 (Cellulose synthase), IPR013083 (Zinc finger, RING/FYVE/PHD-type), IPR029044 (Nucleotide-diphospho-sugar transferases) | GO:0005515 (protein binding), GO:0008270 (zinc ion binding), GO:0016020 (membrane), GO:0016760 (cellulose synthase (UDP-forming) activity), GO:0030244 (cellulose biosynthetic process) |
| Niben101Scf08533g03005.1 | 22 | 6 | -1.778 | 0.00716798 | sp\|Q9Y618\|NCOR2_HUMAN | Nuclear receptor corepressor 2 | IPR009057 (Homeodomain-like) | GO:0003677 (DNA binding), GO:0003682 (chromatin binding) |
| Niben101Scf10989g00004.1 | 12 | 3 | -1.774 | 0.035127249 | sp\|P92942\|CLCB_ARATH | Chloride channel protein CLC-b | IPR000644 (CBS domain), IPR014743 (Chloride channel, core) | GO:0005216 (ion channel activity), GO:0055085 (transmembrane transport) |
| Niben101Scf00175g02019.1 | 14 | 4 | -1.772 | 0.029151641 | AT3G21760.1 | UDP-Glycosyltransferase superfamily protein LENGTH=485 | IPR002213 (UDP-glucuronosyl/UDP-glucosyltransferase) | GO:0008152 (metabolic process), GO:0016758 (transferase activity, transferring hexosyl groups) |
| Niben101Scf00868g07005.1 | 24 | 6 | -1.771 | 0.000945574 | sp\|Q9LIG2\|RLK6_ARATH | Receptor-like protein kinase | IPR011009 (Protein kinase-like domain) | GO:0004672 (protein kinase activity), GO:0004674 (protein serine/threonine kinase activity), GO:0005524 (ATP binding), GO:0006468 (protein phosphorylation), GO:0016772 (transferase activity, transferring phosphorus-containing groups) |
| Niben101Scf00650g02002.1 | 33 | 9 | -1.771 | 0.000415211 | gb\|AAT39951.2\| | Disease resistance protein, putative [Solanum demissum] | IPR002182 (NB-ARC), IPR021929 (Late blight resistance protein R1) | GO:0043531 (ADP binding) |
| Niben101Ctg13092g00001.1 | 12 | 3 | -1.767 | 0.026524883 | emb\|CAC83580.1\| | major latex-like protein [Arabidopsis thaliana] | IPR000916 (Bet v I domain), IPR023393 (START-like domain) | GO:0006952 (defense response), GO:0009607 (response to biotic stimulus) |
| Niben101Scf03138g01018.1 | 18 | 5 | -1.767 | 0.005640986 | gb\|AAT77883.1\| | expressed protein [Oryza sativa Japonica Group] gb\|ABF98697.1\| expressed protein [Oryza sativa Japonica Group] gb\|EAZ28473.1\| hypothetical protein OsJ_12454 [Oryza sativa Japonica Group] | IPR008796 (Photosystem I PsaN, reaction centre subunit N) | GO:0005516 (calmodulin binding), GO:0009522 (photosystem I), GO:0015979 (photosynthesis), GO:0042651 (thylakoid membrane) |
| Niben101Scf02108g02018.1 | 29 | 8 | -1.764 | 0.009647661 | AT1G73460.1 | Protein kinase superfamily protein LENGTH=1169 | IPR011009 (Protein kinase-like domain) | GO:0004672 (protein kinase activity), GO:0004674 (protein serine/threonine kinase activity), GO:0005524 (ATP binding), GO:0006468 (protein phosphorylation), GO:0016772 (transferase activity, transferring phosphorus-containing groups) |
| Niben101Scf01814g05011.1 | 25 | 7 | -1.755 | 0.002834005 | AT3G50590.1 | Transducin/WD40 repeat-like superfamily protein LENGTH=1614 | IPR015943 (WD40/YVTN repeat-like-containing domain) | GO:0005515 (protein binding) |
| Niben101Scf00173g09004.1 | 15 | 4 | -1.742 | 0.047153697 | sp\|A4GG89\|RBL_PHAVU | Ribulose bisphosphate carboxylase large chain | IPR020888 (Ribulose bisphosphate carboxylase, large subunit) | GO:0000287 (magnesium ion binding), GO:0015977 (carbon fixation), GO:0016984 (ribulose-bisphosphate carboxylase activity) |
| Niben101Scf09454g00008.1 | 43 | 12 | -1.742 | 0.000310311 | sp\|P48000\|KNAT3_ARATH | Homeobox protein knotted-1-like 3 | IPR005540 (KNOX1), IPR005541 (KNOX2), IPR009057 (Homeodomain-like) | GO:0003677 (DNA binding), GO:0005634 (nucleus), GO:0006355 (regulation of transcription, DNA-templated) |
| Niben101Scf11444g00013.1 | 14 | 4 | -1.737 | 0.032715916 | AT3G02750.3 | Protein phosphatase 2C family protein LENGTH=527 | IPR001932 (Protein phosphatase 2C (PP2C)-like domain), IPR015655 (Protein phosphatase 2C) | GO:0003824 (catalytic activity) |
| Niben101Scf00228g06003.1 | 19 | 5 | -1.736 | 0.027040915 | sp\|Q9SRT0\|PUB9_ARATH | U-box domain-containing protein 9 | IPR013083 (Zinc finger, RING/FYVE/PHD-type), IPR016024 (Armadillo-type fold) | GO:0004842 (ubiquitin-protein transferase activity), GO:0005488 (binding), GO:0005515 (protein binding), GO:0016567 (protein ubiquitination) |
| Niben101Scf02182g13008.1 | 38 | 10 | -1.736 | 0.000105126 | gb\|ABN08421.1\| | nucleic acid binding , related [Medicago truncatula] | IPR004087 (K Homology domain) | GO:0003676 (nucleic acid binding), GO:0003723 (RNA binding) |
| Niben101Scf01020g06001.1 | 57 | 15 | -1.735 | 0.00035957 | sp\|B0C431\|RS11_ACAM1 | 30S ribosomal protein S11 | IPR001971 (Ribosomal protein S11) | GO:0003735 (structural constituent of ribosome), GO:0005840 (ribosome), GO:0006412 (translation) |
| Niben101Scf01631g01001.1 | 18 | 5 | -1.73 | 0.021846818 | NA | Unknown protein | NA | NA |
| Niben101Scf02242g03013.1 | 14 | 4 | -1.728 | 0.031834966 | sp\|Q9FIB6\|PS12A_ARATH | 26S proteasome non-ATPase regulatory subunit 12 homolog A | IPR000717 (Proteasome component (PCI) domain) | GO:0005515 (protein binding) |
| Niben101Scf02207g11009.1 | 62 | 17 | -1.728 | 6.88008E-05 | emb\|CAC86897.1\| | allene oxide synthase [Medicago truncatula] gb\|KEH40077.1\| cytochrome P450 family fatty acid hydroperoxide lyase [Medicago truncatula] | IPR001128 (Cytochrome P450) | GO:0004497 (monooxygenase activity), GO:0005506 (iron ion binding), GO:0016705 (oxidoreductase activity, acting on paired donors, with incorporation or reduction of molecular oxygen), GO:0020037 (heme binding), GO:0055114 (oxidation-reduction process) |
| Niben101Scf09459g00002.1 | 22 | 6 | -1.721 | 0.011273286 | sp\|P0A3V1\|SLEB_BACCE | Spore cortex-lytic enzyme | IPR001305 (Heat shock protein DnaJ, cysteine-rich domain), IPR002477 (Peptidoglycan binding-like) | GO:0031072 (heat shock protein binding), GO:0051082 (unfolded protein binding) |
| Niben101Scf02937g00003.1 | 21 | 6 | -1.717 | 0.040338499 | sp\|Q9STV0\|GWD2_ARATH | Alpha-glucan water dikinase 2 | IPR002192 (Pyruvate phosphate dikinase, PEP/pyruvate-binding), IPR013816 (ATP-grasp fold, subdomain 2) | GO:0003824 (catalytic activity), GO:0005524 (ATP binding), GO:0016301 (kinase activity), GO:0016310 (phosphorylation) |
| Niben101Scf06964g00007.1 | 29 | 8 | -1.714 | 0.002485114 | sp\|Q7YJX7\|PSBD_CALFG | Photosystem II D2 protein | IPR000484 (Photosynthetic reaction centre, L/M), IPR000932 (Photosystem antenna protein-like) | GO:0009521 (photosystem), GO:0009523 (photosystem II), GO:0009767 (photosynthetic electron transport chain), GO:0009772 (photosynthetic electron transport in photosystem II), GO:0015979 (photosynthesis), GO:0016020 (membrane), GO:0016168 (chlorophyll binding), GO:0019684 (photosynthesis, light reaction), GO:0045156 (electron transporter, transferring electrons within the cyclic electron transport pathway of photosynthesis activity) |
| Niben101Scf05855g06010.1 | 14 | 4 | -1.709 | 0.024134597 | sp\|B2S9C2\|DNAJ_BRUA1 | Chaperone protein DnaJ | IPR008971 (HSP40/DnaJ peptide-binding) | GO:0006457 (protein folding), GO:0051082 (unfolded protein binding) |
| Niben101Scf04276g04022.1 | 182 | 50 | -1.708 | 8.37297E-12 | sp\|Q823T2\|DNAJ_CHLCV | Chaperone protein DnaJ | IPR001623 (DnaJ domain) | NA |
| Niben101Scf03330g00013.1 | 17 | 5 | -1.706 | 0.032881739 | AT2G04540.1 | Beta-ketoacyl synthase LENGTH=461 | IPR017568 (3-oxoacyl-[acyl-carrier-protein] synthase 2), IPR020841 (Polyketide synthase, beta-ketoacyl synthase domain) | GO:0003824 (catalytic activity), GO:0006633 (fatty acid biosynthetic process), GO:0008152 (metabolic process), GO:0016747 (transferase activity, transferring acyl groups other than amino-acyl groups) |
| Niben101Scf08929g00003.1 | 38 | 10 | -1.705 | 5.27031E-05 | ref\|NP_001274932.1\| | zinc finger protein NUTCRACKER-like [Solanum tuber [Solanum tuberosum] | NA | NA |
| Niben101Scf03307g04014.1 | 16 | 4 | -1.699 | 0.039267785 | sp\|P27061\|PPA1_SOLLC | Acid phosphatase 1 | IPR005519 (Acid phosphatase (Class B)), IPR023214 (HAD-like domain) | GO:0003993 (acid phosphatase activity) |
| Niben101Scf00750g02005.1 | 236 | 66 | -1.694 | 0.038665536 | sp\|Q00783\|ICI1_SOLTU | Proteinase inhibitor 1 | IPR000864 (Proteinase inhibitor I13, potato inhibitor I) | GO:0004867 (serine-type endopeptidase inhibitor activity), GO:0009611 (response to wounding) |
| Niben101Scf00495g02014.1 | 12 | 3 | -1.691 | 0.042625563 | gb\|KEH21698.1\| | NLI interacting factor-like phosphatase [Medicago truncatula] | IPR004274 (NLI interacting factor), IPR023214 (HAD-like domain) | GO:0005515 (protein binding) |
| Niben101Scf02078g05004.1 | 2881 | 795 | -1.689 | 5.79868E-10 | sp\|A1EA06\|RR14_AGRST | 30S ribosomal protein S14, chloroplastic | IPR001209 (Ribosomal protein S14) | GO:0003735 (structural constituent of ribosome), GO:0005622 (intracellular), GO:0005840 (ribosome), GO:0006412 (translation) |
| Niben101Scf04804g01002.1 | 20 | 6 | -1.67 | 0.026506074 | sp\|P93788\|REMO_SOLTU | Remorin | IPR005516 (Remorin, C-terminal), IPR005518 (Remorin, N-terminal) | NA |
| Niben101Scf03832g02016.1 | 14 | 4 | -1.659 | 0.040531062 | AT4G32390.1 | Nucleotide-sugar transporter family protein LENGTH=350 | IPR004853 (Triose-phosphate transporter domain) | NA |
| Niben101Scf06369g04008.1 | 14 | 4 | -1.657 | 0.033734963 | AT4G23100.1 | glutamate-cysteine ligase LENGTH=522 | IPR006336 (Glutamate--cysteine ligase, GCS2) | GO:0004357 (glutamate-cysteine ligase activity), GO:0006750 (glutathione biosynthetic process), GO:0042398 (cellular modified amino acid biosynthetic process) |
| Niben101Scf00634g00003.1 | 18 | 5 | -1.657 | 0.048362566 | sp\|A4QLH5\|PSBA_LOBMA | Photosystem Q(B) protein | IPR000484 (Photosynthetic reaction centre, L/M) | GO:0009772 (photosynthetic electron transport in photosystem II), GO:0019684 (photosynthesis, light reaction), GO:0045156 (electron transporter, transferring electrons within the cyclic electron transport pathway of photosynthesis activity) |
| Niben101Scf01071g03004.1 | 44 | 13 | -1.652 | 0.000162923 | AT5G64740.1 | cellulose synthase 6 LENGTH=1084 | IPR005150 (Cellulose synthase), IPR013083 (Zinc finger, RING/FYVE/PHD-type), IPR029044 (Nucleotide-diphospho-sugar transferases) | GO:0005515 (protein binding), GO:0008270 (zinc ion binding), GO:0016020 (membrane), GO:0016760 (cellulose synthase (UDP-forming) activity), GO:0030244 (cellulose biosynthetic process) |
| Niben101Scf07242g07006.1 | 97 | 28 | -1.65 | 1.21287E-07 | AT3G28740.1 | Cytochrome P450 superfamily protein LENGTH=509 | IPR001128 (Cytochrome P450) | GO:0005506 (iron ion binding), GO:0016705 (oxidoreductase activity, acting on paired donors, with incorporation or reduction of molecular oxygen), GO:0020037 (heme binding), GO:0055114 (oxidation-reduction process) |
| Niben101Scf01710g01013.1 | 15 | 4 | -1.646 | 0.028434722 | AT2G42590.3 | general regulatory factor 9 LENGTH=276 | IPR000308 (14-3-3 protein), IPR023409 (14-3-3 protein, conserved site), IPR023410 (14-3-3 domain) | GO:0019904 (protein domain specific binding) |
| Niben101Scf01817g00001.1 | 15 | 4 | -1.635 | 0.032757289 | ref\|XP_007040444.1\| | Ubiquitin ligase protein cop1, putative isoform 5 [Theobroma cacao] gb\|EOY24945.1\| Ubiquitin ligase protein cop1, putative isoform 5 [Theobroma cacao] | IPR011009 (Protein kinase-like domain) | GO:0016772 (transferase activity, transferring phosphorus-containing groups) |
| Niben101Scf15568g00037.1 | 20 | 6 | -1.634 | 0.036001425 | sp\|P93788\|REMO_SOLTU | Remorin | IPR005516 (Remorin, C-terminal), IPR005518 (Remorin, N-terminal) | NA |
| Niben101Scf01036g03001.1 | 45 | 13 | -1.633 | 3.10179E-05 | sp\|P15904\|POR_AVESA | Protochlorophyllide reductase | IPR002347 (Glucose/ribitol dehydrogenase) | GO:0008152 (metabolic process), GO:0016491 (oxidoreductase activity), GO:0016630 (protochlorophyllide reductase activity), GO:0055114 (oxidation-reduction process) |
| Niben101Scf03444g04001.1 | 14 | 4 | -1.632 | 0.028353374 | sp\|P57758\|CTNS_ARATH | Cystinosin homolog | IPR005282 (Lysosomal cystine transporter) | NA |
| Niben101Scf06183g03009.1 | 19 | 5 | -1.632 | 0.027899124 | AT2G42590.3 | general regulatory factor 9 LENGTH=276 | IPR000308 (14-3-3 protein), IPR023409 (14-3-3 protein, conserved site), IPR023410 (14-3-3 domain) | GO:0019904 (protein domain specific binding) |
| Niben101Scf02015g01007.1 | 26 | 7 | -1.632 | 0.002997636 | sp\|Q9LEC8\|DHEB_NICPL | Glutamate dehydrogenase B | IPR006095 (Glutamate/phenylalanine/leucine/valine dehydrogenase) | GO:0006520 (cellular amino acid metabolic process), GO:0016491 (oxidoreductase activity), GO:0055114 (oxidation-reduction process) |
| Niben101Scf09436g01003.1 | 20 | 6 | -1.628 | 0.032009867 | AT2G31510.1 | IBR domain-containing protein LENGTH=562 | IPR002867 (Zinc finger, C6HC-type), IPR013083 (Zinc finger, RING/FYVE/PHD-type) | GO:0005515 (protein binding), GO:0008270 (zinc ion binding), GO:0046872 (metal ion binding) |
| Niben101Scf23606g00007.1 | 60 | 17 | -1.626 | 0.000286692 | sp\|C5BDY7\|MDTK_EDWI9 | Multidrug resistance protein MdtK | IPR002528 (Multi antimicrobial extrusion protein) | GO:0006855 (drug transmembrane transport), GO:0015238 (drug transmembrane transporter activity), GO:0015297 (antiporter activity), GO:0016020 (membrane), GO:0055085 (transmembrane transport) |
| Niben101Scf19057g00004.1 | 64 | 19 | -1.619 | 0.024270484 | sp\|Q39002\|TLC1_ARATH | ADP,ATP carrier protein 1, chloroplastic | NA | NA |
| Niben101Scf01328g01012.1 | 25 | 7 | -1.616 | 0.002792978 | sp\|P27522\|CB13_SOLLC | Chlorophyll a-b binding protein 8, chloroplastic | IPR022796 (Chlorophyll A-B binding protein), IPR023329 (Chlorophyll a/b binding protein domain) | GO:0009765 (photosynthesis, light harvesting), GO:0016020 (membrane) |
| Niben101Scf09107g01004.1 | 43 | 13 | -1.615 | 0.001535759 | sp\|Q9LIB2\|PHS1_ARATH | Alpha-glucan phosphorylase 1 | IPR000811 (Glycosyl transferase, family 35) | GO:0005975 (carbohydrate metabolic process), GO:0008184 (glycogen phosphorylase activity) |
| Niben101Scf02026g04007.1 | 88 | 26 | -1.614 | 7.5691E-05 | AT2G25180.1 | response regulator 12 LENGTH=596 | IPR010402 (CCT domain), IPR011006 (CheY-like superfamily) | GO:0000160 (phosphorelay signal transduction system), GO:0005515 (protein binding) |
| Niben101Scf04422g10004.1 | 48 | 14 | -1.608 | 8.49798E-06 | emb\|CDX78779.1\| | BnaA03g06910D [Brassica napus] | NA | NA |
| Niben101Scf01994g02012.1 | 21 | 6 | -1.601 | 0.030875757 | sp\|Q9XF97\|RL4_PRUAR | 60S ribosomal protein L4 | IPR002136 (Ribosomal protein L4/L1e), IPR023574 (Ribosomal protein L4 domain), IPR025755 (60S ribosomal protein L4, C-terminal domain) | GO:0003735 (structural constituent of ribosome), GO:0005840 (ribosome), GO:0006412 (translation) |
| Niben101Scf09590g03007.1 | 565 | 169 | -1.596 | 4.53865E-09 | sp\|Q08507\|ACCO3_PETHY | 1-aminocyclopropane-1-carboxylate oxidase 3 | IPR026992 (Non-haem dioxygenase N-terminal domain), IPR027443 (Isopenicillin N synthase-like) | NA |
| Niben101Scf08041g01021.1 | 29 | 9 | -1.592 | 0.004847971 | ref\|XP_007042708.1\| | NAD(P)-binding R isoform 1 [Theobroma cacao] | IPR016040 (NAD(P)-binding domain) | NA |
| Niben101Scf02942g00020.1 | 35 | 10 | -1.591 | 0.001116979 | emb\|CDY44410.1\| | BnaA02g29670D [Brassica napus] | NA | NA |
| Niben101Scf00887g02006.1 | 45 | 14 | -1.591 | 0.004143359 | sp\|P50134\|DCOR_DATST | Ornithine decarboxylase | IPR000183 (Ornithine/DAP/Arg decarboxylase), IPR029066 (PLP-binding barrel) | GO:0003824 (catalytic activity), GO:0006596 (polyamine biosynthetic process) |
| Niben101Scf06088g00005.1 | 20 | 6 | -1.584 | 0.038937693 | ref\|XP_002525522.1\| | nucleic acid binding protein, putative [Ricinus communis] gb\|EEF36880.1\| nucleic acid binding protein, putative [Ricinus communis] | IPR004087 (K Homology domain) | GO:0003676 (nucleic acid binding), GO:0003723 (RNA binding) |
| Niben101Scf09725g00020.1 | 22 | 7 | -1.583 | 0.005837906 | ref\|XP_007030054.1\| | Nucleic acid binding protein, putative isoform 2 [Theobroma cacao] gb\|EOY10556.1\| Nucleic acid binding protein, putative isoform 2 [Theobroma cacao] | IPR012677 (Nucleotide-binding alpha-beta plait domain) | GO:0000166 (nucleotide binding), GO:0003676 (nucleic acid binding) |
| Niben101Scf06923g00002.1 | 35 | 10 | -1.58 | 0.000982481 | AT3G60390.1 | homeobox-leucine zipper protein 3 LENGTH=315 | IPR003106 (Leucine zipper, homeobox-associated), IPR006712 (HD-ZIP protein, N-terminal), IPR009057 (Homeodomain-like) | GO:0003677 (DNA binding), GO:0003700 (sequence-specific DNA binding transcription factor activity), GO:0005634 (nucleus), GO:0006355 (regulation of transcription, DNA-templated), GO:0043565 (sequence-specific DNA binding) |
| Niben101Scf00077g05011.1 | 18 | 5 | -1.579 | 0.030074505 | sp\|Q9CR16\|PPID_MOUSE | Peptidyl-prolyl cis-trans isomerase D | IPR001179 (Peptidyl-prolyl cis-trans isomerase, FKBP-type, domain), IPR011990 (Tetratricopeptide-like helical domain), IPR023566 (Peptidyl-prolyl cis-trans isomerase, FKBP-type) | GO:0005515 (protein binding), GO:0006457 (protein folding) |
| Niben101Scf01718g01010.1 | 48 | 14 | -1.578 | 0.000457361 | AT2G37180.1 | Aquaporin-like superfamily protein LENGTH=285 | IPR000425 (Major intrinsic protein), IPR023271 (Aquaporin-like) | GO:0005215 (transporter activity), GO:0006810 (transport), GO:0016020 (membrane) |
| Niben101Scf04641g03006.1 | 65 | 20 | -1.576 | 2.99628E-06 | sp\|Q9SQI2\|GIGAN_ARATH | Protein GIGANTEA | IPR026211 (GIGANTEA) | GO:2000028 (regulation of photoperiodism, flowering) |
| Niben101Scf03251g00008.1 | 16 | 5 | -1.571 | 0.030475301 | sp\|Q40831\|TBA1_PELFA | Tubulin alpha-1 chain | IPR000217 (Tubulin), IPR023123 (Tubulin, C-terminal) | GO:0003924 (GTPase activity), GO:0005200 (structural constituent of cytoskeleton), GO:0005525 (GTP binding), GO:0005874 (microtubule), GO:0006184 (GTP catabolic process), GO:0007017 (microtubule-based process), GO:0043234 (protein complex), GO:0051258 (protein polymerization) |
| Niben101Scf12383g01002.1 | 39 | 12 | -1.569 | 0.000329616 | sp\|P49756\|RBM25_HUMAN | RNA-binding protein 25 | IPR002483 (PWI domain), IPR012677 (Nucleotide-binding alpha-beta plait domain) | GO:0000166 (nucleotide binding), GO:0003676 (nucleic acid binding), GO:0006397 (mRNA processing) |
| Niben101Scf03365g02002.1 | 15 | 4 | -1.567 | 0.044955238 | sp\|Q4KLU2\|PRP39_XENLA | Pre-mRNA-processing factor 39 | IPR011990 (Tetratricopeptide-like helical domain) | GO:0005515 (protein binding), GO:0005622 (intracellular), GO:0006396 (RNA processing) |
| Niben101Scf02182g13008.1 | 23 | 7 | -1.567 | 0.003383588 | gb\|ABN08421.1\| | nucleic acid binding , related [Medicago truncatula] | IPR004087 (K Homology domain) | GO:0003676 (nucleic acid binding), GO:0003723 (RNA binding) |
| Niben101Scf06848g08022.1 | 27 | 8 | -1.567 | 0.003004202 | sp\|Q58D06\|WDR74_BOVIN | WD repeat-containing protein 74 | IPR011047 (Quinonprotein alcohol dehydrogenase-like superfamily), IPR015943 (WD40/YVTN repeat-like-containing domain) | GO:0005515 (protein binding) |
| Niben101Scf00671g00011.1 | 59 | 18 | -1.558 | 2.55128E-06 | sp\|P93047\|HMGB3_ARATH | High mobility group B protein 3 | IPR009071 (High mobility group box domain) | NA |
| Niben101Scf00635g09016.1 | 21 | 6 | -1.555 | 0.006697543 | sp\|Q9M069\|E137_ARATH | Glucan endo-1,3-beta-glucosidase 7 | IPR012946 (X8 domain) | NA |
| Niben101Scf23111g00012.1 | 474 | 146 | -1.554 | 2.89853E-18 | gb\|AIT42219.1\| | metallocarboxypeptidase inhibitor [Solanum tuberosum] | IPR004231 (Carboxypeptidase A inhibitor-like) | NA |
| Niben101Scf03621g01017.1 | 27 | 8 | -1.55 | 0.000812181 | sp\|Q8L7S4\|MP702_ARATH | Microtubule-associated protein 70-2 | IPR009768 (Microtubule-associated protein 70) | GO:0007010 (cytoskeleton organization), GO:0008017 (microtubule binding) |
| Niben101Scf15022g00007.1 | 14 | 4 | -1.548 | 0.034022357 | sp\|Q8W4S4\|VHAA3_ARATH | V-type proton ATPase subunit a3 | IPR002490 (V-type ATPase, V0 complex, 116kDa subunit family) | GO:0000220 (vacuolar proton-transporting V-type ATPase, V0 domain), GO:0015078 (hydrogen ion transmembrane transporter activity), GO:0015991 (ATP hydrolysis coupled proton transport), GO:0033179 (proton-transporting V-type ATPase, V0 domain) |
| Niben101Scf09115g00014.1 | 89 | 27 | -1.547 | 2.16463E-08 | AT3G30390.1 | Transmembrane amino acid transporter family protein LENGTH=460 | IPR013057 (Amino acid transporter, transmembrane) | NA |
| Niben101Scf05300g01001.1 | 20 | 6 | -1.538 | 0.011781521 | sp\|Q54UC9\|KIF3_DICDI | Kinesin-related protein 3 | IPR001752 (Kinesin motor domain), IPR027417 (P-loop containing nucleoside triphosphate hydrolase), IPR027640 (Kinesin-like protein) | GO:0003777 (microtubule motor activity), GO:0005524 (ATP binding), GO:0005871 (kinesin complex), GO:0007018 (microtubule-based movement), GO:0008017 (microtubule binding) |
| Niben101Scf03449g02015.1 | 28 | 9 | -1.536 | 0.002686429 | sp\|Q9DC07\|LNEBL_MOUSE | LIM zinc-binding domain-containing Nebulette | IPR001781 (Zinc finger, LIM-type) | GO:0008270 (zinc ion binding) |
| Niben101Scf03224g06001.1 | 27 | 8 | -1.531 | 0.007707323 | AT1G15910.1 | XH/XS domain-containing protein LENGTH=634 | IPR005379 (Uncharacterised domain XH), IPR005380 (XS domain), IPR005381 (Zinc finger-XS domain) | GO:0031047 (gene silencing by RNA) |
| Niben101Scf02009g00007.1 | 17 | 5 | -1.527 | 0.045615299 | sp\|C5D6U5\|MENE_GEOSW | 2-succinylbenzoate--CoA ligase | IPR000873 (AMP-dependent synthetase/ligase), IPR025110 (AMP-binding enzyme C-terminal domain) | GO:0003824 (catalytic activity), GO:0008152 (metabolic process) |
| Niben101Scf02594g01007.1 | 26 | 8 | -1.52 | 0.006494171 | sp\|Q9FRL3\|ERDL6_ARATH | Sugar transporter ERD6-like 6 | IPR005828 (General substrate transporter), IPR020846 (Major facilitator superfamily domain) | GO:0005215 (transporter activity), GO:0016021 (integral component of membrane), GO:0022857 (transmembrane transporter activity), GO:0055085 (transmembrane transport) |
| Niben101Scf00324g00004.1 | 27 | 8 | -1.516 | 0.016227779 | ref\|WP_025921648.1\| | MULTISPECIES: hypothetical protein [Prochlorococcus] | IPR021659 (Protein of unknown function DUF3252) | NA |
| Niben101Scf06112g00003.1 | 22 | 7 | -1.515 | 0.019757557 | gb\|ADK97920.1\| | LuxR family transcriptional regulator [Schiedea globosa] | NA | NA |
| Niben101Scf11795g00014.1 | 24 | 7 | -1.512 | 0.017195061 | sp\|Q9FN94\|RLK7_ARATH | Receptor-like protein kinase | IPR011009 (Protein kinase-like domain), IPR013320 (Concanavalin A-like lectin/glucanase domain) | GO:0004672 (protein kinase activity), GO:0004674 (protein serine/threonine kinase activity), GO:0005524 (ATP binding), GO:0006468 (protein phosphorylation), GO:0016772 (transferase activity, transferring phosphorus-containing groups) |
| Niben101Scf02123g02006.1 | 16 | 5 | -1.501 | 0.031099847 | sp\|Q7F8R0\|C3H14_ORYSJ | Zinc finger CCCH domain-containing protein 14 | IPR000571 (Zinc finger, CCCH-type), IPR004087 (K Homology domain) | GO:0003676 (nucleic acid binding), GO:0003723 (RNA binding), GO:0046872 (metal ion binding) |
| Niben101Scf02824g03017.1 | 33 | 11 | -1.5 | 0.010527456 | AT2G04030.1 | Chaperone protein htpG family protein LENGTH=780 | IPR001404 (Heat shock protein Hsp90 family) | GO:0005524 (ATP binding), GO:0006457 (protein folding), GO:0006950 (response to stress), GO:0051082 (unfolded protein binding) |
| Niben101Scf05732g03012.1 | 38 | 12 | -1.493 | 0.000897539 | AT5G24690.1 | Protein of unknown function (DUF3411) LENGTH=521 | IPR021825 (Protein of unknown function DUF3411, plant) | NA |
| Niben101Scf02455g03033.1 | 59 | 19 | -1.488 | 0.000230728 | AT1G80530.1 | Major facilitator superfamily protein LENGTH=561 | IPR010658 (Nodulin-like), IPR020846 (Major facilitator superfamily domain) | NA |
| Niben101Scf04964g00002.1 | 142 | 45 | -1.487 | 2.97497E-08 | sp\|P51193\|PSAF_PORPU | Photosystem I reaction center subunit III | IPR003666 (Photosystem I PsaF, reaction centre subunit III) | GO:0009522 (photosystem I), GO:0009538 (photosystem I reaction center), GO:0015979 (photosynthesis) |
| Niben101Scf07383g07006.1 | 31 | 10 | -1.485 | 0.004120102 | emb\|CDY19080.1\| | BnaA04g18200D [Brassica napus] | IPR012881 (Protein of unknown function DUF1685) | NA |
| Niben101Scf05298g06005.1 | 822 | 265 | -1.476 | 0.027479665 | AT1G56220.3 | Dormancy/auxin associated family protein LENGTH=140 | IPR008406 (Dormancyauxin associated) | NA |
| Niben101Scf11162g00002.1 | 23 | 7 | -1.475 | 0.044392545 | AT1G68910.1 | WPP domain-interacting protein 2 LENGTH=627 | NA | NA |
| Niben101Scf01962g01006.1 | 27 | 9 | -1.474 | 0.048253822 | sp\|Q948T6\|LGUL_ORYSJ | Lactoylglutathione lyase | IPR004361 (Glyoxalase I), IPR029068 (Glyoxalase/Bleomycin resistance protein/Dihydroxybiphenyl dioxygenase) | GO:0004462 (lactoylglutathione lyase activity), GO:0046872 (metal ion binding) |
| Niben101Scf08108g00003.1 | 39 | 13 | -1.474 | 0.000122964 | AT2G29670.1 | Tetratricopeptide repeat (TPR)-like superfamily protein LENGTH=536 | IPR011990 (Tetratricopeptide-like helical domain) | GO:0005515 (protein binding) |
| Niben101Scf09997g00025.1 | 37 | 12 | -1.471 | 0.000145798 | AT3G24630.1 | unknown protein; Has 5348 Blast hits to 3182 proteins in 353 species: Archae - 0; Bacteria - 481; Metazoa - 1959; Fungi - 405; Plants - 180; Viruses - 10; Other Eukaryotes - 2313 (source: NCBI BLink). LENGTH=724 | IPR025486 (Domain of unknown function DUF4378) | NA |
| Niben101Scf00616g00001.1 | 25 | 8 | -1.47 | 0.010334569 | AT5G41460.1 | Protein of unknown function (DUF604) LENGTH=524 | IPR006740 (Protein of unknown function DUF604) | NA |
| Niben101Scf12160g01023.1 | 18 | 6 | -1.466 | 0.020713286 | sp\|Q5L0K0\|PYRH_GEOKA | Uridylate kinase | IPR001048 (Aspartate/glutamate/uridylate kinase), IPR015963 (Uridylate kinase, bacteria) | GO:0005737 (cytoplasm), GO:0006221 (pyrimidine nucleotide biosynthetic process), GO:0033862 (UMP kinase activity) |
| Niben101Scf00378g01009.1 | 20 | 6 | -1.465 | 0.044429899 | sp\|Q8DF81\|PUR4_VIBVU | Phosphoribosylformylglycinamidine synthase | IPR010073 (Phosphoribosylformylglycinamidine synthase), IPR016188 (PurM, N-terminal-like), IPR029062 (Class I glutamine amidotransferase-like) | GO:0003824 (catalytic activity), GO:0004642 (phosphoribosylformylglycinamidine synthase activity), GO:0006189 ('de novo' IMP biosynthetic process) |
| Niben101Scf01102g03004.1 | 20 | 6 | -1.459 | 0.030587024 | sp\|B1NWE1\|RPOC1_MANES | DNA-directed RNA polymerase subunit beta' | IPR006592 (RNA polymerase, N-terminal), IPR007066 (RNA polymerase Rpb1, domain 3), IPR007080 (RNA polymerase Rpb1, domain 1), IPR007081 (RNA polymerase Rpb1, domain 5), IPR015801 (Copper amine oxidase, N2/N3-terminal), IPR021602 (Protein of unknown function DUF3223) | GO:0003677 (DNA binding), GO:0003899 (DNA-directed RNA polymerase activity), GO:0005507 (copper ion binding), GO:0006351 (transcription, DNA-templated), GO:0009308 (amine metabolic process), GO:0048038 (quinone binding) |
| Niben101Scf08975g01001.1 | 32 | 10 | -1.458 | 0.001959476 | sp\|O49595\|HMGB1_ARATH | High mobility group B protein 1 | IPR009071 (High mobility group box domain) | NA |
| Niben101Scf03832g01006.1 | 26 | 9 | -1.457 | 0.037227186 | sp\|P29545\|EF1B_ORYSJ | Elongation factor 1-beta | IPR014717 (Translation elongation factor EF1B/ribosomal protein S6) | GO:0003746 (translation elongation factor activity), GO:0005853 (eukaryotic translation elongation factor 1 complex), GO:0006414 (translational elongation) |
| Niben101Scf05343g01007.1 | 24 | 8 | -1.451 | 0.012716326 | AT5G53550.1 | YELLOW STRIPE like 3 LENGTH=675 | IPR004813 (Oligopeptide transporter, OPT superfamily) | GO:0055085 (transmembrane transport) |
| Niben101Scf04198g01001.1 | 27 | 9 | -1.451 | 0.007407961 | sp\|P51375\|GLTB_PORPU | Ferredoxin-dependent glutamate synthase | IPR013785 (Aldolase-type TIM barrel), IPR029055 (Nucleophile aminohydrolases, N-terminal) | GO:0003824 (catalytic activity), GO:0006537 (glutamate biosynthetic process), GO:0006807 (nitrogen compound metabolic process), GO:0008152 (metabolic process), GO:0015930 (glutamate synthase activity), GO:0016638 (oxidoreductase activity, acting on the CH-NH2 group of donors), GO:0055114 (oxidation-reduction process) |
| Niben101Scf03705g01011.1 | 16 | 5 | -1.445 | 0.040989911 | AT1G18160.1 | Protein kinase superfamily protein LENGTH=992 | IPR011009 (Protein kinase-like domain) | GO:0004672 (protein kinase activity), GO:0004674 (protein serine/threonine kinase activity), GO:0005524 (ATP binding), GO:0006468 (protein phosphorylation), GO:0016772 (transferase activity, transferring phosphorus-containing groups) |
| Niben101Scf06840g00010.1 | 34 | 11 | -1.444 | 0.000967701 | sp\|A9LYE0\|RK22_ACOAM | 50S ribosomal protein L22, chloroplastic | IPR001063 (Ribosomal protein L22/L17), IPR001351 (Ribosomal protein S3, C-terminal), IPR009019 (K homology domain, prokaryotic type), IPR023575 (Ribosomal protein S19, superfamily) | GO:0003723 (RNA binding), GO:0003735 (structural constituent of ribosome), GO:0005840 (ribosome), GO:0006412 (translation), GO:0015934 (large ribosomal subunit) |
| Niben101Scf06726g00034.1 | 373 | 124 | -1.444 | 4.33547E-09 | sp\|Q942D4\|BURP3_ORYSJ | BURP domain-containing protein 3 | IPR004873 (BURP domain) | NA |
| Niben101Scf13036g01001.1 | 70 | 23 | -1.443 | 2.81586E-05 | dbj\|BAF52855.1\| | repressor of silencing 1 [Nicotiana tabacum] | IPR011257 (DNA glycosylase), IPR023170 (Helix-turn-helix, base-excision DNA repair, C-terminal), IPR028924 (Permuted single zf-CXXC unit), IPR028925 (Demeter, RRM-fold domain) | GO:0003824 (catalytic activity), GO:0006281 (DNA repair), GO:0051539 (4 iron, 4 sulfur cluster binding) |
| Niben101Scf04829g00009.1 | 25 | 8 | -1.442 | 0.002671194 | sp\|Q7RBX5\|KC1_PLAYO | Casein kinase I | IPR011009 (Protein kinase-like domain) | GO:0004672 (protein kinase activity), GO:0004674 (protein serine/threonine kinase activity), GO:0005524 (ATP binding), GO:0006468 (protein phosphorylation), GO:0016772 (transferase activity, transferring phosphorus-containing groups) |
| Niben101Scf03169g05002.1 | 75 | 25 | -1.439 | 0.001185914 | sp\|Q0G9N3\|ATPF_LIRTU | ATP synthase subunit b, chloroplastic | IPR002146 (ATPase, F0 complex, subunit B/B', bacterial/chloroplast), IPR004100 (ATPase, F1 complex alpha/beta subunit, N-terminal domain), IPR023366 (ATP synthase subunit alpha-like domain) | GO:0015078 (hydrogen ion transmembrane transporter activity), GO:0015986 (ATP synthesis coupled proton transport), GO:0015992 (proton transport), GO:0045263 (proton-transporting ATP synthase complex, coupling factor F(o)), GO:0046034 (ATP metabolic process) |
| Niben101Scf03535g01024.1 | 17 | 6 | -1.438 | 0.02582884 | sp\|O60870\|KIN17_HUMAN | DNA/RNA-binding protein KIN17 | IPR019447 (DNA/RNA-binding protein Kin17, conserved domain) | NA |
| Niben101Scf03066g02005.1 | 21 | 7 | -1.436 | 0.022971905 | sp\|P49756\|RBM25_HUMAN | RNA-binding protein 25 | IPR002483 (PWI domain), IPR012677 (Nucleotide-binding alpha-beta plait domain) | GO:0000166 (nucleotide binding), GO:0003676 (nucleic acid binding), GO:0006397 (mRNA processing) |
| Niben101Scf03673g02009.1 | 47 | 16 | -1.433 | 0.004636749 | sp\|P93047\|HMGB3_ARATH | High mobility group B protein 3 | IPR009071 (High mobility group box domain) | NA |
| Niben101Scf00683g03020.1 | 26 | 9 | -1.43 | 0.010881991 | sp\|Q71KN3\|CYB6_KLEBI | Cytochrome b6 | IPR016174 (Di-haem cytochrome, transmembrane), IPR023530 (Cytochrome b6, PetB), IPR027387 (Cytochrome b/b6-like domain) | GO:0005506 (iron ion binding), GO:0009055 (electron carrier activity), GO:0016020 (membrane), GO:0016021 (integral component of membrane), GO:0016491 (oxidoreductase activity), GO:0020037 (heme binding), GO:0022900 (electron transport chain), GO:0022904 (respiratory electron transport chain), GO:0045158 (electron transporter, transferring electrons within cytochrome b6/f complex of photosystem II activity) |
| Niben101Scf02279g08003.1 | 23 | 8 | -1.428 | 0.039686081 | AT1G69780.1 | Homeobox-leucine zipper protein family LENGTH=294 | IPR003106 (Leucine zipper, homeobox-associated), IPR009057 (Homeodomain-like) | GO:0003677 (DNA binding), GO:0003700 (sequence-specific DNA binding transcription factor activity), GO:0005634 (nucleus), GO:0006355 (regulation of transcription, DNA-templated), GO:0043565 (sequence-specific DNA binding) |
| Niben101Scf04104g07002.1 | 18 | 6 | -1.426 | 0.038297417 | AT1G05940.1 | cationic amino acid transporter 9 LENGTH=569 | IPR002293 (Amino acid/polyamine transporter I), IPR029485 (Cationic amino acid transporter, C-terminal) | GO:0003333 (amino acid transmembrane transport), GO:0015171 (amino acid transmembrane transporter activity), GO:0016020 (membrane) |
| Niben101Scf07953g02010.1 | 18 | 6 | -1.422 | 0.030074505 | sp\|Q6K5F8\|CDKG1_ORYSJ | Cyclin-dependent kinase G-1 | IPR011009 (Protein kinase-like domain) | GO:0004672 (protein kinase activity), GO:0004674 (protein serine/threonine kinase activity), GO:0005524 (ATP binding), GO:0006468 (protein phosphorylation), GO:0016772 (transferase activity, transferring phosphorus-containing groups) |
| Niben101Scf17701g01017.1 | 19 | 6 | -1.418 | 0.030074505 | sp\|Q7XA78\|D27_ARATH | Beta-carotene isomerase D27, chloroplastic | IPR025114 (Domain of unknown function DUF4033) | NA |
| Niben101Scf03648g02002.1 | 84 | 28 | -1.418 | 3.62426E-06 | sp\|A8Z5X0\|MNME_SULMW | tRNA modification GTPase MnmE | IPR005690 (Chloroplast protein import component Toc86/159), IPR024283 (Domain of unknown function DUF3406, chloroplast translocase), IPR027417 (P-loop containing nucleoside triphosphate hydrolase) | GO:0005525 (GTP binding), GO:0016817 (hydrolase activity, acting on acid anhydrides) |
| Niben101Scf03039g01020.1 | 17 | 6 | -1.413 | 0.033044044 | sp\|Q9C8K7\|FBK21_ARATH | F-box/kelch-repeat protein | IPR001810 (F-box domain), IPR011043 (Galactose oxidase/kelch, beta-propeller), IPR015915 (Kelch-type beta propeller) | GO:0005515 (protein binding) |
| Niben101Scf01832g00006.1 | 16 | 5 | -1.411 | 0.027995757 | sp\|Q7KKH3\|SDA1_DROME | Protein SDA1 homolog | IPR007949 (SDA1 domain), IPR016024 (Armadillo-type fold), IPR027312 (Sda1) | GO:0000055 (ribosomal large subunit export from nucleus), GO:0005488 (binding), GO:0030036 (actin cytoskeleton organization), GO:0042273 (ribosomal large subunit biogenesis) |
| Niben101Scf17594g00006.1 | 56 | 19 | -1.411 | 0.000294263 | AT4G31500.1 | cytochrome P450, family 83, subfamily B, polypeptide 1 LENGTH=499 | IPR001128 (Cytochrome P450) | GO:0005506 (iron ion binding), GO:0016705 (oxidoreductase activity, acting on paired donors, with incorporation or reduction of molecular oxygen), GO:0020037 (heme binding), GO:0055114 (oxidation-reduction process) |
| Niben101Scf18347g00015.1 | 22 | 7 | -1.409 | 0.0167906 | sp\|Q84MC7\|PYL9_ARATH | Abscisic acid receptor PYL9 | IPR019587 (Polyketide cyclase/dehydrase), IPR023393 (START-like domain) | NA |
| Niben101Scf07631g02024.1 | 15 | 5 | -1.408 | 0.047451637 | sp\|Q9SQI2\|GIGAN_ARATH | Protein GIGANTEA | IPR026211 (GIGANTEA) | GO:2000028 (regulation of photoperiodism, flowering) |
| Niben101Scf12383g01002.1 | 16 | 5 | -1.408 | 0.036644794 | sp\|P49756\|RBM25_HUMAN | RNA-binding protein 25 | IPR002483 (PWI domain), IPR012677 (Nucleotide-binding alpha-beta plait domain) | GO:0000166 (nucleotide binding), GO:0003676 (nucleic acid binding), GO:0006397 (mRNA processing) |
| Niben101Scf21348g00006.1 | 23 | 8 | -1.408 | 0.041243451 | sp\|B0JXB7\|UCRI_MICAN | Cytochrome b6-f complex iron-sulfur subunit | IPR014349 (Rieske iron-sulphur protein), IPR014909 (Cytochrome b6-f complex Fe-S subunit), IPR023960 (Cytochrome b6-f complex iron-sulfur subunit) | GO:0008121 (ubiquinol-cytochrome-c reductase activity), GO:0009496 (plastoquinol--plastocyanin reductase activity), GO:0015979 (photosynthesis), GO:0016020 (membrane), GO:0016491 (oxidoreductase activity), GO:0016679 (oxidoreductase activity, acting on diphenols and related substances as donors), GO:0042651 (thylakoid membrane), GO:0045158 (electron transporter, transferring electrons within cytochrome b6/f complex of photosystem II activity), GO:0051537 (2 iron, 2 sulfur cluster binding), GO:0055114 (oxidation-reduction process) |
| Niben101Scf16888g00015.1 | 36 | 12 | -1.404 | 0.010016217 | AT2G21170.1 | triosephosphate isomerase LENGTH=315 | IPR000652 (Triosephosphate isomerase), IPR013785 (Aldolase-type TIM barrel) | GO:0003824 (catalytic activity), GO:0004807 (triose-phosphate isomerase activity), GO:0006096 (glycolytic process), GO:0008152 (metabolic process) |
| Niben101Scf07608g04004.1 | 49 | 17 | -1.403 | 0.042523861 | sp\|Q9M2V7\|AB16G_ARATH | ABC transporter G family member 16 | IPR003439 (ABC transporter-like), IPR013525 (ABC-2 type transporter), IPR027417 (P-loop containing nucleoside triphosphate hydrolase) | GO:0005524 (ATP binding), GO:0016020 (membrane), GO:0016887 (ATPase activity) |
| Niben101Scf04040g05013.1 | 22 | 7 | -1.399 | 0.032395766 | ref\|XP_007011616.1\| | Centromere-associated protein E, putative isoform 1 [Theobroma cacao] gb\|EOY29235.1\| Centromere-associated protein E, putative isoform 1 [Theobroma cacao] | NA | NA |
| Niben101Scf08975g02018.1 | 44 | 15 | -1.397 | 0.007023056 | sp\|Q93Z81\|CAX3_ARATH | Vacuolar cation/proton exchanger 3 | IPR004713 (Calcium/proton exchanger) | GO:0006812 (cation transport), GO:0006816 (calcium ion transport), GO:0008324 (cation transmembrane transporter activity), GO:0015369 (calcium:proton antiporter activity), GO:0016021 (integral component of membrane), GO:0055085 (transmembrane transport) |
| Niben101Scf00894g01012.1 | 55 | 19 | -1.396 | 7.31124E-05 | gb\|ADK97918.1\| | LuxR family transcriptional regulator [Schiedea globosa] | NA | NA |
| Niben101Scf06537g00004.1 | 50 | 17 | -1.395 | 0.000491529 | sp\|Q9LF10\|CRS1_ARATH | Chloroplastic group IIA intron splicing facilitator CRS1, chloroplastic | IPR001890 (RNA-binding, CRM domain) | GO:0003723 (RNA binding) |
| Niben101Scf12680g03014.1 | 37 | 13 | -1.393 | 0.010732156 | sp\|A1U7J4\|YIDC_MARHV | Membrane protein insertase YidC | IPR001708 (Membrane insertase OXA1/ALB3/YidC), IPR028055 (Membrane insertase YidC/Oxa1, C-terminal) | GO:0016021 (integral component of membrane), GO:0051205 (protein insertion into membrane) |
| Niben101Scf04995g01002.1 | 31 | 11 | -1.389 | 0.017300533 | emb\|CAE01801.2\| | product [Oryza sativa Japonica Group] | IPR004864 (Late embryogenesis abundant protein, LEA-14) | NA |
| Niben101Scf02235g04025.1 | 56 | 19 | -1.389 | 0.000743811 | sp\|Q8S8Y2\|ATPF_ATRBE | ATP synthase subunit b, chloroplastic | NA | NA |
| Niben101Scf02816g16025.1 | 24 | 8 | -1.388 | 0.02583416 | gb\|KEH19038.1\| | brefeldin A-inhibited guanine nucleotide-exchange protein [Medicago truncatula] | IPR000904 (Sec7 domain), IPR011992 (EF-hand domain pair), IPR016024 (Armadillo-type fold), IPR023394 (Sec7 domain, alpha orthogonal bundle) | GO:0005086 (ARF guanyl-nucleotide exchange factor activity), GO:0005488 (binding), GO:0005509 (calcium ion binding), GO:0032012 (regulation of ARF protein signal transduction) |
| Niben101Scf08026g00008.1 | 49 | 17 | -1.388 | 0.00079703 | AT4G17840.1 | FUNCTIONS IN: molecular_function unknown; INVOLVED IN: biological_process unknown; LOCATED IN: chloroplast, membrane; EXPRESSED IN: 23 plant structures; EXPRESSED DURING: 13 growth stages; | IPR003675 (CAAX amino terminal protease) | GO:0016020 (membrane) |
| Niben101Scf00987g02009.1 | 28 | 10 | -1.385 | 0.004795611 | AT5G43310.1 | COP1-interacting protein-related LENGTH=1237 | NA | NA |
| Niben101Scf01734g01046.1 | 433 | 150 | -1.383 | 3.10156E-10 | gb\|AFH58131.1\| | Ycf6, partial (chloroplast) [Dombeya acutangula] | NA | NA |
| Niben101Scf10630g02001.1 | 27 | 9 | -1.381 | 0.047134766 | AT3G14270.1 | phosphatidylinositol-4-phosphate 5-kinase family protein LENGTH=1791 | IPR002423 (Chaperonin Cpn60/TCP-1), IPR013083 (Zinc finger, RING/FYVE/PHD-type), IPR027409 (GroEL-like apical domain) | GO:0005524 (ATP binding), GO:0044267 (cellular protein metabolic process), GO:0046872 (metal ion binding) |
| Niben101Scf01472g02001.1 | 17 | 6 | -1.378 | 0.039751815 | sp\|Q57242\|UUP1_HAEIN | ABC transporter ATP-binding protein uup-1 | IPR003439 (ABC transporter-like), IPR027417 (P-loop containing nucleoside triphosphate hydrolase) | GO:0005524 (ATP binding), GO:0016887 (ATPase activity) |
| Niben101Scf01848g02014.1 | 32 | 11 | -1.377 | 0.008115048 | sp\|P54119\|CDK1_AJECA | Cyclin-dependent kinase 1 | IPR011009 (Protein kinase-like domain) | GO:0004672 (protein kinase activity), GO:0004674 (protein serine/threonine kinase activity), GO:0005524 (ATP binding), GO:0006468 (protein phosphorylation), GO:0016772 (transferase activity, transferring phosphorus-containing groups) |
| Niben101Scf01399g00012.1 | 26 | 9 | -1.376 | 0.028353374 | sp\|A9IYW8\|ATPG_BART1 | ATP synthase gamma chain | IPR000131 (ATPase, F1 complex, gamma subunit), IPR023632 (ATPase, F1 complex, gamma subunit conserved site), IPR023633 (ATPase, F1 complex, gamma subunit domain) | GO:0015986 (ATP synthesis coupled proton transport), GO:0045261 (proton-transporting ATP synthase complex, catalytic core F(1)), GO:0046933 (proton-transporting ATP synthase activity, rotational mechanism), GO:0046961 (proton-transporting ATPase activity, rotational mechanism) |
| Niben101Scf00225g05002.1 | 34 | 12 | -1.376 | 0.00192978 | sp\|Q869S5\|DOKA_DICDI | Hybrid signal transduction protein dokA | IPR011006 (CheY-like superfamily) | GO:0000160 (phosphorelay signal transduction system) |
| Niben101Scf04375g09011.1 | 47 | 16 | -1.375 | 0.001029906 | sp\|Q6DEL2\|CLPT1_DANRE | Cleft lip and palate transmembrane protein 1 homolog | IPR008429 (Cleft lip and palate transmembrane 1) | GO:0016021 (integral component of membrane) |
| Niben101Scf01771g01021.1 | 43 | 15 | -1.367 | 0.001487226 | sp\|Q8L7W2\|NUDT8_ARATH | Nudix hydrolase 8 | IPR003293 (Nudix hydrolase 6-like) | GO:0016787 (hydrolase activity) |
| Niben101Scf05929g02002.1 | 29 | 10 | -1.366 | 0.047464534 | sp\|Q5RC80\|RBM39_PONAB | RNA-binding protein 39 | IPR012677 (Nucleotide-binding alpha-beta plait domain) | GO:0000166 (nucleotide binding), GO:0003676 (nucleic acid binding) |
| Niben101Scf01005g02014.1 | 48 | 17 | -1.364 | 0.003033069 | sp\|Q9LSW7\|ACCH9_ARATH | 1-aminocyclopropane-1-carboxylate oxidase homolog 9 | IPR005123 (Oxoglutarate/iron-dependent dioxygenase), IPR027443 (Isopenicillin N synthase-like) | GO:0016491 (oxidoreductase activity), GO:0016706 (oxidoreductase activity, acting on paired donors, with incorporation or reduction of molecular oxygen, 2-oxoglutarate as one donor, and incorporation of one atom each of oxygen into both donors), GO:0055114 (oxidation-reduction process) |
| Niben101Scf09416g05007.1 | 21 | 7 | -1.363 | 0.039751815 | NA | Unknown protein | NA | NA |
| Niben101Scf05154g01012.1 | 34 | 12 | -1.359 | 0.01663746 | ref\|XP_007039076.1\| | Ribonuclease P protein subunit P38-related isoform 2 [Theobroma cacao] gb\|EOY23577.1\| Ribonuclease P protein subunit P38-related isoform 2 [Theobroma cacao] | IPR020849 (Small GTPase superfamily, Ras type) | GO:0005525 (GTP binding), GO:0006184 (GTP catabolic process), GO:0007165 (signal transduction), GO:0016020 (membrane) |
| Niben101Scf00597g05012.1 | 45 | 16 | -1.359 | 0.000162923 | gb\|ABF95303.1\| | expressed protein [Oryza sativa Japonica Group] | IPR022212 (Protein of unknown function DUF3741), IPR025486 (Domain of unknown function DUF4378) | NA |
| Niben101Scf09559g02008.1 | 15 | 5 | -1.353 | 0.046280709 | AT1G26480.1 | general regulatory factor 12 LENGTH=268 | IPR000308 (14-3-3 protein), IPR023409 (14-3-3 protein, conserved site), IPR023410 (14-3-3 domain) | GO:0019904 (protein domain specific binding) |
| Niben101Scf06856g00002.1 | 22 | 8 | -1.353 | 0.02271915 | sp\|O23693\|MLO4_ARATH | MLO-like protein 4 | IPR004326 (Mlo-related protein) | GO:0006952 (defense response), GO:0016021 (integral component of membrane) |
| Niben101Scf14977g00025.1 | 21 | 7 | -1.352 | 0.028591545 | AT3G05840.2 | Protein kinase superfamily protein LENGTH=409 | IPR011009 (Protein kinase-like domain) | GO:0004672 (protein kinase activity), GO:0004674 (protein serine/threonine kinase activity), GO:0005524 (ATP binding), GO:0006468 (protein phosphorylation), GO:0016772 (transferase activity, transferring phosphorus-containing groups) |
| Niben101Scf01175g03010.1 | 119 | 42 | -1.35 | 1.43628E-05 | sp\|O23144\|PPI1_ARATH | Proton pump-interactor 1 | IPR029669 (Proton pump-interactor) | GO:0005783 (endoplasmic reticulum), GO:0005886 (plasma membrane), GO:0010155 (regulation of proton transport) |
| Niben101Scf01175g03010.1 | 28 | 10 | -1.349 | 0.016245873 | sp\|O23144\|PPI1_ARATH | Proton pump-interactor 1 | IPR029669 (Proton pump-interactor) | GO:0005783 (endoplasmic reticulum), GO:0005886 (plasma membrane), GO:0010155 (regulation of proton transport) |
| Niben101Scf00503g05016.1 | 115 | 40 | -1.349 | 0.000316272 | AT3G55570.1 | unknown protein; BEST Arabidopsis thaliana protein match is: unknown protein . LENGTH=109 | NA | NA |
| Niben101Scf05621g13007.1 | 19 | 7 | -1.337 | 0.049464262 | sp\|P10708\|CB12_SOLLC | Chlorophyll a-b binding protein 7, chloroplastic | IPR022796 (Chlorophyll A-B binding protein), IPR023329 (Chlorophyll a/b binding protein domain) | GO:0009765 (photosynthesis, light harvesting), GO:0016020 (membrane) |
| Niben101Scf08680g00003.1 | 23 | 8 | -1.333 | 0.041531077 | sp\|Q9LQ12\|4CLL1_ARATH | 4-coumarate--CoA ligase-like 1 | IPR000873 (AMP-dependent synthetase/ligase), IPR025110 (AMP-binding enzyme C-terminal domain) | GO:0003824 (catalytic activity), GO:0008152 (metabolic process) |
| Niben101Scf07631g02024.1 | 163 | 59 | -1.333 | 2.21177E-08 | sp\|Q9SQI2\|GIGAN_ARATH | Protein GIGANTEA | IPR026211 (GIGANTEA) | GO:2000028 (regulation of photoperiodism, flowering) |
| Niben101Scf00797g20001.1 | 21 | 7 | -1.331 | 0.027479665 | sp\|Q7KKH3\|SDA1_DROME | Protein SDA1 homolog | IPR007949 (SDA1 domain), IPR016024 (Armadillo-type fold), IPR027312 (Sda1) | GO:0000055 (ribosomal large subunit export from nucleus), GO:0005488 (binding), GO:0030036 (actin cytoskeleton organization), GO:0042273 (ribosomal large subunit biogenesis) |
| Niben101Scf04477g00011.1 | 33 | 12 | -1.331 | 0.005905227 | AT1G50660.1 | unknown protein; INVOLVED IN: biological_process unknown; LOCATED IN: chloroplast; EXPRESSED IN: 22 plant structures; EXPRESSED DURING: 13 growth stages; BEST Arabidopsis thaliana protein match is: unknown protein . LENGTH=725 | NA | NA |
| Niben101Scf07311g01002.1 | 27 | 10 | -1.324 | 0.013300565 | NA | Unknown protein | IPR013880 (Yos1-like) | NA |
| Niben101Scf00383g03001.1 | 25 | 9 | -1.321 | 0.043750454 | sp\|A9KIG5\|FTSH_CLOPH | ATP-dependent zinc metalloprotease FtsH | IPR003959 (ATPase, AAA-type, core), IPR025753 (AAA-type ATPase, N-terminal domain), IPR027417 (P-loop containing nucleoside triphosphate hydrolase) | GO:0005524 (ATP binding) |
| Niben101Scf09253g00007.1 | 37 | 13 | -1.317 | 0.006135229 | sp\|P46824\|KLC_DROME | Kinesin light chain | IPR011990 (Tetratricopeptide-like helical domain) | GO:0005515 (protein binding) |
| Niben101Scf06996g02041.1 | 120 | 44 | -1.316 | 0.000160401 | sp\|B0B7R0\|DNAJ_CHLT2 | Chaperone protein DnaJ | IPR001623 (DnaJ domain) | NA |
| Niben101Scf07944g05004.1 | 31 | 11 | -1.312 | 0.009280507 | sp\|Q54JC8\|PUR4_DICDI | Phosphoribosylformylglycinamidine synthase | NA | NA |
| Niben101Scf05969g00011.1 | 22 | 8 | -1.308 | 0.037783386 | sp\|Q96JC1\|VPS39_HUMAN | Vam6/Vps39-like protein | IPR000547 (Clathrin, heavy chain/VPS, 7-fold repeat), IPR001180 (Citron-like), IPR019452 (Vacuolar sorting protein 39/Transforming growth factor beta receptor-associated domain 1), IPR019453 (Vacuolar sorting protein 39/Transforming growth factor beta receptor-associated domain 2) | GO:0005083 (small GTPase regulator activity), GO:0006886 (intracellular protein transport), GO:0016192 (vesicle-mediated transport) |
| Niben101Scf09698g01005.1 | 65 | 23 | -1.303 | 0.031318826 | AT2G41870.1 | Remorin family protein LENGTH=274 | IPR005516 (Remorin, C-terminal) | NA |
| Niben101Scf05405g00002.1 | 30 | 11 | -1.302 | 0.015511271 | AT3G12620.1 | Protein phosphatase 2C family protein LENGTH=385 | IPR001932 (Protein phosphatase 2C (PP2C)-like domain), IPR015655 (Protein phosphatase 2C) | GO:0003824 (catalytic activity), GO:0004722 (protein serine/threonine phosphatase activity), GO:0006470 (protein dephosphorylation) |
| Niben101Scf02124g00017.1 | 19 | 7 | -1.3 | 0.026321513 | gb\|KEH28577.1\| | Snf1-related kinase interactor 1, putative [Medicago truncatula] | NA | NA |
| Niben101Scf01142g03007.1 | 24 | 9 | -1.3 | 0.041940866 | sp\|Q67PA6\|SYP_SYMTH | Proline--tRNA ligase | IPR002316 (Proline-tRNA ligase, class IIa), IPR017449 (Prolyl-tRNA synthetase, class II) | GO:0000166 (nucleotide binding), GO:0004812 (aminoacyl-tRNA ligase activity), GO:0004827 (proline-tRNA ligase activity), GO:0005524 (ATP binding), GO:0005737 (cytoplasm), GO:0006418 (tRNA aminoacylation for protein translation), GO:0006433 (prolyl-tRNA aminoacylation) |
| Niben101Scf01455g01022.1 | 77 | 28 | -1.299 | 6.88008E-05 | sp\|Q9ZTW3\|VA721_ARATH | Vesicle-associated membrane protein 721 | IPR001388 (Synaptobrevin), IPR011012 (Longin-like domain) | GO:0006810 (transport), GO:0016021 (integral component of membrane), GO:0016192 (vesicle-mediated transport) |
| Niben101Scf18384g00001.1 | 99 | 37 | -1.295 | 0.000275608 | sp\|Q93VH2\|BBD2_ARATH | Bifunctional nuclease 2 | IPR003729 (Bifunctional nuclease domain) | GO:0004518 (nuclease activity) |
| Niben101Scf06996g02041.1 | 40 | 15 | -1.293 | 0.009469483 | sp\|B0B7R0\|DNAJ_CHLT2 | Chaperone protein DnaJ | IPR001623 (DnaJ domain) | NA |
| Niben101Scf05824g12003.1 | 31 | 11 | -1.288 | 0.004541414 | AT4G31820.1 | Phototropic-responsive NPH3 family protein LENGTH=571 | IPR011333 (BTB/POZ fold), IPR027356 (NPH3 domain), IPR029961 (BTB/POZ domain-containing protein NPY1) | GO:0004871 (signal transducer activity), GO:0005515 (protein binding), GO:0048513 (organ development), GO:0060918 (auxin transport) |
| Niben101Scf01998g03011.1 | 27 | 10 | -1.286 | 0.04468457 | AT2G36780.1 | UDP-Glycosyltransferase superfamily protein LENGTH=496 | IPR002213 (UDP-glucuronosyl/UDP-glucosyltransferase) | GO:0008152 (metabolic process), GO:0016758 (transferase activity, transferring hexosyl groups) |
| Niben101Scf08415g01034.1 | 23 | 8 | -1.281 | 0.04662556 | gb\|KEH43713.1\| | myosin heavy chain-like protein [Medicago truncatula] | IPR019448 (EEIG1/EHBP1 N-terminal domain) | NA |
| Niben101Scf00578g00008.1 | 29 | 11 | -1.275 | 0.036471677 | sp\|Q9XFT3\|PSBQ1_ARATH | Oxygen-evolving enhancer protein 3-1, chloroplastic | IPR008797 (Photosystem II PsbQ, oxygen evolving complex), IPR023222 (PsbQ-like domain) | GO:0005509 (calcium ion binding), GO:0009523 (photosystem II), GO:0009654 (photosystem II oxygen evolving complex), GO:0015979 (photosynthesis), GO:0019898 (extrinsic component of membrane) |
| Niben101Scf01453g02010.1 | 92 | 34 | -1.27 | 2.52363E-07 | sp\|Q9SQI2\|GIGAN_ARATH | Protein GIGANTEA | IPR026211 (GIGANTEA) | GO:2000028 (regulation of photoperiodism, flowering) |
| Niben101Scf02078g08043.1 | 35 | 13 | -1.269 | 0.009705938 | sp\|Q6PGE4\|ZF316_MOUSE | Zinc finger protein 316 | IPR013087 (Zinc finger C2H2-type/integrase DNA-binding domain) | GO:0003676 (nucleic acid binding), GO:0046872 (metal ion binding) |
| Niben101Scf02902g01014.1 | 36 | 13 | -1.266 | 0.010854743 | sp\|A0AUR5\|F188A_DANRE | Protein FAM188A | IPR003903 (Ubiquitin interacting motif), IPR011992 (EF-hand domain pair), IPR025257 (Domain of unknown function DUF4205) | NA |
| Niben101Scf01072g03042.1 | 33 | 12 | -1.26 | 0.02869223 | sp\|Q9JUU8\|RLMB_NEIMA | 23S rRNA (guanosine-2'-O-)-methyltransferase RlmB | IPR029028 (Alpha/beta knot methyltransferases) | GO:0003723 (RNA binding), GO:0006396 (RNA processing), GO:0008173 (RNA methyltransferase activity) |
| Niben101Scf03080g04005.1 | 38 | 14 | -1.26 | 0.004120414 | sp\|Q43383\|ACCH5_ARATH | 1-aminocyclopropane-1-carboxylate oxidase homolog 5 | IPR002283 (Isopenicillin N synthase), IPR026992 (Non-haem dioxygenase N-terminal domain), IPR027443 (Isopenicillin N synthase-like) | GO:0005506 (iron ion binding), GO:0016491 (oxidoreductase activity), GO:0016706 (oxidoreductase activity, acting on paired donors, with incorporation or reduction of molecular oxygen, 2-oxoglutarate as one donor, and incorporation of one atom each of oxygen into both donors), GO:0055114 (oxidation-reduction process) |
| Niben101Scf07121g02017.1 | 46 | 17 | -1.255 | 0.006478884 | AT3G54360.1 | zinc ion binding LENGTH=405 | IPR011990 (Tetratricopeptide-like helical domain), IPR013083 (Zinc finger, RING/FYVE/PHD-type) | GO:0005515 (protein binding), GO:0008270 (zinc ion binding) |
| Niben101Scf07339g02015.1 | 461 | 174 | -1.255 | 0.0374643 | sp\|A5UE38\|GLPE_HAEIE | Thiosulfate sulfurtransferase GlpE | IPR001763 (Rhodanese-like domain) | NA |
| Niben101Scf04388g00011.1 | 21 | 8 | -1.252 | 0.042077774 | sp\|O50314\|BCHH_CHLP8 | Magnesium-chelatase subunit H | IPR003672 (CobN/magnesium chelatase) | GO:0009058 (biosynthetic process), GO:0015995 (chlorophyll biosynthetic process), GO:0016851 (magnesium chelatase activity) |
| Niben101Scf11689g02005.1 | 32 | 12 | -1.252 | 0.015476193 | AT2G37180.1 | Aquaporin-like superfamily protein LENGTH=285 | IPR000425 (Major intrinsic protein), IPR023271 (Aquaporin-like) | GO:0005215 (transporter activity), GO:0006810 (transport), GO:0016020 (membrane) |
| Niben101Scf05943g00002.1 | 81 | 30 | -1.25 | 1.70945E-05 | sp\|Q40459\|PSBO_TOBAC | Oxygen-evolving enhancer protein 1, chloroplastic | IPR002628 (Photosystem II PsbO, manganese-stabilising), IPR011250 (Outer membrane protein/outer membrane enzyme PagP , beta-barrel) | GO:0005509 (calcium ion binding), GO:0009279 (cell outer membrane), GO:0009523 (photosystem II), GO:0009654 (photosystem II oxygen evolving complex), GO:0015979 (photosynthesis), GO:0016021 (integral component of membrane), GO:0019898 (extrinsic component of membrane), GO:0042549 (photosystem II stabilization) |
| Niben101Scf11886g03034.1 | 28 | 10 | -1.247 | 0.0374643 | sp\|Q8CCP0\|NEMF_MOUSE | Nuclear export mediator factor Nemf | IPR001878 (Zinc finger, CCHC-type), IPR008532 (Domain of unknown function DUF814), IPR021846 (Protein of unknown function DUF3441) | GO:0003676 (nucleic acid binding), GO:0008270 (zinc ion binding) |
| Niben101Scf03253g02006.1 | 112 | 43 | -1.244 | 2.49571E-05 | sp\|B1L938\|RL1_THESQ | 50S ribosomal protein L1 | IPR016095 (Ribosomal protein L1, 3-layer alpha/beta-sandwich), IPR023673 (Ribosomal protein L1, conserved site), IPR023674 (Ribosomal protein L1-like), IPR028364 (Ribosomal protein L1/ribosomal biogenesis protein) | GO:0003723 (RNA binding), GO:0003735 (structural constituent of ribosome), GO:0006412 (translation), GO:0015934 (large ribosomal subunit) |
| Niben101Scf06655g00013.1 | 40 | 15 | -1.242 | 0.012133009 | gb\|KEH37012.1\| | E3 ubiquitin-protein ligase LIN-like protein, putative [Medicago truncatula] | IPR016024 (Armadillo-type fold) | GO:0005488 (binding) |
| Niben101Scf02730g01006.1 | 112 | 43 | -1.237 | 2.11345E-06 | sp\|Q5E992\|PPIL1_BOVIN | Peptidyl-prolyl cis-trans isomerase-like 1 | IPR002130 (Cyclophilin-type peptidyl-prolyl cis-trans isomerase domain), IPR023222 (PsbQ-like domain), IPR029000 (Cyclophilin-like domain) | GO:0000413 (protein peptidyl-prolyl isomerization), GO:0003755 (peptidyl-prolyl cis-trans isomerase activity), GO:0006457 (protein folding) |
| Niben101Scf08092g00003.1 | 25 | 9 | -1.235 | 0.045070252 | gb\|KEH39902.1\| | Smr (small MutS-related) domain protein [Medicago truncatula] | IPR002625 (Smr protein/MutS2 C-terminal) | NA |
| Niben101Scf01582g03003.1 | 29 | 11 | -1.235 | 0.036788535 | ref\|NP_001049616.1\| | product [Oryza sativa Japonica Group] | IPR007656 (Zein-binding domain) | NA |
| Niben101Scf05142g00023.1 | 1119 | 429 | -1.234 | 2.16463E-08 | sp\|P82129\|RR7_SPIOL | 30S ribosomal protein S7, chloroplastic | IPR000235 (Ribosomal protein S5/S7), IPR023798 (Ribosomal protein S7 domain) | GO:0003723 (RNA binding), GO:0003735 (structural constituent of ribosome), GO:0006412 (translation) |
| Niben101Scf02293g03010.1 | 36 | 14 | -1.231 | 0.005828779 | emb\|CDX89334.1\| | BnaA01g15590D [Brassica napus] | NA | NA |
| Niben101Scf00851g05004.1 | 21 | 8 | -1.228 | 0.043701093 | AT1G04580.1 | aldehyde oxidase 4 LENGTH=1337 | IPR012675 (Beta-grasp domain), IPR016166 (FAD-binding, type 2), IPR016208 (Aldehyde oxidase/xanthine dehydrogenase) | GO:0003824 (catalytic activity), GO:0005506 (iron ion binding), GO:0009055 (electron carrier activity), GO:0016491 (oxidoreductase activity), GO:0016614 (oxidoreductase activity, acting on CH-OH group of donors), GO:0046872 (metal ion binding), GO:0050660 (flavin adenine dinucleotide binding), GO:0051536 (iron-sulfur cluster binding), GO:0051537 (2 iron, 2 sulfur cluster binding), GO:0055114 (oxidation-reduction process) |
| Niben101Scf05056g00005.1 | 20 | 8 | -1.219 | 0.049464262 | sp\|Q8XA87\|DEAD_ECO57 | ATP-dependent RNA helicase DeaD | IPR001650 (Helicase, C-terminal), IPR014001 (Helicase superfamily 1/2, ATP-binding domain), IPR014014 (RNA helicase, DEAD-box type, Q motif), IPR027417 (P-loop containing nucleoside triphosphate hydrolase) | GO:0003676 (nucleic acid binding), GO:0005524 (ATP binding) |
| Niben101Scf10122g00007.1 | 22 | 8 | -1.217 | 0.045003062 | sp\|P05100\|3MG1_ECOLI | DNA-3-methyladenine glycosylase 1 | IPR005019 (Methyladenine glycosylase) | GO:0003824 (catalytic activity), GO:0006281 (DNA repair), GO:0006284 (base-excision repair), GO:0008725 (DNA-3-methyladenine glycosylase activity) |
| Niben101Scf15224g00002.1 | 43 | 17 | -1.214 | 0.00534088 | sp\|Q9R210\|TFEB_MOUSE | Transcription factor EB | IPR011598 (Myc-type, basic helix-loop-helix (bHLH) domain), IPR025610 (Transcription factor MYC/MYB N-terminal) | GO:0046983 (protein dimerization activity) |
| Niben101Scf06059g02006.1 | 46 | 18 | -1.212 | 0.012959501 | sp\|Q9T068\|EPFL2_ARATH | EPIDERMAL PATTERNING FACTOR-like protein 2 | NA | NA |
| Niben101Scf04099g09005.1 | 143 | 56 | -1.208 | 1.1375E-07 | sp\|P74518\|LRTA_SYNY3 | Light-repressed protein A homolog | IPR003489 (Ribosomal protein S30Ae/sigma 54 modulation protein) | GO:0044238 (primary metabolic process) |
| Niben101Scf01143g11003.1 | 40 | 16 | -1.201 | 0.008534612 | ref\|NP_001067105.1\| | product [Oryza sativa Japonica Group] | IPR009500 (Protein of unknown function DUF1118) | NA |
| Niben101Scf00090g04014.1 | 27 | 11 | -1.198 | 0.029855212 | ref\|XP_007017230.1\| | Cold regulated gene 27, putative isoform 3 [Theobroma cacao] gb\|EOY14455.1\| Cold regulated gene 27, putative isoform 3 [Theobroma cacao] | NA | NA |
| Niben101Scf06674g00004.1 | 25 | 10 | -1.197 | 0.036001425 | sp\|Q9SJZ6\|MED18_ARATH | Mediator of RNA polymerase II transcription subunit 18 | NA | NA |
| Niben101Scf24155g00014.1 | 41 | 16 | -1.197 | 0.002199731 | ref\|XP_002875323.1\| | binding protein [Arabidopsis lyrata subsp. lyrata] gb\|EFH51582.1\| binding protein [Arabidopsis lyrata subsp. lyrata] | IPR011990 (Tetratricopeptide-like helical domain) | GO:0005515 (protein binding) |
| Niben101Scf04329g06015.1 | 56 | 22 | -1.194 | 0.0167906 | sp\|P74214\|SRP54_SYNY3 | Signal recognition particle protein | IPR004780 (Signal recognition particle protein Ffh), IPR027417 (P-loop containing nucleoside triphosphate hydrolase) | GO:0003924 (GTPase activity), GO:0005525 (GTP binding), GO:0006614 (SRP-dependent cotranslational protein targeting to membrane), GO:0008312 (7S RNA binding), GO:0048500 (signal recognition particle) |
| Niben101Scf09107g01004.1 | 101 | 40 | -1.194 | 0.000812998 | sp\|Q9LIB2\|PHS1_ARATH | Alpha-glucan phosphorylase 1 | IPR000811 (Glycosyl transferase, family 35) | GO:0005975 (carbohydrate metabolic process), GO:0008184 (glycogen phosphorylase activity) |
| Niben101Scf15067g01008.1 | 25 | 10 | -1.188 | 0.022449429 | AT3G61160.2 | Protein kinase superfamily protein LENGTH=438 | IPR011009 (Protein kinase-like domain) | GO:0016772 (transferase activity, transferring phosphorus-containing groups) |
| Niben101Scf03647g01004.1 | 22 | 9 | -1.184 | 0.047999884 | sp\|P14233\|TGA1B_TOBAC | TGACG-sequence-specific DNA-binding protein TGA-1B | IPR004827 (Basic-leucine zipper domain) | GO:0003700 (sequence-specific DNA binding transcription factor activity), GO:0006355 (regulation of transcription, DNA-templated), GO:0043565 (sequence-specific DNA binding) |
| Niben101Scf03039g01017.1 | 19 | 7 | -1.183 | 0.048037472 | sp\|A3CK48\|VATA_STRSV | V-type ATP synthase alpha chain | IPR005725 (ATPase, V1 complex, subunit A), IPR027417 (P-loop containing nucleoside triphosphate hydrolase) | GO:0005524 (ATP binding), GO:0015991 (ATP hydrolysis coupled proton transport), GO:0015992 (proton transport), GO:0016820 (hydrolase activity, acting on acid anhydrides, catalyzing transmembrane movement of substances), GO:0033178 (proton-transporting two-sector ATPase complex, catalytic domain), GO:0033180 (proton-transporting V-type ATPase, V1 domain), GO:0046034 (ATP metabolic process), GO:0046961 (proton-transporting ATPase activity, rotational mechanism) |
| Niben101Scf00927g00002.1 | 156 | 63 | -1.182 | 0.000146022 | sp\|Q9C8K7\|FBK21_ARATH | F-box/kelch-repeat protein | IPR000014 (PAS domain), IPR001810 (F-box domain), IPR011043 (Galactose oxidase/kelch, beta-propeller), IPR015915 (Kelch-type beta propeller) | GO:0004871 (signal transducer activity), GO:0005515 (protein binding), GO:0007165 (signal transduction) |
| Niben101Scf02408g00020.1 | 31 | 12 | -1.179 | 0.044098424 | AT3G19240.1 | Vacuolar import/degradation, Vid27-related protein LENGTH=648 | IPR013863 (Vacuolar import/degradation, Vid27-related), IPR015943 (WD40/YVTN repeat-like-containing domain) | GO:0005515 (protein binding) |
| Niben101Scf01184g17001.1 | 28 | 11 | -1.175 | 0.043701093 | AT4G35160.1 | O-methyltransferase family protein LENGTH=382 | IPR016461 (O-methyltransferase COMT-type), IPR029063 (S-adenosyl-L-methionine-dependent methyltransferase) | GO:0008168 (methyltransferase activity), GO:0008171 (O-methyltransferase activity), GO:0046983 (protein dimerization activity) |
| Niben101Scf08783g01002.1 | 79 | 31 | -1.172 | 5.84092E-05 | gb\|ADM94222.1\| | TGN-localized SYP41-interacting protein [Arabidopsis thaliana] | NA | NA |
| Niben101Scf08998g02001.1 | 33 | 13 | -1.17 | 0.022318401 | emb\|CDX99035.1\| | BnaC09g48000D [Brassica napus] | NA | NA |
| Niben101Scf26023g00004.1 | 452 | 183 | -1.167 | 0.019809562 | sp\|O24606\|EIN3_ARATH | Protein ETHYLENE INSENSITIVE 3 | NA | NA |
| Niben101Scf01627g03009.1 | 77 | 31 | -1.166 | 6.82946E-05 | sp\|Q9S7Y1\|BH030_ARATH | Transcription factor bHLH30 | IPR011598 (Myc-type, basic helix-loop-helix (bHLH) domain) | GO:0046983 (protein dimerization activity) |
| Niben101Scf04529g00005.1 | 123 | 49 | -1.165 | 7.31124E-05 | sp\|Q8DLP3\|ATPA_THEEB | ATP synthase subunit alpha | IPR005294 (ATPase, F1 complex, alpha subunit), IPR023366 (ATP synthase subunit alpha-like domain), IPR027417 (P-loop containing nucleoside triphosphate hydrolase) | GO:0005524 (ATP binding), GO:0015986 (ATP synthesis coupled proton transport), GO:0015991 (ATP hydrolysis coupled proton transport), GO:0015992 (proton transport), GO:0016820 (hydrolase activity, acting on acid anhydrides, catalyzing transmembrane movement of substances), GO:0033178 (proton-transporting two-sector ATPase complex, catalytic domain), GO:0045261 (proton-transporting ATP synthase complex, catalytic core F(1)), GO:0046034 (ATP metabolic process), GO:0046933 (proton-transporting ATP synthase activity, rotational mechanism), GO:0046961 (proton-transporting ATPase activity, rotational mechanism) |
| Niben101Scf06090g02010.1 | 47 | 19 | -1.159 | 0.020470188 | AT1G70940.1 | Auxin efflux carrier family protein LENGTH=640 | IPR004776 (Auxin efflux carrier) | GO:0016021 (integral component of membrane), GO:0055085 (transmembrane transport) |
| Niben101Scf07682g01012.1 | 23 | 9 | -1.158 | 0.0374643 | AT1G58470.1 | RNA-binding protein 1 LENGTH=360 | IPR012677 (Nucleotide-binding alpha-beta plait domain) | GO:0000166 (nucleotide binding), GO:0003676 (nucleic acid binding) |
| Niben101Scf07247g04003.1 | 38 | 15 | -1.156 | 0.017429683 | sp\|O23144\|PPI1_ARATH | Proton pump-interactor 1 | IPR029669 (Proton pump-interactor) | GO:0005783 (endoplasmic reticulum), GO:0005886 (plasma membrane), GO:0010155 (regulation of proton transport) |
| Niben101Scf05100g01013.1 | 65 | 26 | -1.154 | 0.002949703 | sp\|Q5RC80\|RBM39_PONAB | RNA-binding protein 39 | IPR012677 (Nucleotide-binding alpha-beta plait domain) | GO:0000166 (nucleotide binding), GO:0003676 (nucleic acid binding) |
| Niben101Scf19733g00001.1 | 34 | 14 | -1.152 | 0.034436638 | sp\|Q6K7E6\|ERF1_ORYSJ | Ethylene-responsive transcription factor 1 | IPR016177 (DNA-binding domain) | GO:0003677 (DNA binding), GO:0003700 (sequence-specific DNA binding transcription factor activity), GO:0006355 (regulation of transcription, DNA-templated) |
| Niben101Scf04495g00026.1 | 37 | 15 | -1.152 | 0.03133834 | AT5G38840.1 | SMAD/FHA domain-containing protein LENGTH=735 | IPR008984 (SMAD/FHA domain) | GO:0005515 (protein binding) |
| Niben101Scf05767g01014.1 | 55 | 22 | -1.152 | 0.002446666 | sp\|Q67ZM4\|ADF7_ARATH | Actin-depolymerizing factor 7 | IPR017904 (ADF/Cofilin/Destrin), IPR029006 (ADF-H/Gelsolin-like domain) | GO:0003779 (actin binding), GO:0005622 (intracellular), GO:0015629 (actin cytoskeleton), GO:0030042 (actin filament depolymerization) |
| Niben101Scf14115g00003.1 | 115 | 47 | -1.15 | 7.43862E-05 | sp\|Q8L7W2\|NUDT8_ARATH | Nudix hydrolase 8 | IPR003293 (Nudix hydrolase 6-like) | NA |
| Niben101Scf03753g00018.1 | 44 | 18 | -1.148 | 0.010016766 | sp\|Q38890\|GUN25_ARATH | Endoglucanase 25 | IPR001701 (Glycoside hydrolase, family 9), IPR008928 (Six-hairpin glycosidase-like) | GO:0003824 (catalytic activity), GO:0004553 (hydrolase activity, hydrolyzing O-glycosyl compounds), GO:0005975 (carbohydrate metabolic process) |
| Niben101Scf01175g03010.1 | 168 | 68 | -1.148 | 2.47462E-07 | sp\|O23144\|PPI1_ARATH | Proton pump-interactor 1 | IPR029669 (Proton pump-interactor) | GO:0005783 (endoplasmic reticulum), GO:0005886 (plasma membrane), GO:0010155 (regulation of proton transport) |
| Niben101Scf07109g04011.1 | 32 | 13 | -1.146 | 0.020262571 | sp\|P01122\|RHO_APLCA | Ras-like GTP-binding protein RHO | IPR001806 (Small GTPase superfamily), IPR005225 (Small GTP-binding protein domain), IPR027417 (P-loop containing nucleoside triphosphate hydrolase) | GO:0005525 (GTP binding), GO:0005622 (intracellular), GO:0006184 (GTP catabolic process), GO:0007165 (signal transduction), GO:0007264 (small GTPase mediated signal transduction), GO:0015031 (protein transport), GO:0016020 (membrane) |
| Niben101Scf04529g00012.1 | 47 | 19 | -1.146 | 0.008228055 | sp\|P48188\|PSBK_MAIZE | Photosystem II reaction center protein K | IPR003686 (Photosystem II PsbI), IPR003687 (Photosystem II PsbK) | GO:0009523 (photosystem II), GO:0009539 (photosystem II reaction center), GO:0015979 (photosynthesis), GO:0016020 (membrane) |
| Niben101Scf01374g04004.1 | 25 | 10 | -1.14 | 0.040514849 | AT3G16060.1 | ATP binding microtubule motor family protein LENGTH=684 | IPR001752 (Kinesin motor domain), IPR027417 (P-loop containing nucleoside triphosphate hydrolase), IPR027640 (Kinesin-like protein) | GO:0003777 (microtubule motor activity), GO:0005524 (ATP binding), GO:0005871 (kinesin complex), GO:0007018 (microtubule-based movement), GO:0008017 (microtubule binding) |
| Niben101Scf07103g01015.1 | 161 | 66 | -1.135 | 0.000562692 | sp\|Q7H8L9\|ATPB_IPOOB | ATP synthase subunit beta, chloroplastic | IPR000194 (ATPase, F1/V1/A1 complex, alpha/beta subunit, nucleotide-binding domain), IPR001469 (ATPase, F1 complex, delta/epsilon subunit), IPR027417 (P-loop containing nucleoside triphosphate hydrolase) | GO:0005524 (ATP binding), GO:0015986 (ATP synthesis coupled proton transport), GO:0045261 (proton-transporting ATP synthase complex, catalytic core F(1)), GO:0046933 (proton-transporting ATP synthase activity, rotational mechanism), GO:0046961 (proton-transporting ATPase activity, rotational mechanism) |
| Niben101Scf11383g00015.1 | 35 | 14 | -1.13 | 0.042381889 | sp\|Q8LC83\|RS242_ARATH | 40S ribosomal protein S24-2 | IPR001976 (Ribosomal protein S24e), IPR012677 (Nucleotide-binding alpha-beta plait domain) | GO:0000166 (nucleotide binding), GO:0003735 (structural constituent of ribosome), GO:0005622 (intracellular), GO:0005840 (ribosome), GO:0006412 (translation) |
| Niben101Scf02572g00016.1 | 62 | 25 | -1.13 | 0.002997636 | sp\|Q9M817\|PTR6_ARATH | Protein NRT1/ PTR FAMILY 1.2 | IPR000109 (Proton-dependent oligopeptide transporter family), IPR020846 (Major facilitator superfamily domain) | GO:0005215 (transporter activity), GO:0006810 (transport), GO:0016020 (membrane) |
| Niben101Scf01696g04006.1 | 30 | 12 | -1.129 | 0.034283145 | sp\|O24407\|IAA16_ARATH | Auxin-responsive protein IAA16 | IPR003311 (AUX/IAA protein) | GO:0005634 (nucleus), GO:0006355 (regulation of transcription, DNA-templated), GO:0046983 (protein dimerization activity) |
| Niben101Scf02736g01011.1 | 34 | 14 | -1.128 | 0.018418767 | sp\|P14009\|14KD_DAUCA | 14 kDa proline-rich protein DC2.15 | NA | NA |
| Niben101Scf12383g01002.1 | 33 | 14 | -1.123 | 0.0374643 | sp\|P49756\|RBM25_HUMAN | RNA-binding protein 25 | IPR002483 (PWI domain), IPR012677 (Nucleotide-binding alpha-beta plait domain) | GO:0000166 (nucleotide binding), GO:0003676 (nucleic acid binding), GO:0006397 (mRNA processing) |
| Niben101Scf12318g00007.1 | 46 | 19 | -1.122 | 0.011206409 | sp\|Q9SV43\|PLP7_ARATH | Patatin-like protein 7 | IPR016035 (Acyl transferase/acyl hydrolase/lysophospholipase) | GO:0008152 (metabolic process) |
| Niben101Scf05071g00011.1 | 91 | 38 | -1.122 | 0.002167064 | sp\|Q7NFI1\|NDHO_GLOVI | NAD(P)H-quinone oxidoreductase subunit O | IPR020905 (NAD(P)H-quinone oxidoreductase subunit O) | GO:0005886 (plasma membrane), GO:0016655 (oxidoreductase activity, acting on NAD(P)H, quinone or similar compound as acceptor), GO:0055114 (oxidation-reduction process) |
| Niben101Scf08422g00003.1 | 7102 | 2944 | -1.122 | 0.000636703 | AT2G33830.2 | Dormancy/auxin associated family protein LENGTH=108 | IPR008406 (Dormancyauxin associated) | NA |
| Niben101Scf01897g05020.1 | 40 | 17 | -1.12 | 0.008993586 | sp\|Q8RXQ1\|TBL35_ARATH | Protein trichome birefringence-like 35 | IPR025846 (PMR5 N-terminal domain), IPR026057 (PC-Esterase) | NA |
| Niben101Scf00753g03018.1 | 389 | 161 | -1.12 | 3.57009E-08 | sp\|Q8RWD0\|COL16_ARATH | Zinc finger protein CONSTANS-LIKE 16 | IPR000315 (Zinc finger, B-box), IPR010402 (CCT domain) | GO:0005515 (protein binding), GO:0005622 (intracellular), GO:0008270 (zinc ion binding) |
| Niben101Scf03927g00002.1 | 35 | 14 | -1.114 | 0.04250718 | sp\|Q65VS6\|SECA_MANSM | Protein translocase subunit SecA | NA | NA |
| Niben101Scf04439g01023.1 | 27 | 11 | -1.113 | 0.049464262 | gb\|KEH40500.1\| | zinc finger CCCH domain protein, putative [Medicago truncatula] | NA | NA |
| Niben101Scf18343g00013.1 | 42 | 18 | -1.107 | 0.030074505 | sp\|O65837\|LCYE_SOLLC | Lycopene epsilon cyclase, chloroplastic | IPR008671 (Lycopene cyclase-type, FAD-binding) | GO:0016117 (carotenoid biosynthetic process), GO:0016705 (oxidoreductase activity, acting on paired donors, with incorporation or reduction of molecular oxygen) |
| Niben101Scf01090g04021.1 | 50 | 21 | -1.107 | 0.009429782 | sp\|Q6Q1P4\|SMC1_ARATH | Structural maintenance of chromosomes protein 1 | IPR003395 (RecF/RecN/SMC, N-terminal), IPR010935 (SMCs flexible hinge), IPR024704 (Structural maintenance of chromosomes protein), IPR027417 (P-loop containing nucleoside triphosphate hydrolase) | GO:0003682 (chromatin binding), GO:0005515 (protein binding), GO:0005524 (ATP binding), GO:0005694 (chromosome), GO:0007064 (mitotic sister chromatid cohesion), GO:0008278 (cohesin complex), GO:0046982 (protein heterodimerization activity), GO:0051276 (chromosome organization) |
| Niben101Scf01980g12012.1 | 63 | 26 | -1.106 | 0.001328066 | sp\|Q9S7L9\|CX6B1_ARATH | Cytochrome c oxidase subunit 6b-1 | IPR003213 (Cytochrome c oxidase, subunit VIb) | GO:0004129 (cytochrome-c oxidase activity), GO:0005739 (mitochondrion) |
| Niben101Scf11084g00011.1 | 38 | 16 | -1.102 | 0.010069307 | sp\|Q9HDW7\|ATC2_SCHPO | Calcium-transporting ATPase 2 | IPR001757 (P-type ATPase), IPR023214 (HAD-like domain), IPR023298 (P-type ATPase, transmembrane domain), IPR024750 (Calcium-transporting P-type ATPase, N-terminal autoinhibitory domain) | GO:0000166 (nucleotide binding), GO:0005388 (calcium-transporting ATPase activity), GO:0005516 (calmodulin binding), GO:0005524 (ATP binding), GO:0016020 (membrane), GO:0016021 (integral component of membrane), GO:0046872 (metal ion binding), GO:0070588 (calcium ion transmembrane transport) |
| Niben101Scf01139g00001.1 | 22 | 9 | -1.096 | 0.047464534 | AT5G65380.1 | MATE efflux family protein LENGTH=486 | IPR002528 (Multi antimicrobial extrusion protein) | GO:0006855 (drug transmembrane transport), GO:0015238 (drug transmembrane transporter activity), GO:0015297 (antiporter activity), GO:0016020 (membrane), GO:0055085 (transmembrane transport) |
| Niben101Scf08341g10005.1 | 30 | 12 | -1.096 | 0.045003062 | AT3G24520.1 | heat shock transcription factor C1 LENGTH=330 | IPR011991 (Winged helix-turn-helix DNA-binding domain), IPR027725 (Heat shock transcription factor family) | GO:0003700 (sequence-specific DNA binding transcription factor activity), GO:0005634 (nucleus), GO:0006355 (regulation of transcription, DNA-templated), GO:0009408 (response to heat), GO:0043565 (sequence-specific DNA binding) |
| Niben101Scf00753g03016.1 | 37 | 15 | -1.091 | 0.013669653 | sp\|Q9XF43\|KCS6_ARATH | 3-ketoacyl-CoA synthase 6 | IPR012392 (Very-long-chain 3-ketoacyl-CoA synthase), IPR016039 (Thiolase-like) | GO:0003824 (catalytic activity), GO:0006633 (fatty acid biosynthetic process), GO:0008152 (metabolic process), GO:0008610 (lipid biosynthetic process), GO:0016020 (membrane), GO:0016747 (transferase activity, transferring acyl groups other than amino-acyl groups) |
| Niben101Scf06347g04035.1 | 312 | 132 | -1.091 | 0.000384523 | sp\|C1DFM2\|DNAJ_AZOVD | Chaperone protein DnaJ | IPR001305 (Heat shock protein DnaJ, cysteine-rich domain) | GO:0031072 (heat shock protein binding), GO:0051082 (unfolded protein binding) |
| Niben101Scf10846g00017.1 | 49 | 21 | -1.09 | 0.005315406 | AT1G65790.1 | receptor kinase 1 LENGTH=843 | IPR011009 (Protein kinase-like domain), IPR021820 (S-locus receptor kinase, C-terminal) | GO:0004672 (protein kinase activity), GO:0004674 (protein serine/threonine kinase activity), GO:0005524 (ATP binding), GO:0006468 (protein phosphorylation), GO:0016772 (transferase activity, transferring phosphorus-containing groups) |
| Niben101Scf09363g00018.1 | 100 | 42 | -1.089 | 0.028391526 | sp\|Q3AUZ9\|CH602_SYNS9 | 60 kDa chaperonin 2 | IPR002423 (Chaperonin Cpn60/TCP-1), IPR027409 (GroEL-like apical domain), IPR027413 (GroEL-like equatorial domain) | GO:0005524 (ATP binding), GO:0005737 (cytoplasm), GO:0006457 (protein folding), GO:0042026 (protein refolding), GO:0044267 (cellular protein metabolic process) |
| Niben101Scf00574g03004.1 | 49 | 21 | -1.087 | 0.012263471 | sp\|P24922\|IF5A2_NICPL | Eukaryotic translation initiation factor 5A-2 | IPR001884 (Translation elongation factor IF5A) | GO:0003723 (RNA binding), GO:0003746 (translation elongation factor activity), GO:0006452 (translational frameshifting), GO:0043022 (ribosome binding), GO:0045901 (positive regulation of translational elongation), GO:0045905 (positive regulation of translational termination) |
| Niben101Scf00369g09013.1 | 65 | 28 | -1.084 | 0.000982481 | sp\|Q6IV77\|TAZ_MACMU | Tafazzin | IPR000872 (Tafazzin) | GO:0006644 (phospholipid metabolic process), GO:0008152 (metabolic process), GO:0016746 (transferase activity, transferring acyl groups) |
| Niben101Scf11337g00009.1 | 37 | 16 | -1.077 | 0.01862756 | sp\|Q9M5P8\|SNAA_SOLTU | Alpha-soluble NSF attachment protein | IPR000744 (NSF attachment protein), IPR001252 (Malate dehydrogenase, active site) | GO:0005515 (protein binding), GO:0006108 (malate metabolic process), GO:0006886 (intracellular protein transport), GO:0016615 (malate dehydrogenase activity), GO:0055114 (oxidation-reduction process) |
| Niben101Scf07247g04003.1 | 113 | 48 | -1.075 | 2.11611E-05 | sp\|O23144\|PPI1_ARATH | Proton pump-interactor 1 | IPR029669 (Proton pump-interactor) | GO:0005783 (endoplasmic reticulum), GO:0005886 (plasma membrane), GO:0010155 (regulation of proton transport) |
| Niben101Scf02637g02013.1 | 65 | 28 | -1.07 | 0.007068351 | sp\|A6UWV8\|RS5_META3 | 30S ribosomal protein S5 | IPR000851 (Ribosomal protein S5), IPR014720 (Double-stranded RNA-binding domain) | GO:0003723 (RNA binding), GO:0003735 (structural constituent of ribosome), GO:0005840 (ribosome), GO:0006412 (translation), GO:0015935 (small ribosomal subunit) |
| Niben101Scf03047g00007.1 | 43 | 19 | -1.069 | 0.026776581 | AT1G11430.1 | plastid developmental protein DAG, putative LENGTH=232 | NA | NA |
| Niben101Scf06831g01009.1 | 125 | 54 | -1.069 | 4.26344E-05 | sp\|P22852\|TBB_POLAG | Tubulin beta chain | IPR000217 (Tubulin), IPR023123 (Tubulin, C-terminal) | GO:0003924 (GTPase activity), GO:0005200 (structural constituent of cytoskeleton), GO:0005525 (GTP binding), GO:0005874 (microtubule), GO:0006184 (GTP catabolic process), GO:0007017 (microtubule-based process), GO:0043234 (protein complex), GO:0051258 (protein polymerization) |
| Niben101Scf03309g08032.1 | 53 | 23 | -1.066 | 0.031205626 | sp\|Q5QNA6\|SCRL2_ORYSJ | SCAR-like protein 2 | IPR028288 (SCAR/WAVE family) | GO:0005856 (cytoskeleton), GO:0030036 (actin cytoskeleton organization) |
| Niben101Scf03287g02002.1 | 64 | 27 | -1.065 | 0.002619009 | sp\|O04006\|PSAH_BRACM | Photosystem I reaction center subunit VI, chloroplastic | IPR004928 (Photosystem I PsaH, reaction centre subunit VI), IPR023833 (Signal peptide, camelysin), IPR029027 (Single alpha-helix domain) | GO:0009522 (photosystem I), GO:0009538 (photosystem I reaction center), GO:0015979 (photosynthesis) |
| Niben101Scf04995g00014.1 | 47 | 20 | -1.064 | 0.009512105 | sp\|Q9LIG2\|RLK6_ARATH | Receptor-like protein kinase | IPR011009 (Protein kinase-like domain) | GO:0004672 (protein kinase activity), GO:0004674 (protein serine/threonine kinase activity), GO:0005524 (ATP binding), GO:0006468 (protein phosphorylation), GO:0016772 (transferase activity, transferring phosphorus-containing groups) |
| Niben101Scf12308g00011.1 | 135 | 57 | -1.063 | 0.022354459 | emb\|CCG48010.1\| | sarcoplasmic reticulum histidine-rich calcium-binding protein precursor, putative, expressed [Triticum aestivum] | NA | NA |
| Niben101Scf02857g01010.1 | 62 | 26 | -1.062 | 0.006723065 | AT1G30270.1 | CBL-interacting protein kinase 23 LENGTH=482 | IPR004041 (NAF domain), IPR011009 (Protein kinase-like domain), IPR028375 (KA1 domain/Ssp2 C-terminal domain) | GO:0004672 (protein kinase activity), GO:0004674 (protein serine/threonine kinase activity), GO:0005524 (ATP binding), GO:0006468 (protein phosphorylation), GO:0007165 (signal transduction), GO:0016772 (transferase activity, transferring phosphorus-containing groups) |
| Niben101Scf03926g00001.1 | 218 | 95 | -1.06 | 0.001599214 | sp\|Q93VH2\|BBD2_ARATH | Bifunctional nuclease 2 | IPR003729 (Bifunctional nuclease domain) | GO:0004518 (nuclease activity) |
| Niben101Scf06344g01021.1 | 56 | 24 | -1.059 | 0.020145295 | sp\|F4HT49\|PABP1_ARATH | Polyadenylate-binding protein 1 | IPR006515 (Polyadenylate binding protein, human types 1, 2, 3, 4), IPR012677 (Nucleotide-binding alpha-beta plait domain) | GO:0000166 (nucleotide binding), GO:0003676 (nucleic acid binding), GO:0003723 (RNA binding) |
| Niben101Scf04793g01008.1 | 39 | 17 | -1.057 | 0.011733462 | sp\|Q8IFW4\|THIOT_DROME | Thioredoxin-T | IPR005746 (Thioredoxin), IPR012336 (Thioredoxin-like fold) | GO:0006662 (glycerol ether metabolic process), GO:0015035 (protein disulfide oxidoreductase activity), GO:0045454 (cell redox homeostasis) |
| Niben101Scf05929g02002.1 | 36 | 16 | -1.056 | 0.01076844 | sp\|Q5RC80\|RBM39_PONAB | RNA-binding protein 39 | IPR012677 (Nucleotide-binding alpha-beta plait domain) | GO:0000166 (nucleotide binding), GO:0003676 (nucleic acid binding) |
| Niben101Scf00820g00005.1 | 56 | 24 | -1.056 | 0.002838954 | AT1G65295.1 | unknown protein; LOCATED IN: endomembrane system; EXPRESSED IN: 22 plant structures; EXPRESSED DURING: 13 growth stages; BEST Arabidopsis thaliana protein match is: unknown protein . LENGTH=115 | NA | NA |
| Niben101Scf01987g00015.1 | 57 | 24 | -1.056 | 0.046972241 | sp\|P27494\|CB23_TOBAC | Chlorophyll a-b binding protein 36, chloroplastic | IPR022796 (Chlorophyll A-B binding protein), IPR023329 (Chlorophyll a/b binding protein domain) | GO:0009765 (photosynthesis, light harvesting), GO:0016020 (membrane) |
| Niben101Scf01463g07019.1 | 42 | 18 | -1.049 | 0.017657757 | sp\|Q9DC07\|LNEBL_MOUSE | LIM zinc-binding domain-containing Nebulette | IPR001781 (Zinc finger, LIM-type) | GO:0008270 (zinc ion binding) |
| Niben101Scf00894g01012.1 | 43 | 19 | -1.047 | 0.011733462 | gb\|ADK97918.1\| | LuxR family transcriptional regulator [Schiedea globosa] | NA | NA |
| Niben101Scf02503g00004.1 | 38 | 16 | -1.046 | 0.029558023 | emb\|CDX81975.1\| | BnaC08g36260D [Brassica napus] | IPR006502 (Protein of unknown function DUF506, plant) | NA |
| Niben101Scf01124g23009.1 | 47 | 21 | -1.039 | 0.01298909 | ref\|XP_002513164.1\| | conserved hypothetical protein [Ricinus communis] gb\|EEF49155.1\| conserved hypothetical protein [Ricinus communis] | NA | NA |
| Niben101Scf00024g04005.1 | 57 | 25 | -1.038 | 0.008072065 | AT1G74740.1 | calcium-dependent protein kinase 30 LENGTH=541 | IPR011009 (Protein kinase-like domain), IPR011992 (EF-hand domain pair) | GO:0004672 (protein kinase activity), GO:0004674 (protein serine/threonine kinase activity), GO:0005509 (calcium ion binding), GO:0005524 (ATP binding), GO:0006468 (protein phosphorylation), GO:0016772 (transferase activity, transferring phosphorus-containing groups) |
| Niben101Scf09107g01004.1 | 51 | 22 | -1.036 | 0.029376188 | sp\|Q9LIB2\|PHS1_ARATH | Alpha-glucan phosphorylase 1 | IPR000811 (Glycosyl transferase, family 35) | GO:0005975 (carbohydrate metabolic process), GO:0008184 (glycogen phosphorylase activity) |
| Niben101Scf00637g05012.1 | 41 | 18 | -1.034 | 0.036209084 | emb\|CDY42132.1\| | BnaC05g40870D [Brassica napus] | IPR019149 (Protein of unknown function DUF2048), IPR029058 (Alpha/Beta hydrolase fold) | NA |
| Niben101Scf07391g00016.1 | 81 | 36 | -1.034 | 0.001379066 | sp\|Q9LYP4\|BAG3_ARATH | BAG family molecular chaperone regulator 3 | IPR003103 (BAG domain), IPR029071 (Ubiquitin-related domain) | GO:0005515 (protein binding), GO:0051087 (chaperone binding) |
| Niben101Scf02041g05016.1 | 88 | 38 | -1.032 | 0.000457361 | AT4G35560.2 | Transducin/WD40 repeat-like superfamily protein LENGTH=1050 | NA | NA |
| Niben101Scf02471g05010.1 | 34 | 15 | -1.03 | 0.030007756 | sp\|Q6IQ97\|RBM41_DANRE | RNA-binding protein 4.1 | IPR012677 (Nucleotide-binding alpha-beta plait domain) | GO:0000166 (nucleotide binding), GO:0003676 (nucleic acid binding) |
| Niben101Scf15224g00007.1 | 127 | 56 | -1.03 | 0.001271262 | AT5G46770.1 | unknown protein; Has 30201 Blast hits to 17322 proteins in 780 species: Archae - 12; Bacteria - 1396; Metazoa - 17338; Fungi - 3422; Plants - 5037; Viruses - 0; Other Eukaryotes - 2996 (source: NCBI BLink). LENGTH=133 | NA | NA |
| Niben101Scf07894g00006.1 | 55 | 24 | -1.026 | 0.003724518 | emb\|CDX94651.1\| | BnaC07g10070D [Brassica napus] | IPR025322 (Protein of unknown function DUF4228, plant) | NA |
| Niben101Scf02910g01040.1 | 60 | 27 | -1.026 | 0.012133009 | sp\|P93788\|REMO_SOLTU | Remorin | IPR005516 (Remorin, C-terminal), IPR005518 (Remorin, N-terminal) | NA |
| Niben101Scf06499g01019.1 | 60 | 26 | -1.02 | 0.006978369 | sp\|Q9C8K7\|FBK21_ARATH | F-box/kelch-repeat protein | IPR001810 (F-box domain), IPR011043 (Galactose oxidase/kelch, beta-propeller), IPR015915 (Kelch-type beta propeller) | GO:0005515 (protein binding) |
| Niben101Scf01694g12011.1 | 38 | 17 | -1.018 | 0.029308011 | AT2G29670.1 | Tetratricopeptide repeat (TPR)-like superfamily protein LENGTH=536 | IPR011990 (Tetratricopeptide-like helical domain) | GO:0005515 (protein binding) |
| Niben101Scf09871g00013.1 | 68 | 30 | -1.015 | 0.003623531 | NA | Unknown protein | NA | NA |
| Niben101Scf10910g00006.1 | 276 | 123 | -1.014 | 8.77902E-08 | AT1G26190.1 | Phosphoribulokinase / Uridine kinase family LENGTH=674 | IPR006082 (Phosphoribulokinase), IPR027417 (P-loop containing nucleoside triphosphate hydrolase) | GO:0005524 (ATP binding), GO:0005975 (carbohydrate metabolic process), GO:0008152 (metabolic process), GO:0008974 (phosphoribulokinase activity), GO:0016301 (kinase activity) |
| Niben101Scf00905g13009.1 | 55 | 24 | -1.01 | 0.017062776 | AT1G03730.1 | unknown protein; BEST Arabidopsis thaliana protein match is: unknown protein . LENGTH=167 | NA | NA |
| Niben101Scf04384g00008.1 | 70 | 31 | -1.006 | 0.003382997 | AT4G27430.1 | COP1-interacting protein 7 LENGTH=1058 | NA | NA |
| Niben101Scf18822g02009.1 | 175 | 77 | -1.002 | 0.000954651 | sp\|O65572\|CCD1_ARATH | Carotenoid 9,10(9',10')-cleavage dioxygenase 1 | IPR004294 (Carotenoid oxygenase) | NA |
| Niben101Scf00927g04014.1 | 44 | 20 | -1.001 | 0.030587024 | AT1G60140.1 | trehalose phosphate synthase LENGTH=861 | IPR001830 (Glycosyl transferase, family 20), IPR006379 (HAD-superfamily hydrolase, subfamily IIB), IPR023214 (HAD-like domain) | GO:0003824 (catalytic activity), GO:0005992 (trehalose biosynthetic process), GO:0008152 (metabolic process) |
| Niben101Scf06361g01017.1 | 54 | 24 | -1.001 | 0.007407961 | sp\|Q9NQG5\|RPR1B_HUMAN | Regulation of nuclear pre-mRNA domain-containing protein 1B | IPR006569 (CID domain), IPR008942 (ENTH/VHS) | NA |
| Niben101Scf03506g03001.1 | 84 | 38 | -0.996 | 0.009598873 | sp\|Q9SSE5\|COL9_ARATH | Zinc finger protein CONSTANS-LIKE 9 | IPR000315 (Zinc finger, B-box), IPR010402 (CCT domain) | GO:0005515 (protein binding), GO:0005622 (intracellular), GO:0008270 (zinc ion binding) |
| Niben101Scf02145g04003.1 | 84 | 38 | -0.995 | 0.004456206 | AT4G04330.1 | Chaperonin-like RbcX protein LENGTH=174 | NA | NA |
| Niben101Scf05009g04012.1 | 236 | 106 | -0.995 | 4.56963E-07 | AT5G02810.1 | pseudo-response regulator 7 LENGTH=727 | IPR010402 (CCT domain), IPR011006 (CheY-like superfamily) | GO:0000160 (phosphorelay signal transduction system), GO:0005515 (protein binding) |
| Niben101Scf03226g04021.1 | 49 | 22 | -0.991 | 0.006119011 | sp\|O82337\|AGP16_ARATH | Arabinogalactan peptide 16 | IPR009424 (Arabinogalactan peptide, AGP) | NA |
| Niben101Scf04801g01029.1 | 57 | 26 | -0.987 | 0.032324726 | AT5G62840.1 | Phosphoglycerate mutase family protein LENGTH=297 | IPR029033 (Histidine phosphatase superfamily) | NA |
| Niben101Scf02639g04006.1 | 51 | 23 | -0.984 | 0.011273286 | sp\|Q9LIG2\|RLK6_ARATH | Receptor-like protein kinase | IPR011009 (Protein kinase-like domain), IPR013320 (Concanavalin A-like lectin/glucanase domain), IPR024788 (Malectin-like carbohydrate-binding domain) | GO:0004672 (protein kinase activity), GO:0004674 (protein serine/threonine kinase activity), GO:0005524 (ATP binding), GO:0006468 (protein phosphorylation), GO:0016772 (transferase activity, transferring phosphorus-containing groups) |
| Niben101Scf08613g02006.1 | 125 | 57 | -0.981 | 0.001048277 | sp\|O65572\|CCD1_ARATH | Carotenoid 9,10(9',10')-cleavage dioxygenase 1 | IPR004294 (Carotenoid oxygenase) | NA |
| Niben101Scf07036g02002.1 | 41 | 19 | -0.98 | 0.019209644 | NA | Unknown protein | NA | NA |
| Niben101Scf15594g00018.1 | 47 | 21 | -0.979 | 0.010253843 | sp\|Q7Z5K2\|WAPL_HUMAN | Wings apart-like protein homolog | IPR016024 (Armadillo-type fold), IPR022771 (Wings apart-like protein) | GO:0005488 (binding) |
| Niben101Scf04158g00001.1 | 59 | 27 | -0.979 | 0.008055758 | AT5G19670.1 | Exostosin family protein LENGTH=610 | IPR004263 (Exostosin-like) | NA |
| Niben101Scf10396g00002.1 | 41 | 19 | -0.978 | 0.042585356 | ref\|NP_001047044.1\| | product [Oryza sativa Japonica Group] | NA | NA |
| Niben101Scf13942g00028.1 | 56 | 26 | -0.978 | 0.011733462 | sp\|C1B027\|RL6_RHOOB | 50S ribosomal protein L6 | IPR000702 (Ribosomal protein L6) | GO:0003735 (structural constituent of ribosome), GO:0005622 (intracellular), GO:0005840 (ribosome), GO:0006412 (translation), GO:0019843 (rRNA binding) |
| Niben101Scf12623g00008.1 | 218 | 100 | -0.978 | 0.000424832 | sp\|B7GIU0\|URK_ANOFW | Uridine kinase | IPR006082 (Phosphoribulokinase), IPR027417 (P-loop containing nucleoside triphosphate hydrolase) | GO:0005524 (ATP binding), GO:0005975 (carbohydrate metabolic process), GO:0008152 (metabolic process), GO:0008974 (phosphoribulokinase activity), GO:0016301 (kinase activity) |
| Niben101Scf01374g06005.1 | 77 | 35 | -0.972 | 0.038563996 | sp\|Q3B5F5\|CH60_PELLD | 60 kDa chaperonin | IPR002423 (Chaperonin Cpn60/TCP-1), IPR027409 (GroEL-like apical domain), IPR027413 (GroEL-like equatorial domain) | GO:0005524 (ATP binding), GO:0005737 (cytoplasm), GO:0006457 (protein folding), GO:0042026 (protein refolding), GO:0044267 (cellular protein metabolic process) |
| Niben101Scf01500g06009.1 | 37 | 17 | -0.969 | 0.049843978 | AT5G47390.1 | myb-like transcription factor family protein LENGTH=365 | IPR009057 (Homeodomain-like) | GO:0003677 (DNA binding), GO:0003682 (chromatin binding) |
| Niben101Scf01856g02002.1 | 87 | 40 | -0.968 | 0.006119011 | AT3G06840.1 | unknown protein; BEST Arabidopsis thaliana protein match is: unknown protein . LENGTH=187 | NA | NA |
| Niben101Scf06091g08011.1 | 59 | 27 | -0.967 | 0.017057487 | sp\|Q9LVZ8\|LOR12_ARATH | Protein LURP-one-related 12 | IPR025659 (Tubby C-terminal-like domain) | NA |
| Niben101Scf09018g00001.1 | 239 | 110 | -0.967 | 5.06529E-05 | sp\|Q3C1H3\|ATPF_NICSY | ATP synthase subunit b, chloroplastic | NA | NA |
| Niben101Scf00724g00001.1 | 273 | 126 | -0.966 | 0.028591545 | sp\|Q9LDE3\|FBK9_ARATH | F-box/kelch-repeat protein | IPR001810 (F-box domain), IPR015915 (Kelch-type beta propeller) | GO:0005515 (protein binding) |
| Niben101Scf02159g01014.1 | 62 | 29 | -0.961 | 0.012305518 | sp\|Q9FMK7\|BT1_ARATH | BTB/POZ and TAZ domain-containing protein 1 | IPR000197 (Zinc finger, TAZ-type), IPR011333 (BTB/POZ fold) | GO:0003712 (transcription cofactor activity), GO:0004402 (histone acetyltransferase activity), GO:0005515 (protein binding), GO:0005634 (nucleus), GO:0006355 (regulation of transcription, DNA-templated), GO:0008270 (zinc ion binding) |
| Niben101Scf01139g01002.1 | 64 | 30 | -0.961 | 0.012549203 | gb\|ABS79378.1\| | At4g40080-like protein [Arabidopsis lyrata subsp. petraea] | IPR008942 (ENTH/VHS), IPR011417 (AP180 N-terminal homology (ANTH) domain) | GO:0005543 (phospholipid binding) |
| Niben101Scf00508g06005.1 | 53 | 25 | -0.958 | 0.003749975 | sp\|Q0WSY2\|FPP4_ARATH | Filament-like plant protein 4 | IPR008587 (Filament-like plant protein) | NA |
| Niben101Scf16022g01005.1 | 44 | 20 | -0.956 | 0.024016409 | NA | Unknown protein | NA | NA |
| Niben101Scf01521g09001.1 | 65 | 30 | -0.956 | 0.022047882 | pdb\|2V0U\|A | Chain A, N- And C-terminal Helices Of Oat Lov2 (404-546) Are Involved In Light-induced Signal Transduction (cryo Dark Structure Of Lov2 (404-546)) Chain A, N- And C-Terminal Helices Of Oat Lov2 (404-546) Are Involved In Light-Induced Signal Transduction (Cryo- Trapped Light Structure Of Lov2 (404-546)) | IPR000014 (PAS domain) | GO:0004871 (signal transducer activity), GO:0007165 (signal transduction) |
| Niben101Scf12919g00016.1 | 147 | 68 | -0.955 | 0.000196212 | sp\|F4HT49\|PABP1_ARATH | Polyadenylate-binding protein 1 | IPR006515 (Polyadenylate binding protein, human types 1, 2, 3, 4), IPR012677 (Nucleotide-binding alpha-beta plait domain) | GO:0000166 (nucleotide binding), GO:0003676 (nucleic acid binding), GO:0003723 (RNA binding) |
| Niben101Scf00161g05002.1 | 1170 | 539 | -0.955 | 1.71725E-05 | sp\|Q8RWE8\|GGAP1_ARATH | GDP-L-galactose phosphorylase 1 | IPR026506 (GDP-L-galactose/GDP-D-glucose phosphorylase) | GO:0080048 (GDP-D-glucose phosphorylase activity) |
| Niben101Scf03905g02008.1 | 65 | 30 | -0.954 | 0.003338548 | sp\|Q8LGG8\|USPAL_ARATH | Universal stress protein A-like protein | IPR006015 (Universal stress protein A) | GO:0006950 (response to stress) |
| Niben101Scf01927g09002.1 | 42 | 20 | -0.953 | 0.045867004 | sp\|Q9FK16\|AGP22_ARATH | Arabinogalactan peptide 22 | IPR009424 (Arabinogalactan peptide, AGP) | NA |
| Niben101Scf34590g00006.1 | 95 | 44 | -0.952 | 0.006119011 | AT1G37130.1 | nitrate reductase 2 LENGTH=917 | IPR001834 (NADH:cytochrome b5 reductase (CBR)), IPR008335 (Eukaryotic molybdopterin oxidoreductase), IPR014756 (Immunoglobulin E-set), IPR017938 (Riboflavin synthase-like beta-barrel) | GO:0005506 (iron ion binding), GO:0009055 (electron carrier activity), GO:0016491 (oxidoreductase activity), GO:0020037 (heme binding), GO:0030151 (molybdenum ion binding), GO:0042128 (nitrate assimilation), GO:0043546 (molybdopterin cofactor binding), GO:0046857 (oxidoreductase activity, acting on other nitrogenous compounds as donors, with NAD or NADP as acceptor), GO:0046872 (metal ion binding), GO:0050660 (flavin adenine dinucleotide binding), GO:0055114 (oxidation-reduction process) |
| Niben101Scf02270g02006.1 | 56 | 26 | -0.946 | 0.033893689 | AT1G51100.1 | unknown protein; FUNCTIONS IN: molecular_function unknown; INVOLVED IN: biological_process unknown; LOCATED IN: chloroplast, chloroplast stroma; EXPRESSED IN: 22 plant structures; EXPRESSED DURING: 13 growth stages; Has 26 Blast hits to 26 proteins in 9 species: Archae - 0; Bacteria - 0; Metazoa - 0; Fungi - 0; Plants - 26; Viruses - 0; Other Eukaryotes - 0 (source: NCBI BLink). LENGTH=211 | NA | NA |
| Niben101Scf13141g02002.1 | 357 | 169 | -0.945 | 4.90226E-05 | sp\|Q9SR92\|STR10_ARATH | Rhodanese-like domain-containing protein 10 | IPR001763 (Rhodanese-like domain) | NA |
| Niben101Scf07165g00002.1 | 79 | 37 | -0.944 | 0.011385964 | sp\|B1XJT1\|RL29_SYNP2 | 50S ribosomal protein L29 | IPR001854 (Ribosomal protein L29) | GO:0003735 (structural constituent of ribosome), GO:0005622 (intracellular), GO:0005840 (ribosome), GO:0006412 (translation) |
| Niben101Scf00823g01006.1 | 88 | 41 | -0.942 | 0.001212639 | sp\|O82256\|COL13_ARATH | Zinc finger protein CONSTANS-LIKE 13 | IPR000315 (Zinc finger, B-box), IPR010402 (CCT domain) | GO:0005515 (protein binding), GO:0005622 (intracellular), GO:0008270 (zinc ion binding) |
| Niben101Scf14076g00012.1 | 80 | 38 | -0.94 | 0.005832076 | dbj\|BAF52858.1\| | repressor of silencing 3 [Nicotiana tabacum] | IPR011257 (DNA glycosylase), IPR023170 (Helix-turn-helix, base-excision DNA repair, C-terminal), IPR028924 (Permuted single zf-CXXC unit), IPR028925 (Demeter, RRM-fold domain) | GO:0003824 (catalytic activity), GO:0006281 (DNA repair), GO:0006284 (base-excision repair) |
| Niben101Scf02807g11016.1 | 170 | 81 | -0.94 | 0.000812998 | emb\|CDY48281.1\| | BnaA05g10730D [Brassica napus] | NA | NA |
| Niben101Scf02982g03019.1 | 91 | 43 | -0.938 | 0.002838954 | AT3G23820.1 | UDP-D-glucuronate 4-epimerase 6 LENGTH=460 | IPR001509 (NAD-dependent epimerase/dehydratase, N-terminal domain), IPR008089 (Nucleotide sugar epimerase) | GO:0003824 (catalytic activity), GO:0005975 (carbohydrate metabolic process), GO:0016857 (racemase and epimerase activity, acting on carbohydrates and derivatives), GO:0050662 (coenzyme binding) |
| Niben101Scf04592g00022.1 | 47 | 22 | -0.935 | 0.038297417 | sp\|Q06AT9\|RBM4B_PIG | RNA-binding protein 4B | IPR012677 (Nucleotide-binding alpha-beta plait domain) | GO:0000166 (nucleotide binding), GO:0003676 (nucleic acid binding) |
| Niben101Scf02459g00024.1 | 56 | 26 | -0.933 | 0.006904671 | sp\|P27518\|CB21_GOSHI | Chlorophyll a-b binding protein 151, chloroplastic | IPR022796 (Chlorophyll A-B binding protein), IPR023329 (Chlorophyll a/b binding protein domain) | GO:0009765 (photosynthesis, light harvesting), GO:0016020 (membrane) |
| Niben101Scf10763g00008.1 | 613 | 289 | -0.933 | 0.000687112 | AT4G27450.1 | Aluminium induced protein with YGL and LRDR motifs LENGTH=250 | IPR024286 (Domain of unknown function DUF3700), IPR029055 (Nucleophile aminohydrolases, N-terminal) | NA |
| Niben101Scf08859g00010.1 | 56 | 26 | -0.929 | 0.006301163 | NA | Unknown protein | NA | NA |
| Niben101Scf02572g03009.1 | 520 | 244 | -0.929 | 0.014185258 | ref\|NP_001172203.1\| | OsJ_00601 [Oryza sativa Japonica Group] dbj\|BAH90933.1\| [Oryza sativa Japonica Group] dbj\|BAN78703.1\| cadmium tolerant 3 [Oryza sativa Japonica Group] | NA | NA |
| Niben101Scf04633g01009.1 | 35 | 17 | -0.928 | 0.026321513 | AT3G05840.2 | Protein kinase superfamily protein LENGTH=409 | IPR011009 (Protein kinase-like domain) | GO:0004672 (protein kinase activity), GO:0004674 (protein serine/threonine kinase activity), GO:0005524 (ATP binding), GO:0006468 (protein phosphorylation), GO:0016772 (transferase activity, transferring phosphorus-containing groups) |
| Niben101Scf08368g02011.1 | 56 | 27 | -0.928 | 0.030007756 | sp\|Q9Y6U7\|RN215_HUMAN | RING finger protein 215 | IPR013083 (Zinc finger, RING/FYVE/PHD-type) | GO:0005515 (protein binding), GO:0008270 (zinc ion binding) |
| Niben101Scf07109g01003.1 | 34 | 16 | -0.927 | 0.044147726 | AT2G20990.3 | synaptotagmin A LENGTH=579 | IPR000008 (C2 domain) | GO:0005515 (protein binding) |
| Niben101Scf05541g02009.1 | 34 | 16 | -0.927 | 0.043380403 | sp\|Q5U5Q3\|MEX3C_HUMAN | RNA-binding E3 ubiquitin-protein ligase MEX3C | IPR013083 (Zinc finger, RING/FYVE/PHD-type) | GO:0005515 (protein binding), GO:0008270 (zinc ion binding) |
| Niben101Scf05634g02005.1 | 90 | 43 | -0.926 | 0.023387937 | sp\|Q949M2\|FABG4_BRANA | 3-oxoacyl-[acyl-carrier-protein] reductase 4 | IPR002198 (Short-chain dehydrogenase/reductase SDR), IPR003560 (2,3-dihydro-2,3-dihydroxybenzoate dehydrogenase) | GO:0008152 (metabolic process), GO:0008667 (2,3-dihydro-2,3-dihydroxybenzoate dehydrogenase activity), GO:0016491 (oxidoreductase activity), GO:0019290 (siderophore biosynthetic process), GO:0055114 (oxidation-reduction process) |
| Niben101Scf08522g02011.1 | 61 | 29 | -0.925 | 0.012710474 | AT4G16340.1 | guanyl-nucleotide exchange factors;GTPase binding;GTP binding LENGTH=1830 | IPR010703 (Dedicator of cytokinesis C-terminal), IPR026791 (Dedicator of cytokinesis), IPR027357 (DHR-2 domain) | GO:0005085 (guanyl-nucleotide exchange factor activity), GO:0007264 (small GTPase mediated signal transduction) |
| Niben101Scf10560g00004.1 | 151 | 72 | -0.924 | 0.000502499 | sp\|Q9LIG2\|RLK6_ARATH | Receptor-like protein kinase | IPR011009 (Protein kinase-like domain), IPR013320 (Concanavalin A-like lectin/glucanase domain) | GO:0004672 (protein kinase activity), GO:0004713 (protein tyrosine kinase activity), GO:0005524 (ATP binding), GO:0006468 (protein phosphorylation), GO:0016772 (transferase activity, transferring phosphorus-containing groups) |
| Niben101Scf08222g02003.1 | 315 | 151 | -0.923 | 0.001154023 | sp\|Q9Z8J2\|TLC1_CHLPN | ADP,ATP carrier protein 1 | IPR004667 (ADP/ATP carrier protein) | GO:0005471 (ATP:ADP antiporter activity), GO:0005524 (ATP binding), GO:0006810 (transport), GO:0016021 (integral component of membrane) |
| Niben101Scf00774g00002.1 | 66 | 31 | -0.92 | 0.006205809 | AT5G58000.1 | Reticulon family protein LENGTH=487 | IPR003388 (Reticulon) | NA |
| Niben101Scf01856g02002.1 | 114 | 54 | -0.919 | 0.000926715 | AT3G06840.1 | unknown protein; BEST Arabidopsis thaliana protein match is: unknown protein . LENGTH=187 | NA | NA |
| Niben101Scf02164g04001.1 | 84 | 40 | -0.918 | 0.005566881 | AT1G70320.1 | ubiquitin-protein ligase 2 LENGTH=3658 | IPR000569 (HECT), IPR003903 (Ubiquitin interacting motif), IPR009060 (UBA-like), IPR010309 (E3 ubiquitin ligase, domain of unknown function DUF908), IPR016024 (Armadillo-type fold), IPR025527 (Domain of unknown function DUF4414) | GO:0004842 (ubiquitin-protein transferase activity), GO:0005488 (binding), GO:0005515 (protein binding) |
| Niben101Scf13360g00001.1 | 59 | 28 | -0.917 | 0.027459737 | emb\|CDX96071.1\| | BnaA07g26600D [Brassica napus] | NA | NA |
| Niben101Scf00577g12010.1 | 152 | 73 | -0.916 | 0.004284544 | sp\|Q8RWD0\|COL16_ARATH | Zinc finger protein CONSTANS-LIKE 16 | IPR000315 (Zinc finger, B-box), IPR010402 (CCT domain) | GO:0005515 (protein binding), GO:0005622 (intracellular), GO:0008270 (zinc ion binding) |
| Niben101Scf01834g02011.1 | 361 | 173 | -0.914 | 2.40994E-05 | AT1G49975.1 | INVOLVED IN: photosynthesis; LOCATED IN: photosystem I, chloroplast, thylakoid membrane; EXPRESSED IN: 20 plant structures; EXPRESSED DURING: 13 growth stages; | IPR008796 (Photosystem I PsaN, reaction centre subunit N) | GO:0005516 (calmodulin binding), GO:0009522 (photosystem I), GO:0015979 (photosynthesis), GO:0042651 (thylakoid membrane) |
| Niben101Scf06189g02007.1 | 43 | 21 | -0.913 | 0.043200782 | AT5G47390.1 | myb-like transcription factor family protein LENGTH=365 | IPR009057 (Homeodomain-like) | GO:0003677 (DNA binding), GO:0003682 (chromatin binding) |
| Niben101Scf02909g06003.1 | 88 | 42 | -0.913 | 0.010881991 | emb\|CDY02362.1\| | BnaA09g25400D [Brassica napus] | NA | NA |
| Niben101Scf06726g00034.1 | 2058 | 995 | -0.913 | 0.001567695 | sp\|Q942D4\|BURP3_ORYSJ | BURP domain-containing protein 3 | IPR004873 (BURP domain) | NA |
| Niben101Scf00539g07015.1 | 209 | 100 | -0.91 | 0.000269379 | sp\|Q42403\|TRXH3_ARATH | Thioredoxin H3 | IPR005746 (Thioredoxin), IPR012336 (Thioredoxin-like fold) | GO:0006662 (glycerol ether metabolic process), GO:0015035 (protein disulfide oxidoreductase activity), GO:0045454 (cell redox homeostasis) |
| Niben101Scf01550g01003.1 | 225 | 108 | -0.907 | 4.26344E-05 | sp\|Q94A65\|STR14_ARATH | Rhodanese-like domain-containing protein 14, chloroplastic | IPR001763 (Rhodanese-like domain) | NA |
| Niben101Scf01339g05012.1 | 43 | 21 | -0.906 | 0.030355308 | sp\|Q9VQG2\|APH1_DROME | Gamma-secretase subunit Aph-1 | IPR009294 (Gamma-secretase subunit Aph-1) | GO:0016021 (integral component of membrane), GO:0016485 (protein processing), GO:0043085 (positive regulation of catalytic activity) |
| Niben101Scf02211g03005.1 | 70 | 34 | -0.905 | 0.002298943 | sp\|Q9C8K7\|FBK21_ARATH | F-box/kelch-repeat protein | IPR000014 (PAS domain), IPR001610 (PAC motif), IPR001810 (F-box domain), IPR015915 (Kelch-type beta propeller) | GO:0004871 (signal transducer activity), GO:0005515 (protein binding), GO:0007165 (signal transduction) |
| Niben101Scf07765g01010.1 | 103 | 50 | -0.892 | 0.01677562 | sp\|P60039\|RL73_ARATH | 60S ribosomal protein L7-3 | IPR005998 (Ribosomal protein L7, eukaryotic) | NA |
| Niben101Scf02805g02010.1 | 57 | 27 | -0.887 | 0.043380183 | sp\|O23144\|PPI1_ARATH | Proton pump-interactor 1 | NA | NA |
| Niben101Scf01346g00001.1 | 67 | 32 | -0.887 | 0.005053075 | AT4G11400.1 | ARID/BRIGHT DNA-binding domain;ELM2 domain protein LENGTH=573 | NA | NA |
| Niben101Scf00046g00002.1 | 54 | 26 | -0.886 | 0.042780377 | AT3G27580.1 | Protein kinase superfamily protein LENGTH=578 | IPR011009 (Protein kinase-like domain) | GO:0004672 (protein kinase activity), GO:0004674 (protein serine/threonine kinase activity), GO:0005524 (ATP binding), GO:0006468 (protein phosphorylation), GO:0016772 (transferase activity, transferring phosphorus-containing groups) |
| Niben101Scf08855g00008.1 | 303 | 144 | -0.886 | 0.03130228 | AT1G73260.1 | kunitz trypsin inhibitor 1 LENGTH=215 | IPR002160 (Proteinase inhibitor I3, Kunitz legume) | GO:0004866 (endopeptidase inhibitor activity) |
| Niben101Scf01927g04007.1 | 75 | 37 | -0.884 | 0.037977708 | ref\|XP_002514207.1\| | conserved hypothetical protein [Ricinus communis] gb\|EEF48161.1\| conserved hypothetical protein [Ricinus communis] | NA | NA |
| Niben101Scf01212g01017.1 | 45 | 22 | -0.881 | 0.036383459 | sp\|A4QLH5\|PSBA_LOBMA | Photosystem Q(B) protein | IPR000484 (Photosynthetic reaction centre, L/M) | GO:0009772 (photosynthetic electron transport in photosystem II), GO:0019684 (photosynthesis, light reaction), GO:0045156 (electron transporter, transferring electrons within the cyclic electron transport pathway of photosynthesis activity) |
| Niben101Scf07054g04012.1 | 102 | 50 | -0.881 | 0.004118614 | sp\|Q6YW64\|DRB4_ORYSJ | Double-stranded RNA-binding protein 4 | IPR014720 (Double-stranded RNA-binding domain), IPR026939 (At2g23090 like) | NA |
| Niben101Scf06249g05016.1 | 82 | 40 | -0.879 | 0.023538201 | sp\|P27521\|CA4_ARATH | Chlorophyll a-b binding protein 4, chloroplastic | IPR022796 (Chlorophyll A-B binding protein), IPR023329 (Chlorophyll a/b binding protein domain) | GO:0009765 (photosynthesis, light harvesting), GO:0016020 (membrane) |
| Niben101Scf33781g00006.1 | 84 | 41 | -0.878 | 0.045776188 | gb\|KEH17310.1\| | O-acyltransferase WSD1-like protein [Medicago truncatula] | IPR009721 (O-acyltransferase, WSD1, C-terminal) | GO:0004144 (diacylglycerol O-acyltransferase activity) |
| Niben101Scf02994g03003.1 | 57 | 28 | -0.877 | 0.032081427 | ref\|WP_024571788.1\| | copper oxidase [Bacillus subtilis] | IPR008972 (Cupredoxin) | GO:0005507 (copper ion binding), GO:0016491 (oxidoreductase activity), GO:0055114 (oxidation-reduction process) |
| Niben101Scf03844g01002.1 | 126 | 62 | -0.877 | 0.000737566 | AT1G10940.2 | Protein kinase superfamily protein LENGTH=371 | IPR011009 (Protein kinase-like domain) | GO:0004672 (protein kinase activity), GO:0004674 (protein serine/threonine kinase activity), GO:0005524 (ATP binding), GO:0006468 (protein phosphorylation), GO:0016772 (transferase activity, transferring phosphorus-containing groups) |
| Niben101Scf12858g00003.1 | 102 | 50 | -0.87 | 0.000449914 | sp\|P93332\|NOD3_MEDTR | Bidirectional sugar transporter N3 | NA | NA |
| Niben101Scf08208g02004.1 | 53 | 26 | -0.862 | 0.030355308 | gb\|KEH37079.1\| | maternal effect embryo arrest protein [Medicago truncatula] | NA | NA |
| Niben101Scf04044g19020.1 | 68 | 34 | -0.853 | 0.02497941 | ref\|XP_007013731.1\| | Enhancer of polycomb-like transcription factor protein, putative isoform 5 [Theobroma cacao] gb\|EOY31350.1\| Enhancer of polycomb-like transcription factor protein, putative isoform 5 [Theobroma cacao] | NA | NA |
| Niben101Scf00341g02005.1 | 42 | 21 | -0.85 | 0.03985301 | AT5G55660.1 | DEK domain-containing chromatin associated protein LENGTH=778 | IPR009057 (Homeodomain-like), IPR014876 (DEK, C-terminal) | GO:0003677 (DNA binding) |
| Niben101Scf11137g00006.1 | 52 | 26 | -0.85 | 0.043241003 | AT5G58080.1 | response regulator 18 LENGTH=618 | IPR009057 (Homeodomain-like) | GO:0003677 (DNA binding), GO:0003682 (chromatin binding) |
| Niben101Scf08806g01007.1 | 92 | 46 | -0.85 | 0.003632227 | emb\|CDX96071.1\| | BnaA07g26600D [Brassica napus] | NA | NA |
| Niben101Scf00050g00008.1 | 45 | 22 | -0.849 | 0.045605528 | sp\|P01122\|RHO_APLCA | Ras-like GTP-binding protein RHO | IPR001806 (Small GTPase superfamily), IPR005225 (Small GTP-binding protein domain), IPR027417 (P-loop containing nucleoside triphosphate hydrolase) | GO:0005525 (GTP binding), GO:0005622 (intracellular), GO:0006184 (GTP catabolic process), GO:0007165 (signal transduction), GO:0007264 (small GTPase mediated signal transduction), GO:0015031 (protein transport), GO:0016020 (membrane) |
| Niben101Scf02290g07003.1 | 69 | 35 | -0.849 | 0.006548995 | sp\|O35685\|NUDC_MOUSE | Nuclear migration protein nudC | IPR008978 (HSP20-like chaperone) | NA |
| Niben101Scf33299g00003.1 | 98 | 49 | -0.849 | 0.016117738 | gb\|AFX95943.1\| | green-ripe like protein 1 [Solanum melongena] | IPR008496 (Protein of unknown function DUF778) | NA |
| Niben101Scf00784g00007.1 | 244 | 122 | -0.849 | 8.31342E-05 | emb\|CDY60156.1\| | BnaAnng16520D [Brassica napus] | IPR007033 (Transcriptional activator, plants) | NA |
| Niben101Scf04561g00005.1 | 88 | 44 | -0.847 | 0.041078102 | sp\|P10708\|CB12_SOLLC | Chlorophyll a-b binding protein 7, chloroplastic | IPR022796 (Chlorophyll A-B binding protein), IPR023329 (Chlorophyll a/b binding protein domain) | GO:0009765 (photosynthesis, light harvesting), GO:0016020 (membrane) |
| Niben101Scf01635g05017.1 | 100 | 51 | -0.846 | 0.028779504 | AT5G57660.1 | CONSTANS-like 5 LENGTH=355 | IPR000315 (Zinc finger, B-box), IPR010402 (CCT domain) | GO:0005515 (protein binding), GO:0005622 (intracellular), GO:0008270 (zinc ion binding) |
| Niben101Scf00046g04007.1 | 67 | 34 | -0.844 | 0.016262126 | sp\|Q9T0D6\|PP306_ARATH | Pentatricopeptide repeat-containing protein | IPR002885 (Pentatricopeptide repeat) | NA |
| Niben101Scf02118g00004.1 | 242 | 123 | -0.843 | 0.008738994 | AT1G10940.2 | Protein kinase superfamily protein LENGTH=371 | IPR011009 (Protein kinase-like domain) | GO:0004672 (protein kinase activity), GO:0004674 (protein serine/threonine kinase activity), GO:0005524 (ATP binding), GO:0006468 (protein phosphorylation), GO:0016772 (transferase activity, transferring phosphorus-containing groups) |
| Niben101Scf00324g02004.1 | 43 | 21 | -0.84 | 0.040139764 | sp\|Q9SN58\|UGE5_ARATH | UDP-glucose 4-epimerase 5 | IPR001509 (NAD-dependent epimerase/dehydratase, N-terminal domain), IPR005886 (UDP-glucose 4-epimerase GalE), IPR008089 (Nucleotide sugar epimerase), IPR025308 (UDP-glucose 4-epimerase C-terminal domain) | GO:0003824 (catalytic activity), GO:0003978 (UDP-glucose 4-epimerase activity), GO:0005975 (carbohydrate metabolic process), GO:0006012 (galactose metabolic process), GO:0016857 (racemase and epimerase activity, acting on carbohydrates and derivatives), GO:0050662 (coenzyme binding) |
| Niben101Scf08156g09004.1 | 169 | 85 | -0.84 | 7.96259E-05 | sp\|Q4IB70\|CWC21_GIBZE | Pre-mRNA-splicing factor CWC21 | IPR013170 (mRNA splicing factor, Cwf21) | NA |
| Niben101Scf05689g00008.1 | 52 | 26 | -0.839 | 0.030475301 | ref\|XP_002510763.1\| | conserved hypothetical protein [Ricinus communis] gb\|EEF52950.1\| conserved hypothetical protein [Ricinus communis] | NA | NA |
| Niben101Scf11844g01001.1 | 78 | 39 | -0.839 | 0.007407961 | ref\|WP_018199143.1\| | MULTISPECIES: hypothetical protein [Hydrogenedentes] | IPR025150 (Domain of unknown function DUF4091) | NA |
| Niben101Scf03503g01006.1 | 103 | 52 | -0.839 | 0.002221524 | sp\|Q6Z4I3\|TRH21_ORYSJ | Thioredoxin H2-1 | IPR005746 (Thioredoxin), IPR012336 (Thioredoxin-like fold) | GO:0006662 (glycerol ether metabolic process), GO:0015035 (protein disulfide oxidoreductase activity), GO:0045454 (cell redox homeostasis) |
| Niben101Scf03076g02006.1 | 80 | 41 | -0.837 | 0.018103658 | sp\|Q9LYT2\|PP287_ARATH | Pentatricopeptide repeat-containing protein | IPR002885 (Pentatricopeptide repeat), IPR011990 (Tetratricopeptide-like helical domain) | GO:0005515 (protein binding) |
| Niben101Scf00568g04011.1 | 221 | 111 | -0.837 | 0.000449914 | sp\|B2LMI9\|PSBC_GUIAB | Photosystem II CP43 reaction center protein | IPR000484 (Photosynthetic reaction centre, L/M), IPR000932 (Photosystem antenna protein-like) | GO:0009521 (photosystem), GO:0009523 (photosystem II), GO:0009767 (photosynthetic electron transport chain), GO:0009772 (photosynthetic electron transport in photosystem II), GO:0015979 (photosynthesis), GO:0016020 (membrane), GO:0016168 (chlorophyll binding), GO:0019684 (photosynthesis, light reaction), GO:0030076 (light-harvesting complex), GO:0045156 (electron transporter, transferring electrons within the cyclic electron transport pathway of photosynthesis activity) |
| Niben101Scf15054g01001.1 | 250 | 126 | -0.833 | 0.000316272 | emb\|CDX72133.1\| | BnaC08g27410D [Brassica napus] | NA | NA |
| Niben101Scf05811g00003.1 | 333 | 170 | -0.83 | 0.000718935 | gb\|ADK13066.1\| | conserved hypothetical protein 9 [Hevea brasiliensis] | NA | NA |
| Niben101Scf18637g02010.1 | 63 | 32 | -0.826 | 0.040458236 | AT5G65630.1 | global transcription factor group E7 LENGTH=590 | IPR027353 (NET domain) | NA |
| Niben101Scf01779g00009.1 | 72 | 36 | -0.824 | 0.040781311 | sp\|P0CM95\|CWC21_CRYNB | Pre-mRNA-splicing factor CWC21 | IPR013170 (mRNA splicing factor, Cwf21) | NA |
| Niben101Scf00519g00010.1 | 131 | 67 | -0.821 | 0.007407961 | sp\|P74061\|RPE_SYNY3 | Ribulose-phosphate 3-epimerase | IPR000056 (Ribulose-phosphate 3-epimerase-like), IPR013785 (Aldolase-type TIM barrel) | GO:0003824 (catalytic activity), GO:0005975 (carbohydrate metabolic process), GO:0008152 (metabolic process), GO:0016857 (racemase and epimerase activity, acting on carbohydrates and derivatives) |
| Niben101Scf04847g02012.1 | 54 | 27 | -0.82 | 0.043701093 | sp\|O45766\|CWC15_CAEEL | Protein CWC15 homolog | IPR006973 (Pre-mRNA-splicing factor Cwf15/Cwc15) | GO:0000398 (mRNA splicing, via spliceosome), GO:0005681 (spliceosomal complex) |
| Niben101Scf00672g04011.1 | 50 | 26 | -0.817 | 0.047267818 | sp\|B0R0I6\|CHD8_DANRE | Chromodomain-helicase-DNA-binding protein 8 | IPR000330 (SNF2-related), IPR001650 (Helicase, C-terminal), IPR014012 (Helicase/SANT-associated, DNA binding), IPR027417 (P-loop containing nucleoside triphosphate hydrolase), IPR029295 (Snf2 ATP coupling domain) | GO:0003677 (DNA binding), GO:0005524 (ATP binding), GO:0042393 (histone binding) |
| Niben101Scf02316g03010.1 | 258 | 133 | -0.812 | 0.00409161 | sp\|Q84WM9\|EF1B1_ARATH | Elongation factor 1-beta 1 | IPR014717 (Translation elongation factor EF1B/ribosomal protein S6) | GO:0003746 (translation elongation factor activity), GO:0005853 (eukaryotic translation elongation factor 1 complex), GO:0006414 (translational elongation) |
| Niben101Scf00683g03020.1 | 62975 | 32231 | -0.809 | 3.61436E-07 | sp\|Q71KN3\|CYB6_KLEBI | Cytochrome b6 | IPR016174 (Di-haem cytochrome, transmembrane), IPR023530 (Cytochrome b6, PetB), IPR027387 (Cytochrome b/b6-like domain) | GO:0005506 (iron ion binding), GO:0009055 (electron carrier activity), GO:0016020 (membrane), GO:0016021 (integral component of membrane), GO:0016491 (oxidoreductase activity), GO:0020037 (heme binding), GO:0022900 (electron transport chain), GO:0022904 (respiratory electron transport chain), GO:0045158 (electron transporter, transferring electrons within cytochrome b6/f complex of photosystem II activity) |
| Niben101Scf03939g01001.1 | 181 | 93 | -0.808 | 0.000388799 | AT5G62570.2 | Calmodulin binding protein-like LENGTH=494 | IPR012416 (Calmodulin binding protein-like) | NA |
| Niben101Scf00338g06011.1 | 68 | 35 | -0.805 | 0.007905853 | sp\|P42423\|YXDL_BACSU | ABC transporter ATP-binding protein YxdL | IPR003439 (ABC transporter-like), IPR013525 (ABC-2 type transporter), IPR027417 (P-loop containing nucleoside triphosphate hydrolase) | GO:0005524 (ATP binding), GO:0016020 (membrane), GO:0016887 (ATPase activity) |
| Niben101Scf02927g04002.1 | 97 | 50 | -0.802 | 0.006119011 | sp\|P10639\|THIO_MOUSE | Thioredoxin | IPR005746 (Thioredoxin), IPR012336 (Thioredoxin-like fold) | GO:0006662 (glycerol ether metabolic process), GO:0015035 (protein disulfide oxidoreductase activity), GO:0045454 (cell redox homeostasis) |
| Niben101Scf03732g00001.1 | 137 | 71 | -0.802 | 0.013669653 | sp\|Q1JQE0\|SNW1_BOVIN | SNW domain-containing protein 1 | IPR017862 (SKI-interacting protein, SKIP) | GO:0000398 (mRNA splicing, via spliceosome), GO:0005681 (spliceosomal complex) |
| Niben101Scf02759g01012.1 | 360 | 187 | -0.801 | 0.020004803 | ref\|XP_002532850.1\| | nucleic acid binding protein, putative [Ricinus communis] gb\|EEF29528.1\| nucleic acid binding protein, putative [Ricinus communis] | IPR001878 (Zinc finger, CCHC-type) | GO:0003676 (nucleic acid binding), GO:0008270 (zinc ion binding) |
| Niben101Scf01240g03014.1 | 127 | 66 | -0.798 | 0.000965603 | sp\|Q86JD4\|SEC11_DICDI | Signal peptidase complex catalytic subunit sec11 | IPR001733 (Peptidase S26B, eukaryotic signal peptidase), IPR015927 (Peptidase S24/S26A/S26B/S26C), IPR028360 (Peptidase S24/S26, beta-ribbon domain) | GO:0006465 (signal peptide processing), GO:0008233 (peptidase activity), GO:0016020 (membrane) |
| Niben101Scf08723g02015.1 | 175 | 90 | -0.797 | 0.004081588 | sp\|P84981\|FENR_POPEU | Ferredoxin--NADP reductase, chloroplastic | IPR001433 (Oxidoreductase FAD/NAD(P)-binding), IPR015701 (Ferredoxin--NADP reductase), IPR017938 (Riboflavin synthase-like beta-barrel) | GO:0016491 (oxidoreductase activity), GO:0055114 (oxidation-reduction process) |
| Niben101Scf11137g01017.1 | 60 | 31 | -0.794 | 0.033538421 | gb\|ADY38784.1\| | sequence-specific DNA-binding transcription factor [Coffea arabica] | IPR007759 (DNA-directed RNA polymerase delta subunit/Asxl), IPR009057 (Homeodomain-like), IPR018501 (DDT domain superfamily), IPR028942 (WHIM1 domain) | GO:0003677 (DNA binding), GO:0006351 (transcription, DNA-templated), GO:0006355 (regulation of transcription, DNA-templated) |
| Niben101Scf06249g04003.1 | 117 | 61 | -0.793 | 0.019231189 | AT3G21270.1 | DOF zinc finger protein 2 LENGTH=204 | IPR003851 (Zinc finger, Dof-type) | GO:0003677 (DNA binding), GO:0006355 (regulation of transcription, DNA-templated) |
| Niben101Scf05080g01011.1 | 70 | 36 | -0.79 | 0.013523048 | sp\|Q9FLP0\|MA651_ARATH | 65-kDa microtubule-associated protein 1 | IPR007145 (Microtubule-associated protein, MAP65/Ase1/PRC1) | GO:0000226 (microtubule cytoskeleton organization), GO:0000910 (cytokinesis), GO:0008017 (microtubule binding) |
| Niben101Scf01399g00014.1 | 131 | 68 | -0.787 | 0.006555709 | sp\|A4SIE1\|GLGA_AERS4 | Glycogen synthase | IPR001296 (Glycosyl transferase, family 1) | GO:0009058 (biosynthetic process) |
| Niben101Scf03029g00005.1 | 76 | 40 | -0.786 | 0.026626802 | sp\|P21335\|TADA_BACSU | tRNA-specific adenosine deaminase | IPR016193 (Cytidine deaminase-like), IPR028883 (tRNA-specific adenosine deaminase) | GO:0002100 (tRNA wobble adenosine to inosine editing), GO:0003824 (catalytic activity), GO:0008251 (tRNA-specific adenosine deaminase activity), GO:0008270 (zinc ion binding), GO:0016787 (hydrolase activity) |
| Niben101Scf04386g08002.1 | 82 | 43 | -0.785 | 0.019792908 | sp\|Q9R210\|TFEB_MOUSE | Transcription factor EB | IPR011598 (Myc-type, basic helix-loop-helix (bHLH) domain) | GO:0046983 (protein dimerization activity) |
| Niben101Scf01976g00012.1 | 97 | 51 | -0.784 | 0.033176026 | sp\|Q8GZA8\|ULT1_ARATH | Protein ULTRAPETALA 1 | NA | NA |
| Niben101Scf16714g00001.1 | 797 | 418 | -0.784 | 0.006641103 | sp\|Q38825\|IAA7_ARATH | Auxin-responsive protein IAA7 | IPR003311 (AUX/IAA protein) | GO:0005634 (nucleus), GO:0006355 (regulation of transcription, DNA-templated), GO:0046983 (protein dimerization activity) |
| Niben101Scf22811g01002.1 | 54 | 28 | -0.782 | 0.027790323 | gb\|KEH41854.1\| | START-2 domain protein [Medicago truncatula] | IPR023393 (START-like domain) | NA |
| Niben101Scf05017g06019.1 | 91 | 48 | -0.781 | 0.026644969 | sp\|Q4V0I4\|F16PA_XANC8 | Fructose-1,6-bisphosphatase class 1 | IPR000146 (Fructose-1,6-bisphosphatase class 1/Sedoheputulose-1,7-bisphosphatase) | GO:0005975 (carbohydrate metabolic process), GO:0042132 (fructose 1,6-bisphosphate 1-phosphatase activity), GO:0042578 (phosphoric ester hydrolase activity) |
| Niben101Scf00647g02002.1 | 179 | 94 | -0.781 | 0.000160401 | gb\|KEH21046.1\| | myosin heavy chain-like protein [Medicago truncatula] | IPR019448 (EEIG1/EHBP1 N-terminal domain) | NA |
| Niben101Scf04233g00010.1 | 56 | 29 | -0.776 | 0.046676126 | NA | Unknown protein | NA | NA |
| Niben101Scf02852g03008.1 | 83 | 44 | -0.776 | 0.017300533 | ref\|XP_003610227.1\| | Ycf68 [Medicago truncatula] | NA | NA |
| Niben101Scf02868g02003.1 | 213 | 111 | -0.775 | 0.005053075 | sp\|Q940T9\|COL4_ARATH | Zinc finger protein CONSTANS-LIKE 4 | IPR000315 (Zinc finger, B-box), IPR010402 (CCT domain) | GO:0005515 (protein binding), GO:0005622 (intracellular), GO:0008270 (zinc ion binding) |
| Niben101Scf02864g07005.1 | 114 | 60 | -0.774 | 0.003849161 | sp\|Q9BX46\|RBM24_HUMAN | RNA-binding protein 24 | IPR012677 (Nucleotide-binding alpha-beta plait domain) | GO:0000166 (nucleotide binding), GO:0003676 (nucleic acid binding) |
| Niben101Scf11178g01003.1 | 119 | 62 | -0.773 | 0.046562977 | sp\|P31562\|ACCD_CUSRE | Acetyl-coenzyme A carboxylase carboxyl transferase subunit beta | IPR000438 (Acetyl-CoA carboxylase carboxyl transferase, beta subunit), IPR029045 (ClpP/crotonase-like domain) | GO:0003989 (acetyl-CoA carboxylase activity), GO:0006633 (fatty acid biosynthetic process), GO:0009317 (acetyl-CoA carboxylase complex), GO:0016874 (ligase activity) |
| Niben101Scf02658g01007.1 | 330 | 177 | -0.772 | 0.022354459 | gb\|ACM43526.1\| | phosphatidylinositol-specific phospholipase c [Listeria monocytogenes] | IPR017946 (PLC-like phosphodiesterase, TIM beta/alpha-barrel domain) | GO:0006629 (lipid metabolic process), GO:0008081 (phosphoric diester hydrolase activity) |
| Niben101Scf04629g02001.1 | 371 | 197 | -0.77 | 0.013042628 | sp\|Q93VH2\|BBD2_ARATH | Bifunctional nuclease 2 | IPR003729 (Bifunctional nuclease domain) | GO:0004518 (nuclease activity) |
| Niben101Scf09123g00004.1 | 93 | 49 | -0.767 | 0.032081427 | sp\|B0TI32\|ACCA_HELMI | Acetyl-coenzyme A carboxylase carboxyl transferase subunit alpha | IPR001095 (Acetyl-CoA carboxylase, alpha subunit), IPR011763 (Acetyl-coenzyme A carboxyltransferase, C-terminal), IPR029045 (ClpP/crotonase-like domain) | GO:0003989 (acetyl-CoA carboxylase activity), GO:0006633 (fatty acid biosynthetic process), GO:0009317 (acetyl-CoA carboxylase complex), GO:0016874 (ligase activity) |
| Niben101Scf02757g01001.1 | 142 | 76 | -0.763 | 0.036167029 | AT4G29060.1 | elongation factor Ts family protein LENGTH=953 | IPR001816 (Translation elongation factor EFTs/EF1B), IPR012340 (Nucleic acid-binding, OB-fold) | GO:0003676 (nucleic acid binding), GO:0003746 (translation elongation factor activity), GO:0005515 (protein binding), GO:0005622 (intracellular), GO:0006414 (translational elongation) |
| Niben101Scf03488g08009.1 | 54 | 29 | -0.761 | 0.043321513 | NA | Unknown protein | NA | NA |
| Niben101Scf19423g00008.1 | 200 | 107 | -0.749 | 0.011212582 | sp\|A4QJI0\|PSBA_AETGR | Photosystem Q(B) protein | IPR000484 (Photosynthetic reaction centre, L/M) | GO:0009772 (photosynthetic electron transport in photosystem II), GO:0019684 (photosynthesis, light reaction), GO:0045156 (electron transporter, transferring electrons within the cyclic electron transport pathway of photosynthesis activity) |
| Niben101Scf14009g02003.1 | 94 | 51 | -0.748 | 0.006949852 | ref\|XP_002521463.1\| | Microtubule-associated protein futsch, putative [Ricinus communis] gb\|EEF40953.1\| Microtubule-associated protein futsch, putative [Ricinus communis] | NA | NA |
| Niben101Scf03770g00004.1 | 59 | 32 | -0.746 | 0.025328783 | sp\|Q93Z08\|E136_ARATH | Glucan endo-1,3-beta-glucosidase 6 | IPR000490 (Glycoside hydrolase, family 17), IPR012946 (X8 domain), IPR017853 (Glycoside hydrolase superfamily) | GO:0004553 (hydrolase activity, hydrolyzing O-glycosyl compounds), GO:0005975 (carbohydrate metabolic process) |
| Niben101Scf09321g02026.1 | 81 | 43 | -0.746 | 0.038104386 | sp\|Q9LIG2\|RLK6_ARATH | Receptor-like protein kinase | IPR011009 (Protein kinase-like domain), IPR013320 (Concanavalin A-like lectin/glucanase domain), IPR014729 (Rossmann-like alpha/beta/alpha sandwich fold) | GO:0004672 (protein kinase activity), GO:0005524 (ATP binding), GO:0006468 (protein phosphorylation), GO:0016772 (transferase activity, transferring phosphorus-containing groups) |
| Niben101Scf01520g08031.1 | 138 | 74 | -0.745 | 0.0042229 | AT4G39210.1 | Glucose-1-phosphate adenylyltransferase family protein LENGTH=521 | IPR011831 (Glucose-1-phosphate adenylyltransferase), IPR029044 (Nucleotide-diphospho-sugar transferases) | GO:0005978 (glycogen biosynthetic process), GO:0008878 (glucose-1-phosphate adenylyltransferase activity), GO:0009058 (biosynthetic process), GO:0016779 (nucleotidyltransferase activity) |
| Niben101Scf02576g00011.1 | 335 | 182 | -0.733 | 0.000364519 | AT2G37180.1 | Aquaporin-like superfamily protein LENGTH=285 | IPR000425 (Major intrinsic protein), IPR023271 (Aquaporin-like) | GO:0005215 (transporter activity), GO:0006810 (transport), GO:0016020 (membrane) |
| Niben101Scf11737g00029.1 | 179 | 97 | -0.732 | 0.043241003 | sp\|Q9C9K2\|CP123_ARATH | Calvin cycle protein CP12-3, chloroplastic | IPR003823 (Domain of unknown function CP12) | NA |
| Niben101Scf03893g02007.1 | 552 | 296 | -0.729 | 0.049414698 | sp\|Q9SSK5\|MLP43_ARATH | MLP-like protein 43 | IPR000916 (Bet v I domain), IPR023393 (START-like domain) | GO:0006952 (defense response), GO:0009607 (response to biotic stimulus) |
| Niben101Scf05375g01004.1 | 69 | 37 | -0.728 | 0.037303049 | sp\|F4I2J8\|CATIN_ARATH | Cactin | IPR018816 (Cactin, central domain), IPR019134 (Cactin, C-terminal) | GO:0005515 (protein binding) |
| Niben101Scf04113g07011.1 | 252 | 138 | -0.727 | 0.023538201 | ref\|NP_001051767.1\| | product [Oryza sativa Japonica Group] | NA | NA |
| Niben101Scf07494g00009.1 | 271 | 147 | -0.726 | 0.00021923 | NA | Unknown protein | NA | NA |
| Niben101Scf02583g06003.1 | 333 | 181 | -0.726 | 0.041080667 | sp\|P84981\|FENR_POPEU | Ferredoxin--NADP reductase, chloroplastic | IPR001433 (Oxidoreductase FAD/NAD(P)-binding), IPR015701 (Ferredoxin--NADP reductase), IPR017938 (Riboflavin synthase-like beta-barrel) | GO:0016491 (oxidoreductase activity), GO:0055114 (oxidation-reduction process) |
| Niben101Scf05717g03015.1 | 76 | 41 | -0.724 | 0.020004803 | AT3G08600.1 | Protein of unknown function (DUF1191) LENGTH=316 | IPR010605 (Protein of unknown function DUF1191), IPR013320 (Concanavalin A-like lectin/glucanase domain) | NA |
| Niben101Scf01847g07003.1 | 62 | 34 | -0.723 | 0.049805708 | sp\|Q41009\|TOC34_PEA | Translocase of chloroplast 34 | IPR005690 (Chloroplast protein import component Toc86/159), IPR024283 (Domain of unknown function DUF3406, chloroplast translocase), IPR027417 (P-loop containing nucleoside triphosphate hydrolase) | GO:0005525 (GTP binding), GO:0016817 (hydrolase activity, acting on acid anhydrides) |
| Niben101Scf01036g08009.1 | 455 | 247 | -0.721 | 0.001912481 | sp\|Q9T076\|ENL2_ARATH | Early nodulin-like protein 2 | IPR008972 (Cupredoxin) | GO:0009055 (electron carrier activity) |
| Niben101Scf01781g00009.1 | 92 | 50 | -0.712 | 0.024270484 | sp\|Q8FV29\|YIDC_BRUSU | Membrane protein insertase YidC | IPR001708 (Membrane insertase OXA1/ALB3/YidC), IPR028055 (Membrane insertase YidC/Oxa1, C-terminal) | GO:0016021 (integral component of membrane), GO:0051205 (protein insertion into membrane) |
| Niben101Scf00568g04006.1 | 331 | 180 | -0.709 | 0.028397246 | sp\|Q1ACH1\|CYB6_CHAVU | Cytochrome b6 | IPR016174 (Di-haem cytochrome, transmembrane), IPR027387 (Cytochrome b/b6-like domain) | GO:0009055 (electron carrier activity), GO:0016020 (membrane), GO:0016491 (oxidoreductase activity), GO:0022904 (respiratory electron transport chain) |
| Niben101Scf00309g00001.1 | 159 | 87 | -0.707 | 0.038153886 | gb\|KEH41061.1\| | auxin canalization protein [Medicago truncatula] | IPR008546 (Domain of unknown function DUF828), IPR013666 (Pleckstrin-like, plant) | NA |
| Niben101Scf02156g11001.1 | 90 | 50 | -0.706 | 0.037986509 | AT3G12920.1 | SBP (S-ribonuclease binding protein) family protein LENGTH=335 | IPR013083 (Zinc finger, RING/FYVE/PHD-type) | GO:0005515 (protein binding), GO:0008270 (zinc ion binding) |
| Niben101Scf00715g01003.1 | 300 | 167 | -0.706 | 0.002838954 | NA | Unknown protein | NA | NA |
| Niben101Scf09708g10001.1 | 117 | 65 | -0.701 | 0.045841125 | sp\|Q1C5T7\|HEM6_YERPA | Oxygen-dependent coproporphyrinogen-III oxidase | IPR001260 (Coproporphyrinogen III oxidase, aerobic) | GO:0004109 (coproporphyrinogen oxidase activity), GO:0006779 (porphyrin-containing compound biosynthetic process), GO:0055114 (oxidation-reduction process) |
| Niben101Scf08465g02007.1 | 217 | 120 | -0.696 | 0.036533653 | sp\|Q00423\|HMGYA_SOYBN | HMG-Y-related protein A | IPR011991 (Winged helix-turn-helix DNA-binding domain), IPR020478 (AT hook-like) | GO:0000786 (nucleosome), GO:0003677 (DNA binding), GO:0005634 (nucleus), GO:0006334 (nucleosome assembly) |
| Niben101Scf01718g07005.1 | 136 | 76 | -0.691 | 0.017195061 | AT1G20330.1 | sterol methyltransferase 2 LENGTH=361 | IPR013705 (Sterol methyltransferase C-terminal), IPR029063 (S-adenosyl-L-methionine-dependent methyltransferase) | GO:0006694 (steroid biosynthetic process), GO:0008152 (metabolic process), GO:0008168 (methyltransferase activity) |
| Niben101Scf10608g01005.1 | 349 | 193 | -0.688 | 0.008197815 | sp\|O23264\|SEBP1_ARATH | Selenium-binding protein 1 | IPR008826 (Selenium-binding protein) | GO:0005515 (protein binding), GO:0008430 (selenium binding) |
| Niben101Scf04444g09013.1 | 488 | 273 | -0.685 | 0.003990172 | sp\|Q6NUB2\|CW15A_XENLA | Protein CWC15 homolog A | IPR006973 (Pre-mRNA-splicing factor Cwf15/Cwc15) | GO:0000398 (mRNA splicing, via spliceosome), GO:0005681 (spliceosomal complex) |
| Niben101Scf01001g07022.1 | 110 | 62 | -0.679 | 0.043251595 | sp\|Q9C652\|RBP1_ARATH | RNA-binding protein 1 | IPR012677 (Nucleotide-binding alpha-beta plait domain) | GO:0000166 (nucleotide binding), GO:0003676 (nucleic acid binding) |
| Niben101Scf10688g03004.1 | 92 | 52 | -0.673 | 0.033187546 | sp\|Q9XI36\|MBD10_ARATH | Methyl-CpG-binding domain-containing protein 10 | IPR016177 (DNA-binding domain) | GO:0003677 (DNA binding), GO:0005634 (nucleus) |
| Niben101Scf04869g05006.1 | 105 | 59 | -0.671 | 0.033917134 | ref\|XP_003598002.1\| | Dihydroflavonol-4-reductase [Medicago truncatula] gb\|AES68253.1\| cinnamyl-alcohol dehydrogenase-like protein [Medicago truncatula] | IPR001509 (NAD-dependent epimerase/dehydratase, N-terminal domain), IPR016040 (NAD(P)-binding domain) | GO:0003824 (catalytic activity), GO:0050662 (coenzyme binding) |
| Niben101Scf01002g00008.1 | 162 | 92 | -0.667 | 0.022426436 | ref\|NP_001118630.1\| | high chlorophyll fluorescence 243 [Arabidopsis thaliana] gb\|AEE75618.1\| high chlorophyll fluorescence 243 [Arabidopsis thaliana] | NA | NA |
| Niben101Scf03374g09004.1 | 100 | 57 | -0.666 | 0.03059787 | gb\|KEH21046.1\| | myosin heavy chain-like protein [Medicago truncatula] | IPR019448 (EEIG1/EHBP1 N-terminal domain) | NA |
| Niben101Scf00119g02012.1 | 229 | 131 | -0.664 | 0.006548995 | sp\|O25989\|YIDC_HELPY | Membrane protein insertase YidC | IPR001708 (Membrane insertase OXA1/ALB3/YidC), IPR028055 (Membrane insertase YidC/Oxa1, C-terminal) | GO:0016021 (integral component of membrane), GO:0051205 (protein insertion into membrane) |
| Niben101Scf03538g02003.1 | 103 | 58 | -0.661 | 0.027862352 | sp\|P43672\|UUP_ECOLI | ABC transporter ATP-binding protein uup | IPR003439 (ABC transporter-like), IPR027417 (P-loop containing nucleoside triphosphate hydrolase) | GO:0005524 (ATP binding), GO:0016887 (ATPase activity) |
| Niben101Scf00753g03018.1 | 156 | 88 | -0.661 | 0.041674684 | sp\|Q8RWD0\|COL16_ARATH | Zinc finger protein CONSTANS-LIKE 16 | IPR000315 (Zinc finger, B-box), IPR010402 (CCT domain) | GO:0005515 (protein binding), GO:0005622 (intracellular), GO:0008270 (zinc ion binding) |
| Niben101Scf05487g01009.1 | 151 | 86 | -0.659 | 0.044047111 | sp\|P37218\|H1_SOLLC | Histone H1 | IPR005819 (Histone H5) | GO:0000786 (nucleosome), GO:0003677 (DNA binding), GO:0005634 (nucleus), GO:0006334 (nucleosome assembly) |
| Niben101Scf01761g05001.1 | 83 | 47 | -0.656 | 0.028365031 | sp\|B7ZUF3\|SETD3_XENTR | Histone-lysine N-methyltransferase setd3 | IPR015353 (Rubisco LSMT, substrate-binding domain) | NA |
| Niben101Scf03287g02002.1 | 122 | 69 | -0.654 | 0.032890027 | sp\|O04006\|PSAH_BRACM | Photosystem I reaction center subunit VI, chloroplastic | IPR004928 (Photosystem I PsaH, reaction centre subunit VI), IPR023833 (Signal peptide, camelysin), IPR029027 (Single alpha-helix domain) | GO:0009522 (photosystem I), GO:0009538 (photosystem I reaction center), GO:0015979 (photosynthesis) |
| Niben101Scf01276g00005.1 | 263 | 150 | -0.648 | 0.012689829 | sp\|P12360\|CB11_SOLLC | Chlorophyll a-b binding protein 6A, chloroplastic | IPR022796 (Chlorophyll A-B binding protein), IPR023329 (Chlorophyll a/b binding protein domain) | GO:0009765 (photosynthesis, light harvesting), GO:0016020 (membrane) |
| Niben101Scf06999g01002.1 | 119 | 69 | -0.634 | 0.028492525 | sp\|O04375\|2A5A_ARATH | Serine/threonine protein phosphatase 2A 57 kDa regulatory subunit B' alpha isoform | IPR002554 (Protein phosphatase 2A, regulatory B subunit, B56), IPR016024 (Armadillo-type fold) | GO:0000159 (protein phosphatase type 2A complex), GO:0005488 (binding), GO:0007165 (signal transduction), GO:0008601 (protein phosphatase type 2A regulator activity) |
| Niben101Scf02594g01007.1 | 131 | 77 | -0.63 | 0.042100909 | sp\|Q9FRL3\|ERDL6_ARATH | Sugar transporter ERD6-like 6 | IPR005828 (General substrate transporter), IPR020846 (Major facilitator superfamily domain) | GO:0005215 (transporter activity), GO:0016021 (integral component of membrane), GO:0022857 (transmembrane transporter activity), GO:0055085 (transmembrane transport) |
| Niben101Scf00332g04003.1 | 210 | 122 | -0.627 | 0.01066517 | emb\|CDX76648.1\| | BnaC08g32340D [Brassica napus] | IPR012442 (Protein of unknown function DUF1645, plant) | NA |
| Niben101Scf09447g01014.1 | 429 | 253 | -0.622 | 0.013086019 | sp\|Q9DC07\|LNEBL_MOUSE | LIM zinc-binding domain-containing Nebulette | IPR001781 (Zinc finger, LIM-type) | GO:0008270 (zinc ion binding) |
| Niben101Scf00160g10022.1 | 155 | 90 | -0.621 | 0.036644617 | AT3G23550.1 | MATE efflux family protein LENGTH=469 | IPR002528 (Multi antimicrobial extrusion protein) | GO:0006855 (drug transmembrane transport), GO:0015238 (drug transmembrane transporter activity), GO:0015297 (antiporter activity), GO:0016020 (membrane), GO:0055085 (transmembrane transport) |
| Niben101Scf01899g02003.1 | 678 | 402 | -0.619 | 0.037425584 | AT1G30360.1 | Early-responsive to dehydration stress protein (ERD4) LENGTH=724 | IPR003864 (Domain of unknown function DUF221), IPR027815 (Domain of unknown function DUF4463) | GO:0016020 (membrane) |
| Niben101Scf05030g02010.1 | 273 | 162 | -0.617 | 0.03454746 | AT5G62360.1 | Plant invertase/pectin methylesterase inhibitor superfamily protein LENGTH=203 | IPR006501 (Pectinesterase inhibitor domain) | GO:0004857 (enzyme inhibitor activity), GO:0030599 (pectinesterase activity) |
| Niben101Scf01036g03001.1 | 9576 | 5601 | -0.617 | 0.000149652 | sp\|P15904\|POR_AVESA | Protochlorophyllide reductase | IPR002347 (Glucose/ribitol dehydrogenase) | GO:0008152 (metabolic process), GO:0016491 (oxidoreductase activity), GO:0016630 (protochlorophyllide reductase activity), GO:0055114 (oxidation-reduction process) |
| Niben101Scf03142g02005.1 | 267 | 158 | -0.615 | 0.032765165 | AT4G29060.1 | elongation factor Ts family protein LENGTH=953 | IPR001816 (Translation elongation factor EFTs/EF1B), IPR012340 (Nucleic acid-binding, OB-fold) | GO:0003676 (nucleic acid binding), GO:0003746 (translation elongation factor activity), GO:0005515 (protein binding), GO:0005622 (intracellular), GO:0006414 (translational elongation) |
| Niben101Scf10048g02013.1 | 168 | 98 | -0.612 | 0.030433314 | sp\|Q85A24\|CYB6_ANTFO | Cytochrome b6 | IPR016174 (Di-haem cytochrome, transmembrane), IPR023530 (Cytochrome b6, PetB), IPR027387 (Cytochrome b/b6-like domain) | GO:0005506 (iron ion binding), GO:0009055 (electron carrier activity), GO:0016020 (membrane), GO:0016021 (integral component of membrane), GO:0016491 (oxidoreductase activity), GO:0020037 (heme binding), GO:0022900 (electron transport chain), GO:0022904 (respiratory electron transport chain), GO:0045158 (electron transporter, transferring electrons within cytochrome b6/f complex of photosystem II activity) |
| Niben101Scf05216g09026.1 | 249 | 148 | -0.61 | 0.006857673 | AT1G55480.1 | protein containing PDZ domain, a K-box domain, and a TPR region LENGTH=335 | IPR001478 (PDZ domain), IPR011990 (Tetratricopeptide-like helical domain) | GO:0005515 (protein binding) |
| Niben101Scf01145g02013.1 | 335 | 198 | -0.608 | 0.042780377 | ref\|XP_002532850.1\| | nucleic acid binding protein, putative [Ricinus communis] gb\|EEF29528.1\| nucleic acid binding protein, putative [Ricinus communis] | IPR001878 (Zinc finger, CCHC-type) | GO:0003676 (nucleic acid binding), GO:0008270 (zinc ion binding) |
| Niben101Scf05519g01004.1 | 496 | 295 | -0.604 | 0.016185093 | sp\|Q76LC6\|RBM24_DANRE | RNA-binding protein 24 | IPR012677 (Nucleotide-binding alpha-beta plait domain) | GO:0000166 (nucleotide binding), GO:0003676 (nucleic acid binding) |
| Niben101Scf01409g06005.1 | 166 | 99 | -0.596 | 0.044331393 | sp\|Q9FHH8\|COL5_ARATH | Zinc finger protein CONSTANS-LIKE 5 | IPR000315 (Zinc finger, B-box), IPR010402 (CCT domain) | GO:0005515 (protein binding), GO:0005622 (intracellular), GO:0008270 (zinc ion binding) |
| Niben101Scf02406g04044.1 | 550 | 330 | -0.595 | 0.016528874 | gb\|AER28834.1\| | COSII_At2g15890-like protein [Solanum volubile] | NA | NA |
| Niben101Scf01472g02001.1 | 205 | 122 | -0.591 | 0.020204016 | sp\|Q57242\|UUP1_HAEIN | ABC transporter ATP-binding protein uup-1 | IPR003439 (ABC transporter-like), IPR027417 (P-loop containing nucleoside triphosphate hydrolase) | GO:0005524 (ATP binding), GO:0016887 (ATPase activity) |
| Niben101Scf06906g01008.1 | 177 | 105 | -0.588 | 0.026776581 | emb\|CDX89516.1\| | BnaC04g35350D [Brassica napus] | NA | NA |
| Niben101Scf05343g01007.1 | 271 | 162 | -0.588 | 0.020470188 | AT5G53550.1 | YELLOW STRIPE like 3 LENGTH=675 | IPR004813 (Oligopeptide transporter, OPT superfamily) | GO:0055085 (transmembrane transport) |
| Niben101Scf07000g01005.1 | 223 | 133 | -0.584 | 0.046596961 | sp\|P82658\|TL19_ARATH | Thylakoid lumenal 19 kDa protein, chloroplastic | IPR002683 (Photosystem II PsbP, oxygen evolving complex) | GO:0005509 (calcium ion binding), GO:0009523 (photosystem II), GO:0009654 (photosystem II oxygen evolving complex), GO:0015979 (photosynthesis), GO:0019898 (extrinsic component of membrane) |
| Niben101Scf04529g00012.1 | 484 | 291 | -0.583 | 0.032898853 | sp\|P48188\|PSBK_MAIZE | Photosystem II reaction center protein K | IPR003686 (Photosystem II PsbI), IPR003687 (Photosystem II PsbK) | GO:0009523 (photosystem II), GO:0009539 (photosystem II reaction center), GO:0015979 (photosynthesis), GO:0016020 (membrane) |
| Niben101Scf02807g11007.1 | 124 | 75 | -0.582 | 0.0458623 | AT3G22200.2 | Pyridoxal phosphate (PLP)-dependent transferases superfamily protein LENGTH=513 | IPR005814 (Aminotransferase class-III), IPR015424 (Pyridoxal phosphate-dependent transferase) | GO:0003824 (catalytic activity), GO:0008483 (transaminase activity), GO:0030170 (pyridoxal phosphate binding) |
| Niben101Scf01964g02018.1 | 164 | 99 | -0.581 | 0.036637245 | NA | Unknown protein | NA | NA |
| Niben101Scf15666g00009.1 | 762 | 463 | -0.579 | 0.025054234 | sp\|P74518\|LRTA_SYNY3 | Light-repressed protein A homolog | IPR003489 (Ribosomal protein S30Ae/sigma 54 modulation protein) | GO:0044238 (primary metabolic process) |
| Niben101Scf00887g04013.1 | 174 | 106 | -0.575 | 0.036533653 | sp\|Q99614\|TTC1_HUMAN | Tetratricopeptide repeat protein 1 | IPR011990 (Tetratricopeptide-like helical domain), IPR012336 (Thioredoxin-like fold) | GO:0005515 (protein binding) |
| Niben101Scf00747g00005.1 | 139 | 84 | -0.572 | 0.043321513 | emb\|CDY11893.1\| | BnaC03g13190D [Brassica napus] | NA | NA |
| Niben101Scf15187g03007.1 | 1133 | 686 | -0.568 | 0.04468457 | sp\|P18212\|PSBP2_TOBAC | Oxygen-evolving enhancer protein 2-2, chloroplastic | IPR002683 (Photosystem II PsbP, oxygen evolving complex) | GO:0005509 (calcium ion binding), GO:0009523 (photosystem II), GO:0009654 (photosystem II oxygen evolving complex), GO:0015979 (photosynthesis), GO:0019898 (extrinsic component of membrane) |
| Niben101Scf00897g03002.1 | 501 | 305 | -0.566 | 0.020716644 | sp\|P46417\|GSTX3_SOYBN | Glutathione S-transferase 3 | IPR010987 (Glutathione S-transferase, C-terminal-like), IPR012336 (Thioredoxin-like fold) | GO:0005515 (protein binding) |
| Niben101Scf02237g04009.1 | 183 | 112 | -0.561 | 0.045776188 | AT5G27330.1 | Prefoldin chaperone subunit family protein LENGTH=628 | NA | NA |
| Niben101Scf01550g01003.1 | 153 | 94 | -0.559 | 0.044005531 | sp\|Q94A65\|STR14_ARATH | Rhodanese-like domain-containing protein 14, chloroplastic | IPR001763 (Rhodanese-like domain) | NA |
| Niben101Scf07355g01005.1 | 198 | 121 | -0.556 | 0.028391186 | emb\|CDX94272.1\| | BnaC02g29320D [Brassica napus] | NA | NA |
| Niben101Scf01694g12011.1 | 273 | 168 | -0.556 | 0.041496045 | AT2G29670.1 | Tetratricopeptide repeat (TPR)-like superfamily protein LENGTH=536 | IPR011990 (Tetratricopeptide-like helical domain) | GO:0005515 (protein binding) |
| Niben101Scf02160g06059.1 | 151 | 92 | -0.554 | 0.038628626 | sp\|P08742\|COX1_MAIZE | Cytochrome c oxidase subunit 1 | IPR000883 (Cytochrome c oxidase subunit I), IPR023615 (Cytochrome c oxidase, subunit I, copper-binding site), IPR023616 (Cytochrome c oxidase, subunit I domain) | GO:0004129 (cytochrome-c oxidase activity), GO:0005506 (iron ion binding), GO:0009055 (electron carrier activity), GO:0009060 (aerobic respiration), GO:0016021 (integral component of membrane), GO:0020037 (heme binding), GO:0055114 (oxidation-reduction process) |
| Niben101Scf01184g14003.1 | 142 | 87 | -0.547 | 0.042607504 | emb\|CDX94272.1\| | BnaC02g29320D [Brassica napus] | NA | NA |
| Niben101Scf00860g03018.1 | 657 | 406 | -0.546 | 0.027040915 | AT5G51100.1 | Fe superoxide dismutase 2 LENGTH=305 | IPR001189 (Manganese/iron superoxide dismutase) | GO:0004784 (superoxide dismutase activity), GO:0006801 (superoxide metabolic process), GO:0046872 (metal ion binding), GO:0055114 (oxidation-reduction process) |
| Niben101Scf04528g09004.1 | 286 | 179 | -0.523 | 0.048205217 | AT2G28260.1 | cyclic nucleotide-gated channel 15 LENGTH=678 | IPR014710 (RmlC-like jelly roll fold) | NA |
| Niben101Scf18637g02010.1 | 324 | 205 | -0.509 | 0.032081427 | AT5G65630.1 | global transcription factor group E7 LENGTH=590 | IPR027353 (NET domain) | NA |
| Niben101Scf03251g00008.1 | 485 | 306 | -0.505 | 0.028591545 | sp\|Q40831\|TBA1_PELFA | Tubulin alpha-1 chain | IPR000217 (Tubulin), IPR023123 (Tubulin, C-terminal) | GO:0003924 (GTPase activity), GO:0005200 (structural constituent of cytoskeleton), GO:0005525 (GTP binding), GO:0005874 (microtubule), GO:0006184 (GTP catabolic process), GO:0007017 (microtubule-based process), GO:0043234 (protein complex), GO:0051258 (protein polymerization) |

b. **Downregulated**

| **ID** | **Cas9 av. reads** | ***virp1* av. reads** | **log2FC** | **padj** | **Blast-Hit-Accession** | **Human-Readable-Description** | **Interpro-ID (Description)** | **Gene-Ontology-ID (Name)** |
| --- | --- | --- | --- | --- | --- | --- | --- | --- |
| Niben101Scf00683g03048.1 | 35 | 2 | -3.937 | 0.0044444 | sp\|Q9MAU3\|TAF6_ARATH | Transcription initiation factor TFIID subunit 6 | IPR009072 (Histone-fold), IPR011442 (Domain of unknown function DUF1546) | GO:0005634 (nucleus), GO:0006352 (DNA-templated transcription, initiation), GO:0046982 (protein heterodimerization activity), GO:0051090 (regulation of sequence-specific DNA binding transcription factor activity) |
| Niben101Scf01008g01008.1 | 356 | 27 | -3.704 | 0.001383597 | AT2G33830.2 | Dormancy/auxin associated family protein LENGTH=108 | IPR008406 (Dormancyauxin associated) |  |
| Niben101Scf04327g02021.1 | 20 | 2 | -3.433 | 0.000872599 | gb\|ACM43526.1\| | phosphatidylinositol-specific phospholipase c [Listeria monocytogenes] | IPR017946 (PLC-like phosphodiesterase, TIM beta/alpha-barrel domain) | GO:0006629 (lipid metabolic process), GO:0008081 (phosphoric diester hydrolase activity) |
| Niben101Scf01084g06002.1 | 75 | 7 | -3.43 | 0.000359425 | sp\|P23137\|GRP1_TOBAC | Glycine-rich protein |  |  |
| Niben101Scf03068g00011.1 | 446 | 47 | -3.255 | 0.021792885 | sp\|Q3C1J5\|ATPB_NICSY | ATP synthase subunit beta, chloroplastic | IPR000793 (ATPase, F1/V1/A1 complex, alpha/beta subunit, C-terminal), IPR001469 (ATPase, F1 complex, delta/epsilon subunit), IPR020003 (ATPase, alpha/beta subunit, nucleotide-binding domain, active site) | GO:0005524 (ATP binding), GO:0015986 (ATP synthesis coupled proton transport), GO:0015991 (ATP hydrolysis coupled proton transport), GO:0016820 (hydrolase activity, acting on acid anhydrides, catalyzing transmembrane movement of substances), GO:0033178 (proton-transporting two-sector ATPase complex, catalytic domain), GO:0045261 (proton-transporting ATP synthase complex, catalytic core F(1)), GO:0046933 (proton-transporting ATP synthase activity, rotational mechanism), GO:0046961 (proton-transporting ATPase activity, rotational mechanism) |
| Niben101Scf03975g01004.1 | 20 | 2 | -3.205 | 0.006303153 | emb\|CDY10237.1\| | BnaC05g04800D [Brassica napus] | IPR025322 (Protein of unknown function DUF4228, plant) |  |
| Niben101Scf03098g00008.1 | 456 | 57 | -2.996 | 0.008100188 | sp\|Q14F96\|RK23_POPAL | 50S ribosomal protein L23, chloroplastic | IPR013025 (Ribosomal protein L25/L23) | GO:0000166 (nucleotide binding), GO:0003735 (structural constituent of ribosome), GO:0005622 (intracellular), GO:0005840 (ribosome), GO:0006412 (translation) |
| Niben101Scf04209g01002.1 | 2206 | 312 | -2.821 | 0.000216128 | gb\|ACY74744.2\| | methanol inducible protein [Nicotiana benthamiana] | |  |
| Niben101Scf00621g05002.1 | 34 | 5 | -2.685 | 0.014360726 | sp\|Q75KR1\|PPDK2_ORYSJ | Pyruvate, phosphate dikinase 2 | IPR010121 (Pyruvate, phosphate dikinase), IPR015813 (Pyruvate/Phosphoenolpyruvate kinase-like domain) | GO:0003824 (catalytic activity), GO:0005524 (ATP binding), GO:0006090 (pyruvate metabolic process), GO:0016301 (kinase activity), GO:0016310 (phosphorylation), GO:0016772 (transferase activity, transferring phosphorus-containing groups), GO:0050242 (pyruvate, phosphate dikinase activity) |
| Niben101Scf02078g05004.1 | 883 | 152 | -2.537 | 0.023614628 | sp\|A1EA06\|RR14_AGRST | 30S ribosomal protein S14, chloroplastic | IPR001209 (Ribosomal protein S14) | GO:0003735 (structural constituent of ribosome), GO:0005622 (intracellular), GO:0005840 (ribosome), GO:0006412 (translation) |
| Niben101Scf02290g07003.1 | 57 | 10 | -2.487 | 0.012644511 | sp\|O35685\|NUDC_MOUSE | Nuclear migration protein nudC | IPR008978 (HSP20-like chaperone) |  |
| Niben101Scf09596g00002.1 | 52 | 9 | -2.477 | 0.012961021 | AT2G37460.1 | nodulin MtN21 /EamA-like transporter family protein LENGTH=380 | IPR000620 (EamA domain), IPR030184 (WAT1-related protein) | GO:0016020 (membrane), GO:0016021 (integral component of membrane), GO:0022857 (transmembrane transporter activity) |
| Niben101Scf04133g02029.1 | 30 | 5 | -2.468 | 0.001938212 | sp\|Q9FLJ2\|NC100_ARATH | NAC domain-containing protein 100 | IPR003441 (NAC domain) | GO:0003677 (DNA binding), GO:0006355 (regulation of transcription, DNA-templated) |
| Niben101Scf00383g04011.1 | 25 | 5 | -2.446 | 0.008754985 | AT1G10210.1 | mitogen-activated protein kinase 1 LENGTH=370 | IPR011009 (Protein kinase-like domain) | GO:0004672 (protein kinase activity), GO:0004674 (protein serine/threonine kinase activity), GO:0004707 (MAP kinase activity), GO:0005524 (ATP binding), GO:0006468 (protein phosphorylation), GO:0016772 (transferase activity, transferring phosphorus-containing groups) |
| Niben101Scf03932g05003.1 | 16 | 3 | -2.332 | 0.025895625 | AT3G15030.1 | TCP family transcription factor 4 LENGTH=420 | IPR005333 (Transcription factor, TCP) |  |
| Niben101Scf01049g05011.1 | 14 | 3 | -2.28 | 0.037139073 | sp\|Q9SXA1\|P4KA1_ARATH | Phosphatidylinositol 4-kinase alpha 1 | IPR015433 (Phosphatidylinositol Kinase), IPR016024 (Armadillo-type fold) | GO:0005488 (binding), GO:0046854 (phosphatidylinositol phosphorylation), GO:0048015 (phosphatidylinositol-mediated signaling) |
| Niben101Scf02280g03001.1 | 32 | 7 | -2.249 | 0.009912347 | AT2G40095.1 | Alpha/beta hydrolase related protein LENGTH=209 | IPR019498 (MENTAL domain) |  |
| Niben101Scf11126g02001.1 | 75 | 16 | -2.215 | 0.002585635 | sp\|Q9SXZ2\|FT_ARATH | Protein FLOWERING LOCUS T | IPR008914 (Phosphatidylethanolamine-binding protein PEBP) | |
| Niben101Scf05534g01008.1 | 38 | 8 | -2.211 | 0.002342568 | AT1G66220.1 | Subtilase family protein LENGTH=753 | IPR015500 (Peptidase S8, subtilisin-related) |  |
| Niben101Scf18822g02016.1 | 25 | 5 | -2.165 | 0.00316693 | sp\|Q84KG5\|CCD_CROSA | Carotenoid 9,10(9',10')-cleavage dioxygenase | IPR004294 (Carotenoid oxygenase) |  |
| Niben101Scf02594g07008.1 | 54 | 12 | -2.148 | 0.010476828 | sp\|Q93VH2\|BBD2_ARATH | Bifunctional nuclease 2 | IPR003729 (Bifunctional nuclease domain) | GO:0004518 (nuclease activity) |
| Niben101Scf02793g27005.1 | 59 | 13 | -2.147 | 0.04336653 | sp\|Q94KU5\|PAP3_BRACM | Plastid lipid-associated protein 3, chloroplastic | |  |
| Niben101Scf03985g04023.1 | 111 | 25 | -2.125 | 4.95487E-06 | ref\|XP_001288151.1\| | ankyrin repeat protein [Trichomonas vaginalis G3] gb\|EAX75221.1\| ankyrin repeat protein, putative [Trichomonas vaginalis G3] | | |
| Niben101Scf03732g02011.1 | 16 | 3 | -2.124 | 0.042010747 | ref\|XP_002531664.1\| | conserved hypothetical protein [Ricinus communis] gb\|EEF30708.1\| conserved hypothetical protein [Ricinus communis] | | |
| Niben101Scf06248g00007.1 | 871 | 200 | -2.124 | 0.034017117 | dbj\|BAA21495.1\| | metallocarboxypeptidase inhibitor [Solanum tuberosum] | |  |
| Niben101Scf03923g03003.1 | 19 | 4 | -2.123 | 0.035805931 | sp\|Q9LUH8\|HFA6B_ARATH | Heat stress transcription factor A-6b | IPR011991 (Winged helix-turn-helix DNA-binding domain), IPR027725 (Heat shock transcription factor family) | GO:0003700 (sequence-specific DNA binding transcription factor activity), GO:0005634 (nucleus), GO:0006355 (regulation of transcription, DNA-templated), GO:0009408 (response to heat), GO:0043565 (sequence-specific DNA binding) |
| Niben101Scf01166g17002.1 | 18 | 4 | -2.103 | 0.024182548 | sp\|Q82BY4\|DNAJ2_STRAW | Chaperone protein DnaJ 2 | IPR001623 (DnaJ domain), IPR022226 (Protein of unknown function DUF3752) | |
| Niben101Scf04133g01029.1 | 24 | 6 | -2.066 | 0.006449087 | sp\|Q94A21\|SPX4_ARATH | SPX domain-containing protein 4 | IPR004331 (SPX, N-terminal) |  |
| Niben101Scf18639g00021.1 | 14 | 3 | -2.047 | 0.032130771 | sp\|Q9LDF2\|LAG11_ARATH | LAG1 longevity assurance homolog 1 | IPR006634 (TRAM/LAG1/CLN8 homology domain) | GO:0016021 (integral component of membrane) |
| Niben101Scf09973g00011.1 | 52 | 13 | -2.034 | 0.000392631 | sp\|O82012\|HSP12_SOLPE | 17.6 kDa class I heat shock protein | IPR008978 (HSP20-like chaperone) |  |
| Niben101Scf08130g00015.1 | 27 | 7 | -2.02 | 0.048172688 | sp\|Q9SR36\|GSTU8_ARATH | Glutathione S-transferase U8 | IPR010987 (Glutathione S-transferase, C-terminal-like), IPR012336 (Thioredoxin-like fold) | GO:0005515 (protein binding) |
| Niben101Scf07788g00007.1 | 129 | 32 | -1.997 | 0.002483337 | ref\|XP_002521463.1\| | Microtubule-associated protein futsch, putative [Ricinus communis] gb\|EEF40953.1\| Microtubule-associated protein futsch, putative [Ricinus communis] | | |
| Niben101Scf02069g00018.1 | 647 | 165 | -1.974 | 0.02441887 | gb\|ACY74744.2\| | methanol inducible protein [Nicotiana benthamiana] | |  |
| Niben101Scf05329g01008.1 | 30 | 8 | -1.959 | 0.026041153 | AT3G56200.1 | Transmembrane amino acid transporter family protein LENGTH=435 | IPR013057 (Amino acid transporter, transmembrane) |  |
| Niben101Scf01580g02008.1 | 39 | 10 | -1.934 | 0.012655353 | sp\|Q5CZL1\|GDAP2_XENTR | Ganglioside-induced differentiation-associated protein 2 | IPR001251 (CRAL-TRIO domain) |  |
| Niben101Scf04499g00008.1 | 20 | 5 | -1.932 | 0.022950795 | AT3G12920.1 | SBP (S-ribonuclease binding protein) family protein LENGTH=335 | IPR013083 (Zinc finger, RING/FYVE/PHD-type) | GO:0005515 (protein binding), GO:0008270 (zinc ion binding) |
| Niben101Scf01734g01046.1 | 234 | 62 | -1.914 | 0.014132391 | gb\|AFH58131.1\| | Ycf6, partial (chloroplast) [Dombeya acutangula] | |  |
| Niben101Scf01731g05006.1 | 37 | 10 | -1.911 | 0.017932526 | sp\|P93332\|NOD3_MEDTR | Bidirectional sugar transporter N3 |  |  |
| Niben101Scf00916g01008.1 | 174 | 47 | -1.904 | 0.001245721 | AT5G58070.1 | temperature-induced lipocalin LENGTH=186 | IPR002446 (Lipocalin, bacterial), IPR011038 (Calycin-like) | GO:0005215 (transporter activity) |
| Niben101Scf07987g01001.1 | 37 | 10 | -1.898 | 0.010057412 | sp\|Q84MB3\|ACCH1_ARATH | 1-aminocyclopropane-1-carboxylate oxidase homolog 1 | IPR005123 (Oxoglutarate/iron-dependent dioxygenase), IPR026992 (Non-haem dioxygenase N-terminal domain), IPR027443 (Isopenicillin N synthase-like) | GO:0016491 (oxidoreductase activity), GO:0016706 (oxidoreductase activity, acting on paired donors, with incorporation or reduction of molecular oxygen, 2-oxoglutarate as one donor, and incorporation of one atom each of oxygen into both donors), GO:0055114 (oxidation-reduction process) |
| Niben101Scf04113g07011.1 | 179 | 48 | -1.89 | 0.0079891 | ref\|NP_001051767.1\| | product [Oryza sativa Japonica Group] |  |  |
| Niben101Scf14009g02003.1 | 322 | 89 | -1.842 | 4.50432E-08 | ref\|XP_002521463.1\| | Microtubule-associated protein futsch, putative [Ricinus communis] gb\|EEF40953.1\| Microtubule-associated protein futsch, putative [Ricinus communis] | | |
| Niben101Scf09321g01008.1 | 68 | 19 | -1.835 | 0.034017117 | sp\|Q67YC0\|PPSP1_ARATH | Inorganic pyrophosphatase 1 | IPR006383 (HAD-superfamily hydrolase, subfamily IB, PSPase-like), IPR023214 (HAD-like domain) | GO:0008152 (metabolic process), GO:0016791 (phosphatase activity) |
| Niben101Scf02819g00005.1 | 119 | 34 | -1.83 | 0.001505348 | sp\|Q9SK27\|ENL1_ARATH | Early nodulin-like protein 1 | IPR008972 (Cupredoxin), IPR028871 (Blue (type 1) copper protein, binding site) | GO:0009055 (electron carrier activity) |
| Niben101Scf01863g04002.1 | 35 | 10 | -1.797 | 0.039649079 | sp\|Q9M2Y0\|PSK3_ARATH | Phytosulfokines 3 | IPR009438 (Phytosulfokine) | GO:0005576 (extracellular region), GO:0008083 (growth factor activity), GO:0008283 (cell proliferation) |
| Niben101Scf06436g00006.1 | 44 | 13 | -1.785 | 0.042010747 | AT5G07990.1 | Cytochrome P450 superfamily protein LENGTH=513 | IPR001128 (Cytochrome P450) | GO:0005506 (iron ion binding), GO:0016705 (oxidoreductase activity, acting on paired donors, with incorporation or reduction of molecular oxygen), GO:0020037 (heme binding), GO:0055114 (oxidation-reduction process) |
| Niben101Scf08106g00006.1 | 18 | 5 | -1.757 | 0.043488602 | sp\|Q9FZ70\|FDL1_ARATH | F-box/FBD/LRR-repeat protein | IPR001810 (F-box domain) | GO:0005515 (protein binding) |
| Niben101Scf11071g02017.1 | 29 | 8 | -1.737 | 0.026603735 | sp\|Q9LIG2\|RLK6_ARATH | Receptor-like protein kinase | IPR011009 (Protein kinase-like domain), IPR013320 (Concanavalin A-like lectin/glucanase domain) | GO:0004672 (protein kinase activity), GO:0004674 (protein serine/threonine kinase activity), GO:0005524 (ATP binding), GO:0006468 (protein phosphorylation), GO:0016772 (transferase activity, transferring phosphorus-containing groups) |
| Niben101Scf07599g00012.1 | 18 | 5 | -1.7 | 0.043850658 | sp\|P57057\|GLPT_HUMAN | Glycerol-3-phosphate transporter | IPR011701 (Major facilitator superfamily), IPR020846 (Major facilitator superfamily domain) | GO:0005215 (transporter activity), GO:0006810 (transport), GO:0016021 (integral component of membrane), GO:0055085 (transmembrane transport) |
| Niben101Scf20652g00003.1 | 28 | 9 | -1.696 | 0.020750481 | sp\|F4HQX1\|JAL3_ARATH | Jacalin-related lectin 3 | IPR001229 (Jacalin-like lectin domain) |  |
| Niben101Scf00349g02005.1 | 298 | 92 | -1.693 | 0.042010747 | sp\|O95372\|LYPA2_HUMAN | Acyl-protein thioesterase 2 | IPR029058 (Alpha/Beta hydrolase fold) | GO:0016787 (hydrolase activity) |
| Niben101Scf19421g00012.1 | 35 | 11 | -1.686 | 0.046484191 | AT1G69830.1 | alpha-amylase-like 3 LENGTH=887 | IPR015902 (Glycoside hydrolase, family 13) | GO:0003824 (catalytic activity), GO:0004556 (alpha-amylase activity), GO:0005509 (calcium ion binding), GO:0005975 (carbohydrate metabolic process), GO:0043169 (cation binding) |
| Niben101Scf07590g01014.1 | 161 | 50 | -1.682 | 0.033396411 |  | Unknown protein |  |  |
| Niben101Scf05543g01003.1 | 95 | 30 | -1.655 | 0.012655353 | AT1G69780.1 | Homeobox-leucine zipper protein family LENGTH=294 | IPR003106 (Leucine zipper, homeobox-associated), IPR009057 (Homeodomain-like) | GO:0003677 (DNA binding), GO:0003700 (sequence-specific DNA binding transcription factor activity), GO:0005634 (nucleus), GO:0006355 (regulation of transcription, DNA-templated), GO:0043565 (sequence-specific DNA binding) |
| Niben101Scf10169g04009.1 | 63 | 20 | -1.652 | 0.000685859 | AT1G07160.1 | Protein phosphatase 2C family protein LENGTH=380 | IPR001932 (Protein phosphatase 2C (PP2C)-like domain), IPR015655 (Protein phosphatase 2C) | GO:0003824 (catalytic activity), GO:0004722 (protein serine/threonine phosphatase activity), GO:0006470 (protein dephosphorylation) |
| Niben101Scf01719g08001.1 | 41 | 13 | -1.65 | 0.011901186 | sp\|Q7PC83\|AB41G_ARATH | ABC transporter G family member 41 | IPR003439 (ABC transporter-like), IPR013525 (ABC-2 type transporter), IPR013581 (Plant PDR ABC transporter associated), IPR027417 (P-loop containing nucleoside triphosphate hydrolase), IPR029481 (ABC-transporter extracellular N-terminal domain) | GO:0005524 (ATP binding), GO:0016020 (membrane), GO:0016887 (ATPase activity) |
| Niben101Scf03842g09017.1 | 41 | 13 | -1.641 | 0.009604444 | sp\|P92985\|RBP1C_ARATH | Ran-binding protein 1 homolog c | IPR011993 (Pleckstrin homology-like domain) | GO:0046907 (intracellular transport) |
| Niben101Scf08422g00003.1 | 4572 | 1474 | -1.633 | 0.005521117 | AT2G33830.2 | Dormancy/auxin associated family protein LENGTH=108 | IPR008406 (Dormancyauxin associated) |  |
| Niben101Scf00159g03005.1 | 89 | 28 | -1.63 | 0.030113958 | sp\|Q9FLP6\|SUMO2_ARATH | Small ubiquitin-related modifier 2 | IPR029071 (Ubiquitin-related domain) | GO:0005515 (protein binding) |
| Niben101Scf05142g00023.1 | 248 | 81 | -1.616 | 0.023766788 | sp\|P82129\|RR7_SPIOL | 30S ribosomal protein S7, chloroplastic | IPR000235 (Ribosomal protein S5/S7), IPR023798 (Ribosomal protein S7 domain) | GO:0003723 (RNA binding), GO:0003735 (structural constituent of ribosome), GO:0006412 (translation) |
| Niben101Scf02336g00010.1 | 42 | 14 | -1.598 | 0.011115033 | sp\|P25792\|CYSP_SCHMA | Cathepsin B-like cysteine proteinase | IPR013128 (Peptidase C1A), IPR025660 (Cysteine peptidase, histidine active site), IPR025661 (Cysteine peptidase, asparagine active site) | GO:0006508 (proteolysis), GO:0008234 (cysteine-type peptidase activity) |
| Niben101Scf03488g03004.1 | 123 | 41 | -1.598 | 0.022515422 | AT3G23820.1 | UDP-D-glucuronate 4-epimerase 6 LENGTH=460 | IPR001509 (NAD-dependent epimerase/dehydratase, N-terminal domain), IPR008089 (Nucleotide sugar epimerase) | GO:0003824 (catalytic activity), GO:0005975 (carbohydrate metabolic process), GO:0016857 (racemase and epimerase activity, acting on carbohydrates and derivatives), GO:0050662 (coenzyme binding) |
| Niben101Scf01495g01001.1 | 51 | 18 | -1.524 | 0.008154138 | sp\|P81391\|MYB05_ANTMA | Myb-related protein 305 | IPR009057 (Homeodomain-like) | GO:0003677 (DNA binding), GO:0003682 (chromatin binding) |
| Niben101Scf05135g05010.1 | 44 | 15 | -1.522 | 0.023614628 | sp\|P52478\|UBC1_CAEEL | Ubiquitin-conjugating enzyme E2 1 | IPR016135 (Ubiquitin-conjugating enzyme/RWD-like), IPR023313 (Ubiquitin-conjugating enzyme, active site), IPR027230 (SUMO-conjugating enzyme Ubc9) | GO:0019789 (SUMO transferase activity) |
| Niben101Scf00897g03002.1 | 446 | 155 | -1.521 | 1.07105E-05 | sp\|P46417\|GSTX3_SOYBN | Glutathione S-transferase 3 | IPR010987 (Glutathione S-transferase, C-terminal-like), IPR012336 (Thioredoxin-like fold) | GO:0005515 (protein binding) |
| Niben101Scf02982g03019.1 | 46 | 16 | -1.507 | 0.032525849 | AT3G23820.1 | UDP-D-glucuronate 4-epimerase 6 LENGTH=460 | IPR001509 (NAD-dependent epimerase/dehydratase, N-terminal domain), IPR008089 (Nucleotide sugar epimerase) | GO:0003824 (catalytic activity), GO:0005975 (carbohydrate metabolic process), GO:0016857 (racemase and epimerase activity, acting on carbohydrates and derivatives), GO:0050662 (coenzyme binding) |
| Niben101Scf26023g00004.1 | 584 | 207 | -1.494 | 0.005688971 | sp\|O24606\|EIN3_ARATH | Protein ETHYLENE INSENSITIVE 3 |  |  |
| Niben101Scf11283g00002.1 | 1441 | 514 | -1.488 | 0.016720726 | sp\|Q9SSK9\|MLP28_ARATH | MLP-like protein 28 | IPR000916 (Bet v I domain), IPR023393 (START-like domain) | GO:0006952 (defense response), GO:0009607 (response to biotic stimulus) |
| Niben101Scf12277g02002.1 | 37 | 13 | -1.487 | 0.021041225 | AT4G08240.2 | unknown protein; Has 33 Blast hits to 33 proteins in 12 species: Archae - 0; Bacteria - 0; Metazoa - 0; Fungi - 0; Plants - 33; Viruses - 0; Other Eukaryotes - 0 (source: NCBI BLink). LENGTH=136 | | |
| Niben101Scf24482g00010.1 | 129 | 46 | -1.471 | 0.018855636 | AT5G35220.1 | Peptidase M50 family protein LENGTH=548 |  |  |
| Niben101Scf09678g00006.1 | 54 | 19 | -1.465 | 0.02274829 | sp\|Q76LC6\|RBM24_DANRE | RNA-binding protein 24 | IPR012677 (Nucleotide-binding alpha-beta plait domain) | GO:0000166 (nucleotide binding), GO:0003676 (nucleic acid binding) |
| Niben101Scf02665g07008.1 | 28 | 10 | -1.464 | 0.042438918 | sp\|Q8L7Y9\|NPC1_ARATH | Non-specific phospholipase C1 | IPR007312 (Phosphoesterase), IPR017850 (Alkaline-phosphatase-like, core domain) | GO:0003824 (catalytic activity), GO:0008152 (metabolic process), GO:0016788 (hydrolase activity, acting on ester bonds) |
| Niben101Scf05298g06005.1 | 1509 | 548 | -1.46 | 0.047042367 | AT1G56220.3 | Dormancy/auxin associated family protein LENGTH=140 | IPR008406 (Dormancyauxin associated) |  |
| Niben101Scf09987g02012.1 | 116 | 43 | -1.437 | 0.007461525 | emb\|CDY54019.1\| | BnaCnng26020D [Brassica napus] |  |  |
| Niben101Scf08266g00003.1 | 41 | 15 | -1.435 | 0.047147775 | sp\|O23482\|OPT3_ARATH | Oligopeptide transporter 3 | IPR004813 (Oligopeptide transporter, OPT superfamily) | GO:0055085 (transmembrane transport) |
| Niben101Scf04071g00009.1 | 33 | 12 | -1.413 | 0.040870656 | AT2G44430.1 | DNA-binding bromodomain-containing protein LENGTH=646 | IPR001487 (Bromodomain), IPR009057 (Homeodomain-like) | GO:0003677 (DNA binding), GO:0003682 (chromatin binding), GO:0005515 (protein binding) |
| Niben101Scf00557g02001.1 | 50 | 19 | -1.405 | 0.029743909 | sp\|A7FUZ6\|MOAA_CLOB1 | Cyclic pyranopterin monophosphate synthase | IPR007197 (Radical SAM), IPR013483 (Molybdenum cofactor biosynthesis protein A) | GO:0003824 (catalytic activity), GO:0006777 (Mo-molybdopterin cofactor biosynthetic process), GO:0019008 (molybdopterin synthase complex), GO:0046872 (metal ion binding), GO:0051536 (iron-sulfur cluster binding), GO:0051539 (4 iron, 4 sulfur cluster binding) |
| Niben101Scf03926g00001.1 | 204 | 77 | -1.404 | 0.00488511 | sp\|Q93VH2\|BBD2_ARATH | Bifunctional nuclease 2 | IPR003729 (Bifunctional nuclease domain) | GO:0004518 (nuclease activity) |
| Niben101Scf00536g05021.1 | 34 | 13 | -1.391 | 0.049069674 | AT5G48850.1 | Tetratricopeptide repeat (TPR)-like superfamily protein LENGTH=306 | IPR011990 (Tetratricopeptide-like helical domain) | GO:0005515 (protein binding) |
| Niben101Scf05713g00004.1 | 55 | 21 | -1.386 | 0.014501877 | sp\|Q9SCV3\|BGAL9_ARATH | Beta-galactosidase 9 | IPR000922 (D-galactoside/L-rhamnose binding SUEL lectin domain), IPR001944 (Glycoside hydrolase, family 35), IPR017853 (Glycoside hydrolase superfamily) | GO:0004553 (hydrolase activity, hydrolyzing O-glycosyl compounds), GO:0005975 (carbohydrate metabolic process), GO:0030246 (carbohydrate binding) |
| Niben101Scf07579g03004.1 | 271 | 104 | -1.385 | 0.019682082 | gb\|KEH31349.1\| | stress up-regulated Nod 19 protein [Medicago truncatula] | IPR011692 (Stress up-regulated Nod 19) |  |
| Niben101Scf18347g00012.1 | 3271 | 1259 | -1.378 | 0.0466297 |  | Unknown protein |  |  |
| Niben101Scf00568g04011.1 | 211 | 81 | -1.377 | 0.011675219 | sp\|B2LMI9\|PSBC_GUIAB | Photosystem II CP43 reaction center protein | IPR000484 (Photosynthetic reaction centre, L/M), IPR000932 (Photosystem antenna protein-like) | GO:0009521 (photosystem), GO:0009523 (photosystem II), GO:0009767 (photosynthetic electron transport chain), GO:0009772 (photosynthetic electron transport in photosystem II), GO:0015979 (photosynthesis), GO:0016020 (membrane), GO:0016168 (chlorophyll binding), GO:0019684 (photosynthesis, light reaction), GO:0030076 (light-harvesting complex), GO:0045156 (electron transporter, transferring electrons within the cyclic electron transport pathway of photosynthesis activity) |
| Niben101Scf30418g00015.1 | 141 | 54 | -1.375 | 0.033125156 | ref\|XP_007009555.1\| | Plant protein 1589 of Uncharacterized protein function [Theobroma cacao] gb\|EOY18365.1\| Plant protein 1589 of Uncharacterized protein function [Theobroma cacao] | IPR006476 (Conserved hypothetical protein CHP01589, plant) | |
| Niben101Scf07339g02015.1 | 661 | 255 | -1.372 | 0.02770556 | sp\|A5UE38\|GLPE_HAEIE | Thiosulfate sulfurtransferase GlpE | IPR001763 (Rhodanese-like domain) |  |
| Niben101Scf13741g02011.1 | 31 | 12 | -1.366 | 0.046751106 | sp\|Q9C6Y3\|CCA11_ARATH | Cyclin-A1-1 | IPR014400 (Cyclin A/B/D/E/F/O) | GO:0000079 (regulation of cyclin-dependent protein serine/threonine kinase activity), GO:0005634 (nucleus), GO:0019901 (protein kinase binding), GO:0051726 (regulation of cell cycle) |
| Niben101Scf07391g04030.1 | 102 | 40 | -1.362 | 0.002422212 | sp\|Q9KDV3\|GALE_BACHD | UDP-glucose 4-epimerase | IPR001509 (NAD-dependent epimerase/dehydratase, N-terminal domain), IPR005886 (UDP-glucose 4-epimerase GalE), IPR025308 (UDP-glucose 4-epimerase C-terminal domain) | GO:0003824 (catalytic activity), GO:0003978 (UDP-glucose 4-epimerase activity), GO:0006012 (galactose metabolic process), GO:0050662 (coenzyme binding) |
| Niben101Scf00747g12004.1 | 171 | 67 | -1.34 | 0.000178118 | sp\|O64982\|PRS7_PRUPE | 26S protease regulatory subunit 7 |  |  |
| Niben101Scf38767g00006.1 | 5882 | 2348 | -1.325 | 0.030355253 | sp\|P20238\|MT1_ERYGU | Metallothionein-like protein 1 | IPR000347 (Metallothionein, family 15, plant) | GO:0046872 (metal ion binding) |
| Niben101Scf02569g00027.1 | 75 | 31 | -1.295 | 0.017932526 | AT4G12290.1 | Copper amine oxidase family protein LENGTH=741 | IPR000269 (Copper amine oxidase) | GO:0005507 (copper ion binding), GO:0008131 (primary amine oxidase activity), GO:0009308 (amine metabolic process), GO:0048038 (quinone binding), GO:0055114 (oxidation-reduction process) |
| Niben101Scf09108g00003.1 | 455 | 186 | -1.286 | 0.012525444 | gb\|ADW66129.1\| | late embryogenesis abundant protein Lea5 [Solanum nigrum] | IPR004926 (Late embryogenesis abundant protein, LEA-5) | GO:0006950 (response to stress) |
| Niben101Scf04654g03001.1 | 1022 | 419 | -1.286 | 0.00065275 | AT3G48140.1 | B12D protein LENGTH=88 | IPR010530 (NADH-ubiquinone reductase complex 1 MLRQ subunit) | |
| Niben101Scf17879g00004.1 | 66 | 27 | -1.276 | 0.02499589 |  | Unknown protein |  |  |
| Niben101Scf02548g06013.1 | 3356 | 1390 | -1.272 | 0.019682082 |  | Unknown protein | IPR006031 (XYPPX repeat) |  |
| Niben101Scf10048g02013.1 | 79 | 33 | -1.266 | 0.020750481 | sp\|Q85A24\|CYB6_ANTFO | Cytochrome b6 | IPR016174 (Di-haem cytochrome, transmembrane), IPR023530 (Cytochrome b6, PetB), IPR027387 (Cytochrome b/b6-like domain) | GO:0005506 (iron ion binding), GO:0009055 (electron carrier activity), GO:0016020 (membrane), GO:0016021 (integral component of membrane), GO:0016491 (oxidoreductase activity), GO:0020037 (heme binding), GO:0022900 (electron transport chain), GO:0022904 (respiratory electron transport chain), GO:0045158 (electron transporter, transferring electrons within cytochrome b6/f complex of photosystem II activity) |
| Niben101Scf31091g00006.1 | 172 | 72 | -1.261 | 0.016870154 |  | Unknown protein |  |  |
| Niben101Scf13371g00003.1 | 163 | 69 | -1.241 | 0.04336653 | AT3G07310.1 | Protein of unknown function (DUF760) LENGTH=368 | IPR008479 (Protein of unknown function DUF760) |  |
| Niben101Scf07326g00003.1 | 57 | 25 | -1.237 | 0.033721483 |  | Unknown protein |  |  |
| Niben101Ctg14642g00001.1 | 23018 | 9765 | -1.237 | 0.042208897 | sp\|Q39459\|MT2_CICAR | Metallothionein-like protein 2 | IPR000347 (Metallothionein, family 15, plant) | GO:0046872 (metal ion binding) |
| Niben101Scf03844g01002.1 | 152 | 65 | -1.225 | 0.003737594 | AT1G10940.2 | Protein kinase superfamily protein LENGTH=371 | IPR011009 (Protein kinase-like domain) | GO:0004672 (protein kinase activity), GO:0004674 (protein serine/threonine kinase activity), GO:0005524 (ATP binding), GO:0006468 (protein phosphorylation), GO:0016772 (transferase activity, transferring phosphorus-containing groups) |
| Niben101Scf01493g00007.1 | 137 | 59 | -1.217 | 0.035805931 | AT1G76130.1 | alpha-amylase-like 2 LENGTH=413 | IPR015902 (Glycoside hydrolase, family 13) | GO:0003824 (catalytic activity), GO:0004556 (alpha-amylase activity), GO:0005509 (calcium ion binding), GO:0005975 (carbohydrate metabolic process), GO:0043169 (cation binding) |
| Niben101Scf00780g03018.1 | 56 | 24 | -1.211 | 0.047093285 |  | Unknown protein |  |  |
| Niben101Scf04430g03012.1 | 195 | 84 | -1.202 | 0.001718749 | ref\|XP_002521818.1\| | conserved hypothetical protein [Ricinus communis] gb\|EEF40628.1\| conserved hypothetical protein [Ricinus communis] | | |
| Niben101Scf09698g01005.1 | 136 | 59 | -1.2 | 0.012118982 | AT2G41870.1 | Remorin family protein LENGTH=274 | IPR005516 (Remorin, C-terminal) |  |
| Niben101Scf05713g05004.1 | 89 | 39 | -1.196 | 0.028037329 | emb\|CDX92863.1\| | BnaC07g41210D [Brassica napus] | IPR006502 (Protein of unknown function DUF506, plant) |  |
| Niben101Scf05158g00008.1 | 67 | 30 | -1.181 | 0.049069674 | sp\|Q9SCV3\|BGAL9_ARATH | Beta-galactosidase 9 | IPR000922 (D-galactoside/L-rhamnose binding SUEL lectin domain), IPR001944 (Glycoside hydrolase, family 35), IPR017853 (Glycoside hydrolase superfamily) | GO:0004553 (hydrolase activity, hydrolyzing O-glycosyl compounds), GO:0005975 (carbohydrate metabolic process), GO:0030246 (carbohydrate binding) |
| Niben101Scf02159g01014.1 | 1372 | 607 | -1.176 | 0.014571586 | sp\|Q9FMK7\|BT1_ARATH | BTB/POZ and TAZ domain-containing protein 1 | IPR000197 (Zinc finger, TAZ-type), IPR011333 (BTB/POZ fold) | GO:0003712 (transcription cofactor activity), GO:0004402 (histone acetyltransferase activity), GO:0005515 (protein binding), GO:0005634 (nucleus), GO:0006355 (regulation of transcription, DNA-templated), GO:0008270 (zinc ion binding) |
| Niben101Scf03022g00004.1 | 147 | 66 | -1.169 | 0.007751665 | sp\|Q8H129\|PPA3_ARATH | Purple acid phosphatase 3 | IPR024927 (Acid phosphatase, type 5), IPR029052 (Metallo-dependent phosphatase-like) | GO:0003993 (acid phosphatase activity), GO:0016787 (hydrolase activity) |
| Niben101Scf04364g01014.1 | 72 | 32 | -1.168 | 0.042364952 | sp\|Q28222\|HSP71_CHLAE | Heat shock 70 kDa protein 1 | IPR013126 (Heat shock protein 70 family), IPR029047 (Heat shock protein 70kD, peptide-binding domain), IPR029048 (Heat shock protein 70kD, C-terminal domain) | |
| Niben101Scf02160g06059.1 | 86 | 38 | -1.161 | 0.017483171 | sp\|P08742\|COX1_MAIZE | Cytochrome c oxidase subunit 1 | IPR000883 (Cytochrome c oxidase subunit I), IPR023615 (Cytochrome c oxidase, subunit I, copper-binding site), IPR023616 (Cytochrome c oxidase, subunit I domain) | GO:0004129 (cytochrome-c oxidase activity), GO:0005506 (iron ion binding), GO:0009055 (electron carrier activity), GO:0009060 (aerobic respiration), GO:0016021 (integral component of membrane), GO:0020037 (heme binding), GO:0055114 (oxidation-reduction process) |
| Niben101Scf02511g05005.1 | 150 | 67 | -1.161 | 0.025471289 | sp\|Q03066\|RPSC_NOSS1 | RNA polymerase sigma-C factor | IPR014284 (RNA polymerase sigma-70 like domain) | GO:0003677 (DNA binding), GO:0003700 (sequence-specific DNA binding transcription factor activity), GO:0006352 (DNA-templated transcription, initiation), GO:0006355 (regulation of transcription, DNA-templated), GO:0016987 (sigma factor activity) |
| Niben101Scf03918g00009.1 | 75 | 33 | -1.152 | 0.034017117 | sp\|Q9LSL9\|PP445_ARATH | Pentatricopeptide repeat-containing protein | IPR002885 (Pentatricopeptide repeat), IPR011990 (Tetratricopeptide-like helical domain) | GO:0005515 (protein binding) |
| Niben101Scf03239g04002.1 | 95 | 42 | -1.148 | 0.037654158 | AT2G46140.1 | Late embryogenesis abundant protein LENGTH=166 | IPR004864 (Late embryogenesis abundant protein, LEA-14), IPR013783 (Immunoglobulin-like fold) | GO:0009269 (response to desiccation) |
| Niben101Scf13268g00002.1 | 243 | 110 | -1.143 | 0.005521117 | sp\|P40972\|PLY_TOBAC | Pectate lyase | IPR011050 (Pectin lyase fold/virulence factor), IPR018082 (AmbAllergen) | |
| Niben101Scf02665g12003.1 | 265 | 121 | -1.133 | 0.029220887 | sp\|Q9ZVI6\|LOR8_ARATH | Protein LURP-one-related 8 | IPR025659 (Tubby C-terminal-like domain) |  |
| Niben101Scf02069g00019.1 | 109 | 50 | -1.126 | 0.021012965 | ref\|WP_011984095.1\| | serine protease [Bacillus cytotoxicus] ref\|YP_001374337.1\| hypothetical protein Bcer98_1016 [Bacillus cytotoxicus NVH 391-98] gb\|ABS21342.1\| SCP-like extracellular [Bacillus cytotoxicus NVH 391-98] | | |
| Niben101Scf00568g04006.1 | 171 | 78 | -1.126 | 0.03734388 | sp\|Q1ACH1\|CYB6_CHAVU | Cytochrome b6 | IPR016174 (Di-haem cytochrome, transmembrane), IPR027387 (Cytochrome b/b6-like domain) | GO:0009055 (electron carrier activity), GO:0016020 (membrane), GO:0016491 (oxidoreductase activity), GO:0022904 (respiratory electron transport chain) |
| Niben101Scf05645g08004.1 | 72 | 33 | -1.124 | 0.029685779 | sp\|P57057\|GLPT_HUMAN | Glycerol-3-phosphate transporter | IPR011701 (Major facilitator superfamily), IPR020846 (Major facilitator superfamily domain) | GO:0005215 (transporter activity), GO:0006810 (transport), GO:0016021 (integral component of membrane), GO:0055085 (transmembrane transport) |
| Niben101Scf00229g00004.1 | 145 | 67 | -1.118 | 0.020599552 | sp\|Q6MDI5\|FTSH_PARUW | ATP-dependent zinc metalloprotease FtsH | IPR003959 (ATPase, AAA-type, core), IPR005936 (Peptidase, FtsH), IPR011546 (Peptidase M41, FtsH extracellular), IPR027417 (P-loop containing nucleoside triphosphate hydrolase) | GO:0004222 (metalloendopeptidase activity), GO:0005524 (ATP binding), GO:0006508 (proteolysis), GO:0008270 (zinc ion binding), GO:0016020 (membrane), GO:0016021 (integral component of membrane) |
| Niben101Scf02004g00014.1 | 193 | 89 | -1.11 | 0.01299456 | gb\|AIU50013.1\| | tunicamycin induced 1, partial [Medicago truncatula] | |  |
| Niben101Scf02482g01016.1 | 164 | 76 | -1.104 | 0.010476828 | AT5G28080.2 | Protein kinase superfamily protein LENGTH=492 | IPR011009 (Protein kinase-like domain) | GO:0004672 (protein kinase activity), GO:0004674 (protein serine/threonine kinase activity), GO:0005524 (ATP binding), GO:0006468 (protein phosphorylation), GO:0016772 (transferase activity, transferring phosphorus-containing groups) |
| Niben101Scf01365g05003.1 | 648 | 302 | -1.1 | 0.013413874 | AT1G09960.1 | sucrose transporter 4 LENGTH=510 | IPR005828 (General substrate transporter), IPR005989 (Sucrose/H+ symporter, plant), IPR020846 (Major facilitator superfamily domain) | GO:0005887 (integral component of plasma membrane), GO:0008515 (sucrose transmembrane transporter activity), GO:0015770 (sucrose transport), GO:0016021 (integral component of membrane), GO:0022857 (transmembrane transporter activity), GO:0055085 (transmembrane transport) |
| Niben101Scf03631g01028.1 | 101 | 48 | -1.079 | 0.042010747 | AT2G30600.5 | BTB/POZ domain-containing protein LENGTH=855 | IPR011333 (BTB/POZ fold), IPR011705 (BTB/Kelch-associated), IPR022041 (Farnesoic acid O-methyl transferase) | GO:0005515 (protein binding) |
| Niben101Scf02990g06003.1 | 7012 | 3327 | -1.075 | 0.030601157 | AT4G37800.1 | xyloglucan endotransglucosylase/hydrolase 7 LENGTH=293 | IPR013320 (Concanavalin A-like lectin/glucanase domain), IPR016455 (Xyloglucan endotransglucosylase/hydrolase) | GO:0004553 (hydrolase activity, hydrolyzing O-glycosyl compounds), GO:0005618 (cell wall), GO:0005975 (carbohydrate metabolic process), GO:0006073 (cellular glucan metabolic process), GO:0016762 (xyloglucan:xyloglucosyl transferase activity), GO:0048046 (apoplast) |
| Niben101Scf06782g01011.1 | 307 | 145 | -1.071 | 0.045763882 | sp\|C5D3F4\|DER_GEOSW | GTPase Der | IPR005225 (Small GTP-binding protein domain), IPR016484 (GTP-binding protein EngA), IPR027417 (P-loop containing nucleoside triphosphate hydrolase) | GO:0005525 (GTP binding) |
| Niben101Scf01617g00005.1 | 125 | 60 | -1.065 | 0.011366567 | sp\|Q6ABX8\|ODPB_LEIXX | Pyruvate dehydrogenase E1 component subunit beta | IPR005475 (Transketolase-like, pyrimidine-binding domain), IPR005476 (Transketolase, C-terminal), IPR009014 (Transketolase, C-terminal/Pyruvate-ferredoxin oxidoreductase, domain II), IPR027110 (Pyruvate dehydrogenase E1 component subunit beta), IPR029061 (Thiamin diphosphate-binding fold) | GO:0003824 (catalytic activity), GO:0004739 (pyruvate dehydrogenase (acetyl-transferring) activity), GO:0006086 (acetyl-CoA biosynthetic process from pyruvate), GO:0008152 (metabolic process) |
| Niben101Scf03403g02027.1 | 271 | 131 | -1.048 | 0.013715994 | emb\|CDY52515.1\| | BnaC03g61080D [Brassica napus] |  |  |
| Niben101Scf00530g06012.1 | 149 | 73 | -1.036 | 0.028889565 | sp\|Q03065\|RPSB_NOSS1 | RNA polymerase sigma-B factor | IPR014284 (RNA polymerase sigma-70 like domain) | GO:0003677 (DNA binding), GO:0003700 (sequence-specific DNA binding transcription factor activity), GO:0006352 (DNA-templated transcription, initiation), GO:0006355 (regulation of transcription, DNA-templated), GO:0016987 (sigma factor activity) |
| Niben101Scf04537g01009.1 | 586 | 286 | -1.032 | 0.037139073 | sp\|O82345\|BAG6_ARATH | BAG family molecular chaperone regulator 6 | IPR000048 (IQ motif, EF-hand binding site), IPR003103 (BAG domain) | GO:0005515 (protein binding), GO:0051087 (chaperone binding) |
| Niben101Scf12216g00004.1 | 142 | 70 | -1.031 | 0.013122007 | emb\|CDY45913.1\| | BnaAnng07940D [Brassica napus] |  |  |
| Niben101Scf03985g04028.1 | 317 | 156 | -1.019 | 0.046484191 | sp\|Q84MC7\|PYL9_ARATH | Abscisic acid receptor PYL9 | IPR019587 (Polyketide cyclase/dehydrase), IPR023393 (START-like domain) | |
| Niben101Scf01463g02009.1 | 68 | 33 | -1.017 | 0.043207672 | AT3G13062.2 | Polyketide cyclase/dehydrase and lipid transport superfamily protein LENGTH=440 | IPR002913 (START domain), IPR023393 (START-like domain) | GO:0008289 (lipid binding) |
| Niben101Scf09876g00009.1 | 106 | 53 | -1.011 | 0.024866782 | sp\|Q68F45\|WIPI3_XENLA | WD repeat domain phosphoinositide-interacting protein 3 | IPR015943 (WD40/YVTN repeat-like-containing domain) | GO:0005515 (protein binding) |
| Niben101Scf07395g00031.1 | 103 | 51 | -0.997 | 0.031746008 | sp\|P54211\|PMA1_DUNBI | Plasma membrane ATPase | IPR001757 (P-type ATPase), IPR004014 (Cation-transporting P-type ATPase, N-terminal), IPR023214 (HAD-like domain), IPR023298 (P-type ATPase, transmembrane domain) | GO:0000166 (nucleotide binding), GO:0016021 (integral component of membrane), GO:0046872 (metal ion binding) |
| Niben101Scf12738g00010.1 | 268 | 136 | -0.978 | 0.01658247 | sp\|P00243\|FER_SYNY4 | Ferredoxin | IPR010241 (Ferredoxin [2Fe-2S], plant), IPR012675 (Beta-grasp domain) | GO:0009055 (electron carrier activity), GO:0022900 (electron transport chain), GO:0051536 (iron-sulfur cluster binding), GO:0051537 (2 iron, 2 sulfur cluster binding) |
| Niben101Scf04292g11008.1 | 318 | 163 | -0.962 | 0.013603443 | AT5G56350.1 | Pyruvate kinase family protein LENGTH=498 | IPR001697 (Pyruvate kinase) | GO:0000287 (magnesium ion binding), GO:0003824 (catalytic activity), GO:0004743 (pyruvate kinase activity), GO:0006096 (glycolytic process), GO:0030955 (potassium ion binding) |
| Niben101Scf00308g01005.1 | 848 | 436 | -0.961 | 0.012006048 | AT5G54130.2 | Calcium-binding endonuclease/exonuclease/phosphatase family LENGTH=436 | IPR005135 (Endonuclease/exonuclease/phosphatase), IPR011992 (EF-hand domain pair) | GO:0005509 (calcium ion binding) |
| Niben101Scf02043g02002.1 | 266 | 138 | -0.949 | 0.012655353 | sp\|B9JZ89\|DNAJ_AGRVS | Chaperone protein DnaJ | IPR001623 (DnaJ domain), IPR002939 (Chaperone DnaJ, C-terminal) | GO:0006457 (protein folding), GO:0051082 (unfolded protein binding) |
| Niben101Scf04943g01018.1 | 187 | 97 | -0.939 | 0.028037329 | sp\|Q54BM3\|MCFG_DICDI | Mitochondrial substrate carrier family protein G | IPR002067 (Mitochondrial carrier protein), IPR023395 (Mitochondrial carrier domain) | GO:0055085 (transmembrane transport) |
| Niben101Scf02831g05014.1 | 208 | 109 | -0.935 | 0.025939948 | sp\|Q254M8\|SURE_CHLFF | 5'-nucleotidase SurE | IPR002828 (Survival protein SurE-like phosphatase/nucleotidase) | GO:0016787 (hydrolase activity) |
| Niben101Scf01834g04028.1 | 903 | 487 | -0.89 | 0.024344606 | gb\|ACG41571.1\| | ATPase inhibitor [Zea mays] |  |  |
| Niben101Scf03300g04003.1 | 139 | 76 | -0.88 | 0.048602897 | AT3G19080.1 | SWIB complex BAF60b domain-containing protein LENGTH=462 | IPR003121 (SWIB/MDM2 domain) | GO:0005515 (protein binding) |
| Niben101Scf02807g11007.1 | 302 | 166 | -0.86 | 0.031054746 | AT3G22200.2 | Pyridoxal phosphate (PLP)-dependent transferases superfamily protein LENGTH=513 | IPR005814 (Aminotransferase class-III), IPR015424 (Pyridoxal phosphate-dependent transferase) | GO:0003824 (catalytic activity), GO:0008483 (transaminase activity), GO:0030170 (pyridoxal phosphate binding) |
| Niben101Scf03307g01015.1 | 299 | 166 | -0.856 | 0.034397422 | sp\|Q41142\|PLDA1_RICCO | Phospholipase D alpha 1 | IPR015679 (Phospholipase D family), IPR024632 (Phospholipase D, C-terminal) | GO:0003824 (catalytic activity), GO:0004630 (phospholipase D activity), GO:0005509 (calcium ion binding), GO:0005515 (protein binding), GO:0008152 (metabolic process), GO:0016020 (membrane), GO:0046470 (phosphatidylcholine metabolic process) |
| Niben101Scf09348g00004.1 | 935 | 525 | -0.833 | 0.035855566 | emb\|CBY05740.1\| | ASR2 protein [Solanum chilense] | IPR003496 (ABA/WDS induced protein) | GO:0006950 (response to stress) |
| Niben101Scf09527g01016.1 | 225 | 127 | -0.823 | 0.0466297 | sp\|Q54DB1\|CM2A2_DICDI | Charged multivesicular body protein 2a homolog 2 | IPR005024 (Snf7 family) | GO:0007034 (vacuolar transport) |
| Niben101Scf03731g01008.1 | 260 | 148 | -0.809 | 0.047745067 | AT4G27390.1 | unknown protein; INVOLVED IN: biological_process unknown; LOCATED IN: chloroplast; EXPRESSED IN: 22 plant structures; EXPRESSED DURING: 13 growth stages; Has 30201 Blast hits to 17322 proteins in 780 species: Archae - 12; Bacteria - 1396; Metazoa - 17338; Fungi - 3422; Plants - 5037; Viruses - 0; Other Eukaryotes - 2996 (source: NCBI BLink). LENGTH=235 | | |
| Niben101Scf00069g12017.1 | 1130 | 647 | -0.804 | 0.043850658 | AT5G26780.3 | serine hydroxymethyltransferase 2 LENGTH=533 | IPR001085 (Serine hydroxymethyltransferase), IPR015424 (Pyridoxal phosphate-dependent transferase) | GO:0003824 (catalytic activity), GO:0004372 (glycine hydroxymethyltransferase activity), GO:0006544 (glycine metabolic process), GO:0006563 (L-serine metabolic process), GO:0030170 (pyridoxal phosphate binding) |
| Niben101Scf00063g03007.1 | 948 | 543 | -0.802 | 0.049098137 | sp\|P69039\|IF5A1_NICPL | Eukaryotic translation initiation factor 5A-1 | IPR001884 (Translation elongation factor IF5A) | GO:0003723 (RNA binding), GO:0003746 (translation elongation factor activity), GO:0006452 (translational frameshifting), GO:0043022 (ribosome binding), GO:0045901 (positive regulation of translational elongation), GO:0045905 (positive regulation of translational termination) |
