## Supplemental Table 5 for "VIRP1 bromodomain shapes nuclear condensate formation and has a positive effect on PSTVd accumulation"

**Table S5. Transcriptome analysis of *VIRP1i N. benthamiana* plants.**

a. **Upregulated**

| **ID** | **WT av. reads** | ***VIRP1i* av. reads** | **log2FC** | **padj** | **Blast-Hit-Accession** | **Human-Readable-Description** | **Interpro-ID (Description)** | **Gene-Ontology-ID (Name)** |
| --- | --- | --- | --- | --- | --- | --- | --- | --- |
| Niben101Scf03768g02017.1 | 0 | 10 | 6.465 | 5.81097E-06 | AT5G54090.1 | DNA mismatch repair protein MutS, type 2 LENGTH=796 | NA | NA |
| Niben101Scf03307g06003.1 | 2 | 26 | 3.542 | 0.010917815 | sp\|O82200\|MIOX2_ARATH | Inositol oxygenase 2 | IPR007828 (Inositol oxygenase) | GO:0005506 (iron ion binding), GO:0005737 (cytoplasm), GO:0019310 (inositol catabolic process), GO:0050113 (inositol oxygenase activity), GO:0055114 (oxidation-reduction process) |
| Niben101Scf23113g00009.1 | 1 | 13 | 3.332 | 0.003857205 | emb\|CDX98828.1\| | BnaC09g50070D [Brassica napus] | NA | NA |
| Niben101Scf01240g10022.1 | 1 | 10 | 3.312 | 0.034022357 | NA | Unknown protein | NA | NA |
| Niben101Ctg13782g00003.1 | 5 | 43 | 3.245 | 2.12462E-05 | sp\|P24806\|XTH24_ARATH | Xyloglucan endotransglucosylase/hydrolase protein 24 | IPR013320 (Concanavalin A-like lectin/glucanase domain) | GO:0004553 (hydrolase activity, hydrolyzing O-glycosyl compounds), GO:0005975 (carbohydrate metabolic process) |
| Niben101Scf01237g13004.1 | 4 | 33 | 3.056 | 3.61436E-07 | emb\|CDX98828.1\| | BnaC09g50070D [Brassica napus] | NA | NA |
| Niben101Scf02562g02012.1 | 2 | 13 | 3.051 | 0.000373536 | AT3G11340.1 | UDP-Glycosyltransferase superfamily protein LENGTH=447 | IPR002213 (UDP-glucuronosyl/UDP-glucosyltransferase) | GO:0008152 (metabolic process), GO:0016758 (transferase activity, transferring hexosyl groups) |
| Niben101Scf03814g00003.1 | 2 | 13 | 3.028 | 0.032708416 | NA | Unknown protein | NA | NA |
| Niben101Scf00231g07011.1 | 1 | 10 | 2.973 | 0.001605157 | sp\|Q9ZU49\|LPP1_ARATH | Lipid phosphate phosphatase 1 | IPR000326 (Phosphatidic acid phosphatase type 2/haloperoxidase), IPR002885 (Pentatricopeptide repeat), IPR028681 (Lipid phosphate phosphatase, plant) | GO:0003824 (catalytic activity), GO:0016020 (membrane) |
| Niben101Scf02565g02003.1 | 3 | 16 | 2.375 | 0.000614048 | sp\|Q6K7E6\|ERF1_ORYSJ | Ethylene-responsive transcription factor 1 | IPR003657 (DNA-binding WRKY), IPR016177 (DNA-binding domain) | GO:0003677 (DNA binding), GO:0003700 (sequence-specific DNA binding transcription factor activity), GO:0006355 (regulation of transcription, DNA-templated), GO:0043565 (sequence-specific DNA binding) |
| Niben101Scf09345g00016.1 | 3 | 12 | 2.275 | 0.013238863 | sp\|F4IQJ4\|GASAB_ARATH | Gibberellin-regulated protein 11 | IPR003854 (Gibberellin regulated protein) | NA |
| Niben101Scf02230g03027.1 | 3 | 12 | 2.196 | 0.033674151 | sp\|Q9LVN7\|DPOD1_ARATH | DNA polymerase delta catalytic subunit | IPR006172 (DNA-directed DNA polymerase, family B), IPR023211 (DNA polymerase, palm domain), IPR025687 (C4-type zinc-finger of DNA polymerase delta) | GO:0000166 (nucleotide binding), GO:0003676 (nucleic acid binding), GO:0003677 (DNA binding), GO:0003887 (DNA-directed DNA polymerase activity) |
| Niben101Scf08566g02013.1 | 3 | 13 | 2.172 | 0.005837906 | sp\|Q7XJE6\|MCA1_ARATH | Metacaspase-1 | IPR029030 (Caspase-like domain) | GO:0004197 (cysteine-type endopeptidase activity), GO:0006508 (proteolysis) |
| Niben101Scf17849g00004.1 | 2 | 10 | 2.158 | 0.038846681 | sp\|Q8VID1\|DHRS4_RAT | Dehydrogenase/reductase SDR family member 4 | IPR002347 (Glucose/ribitol dehydrogenase) | GO:0008152 (metabolic process), GO:0016491 (oxidoreductase activity) |
| Niben101Scf03755g00001.1 | 4 | 15 | 2.115 | 0.001773232 | sp\|P55088\|AQP4_MOUSE | Aquaporin-4 | IPR000425 (Major intrinsic protein), IPR023271 (Aquaporin-like) | GO:0005215 (transporter activity), GO:0006810 (transport), GO:0016020 (membrane) |
| Niben101Scf08465g04007.1 | 4 | 14 | 2.022 | 0.0167906 | sp\|Q9C7U5\|E132_ARATH | Glucan endo-1,3-beta-glucosidase 2 | IPR000490 (Glycoside hydrolase, family 17), IPR012946 (X8 domain), IPR017853 (Glycoside hydrolase superfamily) | GO:0004553 (hydrolase activity, hydrolyzing O-glycosyl compounds), GO:0005975 (carbohydrate metabolic process) |
| Niben101Scf10058g00019.1 | 13 | 46 | 2.017 | 1.02853E-06 | sp\|Q84KG5\|CCD_CROSA | Carotenoid 9,10(9',10')-cleavage dioxygenase | IPR004294 (Carotenoid oxygenase) | NA |
| Niben101Scf02839g02012.1 | 32 | 109 | 1.917 | 0.000449914 | AT5G53450.1 | OBP3-responsive gene 1 LENGTH=670 | IPR006843 (Plastid lipid-associated protein/fibrillin conserved domain), IPR011009 (Protein kinase-like domain) | GO:0004672 (protein kinase activity), GO:0005524 (ATP binding), GO:0006468 (protein phosphorylation), GO:0016772 (transferase activity, transferring phosphorus-containing groups) |
| Niben101Scf08291g00009.1 | 4 | 12 | 1.876 | 0.02017392 | sp\|Q7XUN2\|MAD17_ORYSJ | MADS-box transcription factor 17 | IPR002100 (Transcription factor, MADS-box) | GO:0003677 (DNA binding), GO:0046983 (protein dimerization activity) |
| Niben101Scf07539g00001.1 | 5 | 18 | 1.865 | 0.032757289 | sp\|O04161\|AMT12_SOLLC | Ammonium transporter 1 member 2 | IPR001905 (Ammonium transporter), IPR029020 (Ammonium/urea transporter) | GO:0008519 (ammonium transmembrane transporter activity), GO:0015696 (ammonium transport), GO:0016020 (membrane), GO:0072488 (ammonium transmembrane transport) |
| Niben101Scf02449g02005.1 | 12 | 38 | 1.855 | 0.001487226 | emb\|CDY10434.1\| | BnaCnng03350D [Brassica napus] | NA | NA |
| Niben101Scf02738g01016.1 | 29 | 95 | 1.851 | 0.005053075 | sp\|Q6NKX1\|PROD2_ARATH | Proline dehydrogenase 2, mitochondrial | IPR015659 (Proline oxidase), IPR029041 (FAD-linked oxidoreductase-like) | GO:0004657 (proline dehydrogenase activity), GO:0006562 (proline catabolic process) |
| Niben101Scf04738g03003.1 | 4 | 12 | 1.825 | 0.013755176 | AT5G16010.1 | 3-oxo-5-alpha-steroid 4-dehydrogenase family protein LENGTH=268 | IPR001104 (3-oxo-5-alpha-steroid 4-dehydrogenase, C-terminal) | GO:0005737 (cytoplasm), GO:0006629 (lipid metabolic process), GO:0016021 (integral component of membrane), GO:0016627 (oxidoreductase activity, acting on the CH-CH group of donors) |
| Niben101Scf00381g08001.1 | 4 | 14 | 1.811 | 0.006641103 | AT3G51470.1 | Protein phosphatase 2C family protein LENGTH=361 | IPR001932 (Protein phosphatase 2C (PP2C)-like domain), IPR015655 (Protein phosphatase 2C) | GO:0003824 (catalytic activity), GO:0004722 (protein serine/threonine phosphatase activity), GO:0006470 (protein dephosphorylation) |
| Niben101Scf02974g01001.1 | 4 | 14 | 1.794 | 0.022897638 | sp\|Q03015\|NDUS6_NEUCR | NADH-ubiquinone oxidoreductase 12 kDa subunit, mitochondrial | IPR019377 (NADH-ubiquinone oxidoreductase, subunit 10) | NA |
| Niben101Scf01594g00001.1 | 5 | 17 | 1.756 | 0.042100909 | AT5G53590.1 | SAUR-like auxin-responsive protein family LENGTH=142 | IPR003676 (Auxin-induced protein, ARG7) | NA |
| Niben101Scf06112g01008.1 | 100 | 299 | 1.748 | 1.64958E-11 | AT5G66690.1 | UDP-Glycosyltransferase superfamily protein LENGTH=481 | IPR002213 (UDP-glucuronosyl/UDP-glucosyltransferase) | GO:0008152 (metabolic process), GO:0016758 (transferase activity, transferring hexosyl groups) |
| Niben101Scf00367g08003.1 | 72 | 218 | 1.746 | 1.96955E-13 | NA | Unknown protein | NA | NA |
| Niben101Scf11483g00010.1 | 4 | 13 | 1.737 | 0.033737457 | sp\|O59942\|AAP2_NEUCR | Amino-acid permease 2 | IPR002293 (Amino acid/polyamine transporter I), IPR004841 (Amino acid permease/ SLC12A domain) | GO:0003333 (amino acid transmembrane transport), GO:0006810 (transport), GO:0015171 (amino acid transmembrane transporter activity), GO:0016020 (membrane), GO:0055085 (transmembrane transport) |
| Niben101Scf03374g07003.1 | 19 | 58 | 1.722 | 8.49798E-06 | sp\|Q4QJM6\|GLND_HAEI8 | Bifunctional uridylyltransferase/uridylyl-removing enzyme | IPR002912 (ACT domain) | GO:0008152 (metabolic process), GO:0016597 (amino acid binding) |
| Niben101Scf00823g00002.1 | 13 | 40 | 1.712 | 0.034022357 | sp\|Q7G8Y5\|GRXC1_ORYSJ | Glutaredoxin-C1 | IPR011905 (Glutaredoxin-like, plant II), IPR012336 (Thioredoxin-like fold) | GO:0009055 (electron carrier activity), GO:0015035 (protein disulfide oxidoreductase activity), GO:0045454 (cell redox homeostasis) |
| Niben101Scf05862g00017.1 | 9 | 26 | 1.71 | 0.002446666 | sp\|A8GCS6\|NQOR_SERP5 | NAD(P)H dehydrogenase (quinone) | IPR029039 (Flavoprotein-like) | GO:0010181 (FMN binding), GO:0016491 (oxidoreductase activity) |
| Niben101Scf02290g05024.1 | 95 | 281 | 1.708 | 0.000798321 | AT5G53450.1 | OBP3-responsive gene 1 LENGTH=670 | NA | NA |
| Niben101Scf08266g00003.1 | 32 | 95 | 1.708 | 0.002683524 | sp\|O23482\|OPT3_ARATH | Oligopeptide transporter 3 | IPR004813 (Oligopeptide transporter, OPT superfamily) | GO:0055085 (transmembrane transport) |
| Niben101Scf03307g06009.1 | 4 | 13 | 1.708 | 0.041940866 | sp\|B0S230\|PFKA_FINM2 | ATP-dependent 6-phosphofructokinase | IPR000023 (Phosphofructokinase domain), IPR012004 (Pyrophosphate-dependent phosphofructokinase TP0108), IPR022953 (Phosphofructokinase) | GO:0003872 (6-phosphofructokinase activity), GO:0005524 (ATP binding), GO:0006002 (fructose 6-phosphate metabolic process), GO:0006096 (glycolytic process) |
| Niben101Scf06684g03012.1 | 5 | 15 | 1.691 | 0.048114483 | sp\|Q2HBI0\|TOF1_CHAGB | Topoisomerase 1-associated factor 1 | IPR006906 (Timeless protein), IPR007725 (Timeless C-terminal) | NA |
| Niben101Scf10058g00019.1 | 5 | 14 | 1.674 | 0.016227779 | sp\|Q84KG5\|CCD_CROSA | Carotenoid 9,10(9',10')-cleavage dioxygenase | IPR004294 (Carotenoid oxygenase) | NA |
| Niben101Scf07296g00001.1 | 4 | 12 | 1.664 | 0.036533653 | ref\|XP_002510247.1\| | conserved hypothetical protein [Ricinus communis] gb\|EEF52434.1\| conserved hypothetical protein [Ricinus communis] | NA | NA |
| Niben101Scf02044g12019.1 | 4 | 11 | 1.66 | 0.038563996 | NA | Unknown protein | NA | NA |
| Niben101Scf04405g00007.1 | 62 | 173 | 1.658 | 6.68891E-11 | sp\|Q339N5\|CSLH1_ORYSJ | Cellulose synthase-like protein H1 | IPR005150 (Cellulose synthase), IPR029044 (Nucleotide-diphospho-sugar transferases) | GO:0016020 (membrane), GO:0016760 (cellulose synthase (UDP-forming) activity), GO:0030244 (cellulose biosynthetic process) |
| Niben101Scf02348g00008.1 | 17 | 48 | 1.655 | 0.001212639 | AT4G24510.1 | HXXXD-type acyl-transferase family protein LENGTH=421 | IPR003480 (Transferase), IPR023213 (Chloramphenicol acetyltransferase-like domain) | GO:0016747 (transferase activity, transferring acyl groups other than amino-acyl groups) |
| Niben101Scf05283g00018.1 | 5 | 14 | 1.646 | 0.043123656 | sp\|P83774\|GBLP_CANAL | Guanine nucleotide-binding protein subunit beta-like protein | IPR015943 (WD40/YVTN repeat-like-containing domain), IPR020472 (G-protein beta WD-40 repeat) | GO:0005515 (protein binding) |
| Niben101Scf00712g14003.1 | 5 | 14 | 1.61 | 0.044331393 | AT1G20640.1 | Plant regulator RWP-RK family protein LENGTH=844 | IPR000270 (Phox/Bem1p), IPR003035 (RWP-RK domain) | GO:0005515 (protein binding) |
| Niben101Scf08977g00006.1 | 6 | 16 | 1.601 | 0.030074505 | emb\|CDY56816.1\| | BnaCnng31190D [Brassica napus] | IPR024489 (Organ specific protein) | NA |
| Niben101Scf03307g04014.1 | 11 | 30 | 1.586 | 0.00168661 | sp\|P27061\|PPA1_SOLLC | Acid phosphatase 1 | IPR005519 (Acid phosphatase (Class B)), IPR023214 (HAD-like domain) | GO:0003993 (acid phosphatase activity) |
| Niben101Scf08873g01026.1 | 4 | 11 | 1.581 | 0.040139764 | sp\|Q8GUH1\|PUB33_ARATH | U-box domain-containing protein 33 | IPR014729 (Rossmann-like alpha/beta/alpha sandwich fold) | GO:0006950 (response to stress) |
| Niben101Scf01269g01018.1 | 50 | 135 | 1.57 | 1.31958E-11 | sp\|Q9SZ42\|FAD4_ARATH | Fatty acid desaturase 4, chloroplastic | IPR019547 (B-domain containing protein Kua) | NA |
| Niben101Scf09260g03002.1 | 12 | 32 | 1.57 | 0.006641103 | AT1G27320.1 | histidine kinase 3 LENGTH=1036 | IPR003594 (Histidine kinase-like ATPase, C-terminal domain), IPR004358 (Signal transduction histidine kinase-related protein, C-terminal), IPR005467 (Signal transduction histidine kinase, core), IPR011006 (CheY-like superfamily) | GO:0000160 (phosphorelay signal transduction system), GO:0016310 (phosphorylation), GO:0016772 (transferase activity, transferring phosphorus-containing groups) |
| Niben101Scf03263g03004.1 | 6 | 17 | 1.57 | 0.040139764 | sp\|Q9ZW24\|GSTU7_ARATH | Glutathione S-transferase U7 | IPR010987 (Glutathione S-transferase, C-terminal-like), IPR012336 (Thioredoxin-like fold) | GO:0005515 (protein binding) |
| Niben101Scf05584g02018.1 | 12 | 32 | 1.561 | 0.000705968 | sp\|Q9FK13\|PDV1_ARATH | Plastid division protein PDV1 | NA | NA |
| Niben101Scf00148g01023.1 | 11 | 29 | 1.561 | 0.011206409 | AT5G62360.1 | Plant invertase/pectin methylesterase inhibitor superfamily protein LENGTH=203 | IPR006501 (Pectinesterase inhibitor domain) | GO:0004857 (enzyme inhibitor activity), GO:0030599 (pectinesterase activity) |
| Niben101Scf01847g01001.1 | 21 | 54 | 1.56 | 0.017619865 | NA | Unknown protein | NA | NA |
| Niben101Scf11491g02037.1 | 8 | 20 | 1.56 | 0.036797383 | sp\|Q74MI6\|EF1A_NANEQ | Elongation factor 1-alpha | IPR009000 (Translation protein, beta-barrel domain), IPR009001 (Translation elongation factor EF1A/initiation factor IF2gamma, C-terminal) | GO:0005525 (GTP binding) |
| Niben101Scf04407g02027.1 | 9 | 25 | 1.553 | 0.044331393 | sp\|Q8H157\|PTR19_ARATH | Protein NRT1/ PTR FAMILY 4.6 | IPR000109 (Proton-dependent oligopeptide transporter family), IPR020846 (Major facilitator superfamily domain) | GO:0005215 (transporter activity), GO:0006810 (transport), GO:0016020 (membrane) |
| Niben101Scf04992g00013.1 | 20 | 52 | 1.551 | 0.000145798 | AT1G14890.1 | Plant invertase/pectin methylesterase inhibitor superfamily protein LENGTH=219 | IPR006501 (Pectinesterase inhibitor domain) | GO:0004857 (enzyme inhibitor activity), GO:0030599 (pectinesterase activity) |
| Niben101Scf00324g02027.1 | 10 | 28 | 1.541 | 0.006241798 | sp\|Q9SF78\|GDL29_ARATH | GDSL esterase/lipase | IPR013830 (SGNH hydrolase-type esterase domain) | NA |
| Niben101Scf18639g00026.1 | 10 | 27 | 1.541 | 0.000373536 | AT5G23060.1 | calcium sensing receptor LENGTH=387 | IPR001763 (Rhodanese-like domain) | NA |
| Niben101Scf02854g10010.1 | 8 | 22 | 1.539 | 0.004541414 | NA | Unknown protein | IPR001611 (Leucine-rich repeat), IPR003591 (Leucine-rich repeat, typical subtype), IPR025875 (Leucine rich repeat 4) | GO:0005515 (protein binding) |
| Niben101Scf24249g00001.1 | 37 | 97 | 1.528 | 0.000211593 | sp\|Q9SN74\|BH047_ARATH | Transcription factor bHLH47 | IPR011598 (Myc-type, basic helix-loop-helix (bHLH) domain) | GO:0046983 (protein dimerization activity) |
| Niben101Scf08195g07024.1 | 24 | 61 | 1.524 | 1.81995E-06 | sp\|Q9C7N4\|GDL15_ARATH | GDSL esterase/lipase | IPR013830 (SGNH hydrolase-type esterase domain) | GO:0006629 (lipid metabolic process), GO:0016788 (hydrolase activity, acting on ester bonds) |
| Niben101Scf01063g02002.1 | 24 | 63 | 1.52 | 3.51022E-06 | sp\|Q9S9N9\|CCR1_ARATH | Cinnamoyl-CoA reductase 1 | IPR016040 (NAD(P)-binding domain) | NA |
| Niben101Scf05034g05002.1 | 7 | 18 | 1.503 | 0.025641344 | sp\|P0CG15\|CTF8_MOUSE | Chromosome transmission fidelity protein 8 homolog | IPR018607 (Chromosome transmission fidelity protein 8) | NA |
| Niben101Scf01782g11026.1 | 7 | 17 | 1.471 | 0.034748243 | sp\|P0A4H0\|SECY_SYNE7 | Protein translocase subunit SecY | IPR002208 (SecY/SEC61-alpha family), IPR023201 (SecY subunit domain) | GO:0015031 (protein transport), GO:0016020 (membrane) |
| Niben101Scf13823g03007.1 | 10 | 25 | 1.467 | 0.010758452 | AT2G36870.1 | xyloglucan endotransglucosylase/hydrolase 32 LENGTH=299 | IPR013320 (Concanavalin A-like lectin/glucanase domain), IPR016455 (Xyloglucan endotransglucosylase/hydrolase) | GO:0004553 (hydrolase activity, hydrolyzing O-glycosyl compounds), GO:0005618 (cell wall), GO:0005975 (carbohydrate metabolic process), GO:0006073 (cellular glucan metabolic process), GO:0016762 (xyloglucan:xyloglucosyl transferase activity), GO:0048046 (apoplast) |
| Niben101Scf07304g07013.1 | 5 | 13 | 1.467 | 0.041531077 | NA | Unknown protein | NA | NA |
| Niben101Scf01500g00011.1 | 11 | 27 | 1.461 | 0.003241436 | AT1G14890.1 | Plant invertase/pectin methylesterase inhibitor superfamily protein LENGTH=219 | IPR006501 (Pectinesterase inhibitor domain) | GO:0004857 (enzyme inhibitor activity), GO:0030599 (pectinesterase activity) |
| Niben101Scf16068g00007.1 | 22 | 54 | 1.46 | 7.15038E-05 | sp\|B5F4U8\|RPPH_SALA4 | RNA pyrophosphohydrolase | IPR015797 (NUDIX hydrolase domain-like) | GO:0016787 (hydrolase activity) |
| Niben101Scf03326g04010.1 | 19 | 47 | 1.46 | 5.28628E-05 | sp\|Q9S9N9\|CCR1_ARATH | Cinnamoyl-CoA reductase 1 | IPR016040 (NAD(P)-binding domain) | NA |
| Niben101Scf12330g02013.1 | 118 | 289 | 1.441 | 5.99374E-05 | sp\|O23482\|OPT3_ARATH | Oligopeptide transporter 3 | IPR004813 (Oligopeptide transporter, OPT superfamily) | GO:0055085 (transmembrane transport) |
| Niben101Scf06942g02033.1 | 16 | 38 | 1.422 | 0.003614312 | sp\|Q56YT0\|LAC3_ARATH | Laccase-3 | IPR017761 (Laccase) | GO:0005507 (copper ion binding), GO:0016491 (oxidoreductase activity), GO:0046274 (lignin catabolic process), GO:0048046 (apoplast), GO:0052716 (hydroquinone:oxygen oxidoreductase activity), GO:0055114 (oxidation-reduction process) |
| Niben101Scf11709g01031.1 | 8 | 19 | 1.422 | 0.025662251 | sp\|Q6ID77\|MED11_ARATH | Mediator of RNA polymerase II transcription subunit 11 | IPR019404 (Mediator complex, subunit Med11) | GO:0001104 (RNA polymerase II transcription cofactor activity), GO:0006357 (regulation of transcription from RNA polymerase II promoter), GO:0016592 (mediator complex) |
| Niben101Scf04509g01021.1 | 22 | 54 | 1.414 | 0.009546368 | NA | Unknown protein | NA | NA |
| Niben101Scf07509g01001.1 | 8 | 19 | 1.412 | 0.029855212 | AT5G03720.1 | heat shock transcription factor A3 LENGTH=412 | IPR011991 (Winged helix-turn-helix DNA-binding domain), IPR027725 (Heat shock transcription factor family) | GO:0003700 (sequence-specific DNA binding transcription factor activity), GO:0005634 (nucleus), GO:0006355 (regulation of transcription, DNA-templated), GO:0009408 (response to heat), GO:0043565 (sequence-specific DNA binding) |
| Niben101Scf13518g00013.1 | 39 | 93 | 1.408 | 3.70758E-08 | sp\|Q9SZ42\|FAD4_ARATH | Fatty acid desaturase 4, chloroplastic | IPR019547 (B-domain containing protein Kua) | NA |
| Niben101Scf05047g04012.1 | 24 | 56 | 1.392 | 0.000184738 | sp\|P25802\|CYSP1_OSTOS | Cathepsin B-like cysteine proteinase 1 | IPR013128 (Peptidase C1A), IPR025660 (Cysteine peptidase, histidine active site), IPR025661 (Cysteine peptidase, asparagine active site) | GO:0006508 (proteolysis), GO:0008234 (cysteine-type peptidase activity) |
| Niben101Scf00448g18001.1 | 87 | 208 | 1.384 | 0.000946172 | sp\|P09789\|GRP1_PETHY | Glycine-rich cell wall structural protein 1 | NA | NA |
| Niben101Scf02819g02008.1 | 8 | 20 | 1.377 | 0.028591545 | sp\|Q9LND8\|ADSL2_ARATH | Delta-9 desaturase-like 2 protein | IPR015876 (Fatty acid desaturase, type 1, core) | GO:0006629 (lipid metabolic process), GO:0016717 (oxidoreductase activity, acting on paired donors, with oxidation of a pair of donors resulting in the reduction of molecular oxygen to two molecules of water), GO:0055114 (oxidation-reduction process) |
| Niben101Scf20887g00008.1 | 9 | 20 | 1.362 | 0.014626958 | sp\|A2Y5R6\|EXPA4_ORYSI | Expansin-A4 | IPR007118 (Expansin/Lol pI) | GO:0005576 (extracellular region), GO:0009664 (plant-type cell wall organization) |
| Niben101Scf08195g07024.1 | 16 | 37 | 1.355 | 0.010183716 | sp\|Q9C7N4\|GDL15_ARATH | GDSL esterase/lipase | IPR013830 (SGNH hydrolase-type esterase domain) | GO:0006629 (lipid metabolic process), GO:0016788 (hydrolase activity, acting on ester bonds) |
| Niben101Scf03595g08008.1 | 15 | 34 | 1.345 | 0.037303049 | sp\|Q0WPR4\|SCP34_ARATH | Serine carboxypeptidase-like 34 | IPR001563 (Peptidase S10, serine carboxypeptidase), IPR029058 (Alpha/Beta hydrolase fold) | GO:0004185 (serine-type carboxypeptidase activity), GO:0006508 (proteolysis) |
| Niben101Scf10287g01017.1 | 157 | 354 | 1.344 | 4.68535E-10 | sp\|Q9LVE0\|PTR33_ARATH | Protein NRT1/ PTR FAMILY 6.4 | IPR000109 (Proton-dependent oligopeptide transporter family), IPR020846 (Major facilitator superfamily domain) | GO:0005215 (transporter activity), GO:0006810 (transport), GO:0016020 (membrane) |
| Niben101Scf09401g01007.1 | 66 | 149 | 1.335 | 1.20191E-06 | tpg\|DAA57049.1\| | TPA: hypothetical protein ZEAMMB73_793304 [Zea mays] | NA | NA |
| Niben101Scf11790g01008.1 | 20 | 46 | 1.333 | 0.012716326 | AT1G73190.1 | Aquaporin-like superfamily protein LENGTH=268 | IPR000425 (Major intrinsic protein), IPR023271 (Aquaporin-like) | GO:0005215 (transporter activity), GO:0006810 (transport), GO:0016020 (membrane) |
| Niben101Scf07817g02001.1 | 17 | 40 | 1.333 | 0.004118614 | AT5G07990.1 | Cytochrome P450 superfamily protein LENGTH=513 | IPR001128 (Cytochrome P450) | GO:0005506 (iron ion binding), GO:0016705 (oxidoreductase activity, acting on paired donors, with incorporation or reduction of molecular oxygen), GO:0020037 (heme binding), GO:0055114 (oxidation-reduction process) |
| Niben101Scf05553g06001.1 | 38 | 87 | 1.323 | 0.000129816 | sp\|Q9ZPE7\|EXO_ARATH | Protein EXORDIUM | IPR006766 (Phosphate-induced protein 1) | NA |
| Niben101Scf03227g03017.1 | 10 | 23 | 1.319 | 0.008115048 | sp\|P47735\|RLK5_ARATH | Receptor-like protein kinase 5 | IPR001611 (Leucine-rich repeat), IPR011009 (Protein kinase-like domain), IPR013320 (Concanavalin A-like lectin/glucanase domain) | GO:0004672 (protein kinase activity), GO:0005515 (protein binding), GO:0005524 (ATP binding), GO:0006468 (protein phosphorylation), GO:0016772 (transferase activity, transferring phosphorus-containing groups) |
| Niben101Scf02909g09004.1 | 13 | 29 | 1.307 | 0.006205809 | gb\|AGT97382.1\| | EG5651 [Manihot esculenta] | NA | NA |
| Niben101Scf33912g00006.1 | 10 | 23 | 1.301 | 0.02443857 | AT2G41810.1 | Protein of unknown function, DUF642 LENGTH=370 | IPR006946 (Protein of unknown function DUF642), IPR008979 (Galactose-binding domain-like) | NA |
| Niben101Scf10927g00023.1 | 77 | 169 | 1.292 | 9.71165E-09 | sp\|Q54M20\|TTC4_DICDI | Tetratricopeptide repeat protein 4 homolog | IPR011990 (Tetratricopeptide-like helical domain) | GO:0005515 (protein binding) |
| Niben101Scf13227g00026.1 | 85 | 190 | 1.291 | 8.15775E-07 | sp\|Q56254\|RPOM_THECE | DNA-directed RNA polymerase subunit M | IPR001222 (Zinc finger, TFIIS-type), IPR001529 (DNA-directed RNA polymerase, M/15kDa subunit), IPR012164 (DNA-directed RNA polymerase, subunit C11/M/9) | GO:0003676 (nucleic acid binding), GO:0003677 (DNA binding), GO:0003899 (DNA-directed RNA polymerase activity), GO:0006351 (transcription, DNA-templated), GO:0008270 (zinc ion binding) |
| Niben101Scf09648g03008.1 | 14 | 30 | 1.291 | 0.003688236 | AT5G49220.1 | Protein of unknown function (DUF789) LENGTH=409 | IPR008507 (Protein of unknown function DUF789) | NA |
| Niben101Scf00078g02001.1 | 16 | 35 | 1.281 | 0.028434722 | sp\|P49210\|RL9_ORYSJ | 60S ribosomal protein L9 | IPR000702 (Ribosomal protein L6) | GO:0003735 (structural constituent of ribosome), GO:0005622 (intracellular), GO:0005840 (ribosome), GO:0006412 (translation), GO:0019843 (rRNA binding) |
| Niben101Scf02537g06002.1 | 29 | 63 | 1.267 | 0.000132211 | sp\|Q9ZPE7\|EXO_ARATH | Protein EXORDIUM | IPR006766 (Phosphate-induced protein 1) | NA |
| Niben101Scf04915g00003.1 | 10 | 21 | 1.264 | 0.023228639 | AT4G13050.1 | Acyl-ACP thioesterase LENGTH=367 | IPR002864 (Acyl-ACP thioesterase), IPR029069 (HotDog domain) | GO:0006633 (fatty acid biosynthetic process), GO:0016790 (thiolester hydrolase activity) |
| Niben101Scf02335g03005.1 | 35 | 75 | 1.263 | 0.001632035 | sp\|Q9ZQI8\|NLTL2_ARATH | Non-specific lipid-transfer protein-like protein | IPR000528 (Plant lipid transfer protein/Par allergen), IPR016140 (Bifunctional inhibitor/plant lipid transfer protein/seed storage helical domain) | GO:0006869 (lipid transport), GO:0008289 (lipid binding) |
| Niben101Scf04933g00011.1 | 15 | 32 | 1.261 | 0.007407961 | AT2G31010.1 | Protein kinase superfamily protein LENGTH=775 | IPR011009 (Protein kinase-like domain) | GO:0004672 (protein kinase activity), GO:0004674 (protein serine/threonine kinase activity), GO:0005524 (ATP binding), GO:0006468 (protein phosphorylation), GO:0016772 (transferase activity, transferring phosphorus-containing groups) |
| Niben101Scf06263g01004.1 | 23 | 48 | 1.243 | 0.000457361 | AT4G35640.1 | serine acetyltransferase 3;2 LENGTH=355 | IPR005881 (Serine O-acetyltransferase) | GO:0005737 (cytoplasm), GO:0006535 (cysteine biosynthetic process from serine), GO:0009001 (serine O-acetyltransferase activity), GO:0016740 (transferase activity) |
| Niben101Scf05737g00003.1 | 22 | 45 | 1.241 | 0.048445199 | sp\|O78502\|PSAD_GUITH | Photosystem I reaction center subunit II | IPR003685 (Photosystem I PsaD) | GO:0009522 (photosystem I), GO:0009538 (photosystem I reaction center), GO:0015979 (photosynthesis) |
| Niben101Scf07714g00008.1 | 21 | 45 | 1.237 | 0.042780377 | sp\|B9LNJ8\|HIS8_HALLT | Histidinol-phosphate aminotransferase | IPR021178 (Tyrosine transaminase) | GO:0003824 (catalytic activity), GO:0006520 (cellular amino acid metabolic process), GO:0008483 (transaminase activity), GO:0009058 (biosynthetic process), GO:0030170 (pyridoxal phosphate binding) |
| Niben101Scf08819g00017.1 | 10 | 22 | 1.236 | 0.039751815 | prf\|\|1718316A | granule-bound starch synthase | IPR011835 (Glycogen/starch synthase, ADP-glucose type), IPR029056 (Ribokinase-like) | GO:0009011 (starch synthase activity), GO:0009058 (biosynthetic process), GO:0009250 (glucan biosynthetic process) |
| Niben101Scf14996g00009.1 | 143 | 292 | 1.234 | 0.030797601 | sp\|Q9M5L6\|CATA_CAPAN | Catalase | IPR002226 (Catalase haem-binding site), IPR010582 (Catalase immune-responsive domain), IPR011614 (Catalase core domain), IPR018028 (Catalase, mono-functional, haem-containing), IPR020835 (Catalase-like domain), IPR024708 (Catalase active site) | GO:0004096 (catalase activity), GO:0006979 (response to oxidative stress), GO:0020037 (heme binding), GO:0055114 (oxidation-reduction process) |
| Niben101Scf02080g02004.1 | 12 | 26 | 1.229 | 0.012716326 | gb\|AES74971.2\| | membrane protein [Medicago truncatula] | IPR018710 (Protein of unknown function DUF2232) | NA |
| Niben101Scf02583g02013.1 | 4225 | 8678 | 1.206 | 9.06038E-16 | sp\|Q06721\|RCA_ANASC | Ribulose bisphosphate carboxylase/oxygenase activase | IPR003959 (ATPase, AAA-type, core), IPR027417 (P-loop containing nucleoside triphosphate hydrolase) | GO:0005524 (ATP binding) |
| Niben101Scf08368g00010.1 | 33 | 68 | 1.2 | 7.05673E-05 | sp\|Q9ZUA5\|VIT1_ARATH | Vacuolar iron transporter 1 | IPR008217 (Ccc1 family) | NA |
| Niben101Scf15538g00013.1 | 14 | 28 | 1.199 | 0.037923914 | sp\|Q8GXV5\|UGHY_ARATH | (S)-ureidoglycine aminohydrolase | IPR014710 (RmlC-like jelly roll fold) | NA |
| Niben101Scf04306g00002.1 | 14 | 29 | 1.198 | 0.032193971 | sp\|Q9CHL8\|LMRA_LACLA | Multidrug resistance ABC transporter ATP-binding and permease protein | IPR003439 (ABC transporter-like), IPR011527 (ABC transporter type 1, transmembrane domain), IPR027417 (P-loop containing nucleoside triphosphate hydrolase) | GO:0005524 (ATP binding), GO:0006810 (transport), GO:0016021 (integral component of membrane), GO:0016887 (ATPase activity), GO:0042626 (ATPase activity, coupled to transmembrane movement of substances), GO:0055085 (transmembrane transport) |
| Niben101Scf00776g02002.1 | 74 | 154 | 1.196 | 0.000117985 | tpg\|DAA57049.1\| | TPA: hypothetical protein ZEAMMB73_793304 [Zea mays] | NA | NA |
| Niben101Scf00996g02009.1 | 49 | 99 | 1.187 | 0.004044052 | sp\|Q38857\|XTH22_ARATH | Xyloglucan endotransglucosylase/hydrolase protein 22 | IPR008264 (Beta-glucanase), IPR013320 (Concanavalin A-like lectin/glucanase domain), IPR016455 (Xyloglucan endotransglucosylase/hydrolase) | GO:0004553 (hydrolase activity, hydrolyzing O-glycosyl compounds), GO:0005618 (cell wall), GO:0005975 (carbohydrate metabolic process), GO:0006073 (cellular glucan metabolic process), GO:0016762 (xyloglucan:xyloglucosyl transferase activity), GO:0048046 (apoplast) |
| Niben101Scf00279g01014.1 | 35 | 71 | 1.182 | 0.001371951 | sp\|Q9FLG1\|BXL4_ARATH | Beta-D-xylosidase 4 | IPR002772 (Glycoside hydrolase family 3 C-terminal domain), IPR017853 (Glycoside hydrolase superfamily), IPR026891 (Fibronectin type III-like domain), IPR026892 (Glycoside hydrolase family 3) | GO:0004553 (hydrolase activity, hydrolyzing O-glycosyl compounds), GO:0005975 (carbohydrate metabolic process) |
| Niben101Scf06008g00001.1 | 18 | 38 | 1.182 | 0.036001425 | sp\|Q9SD82\|RFA1B_ARATH | Replication protein A 70 kDa DNA-binding subunit B | IPR004591 (Replication factor A protein 1) | GO:0003676 (nucleic acid binding), GO:0003677 (DNA binding), GO:0005634 (nucleus), GO:0006260 (DNA replication), GO:0006281 (DNA repair), GO:0006310 (DNA recombination) |
| Niben101Scf03283g04012.1 | 77 | 159 | 1.181 | 0.000316272 | AT5G46800.1 | Mitochondrial substrate carrier family protein LENGTH=300 | IPR018108 (Mitochondrial substrate/solute carrier), IPR023395 (Mitochondrial carrier domain) | NA |
| Niben101Scf02583g02013.1 | 139 | 282 | 1.172 | 6.68891E-11 | sp\|Q06721\|RCA_ANASC | Ribulose bisphosphate carboxylase/oxygenase activase | IPR003959 (ATPase, AAA-type, core), IPR027417 (P-loop containing nucleoside triphosphate hydrolase) | GO:0005524 (ATP binding) |
| Niben101Scf32212g00015.1 | 18 | 36 | 1.165 | 0.017135045 | sp\|B1Q3J6\|DNM1B_ORYSJ | DNA (cytosine-5)-methyltransferase 1B | IPR001525 (C-5 cytosine methyltransferase), IPR029063 (S-adenosyl-L-methionine-dependent methyltransferase) | GO:0003677 (DNA binding), GO:0003682 (chromatin binding), GO:0003886 (DNA (cytosine-5-)-methyltransferase activity), GO:0005634 (nucleus), GO:0006306 (DNA methylation), GO:0008168 (methyltransferase activity), GO:0090116 (C-5 methylation of cytosine) |
| Niben101Scf00390g02001.1 | 16 | 33 | 1.158 | 0.027479665 | sp\|Q6NKX1\|PROD2_ARATH | Proline dehydrogenase 2, mitochondrial | IPR015659 (Proline oxidase), IPR029041 (FAD-linked oxidoreductase-like) | GO:0004657 (proline dehydrogenase activity), GO:0006562 (proline catabolic process) |
| Niben101Scf04779g01006.1 | 39 | 78 | 1.157 | 0.000792199 | sp\|Q2IJ93\|EFG1_ANADE | Elongation factor G 1 | IPR000795 (Elongation factor, GTP-binding domain), IPR005225 (Small GTP-binding protein domain), IPR009000 (Translation protein, beta-barrel domain), IPR027417 (P-loop containing nucleoside triphosphate hydrolase) | GO:0003924 (GTPase activity), GO:0005525 (GTP binding) |
| Niben101Scf01139g00001.1 | 54 | 106 | 1.142 | 6.98969E-05 | AT5G65380.1 | MATE efflux family protein LENGTH=486 | IPR002528 (Multi antimicrobial extrusion protein) | GO:0006855 (drug transmembrane transport), GO:0015238 (drug transmembrane transporter activity), GO:0015297 (antiporter activity), GO:0016020 (membrane), GO:0055085 (transmembrane transport) |
| Niben101Scf01526g08008.1 | 20 | 39 | 1.14 | 0.026776581 | AT5G46640.1 | AT hook motif DNA-binding family protein LENGTH=386 | IPR005175 (Domain of unknown function DUF296) | NA |
| Niben101Scf10918g00006.1 | 15 | 30 | 1.131 | 0.028831103 | ref\|XP_002869579.1\| | rab GTPase activator [Arabidopsis lyrata subsp. lyrata] gb\|EFH45838.1\| rab GTPase activator [Arabidopsis lyrata subsp. lyrata] | IPR000195 (Rab-GTPase-TBC domain) | GO:0005097 (Rab GTPase activator activity), GO:0032313 (regulation of Rab GTPase activity) |
| Niben101Scf17514g00011.1 | 22 | 43 | 1.13 | 0.0195181 | ref\|XP_002529976.1\| | conserved hypothetical protein [Ricinus communis] gb\|EEF32419.1\| conserved hypothetical protein [Ricinus communis] | NA | NA |
| Niben101Scf02381g04022.1 | 2890 | 5685 | 1.129 | 1.2599E-07 | sp\|P26573\|RBS8_NICPL | Ribulose bisphosphate carboxylase small chain 8B, chloroplastic | IPR000894 (Ribulose bisphosphate carboxylase small chain, domain), IPR024680 (Ribulose-1,5-bisphosphate carboxylase small subunit, N-terminal), IPR024681 (Ribulose bisphosphate carboxylase, small chain) | NA |
| Niben101Scf08334g03002.1 | 12 | 24 | 1.125 | 0.040469735 | AT1G78560.1 | Sodium Bile acid symporter family LENGTH=401 | IPR002657 (Bile acid:sodium symporter/arsenical resistance protein Acr3) | GO:0016020 (membrane) |
| Niben101Scf13167g02003.1 | 42 | 81 | 1.118 | 0.036489553 | sp\|F4KAB8\|MCM6_ARATH | DNA replication licensing factor MCM6 | IPR001208 (Mini-chromosome maintenance, DNA-dependent ATPase), IPR004039 (Rubredoxin-type fold), IPR027417 (P-loop containing nucleoside triphosphate hydrolase), IPR027925 (MCM N-terminal domain) | GO:0003677 (DNA binding), GO:0003678 (DNA helicase activity), GO:0005524 (ATP binding), GO:0005634 (nucleus), GO:0006260 (DNA replication), GO:0006270 (DNA replication initiation), GO:0042555 (MCM complex) |
| Niben101Scf03720g00008.1 | 24 | 48 | 1.113 | 0.010069307 | sp\|Q8VHC5\|CABP4_MOUSE | Calcium-binding protein 4 | IPR011992 (EF-hand domain pair) | GO:0005509 (calcium ion binding) |
| Niben101Scf09387g03016.1 | 29 | 56 | 1.107 | 0.001667663 | sp\|Q11HP9\|EFG_CHESB | Elongation factor G | IPR004540 (Translation elongation factor EFG/EF2), IPR005225 (Small GTP-binding protein domain), IPR027417 (P-loop containing nucleoside triphosphate hydrolase) | GO:0003746 (translation elongation factor activity), GO:0003924 (GTPase activity), GO:0005525 (GTP binding), GO:0005622 (intracellular), GO:0006414 (translational elongation) |
| Niben101Scf06919g00029.1 | 29 | 56 | 1.104 | 0.03106355 | AT1G05990.1 | EF hand calcium-binding protein family LENGTH=150 | IPR011992 (EF-hand domain pair) | GO:0005509 (calcium ion binding) |
| Niben101Scf02638g02016.1 | 17 | 34 | 1.102 | 0.006540027 | sp\|Q885L1\|PHS_PSESM | Pterin-4-alpha-carbinolamine dehydratase | IPR001533 (Transcriptional coactivator/pterin dehydratase) | GO:0006729 (tetrahydrobiopterin biosynthetic process), GO:0008124 (4-alpha-hydroxytetrahydrobiopterin dehydratase activity) |
| Niben101Scf00453g07011.1 | 51 | 98 | 1.097 | 2.63798E-05 | AT5G46800.1 | Mitochondrial substrate carrier family protein LENGTH=300 | IPR018108 (Mitochondrial substrate/solute carrier), IPR023395 (Mitochondrial carrier domain) | NA |
| Niben101Scf04785g01004.1 | 28 | 53 | 1.096 | 0.001953166 | sp\|Q4V8X0\|TPRA1_DANRE | Transmembrane protein adipocyte-associated 1 homolog | IPR018781 (Transmembrane protein adipocyte-associated 1) | NA |
| Niben101Scf07187g01002.1 | 86 | 163 | 1.08 | 4.63476E-07 | AT5G14760.1 | L-aspartate oxidase LENGTH=651 | IPR003953 (FAD binding domain), IPR015939 (Fumarate reductase/succinate dehydrogenase flavoprotein-like, C-terminal), IPR027477 (Succinate dehydrogenase/fumarate reductase flavoprotein, catalytic domain) | GO:0016491 (oxidoreductase activity), GO:0055114 (oxidation-reduction process) |
| Niben101Scf04627g03001.1 | 19 | 37 | 1.078 | 0.013755176 | sp\|Q9FJX6\|FH6_ARATH | Formin-like protein 6 | IPR015425 (Formin, FH2 domain) | NA |
| Niben101Scf02240g03015.1 | 23 | 45 | 1.075 | 0.009657668 | sp\|Q9C8G9\|AB1C_ARATH | ABC transporter C family member 1 | IPR003439 (ABC transporter-like), IPR011527 (ABC transporter type 1, transmembrane domain), IPR027417 (P-loop containing nucleoside triphosphate hydrolase) | GO:0005524 (ATP binding), GO:0006810 (transport), GO:0016021 (integral component of membrane), GO:0016887 (ATPase activity), GO:0042626 (ATPase activity, coupled to transmembrane movement of substances), GO:0055085 (transmembrane transport) |
| Niben101Scf00960g09003.1 | 32 | 60 | 1.074 | 0.002574195 | sp\|O49884\|RL30_LUPLU | 60S ribosomal protein L30 | IPR000231 (Ribosomal protein L30e), IPR022991 (Ribosomal protein L30e, conserved site), IPR029064 (50S ribosomal protein L30e-like) | GO:0003735 (structural constituent of ribosome), GO:0005622 (intracellular), GO:0005840 (ribosome), GO:0006412 (translation) |
| Niben101Ctg16306g00002.1 | 26 | 50 | 1.074 | 0.010522977 | sp\|Q7XSK0\|BGL18_ORYSJ | Beta-glucosidase 18 | IPR001360 (Glycoside hydrolase, family 1), IPR017853 (Glycoside hydrolase superfamily) | GO:0004553 (hydrolase activity, hydrolyzing O-glycosyl compounds), GO:0005975 (carbohydrate metabolic process) |
| Niben101Scf03283g04012.1 | 107 | 201 | 1.066 | 7.31124E-05 | AT5G46800.1 | Mitochondrial substrate carrier family protein LENGTH=300 | IPR018108 (Mitochondrial substrate/solute carrier), IPR023395 (Mitochondrial carrier domain) | NA |
| Niben101Scf08002g01002.1 | 23 | 42 | 1.061 | 0.030158954 | AT1G51700.1 | DOF zinc finger protein 1 LENGTH=194 | IPR003851 (Zinc finger, Dof-type) | GO:0003677 (DNA binding), GO:0006355 (regulation of transcription, DNA-templated) |
| Niben101Scf01326g10014.1 | 15 | 28 | 1.06 | 0.023898007 | sp\|A7Z050\|VIP1_BOVIN | Inositol hexakisphosphate and diphosphoinositol-pentakisphosphate kinase 1 | IPR029033 (Histidine phosphatase superfamily) | GO:0003993 (acid phosphatase activity) |
| Niben101Ctg09403g00001.1 | 47 | 90 | 1.055 | 0.000812998 | NA | Unknown protein | NA | NA |
| Niben101Scf00652g01007.1 | 38 | 70 | 1.055 | 0.020716644 | sp\|P32518\|DUT_SOLLC | Deoxyuridine 5'-triphosphate nucleotidohydrolase | IPR008180 (Deoxyuridine triphosphate nucleotidohydrolase/Deoxycytidine triphosphate deaminase), IPR029054 (dUTPase-like) | GO:0016787 (hydrolase activity), GO:0046080 (dUTP metabolic process) |
| Niben101Scf05412g02002.1 | 19 | 35 | 1.054 | 0.043321513 | sp\|P05618\|CDGT_BACS0 | Cyclomaltodextrin glucanotransferase | IPR013783 (Immunoglobulin-like fold), IPR013784 (Carbohydrate-binding-like fold), IPR015902 (Glycoside hydrolase, family 13) | GO:0003824 (catalytic activity), GO:0030246 (carbohydrate binding), GO:2001070 (starch binding) |
| Niben101Scf04316g00001.1 | 24 | 45 | 1.052 | 0.004746844 | sp\|Q9FIT9\|HS217_ARATH | 21.7 kDa class VI heat shock protein | IPR008978 (HSP20-like chaperone) | NA |
| Niben101Scf01695g02013.1 | 19 | 36 | 1.046 | 0.018801441 | sp\|A4SVA6\|RPPH_POLSQ | RNA pyrophosphohydrolase | IPR003293 (Nudix hydrolase 6-like) | GO:0016787 (hydrolase activity) |
| Niben101Scf03710g06002.1 | 17 | 31 | 1.044 | 0.036140166 | AT1G73190.1 | Aquaporin-like superfamily protein LENGTH=268 | IPR000425 (Major intrinsic protein), IPR023271 (Aquaporin-like) | GO:0005215 (transporter activity), GO:0006810 (transport), GO:0016020 (membrane) |
| Niben101Scf03050g06003.1 | 20 | 38 | 1.043 | 0.042077774 | sp\|Q67YU0\|CKX5_ARATH | Cytokinin dehydrogenase 5 | IPR016164 (FAD-linked oxidase-like, C-terminal), IPR016166 (FAD-binding, type 2), IPR016170 (Vanillyl-alcohol oxidase/Cytokinin dehydrogenase C-terminal domain) | GO:0003824 (catalytic activity), GO:0008762 (UDP-N-acetylmuramate dehydrogenase activity), GO:0009690 (cytokinin metabolic process), GO:0016491 (oxidoreductase activity), GO:0016614 (oxidoreductase activity, acting on CH-OH group of donors), GO:0019139 (cytokinin dehydrogenase activity), GO:0050660 (flavin adenine dinucleotide binding), GO:0055114 (oxidation-reduction process) |
| Niben101Ctg16115g00003.1 | 23 | 42 | 1.04 | 0.0088654 | sp\|Q9C5T4\|WRK18_ARATH | WRKY transcription factor 18 | IPR003657 (DNA-binding WRKY) | GO:0003700 (sequence-specific DNA binding transcription factor activity), GO:0006355 (regulation of transcription, DNA-templated), GO:0043565 (sequence-specific DNA binding) |
| Niben101Scf01409g12028.1 | 27 | 50 | 1.03 | 0.043843734 | sp\|Q49YV3\|PUR8_STAS1 | Adenylosuccinate lyase | IPR000362 (Fumarate lyase family), IPR008948 (L-Aspartase-like), IPR024083 (Fumarase/histidase, N-terminal) | GO:0003824 (catalytic activity), GO:0004018 (N6-(1,2-dicarboxyethyl)AMP AMP-lyase (fumarate-forming) activity), GO:0006188 (IMP biosynthetic process), GO:0009152 (purine ribonucleotide biosynthetic process) |
| Niben101Scf12205g01017.1 | 53 | 96 | 1.024 | 0.001535759 | sp\|Q9C7N4\|GDL15_ARATH | GDSL esterase/lipase | IPR013830 (SGNH hydrolase-type esterase domain) | GO:0006629 (lipid metabolic process), GO:0016788 (hydrolase activity, acting on ester bonds) |
| Niben101Scf00911g02014.1 | 18 | 33 | 1.019 | 0.025990042 | AT1G06820.1 | carotenoid isomerase LENGTH=595 | IPR014101 (Carotene isomerase) | GO:0016117 (carotenoid biosynthetic process), GO:0016853 (isomerase activity) |
| Niben101Scf02451g01004.1 | 97 | 180 | 1.016 | 0.00490263 | sp\|B6EPQ0\|BIOH_ALISL | Pimeloyl-[acyl-carrier protein] methyl ester esterase | IPR029058 (Alpha/Beta hydrolase fold) | NA |
| Niben101Scf00980g01002.1 | 54 | 99 | 1.016 | 0.000283733 | AT2G41250.1 | Haloacid dehalogenase-like hydrolase (HAD) superfamily protein LENGTH=290 | IPR006439 (HAD hydrolase, subfamily IA), IPR023214 (HAD-like domain) | GO:0008152 (metabolic process), GO:0016787 (hydrolase activity) |
| Niben101Scf12858g01026.1 | 68 | 123 | 1.015 | 7.1038E-05 | sp\|O49292\|PPD4_ARATH | PsbP domain-containing protein 4, chloroplastic | IPR002683 (Photosystem II PsbP, oxygen evolving complex) | GO:0005509 (calcium ion binding), GO:0009523 (photosystem II), GO:0009654 (photosystem II oxygen evolving complex), GO:0015979 (photosynthesis), GO:0019898 (extrinsic component of membrane) |
| Niben101Scf05618g03008.1 | 28 | 50 | 1.011 | 0.016007349 | sp\|Q94KT8\|COBRA_ARATH | Protein COBRA | IPR006918 (COBRA, plant) | GO:0010215 (cellulose microfibril organization), GO:0016049 (cell growth), GO:0031225 (anchored component of membrane) |
| Niben101Scf07608g02015.1 | 15 | 27 | 1.008 | 0.028591545 | sp\|Q24849\|MCM3_ENTHI | DNA replication licensing factor MCM3 | IPR001208 (Mini-chromosome maintenance, DNA-dependent ATPase), IPR005637 (TAP C-terminal (TAP-C) domain), IPR027417 (P-loop containing nucleoside triphosphate hydrolase) | GO:0003677 (DNA binding), GO:0005524 (ATP binding), GO:0005634 (nucleus), GO:0006260 (DNA replication), GO:0051028 (mRNA transport) |
| Niben101Scf03121g04010.1 | 57 | 103 | 1.004 | 0.005837906 | ref\|WP_026731445.1\| | GCN5 family acetyltransferase [Fischerella sp. PCC 9605] | IPR016181 (Acyl-CoA N-acyltransferase) | GO:0008080 (N-acetyltransferase activity) |
| Niben101Scf01105g04004.1 | 27 | 49 | 1.004 | 0.006341681 | sp\|Q10184\|HIS4_SCHPO | 1-(5-phosphoribosyl)-5-[(5-phosphoribosylamino)methylideneamino] imidazole-4-carboxamide isomerase | IPR006062 (Histidine biosynthesis), IPR011858 (Phosphoribosylformimino-5-aminoimidazole carboxamide ribotide isomerase, eukaryotic), IPR013785 (Aldolase-type TIM barrel) | GO:0000105 (histidine biosynthetic process), GO:0003824 (catalytic activity), GO:0003949 (1-(5-phosphoribosyl)-5-[(5-phosphoribosylamino)methylideneamino]imidazole-4-carboxamide isomerase activity), GO:0008152 (metabolic process) |
| Niben101Scf04198g07011.1 | 21 | 39 | 1.001 | 0.035010179 | ref\|WP_031065677.1\| | phosphoesterase [Streptomyces sp. NRRL F-5527] | IPR016195 (Polymerase/histidinol phosphatase-like) | GO:0003824 (catalytic activity) |
| Niben101Scf05371g06002.1 | 37 | 68 | 1 | 0.009512105 | ref\|XP_002529976.1\| | conserved hypothetical protein [Ricinus communis] gb\|EEF32419.1\| conserved hypothetical protein [Ricinus communis] | NA | NA |
| Niben101Scf00062g01025.1 | 24 | 44 | 0.994 | 0.021250116 | sp\|B9DUB2\|GUAA_STRU0 | GMP synthase [glutamine-hydrolyzing] | IPR029062 (Class I glutamine amidotransferase-like) | NA |
| Niben101Scf02468g01008.1 | 42 | 75 | 0.987 | 0.011201468 | NA | Unknown protein | NA | NA |
| Niben101Scf10015g05003.1 | 82 | 145 | 0.981 | 0.000812181 | sp\|Q07761\|RL23A_TOBAC | 60S ribosomal protein L23a | IPR005633 (Ribosomal protein L23/L25, N-terminal), IPR013025 (Ribosomal protein L25/L23) | GO:0000166 (nucleotide binding), GO:0003735 (structural constituent of ribosome), GO:0005622 (intracellular), GO:0005840 (ribosome), GO:0006412 (translation) |
| Niben101Scf05928g00005.1 | 33 | 59 | 0.981 | 0.002834005 | gb\|EMS56326.1\| | LON peptidase N-terminal domain and RING finger protein 1 [Triticum urartu] | IPR003111 (Peptidase S16, lon N-terminal), IPR011990 (Tetratricopeptide-like helical domain), IPR013083 (Zinc finger, RING/FYVE/PHD-type), IPR015947 (PUA-like domain) | GO:0004176 (ATP-dependent peptidase activity), GO:0005515 (protein binding), GO:0006508 (proteolysis), GO:0008270 (zinc ion binding) |
| Niben101Scf01822g16013.1 | 16 | 29 | 0.98 | 0.030098874 | sp\|Q2YDC9\|PDCD2_BOVIN | Programmed cell death protein 2 | IPR002893 (Zinc finger, MYND-type), IPR007320 (Programmed cell death protein 2, C-terminal) | GO:0005737 (cytoplasm) |
| Niben101Scf05250g02011.1 | 67 | 118 | 0.975 | 0.022615393 | sp\|B1Z033\|BETB_BURA4 | NAD/NADP-dependent betaine aldehyde dehydrogenase | IPR016161 (Aldehyde/histidinol dehydrogenase), IPR029510 (Aldehyde dehydrogenase, glutamic acid active site) | GO:0008152 (metabolic process), GO:0016491 (oxidoreductase activity), GO:0016620 (oxidoreductase activity, acting on the aldehyde or oxo group of donors, NAD or NADP as acceptor), GO:0055114 (oxidation-reduction process) |
| Niben101Scf09597g01002.1 | 24 | 42 | 0.974 | 0.026078176 | sp\|P42813\|RNS1_ARATH | Ribonuclease 1 | IPR001568 (Ribonuclease T2-like) | GO:0003723 (RNA binding), GO:0033897 (ribonuclease T2 activity) |
| Niben101Scf04329g00013.1 | 19 | 34 | 0.974 | 0.035214705 | AT1G59453.1 | B-block binding subunit of TFIIIC LENGTH=1729 | IPR007309 (B-block binding subunit of TFIIIC), IPR011991 (Winged helix-turn-helix DNA-binding domain) | NA |
| Niben101Scf04505g04002.1 | 295 | 526 | 0.97 | 0.000926715 | ref\|XP_002513164.1\| | conserved hypothetical protein [Ricinus communis] gb\|EEF49155.1\| conserved hypothetical protein [Ricinus communis] | NA | NA |
| Niben101Scf02854g11002.1 | 128 | 225 | 0.968 | 0.008928298 | gb\|KEH28241.1\| | GASA/GAST/Snakin [Medicago truncatula] | NA | NA |
| Niben101Scf25265g00009.1 | 29 | 52 | 0.966 | 0.007273667 | sp\|Q54S83\|NDUF7_DICDI | NADH dehydrogenase [ubiquinone] complex I, assembly factor 7 | IPR003788 (Putative S-adenosyl-L-methionine-dependent methyltransferase MidA), IPR029063 (S-adenosyl-L-methionine-dependent methyltransferase) | NA |
| Niben101Scf06216g00003.1 | 28 | 50 | 0.965 | 0.020973451 | emb\|CDY47534.1\| | BnaC03g69280D [Brassica napus] | NA | NA |
| Niben101Scf06529g02013.1 | 25 | 43 | 0.964 | 0.01342984 | sp\|Q9SGW3\|PSD8A_ARATH | 26S proteasome non-ATPase regulatory subunit 8 homolog A | IPR005062 (SAC3/GANP/Nin1/mts3/eIF-3 p25) | GO:0005838 (proteasome regulatory particle), GO:0006508 (proteolysis) |
| Niben101Scf01785g04007.1 | 27 | 47 | 0.961 | 0.038039812 | AT3G57040.1 | response regulator 9 LENGTH=234 | IPR011006 (CheY-like superfamily) | GO:0000160 (phosphorelay signal transduction system) |
| Niben101Scf02155g01001.1 | 53 | 93 | 0.96 | 0.001633438 | sp\|Q9SW33\|TL1Y_ARATH | Thylakoid lumenal 17.9 kDa protein, chloroplastic | NA | NA |
| Niben101Scf12868g00008.1 | 46 | 80 | 0.957 | 0.00624718 | sp\|Q07439\|HSP71_RAT | Heat shock 70 kDa protein 1A/1B | IPR013126 (Heat shock protein 70 family), IPR029047 (Heat shock protein 70kD, peptide-binding domain), IPR029048 (Heat shock protein 70kD, C-terminal domain) | NA |
| Niben101Scf02990g06003.1 | 65 | 116 | 0.952 | 0.040460802 | AT4G37800.1 | xyloglucan endotransglucosylase/hydrolase 7 LENGTH=293 | IPR013320 (Concanavalin A-like lectin/glucanase domain), IPR016455 (Xyloglucan endotransglucosylase/hydrolase) | GO:0004553 (hydrolase activity, hydrolyzing O-glycosyl compounds), GO:0005618 (cell wall), GO:0005975 (carbohydrate metabolic process), GO:0006073 (cellular glucan metabolic process), GO:0016762 (xyloglucan:xyloglucosyl transferase activity), GO:0048046 (apoplast) |
| Niben101Scf04847g03018.1 | 154 | 269 | 0.95 | 0.001824318 | sp\|Q1CZI7\|HTPG_MYXXD | Chaperone protein HtpG | IPR001404 (Heat shock protein Hsp90 family) | GO:0005524 (ATP binding), GO:0006457 (protein folding), GO:0006950 (response to stress), GO:0051082 (unfolded protein binding) |
| Niben101Scf11790g01009.1 | 30 | 52 | 0.949 | 0.029656293 | sp\|Q66HG1\|DMTF1_RAT | Cyclin-D-binding Myb-like transcription factor 1 | IPR009057 (Homeodomain-like) | GO:0003677 (DNA binding), GO:0003682 (chromatin binding) |
| Niben101Scf09590g03004.1 | 24 | 42 | 0.944 | 0.044693061 | gb\|ADV29615.1\| | At5g37260-like protein [Solanum arcanum] | IPR009057 (Homeodomain-like) | GO:0003677 (DNA binding), GO:0003682 (chromatin binding) |
| Niben101Scf00173g00010.1 | 79 | 136 | 0.942 | 0.011726394 | sp\|A4IF94\|HIAL1_BOVIN | Hippocampus abundant transcript-like protein 1 | IPR011701 (Major facilitator superfamily), IPR020846 (Major facilitator superfamily domain) | GO:0016021 (integral component of membrane), GO:0055085 (transmembrane transport) |
| Niben101Scf03978g04007.1 | 18 | 31 | 0.942 | 0.032902691 | sp\|F4IAT2\|THOC2_ARATH | THO complex subunit 2 | IPR008896 (Uncharacterised protein family Ycf1), IPR018781 (Transmembrane protein adipocyte-associated 1), IPR021418 (THO complex, subunitTHOC2, C-terminal), IPR021726 (THO complex, subunitTHOC2, N-terminal) | NA |
| Niben101Scf00282g02013.1 | 53 | 91 | 0.932 | 0.0002778 | AT5G62140.1 | unknown protein; FUNCTIONS IN: molecular_function unknown; INVOLVED IN: biological_process unknown; LOCATED IN: chloroplast; EXPRESSED IN: 19 plant structures; EXPRESSED DURING: 13 growth stages; Has 60 Blast hits to 60 proteins in 24 species: Archae - 0; Bacteria - 14; Metazoa - 0; Fungi - 0; Plants - 45; Viruses - 0; Other Eukaryotes - 1 (source: NCBI BLink). LENGTH=241 | NA | NA |
| Niben101Scf04847g03018.1 | 97 | 164 | 0.93 | 0.016245873 | sp\|Q1CZI7\|HTPG_MYXXD | Chaperone protein HtpG | IPR001404 (Heat shock protein Hsp90 family) | GO:0005524 (ATP binding), GO:0006457 (protein folding), GO:0006950 (response to stress), GO:0051082 (unfolded protein binding) |
| Niben101Scf09417g01008.1 | 31 | 53 | 0.922 | 0.018895921 | AT3G12120.1 | fatty acid desaturase 2 LENGTH=383 | IPR005804 (Fatty acid desaturase, type 1), IPR021863 (Protein of unknown function DUF3474) | GO:0006629 (lipid metabolic process), GO:0016717 (oxidoreductase activity, acting on paired donors, with oxidation of a pair of donors resulting in the reduction of molecular oxygen to two molecules of water), GO:0055114 (oxidation-reduction process) |
| Niben101Scf20614g00017.1 | 45 | 77 | 0.921 | 0.003170517 | sp\|Q9SYM4\|TPS1_ARATH | Alpha,alpha-trehalose-phosphate synthase [UDP-forming] 1 | IPR001830 (Glycosyl transferase, family 20), IPR003337 (Trehalose-phosphatase), IPR023214 (HAD-like domain) | GO:0003824 (catalytic activity), GO:0003825 (alpha,alpha-trehalose-phosphate synthase (UDP-forming) activity), GO:0005992 (trehalose biosynthetic process) |
| Niben101Scf08002g01012.1 | 537 | 909 | 0.918 | 0.002683524 | sp\|B7LPR5\|YQGE_ESCF3 | UPF0301 protein YqgE | IPR003774 (Protein of unknown function UPF0301) | NA |
| Niben101Scf14098g00007.1 | 566 | 970 | 0.917 | 0.000347589 | AT2G10940.1 | Bifunctional inhibitor/lipid-transfer protein/seed storage 2S albumin superfamily protein LENGTH=291 | IPR016140 (Bifunctional inhibitor/plant lipid transfer protein/seed storage helical domain) | NA |
| Niben101Scf06928g00008.1 | 1452 | 2467 | 0.916 | 0.002715009 | AT2G37180.1 | Aquaporin-like superfamily protein LENGTH=285 | IPR000425 (Major intrinsic protein), IPR023271 (Aquaporin-like) | GO:0005215 (transporter activity), GO:0006810 (transport), GO:0016020 (membrane) |
| Niben101Scf00340g03006.1 | 66 | 112 | 0.912 | 0.00035957 | sp\|P73016\|FABI_SYNY3 | Enoyl-[acyl-carrier-protein] reductase [NADH] FabI | IPR002347 (Glucose/ribitol dehydrogenase) | NA |
| Niben101Scf03706g02006.1 | 37 | 62 | 0.904 | 0.017057487 | sp\|P83948\|PME3_CITSI | Pectinesterase 3 | IPR006501 (Pectinesterase inhibitor domain), IPR011050 (Pectin lyase fold/virulence factor) | GO:0004857 (enzyme inhibitor activity), GO:0005618 (cell wall), GO:0030599 (pectinesterase activity), GO:0042545 (cell wall modification) |
| Niben101Scf08162g00028.1 | 60 | 100 | 0.901 | 0.00602876 | AT5G66550.1 | Maf-like protein LENGTH=207 | IPR029001 (Inosine triphosphate pyrophosphatase-like) | GO:0005737 (cytoplasm) |
| Niben101Scf08191g02001.1 | 50 | 85 | 0.901 | 0.000559185 | sp\|P16097\|CAPP2_MESCR | Phosphoenolpyruvate carboxylase 2 | IPR021135 (Phosphoenolpyruvate carboxylase) | GO:0003824 (catalytic activity), GO:0006099 (tricarboxylic acid cycle), GO:0008964 (phosphoenolpyruvate carboxylase activity), GO:0015977 (carbon fixation) |
| Niben101Scf00466g00012.1 | 3983 | 6616 | 0.899 | 6.76297E-08 | AT2G21330.1 | fructose-bisphosphate aldolase 1 LENGTH=399 | IPR000741 (Fructose-bisphosphate aldolase, class-I), IPR013785 (Aldolase-type TIM barrel), IPR029768 (Fructose-bisphosphate aldolase class-I active site) | GO:0003824 (catalytic activity), GO:0004332 (fructose-bisphosphate aldolase activity), GO:0006096 (glycolytic process) |
| Niben101Scf01836g00002.1 | 319 | 541 | 0.899 | 0.005837906 | sp\|P24806\|XTH24_ARATH | Xyloglucan endotransglucosylase/hydrolase protein 24 | IPR013320 (Concanavalin A-like lectin/glucanase domain), IPR016455 (Xyloglucan endotransglucosylase/hydrolase) | GO:0004553 (hydrolase activity, hydrolyzing O-glycosyl compounds), GO:0005618 (cell wall), GO:0005975 (carbohydrate metabolic process), GO:0006073 (cellular glucan metabolic process), GO:0016762 (xyloglucan:xyloglucosyl transferase activity), GO:0048046 (apoplast) |
| Niben101Scf05351g00032.1 | 64 | 106 | 0.898 | 0.000207383 | AT1G07420.1 | sterol 4-alpha-methyl-oxidase 2-1 LENGTH=266 | IPR006694 (Fatty acid hydroxylase) | GO:0005506 (iron ion binding), GO:0006633 (fatty acid biosynthetic process), GO:0016491 (oxidoreductase activity), GO:0055114 (oxidation-reduction process) |
| Niben101Scf09417g01013.1 | 54 | 91 | 0.894 | 0.010334569 | gb\|AAA34063.1\| | cysteine-rich extensin-like protein [Nicotiana tabacum] | NA | NA |
| Niben101Scf01653g02013.1 | 7045 | 11670 | 0.893 | 2.58853E-10 | sp\|Q06721\|RCA_ANASC | Ribulose bisphosphate carboxylase/oxygenase activase | IPR003959 (ATPase, AAA-type, core), IPR027417 (P-loop containing nucleoside triphosphate hydrolase) | GO:0005524 (ATP binding) |
| Niben101Scf02581g00006.1 | 84 | 141 | 0.893 | 0.005369887 | AT5G51010.1 | Rubredoxin-like superfamily protein LENGTH=154 | IPR004039 (Rubredoxin-type fold) | GO:0005506 (iron ion binding) |
| Niben101Scf02107g04006.1 | 26 | 44 | 0.893 | 0.027392368 | sp\|Q6DN52\|CBPL_DICDI | Calcium-binding protein L | IPR011992 (EF-hand domain pair) | GO:0005509 (calcium ion binding) |
| Niben101Scf01653g02013.1 | 199 | 331 | 0.892 | 2.4593E-05 | sp\|Q06721\|RCA_ANASC | Ribulose bisphosphate carboxylase/oxygenase activase | IPR003959 (ATPase, AAA-type, core), IPR027417 (P-loop containing nucleoside triphosphate hydrolase) | GO:0005524 (ATP binding) |
| Niben101Scf07437g01012.1 | 101 | 168 | 0.892 | 0.000113271 | AT1G70000.1 | myb-like transcription factor family protein LENGTH=261 | IPR009057 (Homeodomain-like) | GO:0003677 (DNA binding), GO:0003682 (chromatin binding) |
| Niben101Scf14859g00005.1 | 1363 | 2276 | 0.888 | 5.89331E-05 | ref\|WP_014243176.1\| | 2-oxoglutarate translocator [Bacteriovorax marinus] ref\|YP_005034360.1\| putative membrane transport protein [Bacteriovorax marinus SJ] emb\|CBW25389.1\| putative membrane transport protein [Bacteriovorax marinus SJ] | IPR001898 (Sodium/sulphate symporter) | GO:0005215 (transporter activity), GO:0006814 (sodium ion transport), GO:0016020 (membrane), GO:0055085 (transmembrane transport) |
| Niben101Scf11251g00008.1 | 60 | 101 | 0.888 | 0.020004803 | sp\|Q6NMQ7\|GASA6_ARATH | Gibberellin-regulated protein 6 | IPR003854 (Gibberellin regulated protein) | NA |
| Niben101Scf00550g00014.1 | 60 | 100 | 0.884 | 0.010249262 | sp\|O22126\|FLA8_ARATH | Fasciclin-like arabinogalactan protein 8 | IPR000782 (FAS1 domain) | NA |
| Niben101Scf13710g03002.1 | 33 | 54 | 0.884 | 0.031473618 | sp\|P48368\|CRTE_CYAPA | Geranylgeranyl diphosphate synthase | IPR017446 (Polyprenyl synthetase-related) | GO:0008299 (isoprenoid biosynthetic process) |
| Niben101Scf03341g05001.1 | 97 | 161 | 0.882 | 0.028067933 | sp\|P11898\|GRP1_OXYRB | Glycine-rich protein HC1 | IPR010800 (Glycine rich protein) | NA |
| Niben101Scf02752g11002.1 | 111 | 186 | 0.881 | 0.001028139 | sp\|B6T959\|CASP1_MAIZE | Casparian strip membrane protein 1 | IPR006702 (Uncharacterised protein family UPF0497, trans-membrane plant) | NA |
| Niben101Scf01036g03021.1 | 22 | 37 | 0.878 | 0.028094655 | sp\|P22682\|CBL_MOUSE | E3 ubiquitin-protein ligase CBL | IPR013083 (Zinc finger, RING/FYVE/PHD-type) | GO:0005515 (protein binding), GO:0008270 (zinc ion binding) |
| Niben101Scf01175g05011.1 | 51 | 84 | 0.876 | 0.016245873 | sp\|Q3SWX8\|RBBP7_BOVIN | Histone-binding protein RBBP7 | IPR015943 (WD40/YVTN repeat-like-containing domain) | GO:0005515 (protein binding) |
| Niben101Scf00919g03002.1 | 28 | 46 | 0.875 | 0.046567631 | AT3G07700.3 | Protein kinase superfamily protein LENGTH=724 | IPR011009 (Protein kinase-like domain) | GO:0016772 (transferase activity, transferring phosphorus-containing groups) |
| Niben101Scf01283g02002.1 | 35 | 58 | 0.874 | 0.047999884 | AT4G14210.1 | phytoene desaturase 3 LENGTH=566 | IPR014102 (Phytoene desaturase), IPR016040 (NAD(P)-binding domain) | GO:0016117 (carotenoid biosynthetic process), GO:0016491 (oxidoreductase activity), GO:0016705 (oxidoreductase activity, acting on paired donors, with incorporation or reduction of molecular oxygen), GO:0055114 (oxidation-reduction process) |
| Niben101Scf11341g00021.1 | 74 | 120 | 0.865 | 0.000993689 | AT5G23060.1 | calcium sensing receptor LENGTH=387 | IPR001763 (Rhodanese-like domain) | NA |
| Niben101Scf01870g03002.1 | 259 | 427 | 0.86 | 0.00523779 | sp\|O04722\|SUT21_ARATH | Sulfate transporter 2.1 | IPR001902 (Sulphate anion transporter), IPR030402 (Sulfate transporter N-terminal domain with GLY motif) | GO:0008271 (secondary active sulfate transmembrane transporter activity), GO:0008272 (sulfate transport), GO:0015116 (sulfate transmembrane transporter activity), GO:0016020 (membrane), GO:0016021 (integral component of membrane), GO:0055085 (transmembrane transport) |
| Niben101Scf02886g02004.1 | 76 | 123 | 0.859 | 0.018363397 | sp\|O48818\|EXPA4_ARATH | Expansin-A4 | IPR007118 (Expansin/Lol pI) | GO:0005576 (extracellular region), GO:0009664 (plant-type cell wall organization) |
| Niben101Scf00897g05001.1 | 79 | 130 | 0.857 | 0.001535759 | sp\|P94111\|STS1_ARATH | Strictosidine synthase 1 | IPR004141 (Strictosidine synthase) | GO:0009058 (biosynthetic process), GO:0016844 (strictosidine synthase activity) |
| Niben101Scf01848g02002.1 | 32 | 52 | 0.85 | 0.032324726 | AT2G48070.3 | resistance to phytophthora 1 LENGTH=201 | NA | NA |
| Niben101Scf04556g04012.1 | 92 | 146 | 0.847 | 0.006949852 | sp\|Q9ZQI8\|NLTL2_ARATH | Non-specific lipid-transfer protein-like protein | IPR000528 (Plant lipid transfer protein/Par allergen), IPR016140 (Bifunctional inhibitor/plant lipid transfer protein/seed storage helical domain) | GO:0006869 (lipid transport), GO:0008289 (lipid binding) |
| Niben101Scf02665g02009.1 | 80 | 132 | 0.847 | 0.049943648 | sp\|P48620\|FAD3C_SESIN | Omega-3 fatty acid desaturase, chloroplastic | IPR005804 (Fatty acid desaturase, type 1), IPR021863 (Protein of unknown function DUF3474) | GO:0006629 (lipid metabolic process), GO:0016717 (oxidoreductase activity, acting on paired donors, with oxidation of a pair of donors resulting in the reduction of molecular oxygen to two molecules of water), GO:0055114 (oxidation-reduction process) |
| Niben101Scf04011g08004.1 | 43 | 70 | 0.847 | 0.003925001 | ref\|XP_005651523.1\| | iron-sulfur cluster assembly protein [Coccomyxa subellipsoidea C-169] gb\|EIE26979.1\| iron-sulfur cluster assembly protein [Coccomyxa subellipsoidea C-169] | IPR001075 (NIF system FeS cluster assembly, NifU, C-terminal) | GO:0005506 (iron ion binding), GO:0016226 (iron-sulfur cluster assembly), GO:0051536 (iron-sulfur cluster binding) |
| Niben101Scf05226g04004.1 | 758 | 1228 | 0.845 | 0.000171151 | AT1G42960.1 | expressed protein localized to the inner membrane of the chloroplast. LENGTH=168 | NA | NA |
| Niben101Scf04436g08005.1 | 101 | 162 | 0.838 | 0.00114196 | sp\|Q9LYK9\|RS263_ARATH | 40S ribosomal protein S26-3 | IPR000892 (Ribosomal protein S26e) | GO:0003735 (structural constituent of ribosome), GO:0005622 (intracellular), GO:0005840 (ribosome), GO:0006412 (translation) |
| Niben101Scf04741g02009.1 | 67 | 108 | 0.834 | 0.001526258 | AT1G53510.1 | mitogen-activated protein kinase 18 LENGTH=615 | IPR011009 (Protein kinase-like domain) | GO:0004672 (protein kinase activity), GO:0005524 (ATP binding), GO:0006468 (protein phosphorylation), GO:0016772 (transferase activity, transferring phosphorus-containing groups) |
| Niben101Scf02824g03017.1 | 62 | 99 | 0.827 | 0.036533653 | AT2G04030.1 | Chaperone protein htpG family protein LENGTH=780 | IPR001404 (Heat shock protein Hsp90 family) | GO:0005524 (ATP binding), GO:0006457 (protein folding), GO:0006950 (response to stress), GO:0051082 (unfolded protein binding) |
| Niben101Scf03320g03004.1 | 35 | 55 | 0.827 | 0.012716326 | sp\|Q6DW76\|DGDG1_SOYBN | Digalactosyldiacylglycerol synthase 1, chloroplastic | NA | NA |
| Niben101Scf02764g05013.1 | 353 | 560 | 0.826 | 1.38487E-05 | sp\|P48620\|FAD3C_SESIN | Omega-3 fatty acid desaturase, chloroplastic | IPR005804 (Fatty acid desaturase, type 1), IPR021863 (Protein of unknown function DUF3474) | GO:0006629 (lipid metabolic process), GO:0016717 (oxidoreductase activity, acting on paired donors, with oxidation of a pair of donors resulting in the reduction of molecular oxygen to two molecules of water), GO:0055114 (oxidation-reduction process) |
| Niben101Scf00709g07009.1 | 217 | 342 | 0.822 | 0.001328066 | AT5G67370.1 | Protein of unknown function (DUF1230) LENGTH=327 | IPR009631 (Uncharacterised protein family Ycf36) | NA |
| Niben101Scf01155g01009.1 | 169 | 268 | 0.82 | 0.000153522 | sp\|P46512\|CAH1_FLALI | Carbonic anhydrase 1 | IPR001765 (Carbonic anhydrase) | GO:0004089 (carbonate dehydratase activity), GO:0008270 (zinc ion binding), GO:0015976 (carbon utilization) |
| Niben101Scf10203g00008.1 | 105 | 168 | 0.819 | 0.001379066 | ref\|WP_015115534.1\| | phage shock protein A (PspA) family protein [N [Nostoc sp. PCC 7107] | IPR007157 (PspA/IM30) | NA |
| Niben101Scf04182g02003.1 | 61 | 97 | 0.818 | 0.001964715 | sp\|O48818\|EXPA4_ARATH | Expansin-A4 | IPR007118 (Expansin/Lol pI) | GO:0005576 (extracellular region), GO:0009664 (plant-type cell wall organization) |
| Niben101Scf01339g06010.1 | 32 | 51 | 0.817 | 0.028591545 | sp\|Q97BW2\|RS14Z_THEVO | 30S ribosomal protein S14 type Z | IPR001209 (Ribosomal protein S14) | GO:0003735 (structural constituent of ribosome), GO:0005622 (intracellular), GO:0005840 (ribosome), GO:0006412 (translation) |
| Niben101Scf02706g03008.1 | 27 | 43 | 0.816 | 0.023100173 | gb\|ADL36873.1\| | At1g62390-like protein [Arabidopsis lyrata subsp. lyrata] | IPR000270 (Phox/Bem1p), IPR011990 (Tetratricopeptide-like helical domain) | GO:0005515 (protein binding) |
| Niben101Scf30335g00004.1 | 114 | 181 | 0.814 | 0.002303751 | NA | Unknown protein | NA | NA |
| Niben101Scf09417g01013.1 | 142 | 225 | 0.811 | 0.001683204 | gb\|AAA34063.1\| | cysteine-rich extensin-like protein [Nicotiana tabacum] | NA | NA |
| Niben101Scf11380g03009.1 | 60 | 95 | 0.811 | 0.012579582 | AT1G02910.1 | tetratricopeptide repeat (TPR)-containing protein LENGTH=453 | IPR011990 (Tetratricopeptide-like helical domain), IPR021883 (Protein of unknown function DUF3493) | GO:0005515 (protein binding) |
| Niben101Scf02610g00005.1 | 47 | 74 | 0.809 | 0.002715009 | AT1G70750.1 | Protein of unknown function, DUF593 LENGTH=749 | IPR007656 (Zein-binding domain) | NA |
| Niben101Scf07309g00003.1 | 776 | 1231 | 0.808 | 0.006429258 | sp\|Q7V643\|CH601_PROMM | 60 kDa chaperonin 1 | IPR002423 (Chaperonin Cpn60/TCP-1), IPR027409 (GroEL-like apical domain), IPR027413 (GroEL-like equatorial domain) | GO:0005524 (ATP binding), GO:0005737 (cytoplasm), GO:0006457 (protein folding), GO:0042026 (protein refolding), GO:0044267 (cellular protein metabolic process) |
| Niben101Scf10796g00009.1 | 160 | 252 | 0.808 | 0.003749975 | sp\|Q1CZI7\|HTPG_MYXXD | Chaperone protein HtpG | IPR001404 (Heat shock protein Hsp90 family) | GO:0005524 (ATP binding), GO:0006457 (protein folding), GO:0006950 (response to stress), GO:0051082 (unfolded protein binding) |
| Niben101Scf00089g04004.1 | 45 | 70 | 0.808 | 0.03985301 | AT2G28510.1 | Dof-type zinc finger DNA-binding family protein LENGTH=288 | IPR003851 (Zinc finger, Dof-type) | GO:0003677 (DNA binding), GO:0006355 (regulation of transcription, DNA-templated) |
| Niben101Scf01823g00003.1 | 27 | 43 | 0.808 | 0.048205217 | sp\|A1TYH3\|ERPA_MARHV | Iron-sulfur cluster insertion protein ErpA | IPR000361 (FeS cluster biogenesis), IPR016092 (FeS cluster insertion protein) | GO:0005198 (structural molecule activity), GO:0016226 (iron-sulfur cluster assembly), GO:0051536 (iron-sulfur cluster binding) |
| Niben101Scf12205g01018.1 | 185 | 290 | 0.804 | 0.001054079 | sp\|Q9FK75\|GDL82_ARATH | GDSL esterase/lipase | IPR013830 (SGNH hydrolase-type esterase domain) | GO:0006629 (lipid metabolic process), GO:0016788 (hydrolase activity, acting on ester bonds) |
| Niben101Scf25430g00005.1 | 39 | 61 | 0.802 | 0.011454175 | sp\|Q54FE8\|RPA49_DICDI | DNA-directed RNA polymerase I subunit rpa49 | IPR009668 (RNA polymerase I associated factor, A49-like) | GO:0003677 (DNA binding), GO:0003899 (DNA-directed RNA polymerase activity), GO:0005634 (nucleus), GO:0006351 (transcription, DNA-templated) |
| Niben101Scf00683g02004.1 | 32 | 50 | 0.8 | 0.046205068 | AT5G55530.1 | Calcium-dependent lipid-binding (CaLB domain) family protein LENGTH=405 | IPR000008 (C2 domain) | GO:0005515 (protein binding) |
| Niben101Scf03208g05006.1 | 124 | 193 | 0.796 | 0.003033069 | NA | Unknown protein | NA | NA |
| Niben101Scf03978g07013.1 | 40 | 63 | 0.786 | 0.034729101 | AT5G17020.1 | exportin 1A LENGTH=1075 | IPR016024 (Armadillo-type fold) | GO:0005488 (binding), GO:0006886 (intracellular protein transport), GO:0008536 (Ran GTPase binding) |
| Niben101Scf07109g05001.1 | 28 | 44 | 0.786 | 0.040458236 | sp\|Q9LIG2\|RLK6_ARATH | Receptor-like protein kinase | IPR011009 (Protein kinase-like domain), IPR014729 (Rossmann-like alpha/beta/alpha sandwich fold) | GO:0004672 (protein kinase activity), GO:0004674 (protein serine/threonine kinase activity), GO:0005524 (ATP binding), GO:0006468 (protein phosphorylation), GO:0006950 (response to stress), GO:0016772 (transferase activity, transferring phosphorus-containing groups) |
| Niben101Scf16592g00013.1 | 120 | 188 | 0.784 | 0.031305686 | sp\|P49637\|R27A3_ARATH | 60S ribosomal protein L27a-3 | IPR021131 (Ribosomal protein L18e/L15P) | NA |
| Niben101Scf05912g02024.1 | 32 | 50 | 0.777 | 0.026644969 | sp\|Q93Y09\|SCP45_ARATH | Serine carboxypeptidase-like 45 | IPR001563 (Peptidase S10, serine carboxypeptidase), IPR029058 (Alpha/Beta hydrolase fold) | GO:0004185 (serine-type carboxypeptidase activity), GO:0006508 (proteolysis) |
| Niben101Scf01655g01015.1 | 160 | 243 | 0.773 | 0.003897746 | sp\|Q7M1E7\|PGLR2_CHAOB | Polygalacturonase | IPR000743 (Glycoside hydrolase, family 28), IPR011050 (Pectin lyase fold/virulence factor) | GO:0004650 (polygalacturonase activity), GO:0005975 (carbohydrate metabolic process) |
| Niben101Scf08195g07014.1 | 382 | 586 | 0.77 | 0.002511193 | sp\|Q9FK75\|GDL82_ARATH | GDSL esterase/lipase | IPR013830 (SGNH hydrolase-type esterase domain) | GO:0006629 (lipid metabolic process), GO:0016788 (hydrolase activity, acting on ester bonds) |
| Niben101Scf04779g01006.1 | 292 | 450 | 0.77 | 0.000761229 | sp\|Q2IJ93\|EFG1_ANADE | Elongation factor G 1 | IPR000795 (Elongation factor, GTP-binding domain), IPR005225 (Small GTP-binding protein domain), IPR009000 (Translation protein, beta-barrel domain), IPR027417 (P-loop containing nucleoside triphosphate hydrolase) | GO:0003924 (GTPase activity), GO:0005525 (GTP binding) |
| Niben101Scf02990g06003.1 | 2666 | 4164 | 0.769 | 0.009647661 | AT4G37800.1 | xyloglucan endotransglucosylase/hydrolase 7 LENGTH=293 | IPR013320 (Concanavalin A-like lectin/glucanase domain), IPR016455 (Xyloglucan endotransglucosylase/hydrolase) | GO:0004553 (hydrolase activity, hydrolyzing O-glycosyl compounds), GO:0005618 (cell wall), GO:0005975 (carbohydrate metabolic process), GO:0006073 (cellular glucan metabolic process), GO:0016762 (xyloglucan:xyloglucosyl transferase activity), GO:0048046 (apoplast) |
| Niben101Scf02842g00014.1 | 101 | 154 | 0.769 | 0.030098874 | sp\|Q98AX9\|CH603_RHILO | 60 kDa chaperonin 3 | IPR002423 (Chaperonin Cpn60/TCP-1), IPR027409 (GroEL-like apical domain), IPR027413 (GroEL-like equatorial domain) | GO:0005524 (ATP binding), GO:0005737 (cytoplasm), GO:0006457 (protein folding), GO:0042026 (protein refolding), GO:0044267 (cellular protein metabolic process) |
| Niben101Scf07234g00008.1 | 45 | 70 | 0.769 | 0.01340991 | AT4G34980.1 | subtilisin-like serine protease 2 LENGTH=764 | IPR015500 (Peptidase S8, subtilisin-related), IPR023828 (Peptidase S8, subtilisin, Ser-active site) | GO:0004252 (serine-type endopeptidase activity), GO:0006508 (proteolysis) |
| Niben101Scf00897g02011.1 | 50 | 76 | 0.767 | 0.037179774 | sp\|Q87G18\|LLDD_VIBPA | L-lactate dehydrogenase | IPR012133 (Alpha-hydroxy acid dehydrogenase, FMN-dependent), IPR013785 (Aldolase-type TIM barrel) | GO:0003824 (catalytic activity), GO:0010181 (FMN binding), GO:0016491 (oxidoreductase activity), GO:0055114 (oxidation-reduction process) |
| Niben101Scf11178g01004.1 | 117 | 178 | 0.764 | 0.022354459 | sp\|Q8ZSB0\|MNTH_NOSS1 | Divalent metal cation transporter MntH | IPR001046 (NRAMP family) | GO:0005215 (transporter activity), GO:0006810 (transport), GO:0016020 (membrane) |
| Niben101Scf04875g02008.1 | 68 | 103 | 0.764 | 0.01663746 | AT2G36780.1 | UDP-Glycosyltransferase superfamily protein LENGTH=496 | IPR002213 (UDP-glucuronosyl/UDP-glucosyltransferase) | GO:0008152 (metabolic process), GO:0016758 (transferase activity, transferring hexosyl groups) |
| Niben101Scf06216g01013.1 | 40 | 62 | 0.764 | 0.033893689 | sp\|Q8LEM8\|RL373_ARATH | 60S ribosomal protein L37-3 | IPR001569 (Ribosomal protein L37e), IPR011332 (Zinc-binding ribosomal protein) | GO:0003735 (structural constituent of ribosome), GO:0005622 (intracellular), GO:0005840 (ribosome), GO:0006412 (translation) |
| Niben101Scf06910g03007.1 | 40 | 62 | 0.762 | 0.044083678 | sp\|A6UUQ4\|GLMU_META3 | Bifunctional protein GlmU | IPR001451 (Bacterial transferase hexapeptide repeat), IPR029044 (Nucleotide-diphospho-sugar transferases) | GO:0009058 (biosynthetic process), GO:0016779 (nucleotidyltransferase activity) |
| Niben101Scf18639g00026.1 | 1737 | 2646 | 0.759 | 7.05673E-05 | AT5G23060.1 | calcium sensing receptor LENGTH=387 | IPR001763 (Rhodanese-like domain) | NA |
| Niben101Scf08457g00004.1 | 71 | 108 | 0.759 | 0.001407261 | sp\|Q3J8A0\|GPH_NITOC | Phosphoglycolate phosphatase | IPR006439 (HAD hydrolase, subfamily IA), IPR023214 (HAD-like domain) | GO:0008152 (metabolic process), GO:0016787 (hydrolase activity) |
| Niben101Scf15824g00019.1 | 28 | 42 | 0.755 | 0.045840285 | ref\|XP_005651523.1\| | iron-sulfur cluster assembly protein [Coccomyxa subellipsoidea C-169] gb\|EIE26979.1\| iron-sulfur cluster assembly protein [Coccomyxa subellipsoidea C-169] | IPR001075 (NIF system FeS cluster assembly, NifU, C-terminal) | GO:0005506 (iron ion binding), GO:0016226 (iron-sulfur cluster assembly), GO:0051536 (iron-sulfur cluster binding) |
| Niben101Scf08328g00011.1 | 70 | 105 | 0.754 | 0.013934966 | AT5G20140.2 | SOUL heme-binding family protein LENGTH=395 | IPR006917 (SOUL haem-binding protein), IPR011256 (Regulatory factor, effector binding domain), IPR018790 (Protein of unknown function DUF2358) | NA |
| Niben101Scf03414g01004.1 | 94 | 143 | 0.747 | 0.0042229 | sp\|P48621\|FAD3C_SOYBN | Omega-3 fatty acid desaturase, chloroplastic | IPR005804 (Fatty acid desaturase, type 1), IPR021863 (Protein of unknown function DUF3474) | GO:0006629 (lipid metabolic process), GO:0016717 (oxidoreductase activity, acting on paired donors, with oxidation of a pair of donors resulting in the reduction of molecular oxygen to two molecules of water), GO:0055114 (oxidation-reduction process) |
| Niben101Scf10556g00029.1 | 114 | 171 | 0.746 | 0.025662251 | emb\|CDX99019.1\| | BnaC09g48160D [Brassica napus] | NA | NA |
| Niben101Scf13379g00006.1 | 71 | 108 | 0.746 | 0.009714094 | sp\|P17198\|RS7_SULAC | 30S ribosomal protein S7 | IPR000235 (Ribosomal protein S5/S7), IPR023798 (Ribosomal protein S7 domain) | GO:0003723 (RNA binding), GO:0003735 (structural constituent of ribosome), GO:0006412 (translation) |
| Niben101Scf17024g00002.1 | 50 | 76 | 0.744 | 0.012579582 | AT1G02205.3 | Fatty acid hydroxylase superfamily LENGTH=630 | IPR006694 (Fatty acid hydroxylase), IPR021940 (Uncharacterised domain Wax2, C-terminal) | GO:0005506 (iron ion binding), GO:0006633 (fatty acid biosynthetic process), GO:0016491 (oxidoreductase activity), GO:0055114 (oxidation-reduction process) |
| Niben101Scf03133g00003.1 | 83 | 124 | 0.741 | 0.005053075 | sp\|Q5HF86\|G3P2_STAAC | Glyceraldehyde-3-phosphate dehydrogenase 2 | IPR003823 (Domain of unknown function CP12), IPR020831 (Glyceraldehyde/Erythrose phosphate dehydrogenase family) | GO:0016620 (oxidoreductase activity, acting on the aldehyde or oxo group of donors, NAD or NADP as acceptor), GO:0055114 (oxidation-reduction process) |
| Niben101Scf03185g01009.1 | 44 | 66 | 0.74 | 0.017426435 | sp\|B6EGU1\|FADA_ALISL | 3-ketoacyl-CoA thiolase | IPR002155 (Thiolase), IPR016039 (Thiolase-like) | GO:0003824 (catalytic activity), GO:0008152 (metabolic process), GO:0016747 (transferase activity, transferring acyl groups other than amino-acyl groups) |
| Niben101Scf00853g07006.1 | 145 | 218 | 0.739 | 0.032708416 | sp\|Q30Z53\|RL24_DESAG | 50S ribosomal protein L24 | IPR005756 (Ribosomal protein L26/L24P, eukaryotic/archaeal), IPR008991 (Translation protein SH3-like domain) | GO:0003735 (structural constituent of ribosome), GO:0005622 (intracellular), GO:0005840 (ribosome), GO:0006412 (translation), GO:0015934 (large ribosomal subunit) |
| Niben101Scf00013g03001.1 | 64 | 97 | 0.739 | 0.012579582 | sp\|B9L3S8\|FTSH2_THERP | ATP-dependent zinc metalloprotease FtsH 2 | IPR000642 (Peptidase M41), IPR003959 (ATPase, AAA-type, core), IPR027417 (P-loop containing nucleoside triphosphate hydrolase) | GO:0004222 (metalloendopeptidase activity), GO:0005524 (ATP binding), GO:0006508 (proteolysis) |
| Niben101Scf01046g03010.1 | 82 | 123 | 0.732 | 0.039634023 | sp\|P93332\|NOD3_MEDTR | Bidirectional sugar transporter N3 | IPR018179 (SWEET sugar transporter, plants) | GO:0016021 (integral component of membrane) |
| Niben101Scf01451g04016.1 | 84 | 125 | 0.731 | 0.008122433 | ref\|WP_009632881.1\| | Protein of unknown function (DUF2996) [Synechocystis sp. PCC 7509] | IPR021374 (Protein of unknown function DUF2996) | NA |
| Niben101Scf18348g01035.1 | 128 | 191 | 0.73 | 0.009353115 | AT4G01070.1 | UDP-Glycosyltransferase superfamily protein LENGTH=480 | IPR002213 (UDP-glucuronosyl/UDP-glucosyltransferase) | GO:0008152 (metabolic process), GO:0016758 (transferase activity, transferring hexosyl groups) |
| Niben101Scf05469g01024.1 | 42 | 62 | 0.73 | 0.029007938 | sp\|P25792\|CYSP_SCHMA | Cathepsin B-like cysteine proteinase | IPR013128 (Peptidase C1A), IPR025660 (Cysteine peptidase, histidine active site) | GO:0006508 (proteolysis), GO:0008234 (cysteine-type peptidase activity) |
| Niben101Scf03671g02003.1 | 354 | 525 | 0.727 | 0.002522179 | emb\|CDY27712.1\| | BnaA05g31210D [Brassica napus] | NA | NA |
| Niben101Scf01927g01013.1 | 46 | 68 | 0.727 | 0.047999884 | sp\|O22704\|CYP5F_ARATH | Cytochrome B5-like protein | IPR001199 (Cytochrome b5-like heme/steroid binding domain) | GO:0020037 (heme binding) |
| Niben101Scf04640g04002.1 | 165 | 244 | 0.726 | 0.021317731 | sp\|Q9ZNS1\|RS7_AVIMR | 40S ribosomal protein S7 | IPR000554 (Ribosomal protein S7e) | GO:0003735 (structural constituent of ribosome), GO:0005622 (intracellular), GO:0005840 (ribosome), GO:0006412 (translation) |
| Niben101Scf09554g01001.1 | 46 | 69 | 0.726 | 0.028353374 | gb\|ABR25403.1\| | transferase (transferring glycosyl group) [Oryza sativa Indica Group] | IPR002495 (Glycosyl transferase, family 8), IPR029044 (Nucleotide-diphospho-sugar transferases) | GO:0016757 (transferase activity, transferring glycosyl groups), GO:0047262 (polygalacturonate 4-alpha-galacturonosyltransferase activity) |
| Niben101Scf04300g00008.1 | 74 | 109 | 0.724 | 0.014363302 | sp\|A3CTG2\|RS10_METMJ | 30S ribosomal protein S10 | IPR001848 (Ribosomal protein S10), IPR027486 (Ribosomal protein S10 domain) | GO:0003723 (RNA binding), GO:0003735 (structural constituent of ribosome), GO:0005622 (intracellular), GO:0005840 (ribosome), GO:0006412 (translation) |
| Niben101Scf03962g01016.1 | 41 | 61 | 0.724 | 0.035923313 | AT4G02990.1 | Mitochondrial transcription termination factor family protein LENGTH=541 | IPR003690 (Mitochodrial transcription termination factor) | GO:0003690 (double-stranded DNA binding), GO:0005739 (mitochondrion), GO:0006355 (regulation of transcription, DNA-templated) |
| Niben101Scf12017g00012.1 | 310 | 461 | 0.722 | 0.013669653 | sp\|Q9ATF5\|RL18A_CASSA | 60S ribosomal protein L18a | IPR023573 (Ribosomal protein L18a/LX), IPR028877 (50S ribosomal protein L18Ae/60S ribosomal protein L20 and L18a) | GO:0003735 (structural constituent of ribosome), GO:0005840 (ribosome), GO:0006412 (translation) |
| Niben101Scf02610g00005.1 | 67 | 99 | 0.722 | 0.008929114 | AT1G70750.1 | Protein of unknown function, DUF593 LENGTH=749 | IPR007656 (Zein-binding domain) | NA |
| Niben101Scf04113g07009.1 | 79 | 117 | 0.721 | 0.029151641 | sp\|E1U332\|ALL12_OLEEU | Isoflavone reductase-like protein | IPR016040 (NAD(P)-binding domain) | NA |
| Niben101Scf06113g00012.1 | 60 | 89 | 0.721 | 0.028353374 | ref\|WP_014109590.1\| | branched-chain amino acid aminotransferase-like protein [Glaciecola nitratireducens] ref\|YP_004872710.1\| branched-chain amino acid aminotransferase-like protein [Glaciecola nitratireducens FR1064] gb\|AEP30717.1\| branched-chain amino acid aminotransferase-like protein [Glaciecola nitratireducens FR1064] | NA | NA |
| Niben101Scf04347g01029.1 | 71 | 105 | 0.719 | 0.038307416 | AT3G61770.1 | Acid phosphatase/vanadium-dependent haloperoxidase-related protein LENGTH=284 | IPR003832 (Acid phosphatase/vanadium-dependent haloperoxidase-related) | NA |
| Niben101Scf02383g00003.1 | 70 | 104 | 0.716 | 0.013669653 | sp\|P50346\|RLA0_SOYBN | 60S acidic ribosomal protein P0 | IPR001790 (Ribosomal protein L10/acidic P0) | GO:0005622 (intracellular), GO:0042254 (ribosome biogenesis) |
| Niben101Scf08272g00019.1 | 65 | 96 | 0.715 | 0.046205068 | sp\|P46291\|RL38_SOLLC | 60S ribosomal protein L38 | IPR002675 (Ribosomal protein L38e) | GO:0003735 (structural constituent of ribosome), GO:0005622 (intracellular), GO:0005840 (ribosome), GO:0006412 (translation) |
| Niben101Scf10796g00009.1 | 119 | 176 | 0.714 | 0.014480119 | sp\|Q1CZI7\|HTPG_MYXXD | Chaperone protein HtpG | IPR001404 (Heat shock protein Hsp90 family) | GO:0005524 (ATP binding), GO:0006457 (protein folding), GO:0006950 (response to stress), GO:0051082 (unfolded protein binding) |
| Niben101Scf03253g00017.1 | 62 | 91 | 0.714 | 0.00703408 | sp\|Q9FGZ9\|UBL5_ARATH | Ubiquitin-like protein 5 | IPR029071 (Ubiquitin-related domain) | GO:0005515 (protein binding) |
| Niben101Scf08488g02011.1 | 13841 | 20428 | 0.713 | 1.07494E-05 | sp\|Q06721\|RCA_ANASC | Ribulose bisphosphate carboxylase/oxygenase activase | IPR003959 (ATPase, AAA-type, core), IPR027417 (P-loop containing nucleoside triphosphate hydrolase) | GO:0005524 (ATP binding) |
| Niben101Scf03894g02023.1 | 80 | 119 | 0.713 | 0.00899827 | sp\|Q8WU67\|ABHD3_HUMAN | Abhydrolase domain-containing protein 3 | IPR029058 (Alpha/Beta hydrolase fold) | NA |
| Niben101Scf05438g02004.1 | 50 | 73 | 0.713 | 0.016642228 | AT2G27820.1 | prephenate dehydratase 1 LENGTH=424 | IPR001086 (Prephenate dehydratase), IPR002912 (ACT domain) | GO:0004664 (prephenate dehydratase activity), GO:0008152 (metabolic process), GO:0009094 (L-phenylalanine biosynthetic process), GO:0016597 (amino acid binding) |
| Niben101Scf00536g04007.1 | 118 | 174 | 0.712 | 0.01677562 | AT1G11545.1 | xyloglucan endotransglucosylase/hydrolase 8 LENGTH=305 | IPR008264 (Beta-glucanase), IPR013320 (Concanavalin A-like lectin/glucanase domain), IPR016455 (Xyloglucan endotransglucosylase/hydrolase) | GO:0004553 (hydrolase activity, hydrolyzing O-glycosyl compounds), GO:0005618 (cell wall), GO:0005975 (carbohydrate metabolic process), GO:0006073 (cellular glucan metabolic process), GO:0016762 (xyloglucan:xyloglucosyl transferase activity), GO:0048046 (apoplast) |
| Niben101Scf03030g00005.1 | 91 | 135 | 0.712 | 0.017478837 | AT5G39530.1 | Protein of unknown function (DUF1997) LENGTH=239 | IPR018971 (Protein of unknown function DUF1997) | NA |
| Niben101Scf03839g04019.1 | 1360 | 1994 | 0.71 | 0.000294263 | sp\|A9I7K9\|GCSP_BORPD | Glycine dehydrogenase (decarboxylating) | IPR020581 (Glycine cleavage system P protein) | GO:0003824 (catalytic activity), GO:0004375 (glycine dehydrogenase (decarboxylating) activity), GO:0006544 (glycine metabolic process), GO:0006546 (glycine catabolic process), GO:0030170 (pyridoxal phosphate binding), GO:0055114 (oxidation-reduction process) |
| Niben101Scf02819g02008.1 | 1300 | 1907 | 0.71 | 0.000464203 | sp\|Q9LND8\|ADSL2_ARATH | Delta-9 desaturase-like 2 protein | IPR015876 (Fatty acid desaturase, type 1, core) | GO:0006629 (lipid metabolic process), GO:0016717 (oxidoreductase activity, acting on paired donors, with oxidation of a pair of donors resulting in the reduction of molecular oxygen to two molecules of water), GO:0055114 (oxidation-reduction process) |
| Niben101Scf00578g06007.1 | 71 | 105 | 0.709 | 0.008887623 | sp\|Q9XEA8\|CYSK2_ORYSJ | Cysteine synthase | IPR005856 (Cysteine synthase K/M) | GO:0004124 (cysteine synthase activity), GO:0006535 (cysteine biosynthetic process from serine) |
| Niben101Scf06562g00007.1 | 52 | 77 | 0.709 | 0.019516741 | AT5G63060.1 | Sec14p-like phosphatidylinositol transfer family protein LENGTH=263 | IPR001251 (CRAL-TRIO domain), IPR011074 (CRAL/TRIO, N-terminal domain) | NA |
| Niben101Scf03538g02003.1 | 337 | 490 | 0.708 | 0.002074021 | sp\|P43672\|UUP_ECOLI | ABC transporter ATP-binding protein uup | IPR003439 (ABC transporter-like), IPR027417 (P-loop containing nucleoside triphosphate hydrolase) | GO:0005524 (ATP binding), GO:0016887 (ATPase activity) |
| Niben101Scf02886g01003.1 | 210 | 306 | 0.708 | 0.006368527 | NA | Unknown protein | NA | NA |
| Niben101Scf02911g03002.1 | 156 | 228 | 0.708 | 0.049414698 | sp\|P34091\|RL6_MESCR | 60S ribosomal protein L6 | IPR000915 (60S ribosomal protein L6E), IPR005568 (Ribosomal protein L6, N-terminal), IPR008991 (Translation protein SH3-like domain) | GO:0003735 (structural constituent of ribosome), GO:0005622 (intracellular), GO:0005840 (ribosome), GO:0006412 (translation) |
| Niben101Scf02027g00001.1 | 111 | 164 | 0.708 | 0.017619865 | AT5G51010.1 | Rubredoxin-like superfamily protein LENGTH=154 | IPR004039 (Rubredoxin-type fold) | GO:0005506 (iron ion binding) |
| Niben101Scf02911g04009.1 | 1794 | 2646 | 0.706 | 0.003741639 | NA | Unknown protein | NA | NA |
| Niben101Scf34874g00008.1 | 89 | 131 | 0.706 | 0.008072065 | sp\|Q01IH3\|PX114_ORYSI | Peroxisomal membrane protein 11-4 | IPR008733 (Peroxisomal biogenesis factor 11) | GO:0005779 (integral component of peroxisomal membrane), GO:0016559 (peroxisome fission) |
| Niben101Scf05767g02003.1 | 80 | 117 | 0.705 | 0.013996106 | sp\|O49884\|RL30_LUPLU | 60S ribosomal protein L30 | IPR000231 (Ribosomal protein L30e), IPR022991 (Ribosomal protein L30e, conserved site), IPR029064 (50S ribosomal protein L30e-like) | GO:0003735 (structural constituent of ribosome), GO:0005622 (intracellular), GO:0005840 (ribosome), GO:0006412 (translation) |
| Niben101Scf01433g08029.1 | 496 | 722 | 0.704 | 0.004118614 | AT4G38970.1 | fructose-bisphosphate aldolase 2 LENGTH=398 | IPR000741 (Fructose-bisphosphate aldolase, class-I), IPR013785 (Aldolase-type TIM barrel), IPR029768 (Fructose-bisphosphate aldolase class-I active site) | GO:0003824 (catalytic activity), GO:0004332 (fructose-bisphosphate aldolase activity), GO:0006096 (glycolytic process) |
| Niben101Scf10055g07005.1 | 81 | 118 | 0.704 | 0.009512105 | sp\|Q6FFL8\|MAO1_ACIAD | NAD-dependent malic enzyme | IPR001891 (Malic oxidoreductase) | GO:0004470 (malic enzyme activity), GO:0004471 (malate dehydrogenase (decarboxylating) (NAD+) activity), GO:0006108 (malate metabolic process), GO:0051287 (NAD binding), GO:0055114 (oxidation-reduction process) |
| Niben101Scf04749g00009.1 | 63 | 91 | 0.704 | 0.026573347 | sp\|A2Y0J7\|RU1A_ORYSI | U1 small nuclear ribonucleoprotein A | IPR012677 (Nucleotide-binding alpha-beta plait domain), IPR024888 (U1 small nuclear ribonucleoprotein A/U2 small nuclear ribonucleoprotein B'') | GO:0000166 (nucleotide binding), GO:0000398 (mRNA splicing, via spliceosome), GO:0003676 (nucleic acid binding), GO:0017069 (snRNA binding) |
| Niben101Scf01383g05012.1 | 449 | 656 | 0.703 | 0.000975519 | AT3G43720.1 | Bifunctional inhibitor/lipid-transfer protein/seed storage 2S albumin superfamily protein LENGTH=193 | IPR016140 (Bifunctional inhibitor/plant lipid transfer protein/seed storage helical domain) | NA |
| Niben101Scf22564g00001.1 | 147 | 217 | 0.703 | 0.002499248 | sp\|Q9FVI2\|ADF1_PETHY | Actin-depolymerizing factor 1 | IPR017904 (ADF/Cofilin/Destrin), IPR029006 (ADF-H/Gelsolin-like domain) | GO:0003779 (actin binding), GO:0005622 (intracellular), GO:0015629 (actin cytoskeleton), GO:0030042 (actin filament depolymerization) |
| Niben101Scf00735g06024.1 | 87 | 128 | 0.703 | 0.010904347 | dbj\|BAM13304.1\| | ozone-responsive stress related protein [Oryza officinalis] | IPR009515 (Protein of unknown function DUF1138) | NA |
| Niben101Scf00774g01001.1 | 401 | 581 | 0.702 | 0.002683524 | AT4G30950.1 | fatty acid desaturase 6 LENGTH=448 | IPR005804 (Fatty acid desaturase, type 1) | GO:0006629 (lipid metabolic process) |
| Niben101Scf11991g01002.1 | 314 | 454 | 0.701 | 0.018606179 | AT4G30950.1 | fatty acid desaturase 6 LENGTH=448 | IPR005804 (Fatty acid desaturase, type 1) | GO:0006629 (lipid metabolic process) |
| Niben101Scf09387g03016.1 | 619 | 907 | 0.7 | 0.001185914 | sp\|Q11HP9\|EFG_CHESB | Elongation factor G | IPR004540 (Translation elongation factor EFG/EF2), IPR005225 (Small GTP-binding protein domain), IPR027417 (P-loop containing nucleoside triphosphate hydrolase) | GO:0003746 (translation elongation factor activity), GO:0003924 (GTPase activity), GO:0005525 (GTP binding), GO:0005622 (intracellular), GO:0006414 (translational elongation) |
| Niben101Scf32746g00006.1 | 448 | 656 | 0.697 | 0.012716326 | sp\|A8ACD2\|RL14_IGNH4 | 50S ribosomal protein L14 | IPR000218 (Ribosomal protein L14b/L23e), IPR023571 (Ribosomal protein L14 domain) | GO:0003735 (structural constituent of ribosome), GO:0005840 (ribosome), GO:0006412 (translation) |
| Niben101Scf02937g04001.1 | 118 | 172 | 0.694 | 0.016560099 | AT1G72330.3 | alanine aminotransferase 2 LENGTH=553 | IPR015424 (Pyridoxal phosphate-dependent transferase) | GO:0003824 (catalytic activity), GO:0009058 (biosynthetic process), GO:0030170 (pyridoxal phosphate binding) |
| Niben101Scf00176g05007.1 | 43 | 63 | 0.694 | 0.041737241 | ref\|WP_016897861.1\| | MULTISPECIES: phospho-3-sulfolactate synthase [Aerococcus] | IPR003830 ((2R)-phospho-3-sulpholactate synthase, ComA), IPR013785 (Aldolase-type TIM barrel) | GO:0003824 (catalytic activity), GO:0019295 (coenzyme M biosynthetic process) |
| Niben101Scf03094g05009.1 | 109 | 160 | 0.692 | 0.033674151 | sp\|Q6AYK6\|CYBP_RAT | Calcyclin-binding protein | IPR007699 (SGS), IPR008978 (HSP20-like chaperone), IPR015120 (Siah interacting protein, N-terminal) | NA |
| Niben101Scf09792g03014.1 | 45 | 65 | 0.692 | 0.041496045 | sp\|Q93XX8\|NOP10_ARATH | H/ACA ribonucleoprotein complex subunit 3-like protein | IPR007264 (H/ACA ribonucleoprotein complex, subunit Nop10) | GO:0001522 (pseudouridine synthesis), GO:0030515 (snoRNA binding), GO:0042254 (ribosome biogenesis), GO:0072588 (box H/ACA RNP complex) |
| Niben101Scf04838g02010.1 | 88 | 127 | 0.69 | 0.018606179 | sp\|Q9SX20\|PTR18_ARATH | Protein NRT1/ PTR FAMILY 3.1 | IPR000109 (Proton-dependent oligopeptide transporter family), IPR020846 (Major facilitator superfamily domain) | GO:0005215 (transporter activity), GO:0006810 (transport), GO:0016020 (membrane) |
| Niben101Scf03113g02012.1 | 165 | 242 | 0.689 | 0.028338156 | sp\|P41101\|RL27_SOLTU | 60S ribosomal protein L27 | IPR001141 (Ribosomal protein L27e), IPR008991 (Translation protein SH3-like domain) | GO:0003735 (structural constituent of ribosome), GO:0005622 (intracellular), GO:0005840 (ribosome), GO:0006412 (translation) |
| Niben101Scf00132g01003.1 | 49 | 70 | 0.689 | 0.029308011 | AT1G64500.1 | Glutaredoxin family protein LENGTH=368 | IPR012336 (Thioredoxin-like fold) | GO:0009055 (electron carrier activity), GO:0015035 (protein disulfide oxidoreductase activity), GO:0045454 (cell redox homeostasis) |
| Niben101Scf03396g01003.1 | 508 | 745 | 0.685 | 0.042077774 | sp\|A9NA82\|CH60_COXBR | 60 kDa chaperonin | IPR002423 (Chaperonin Cpn60/TCP-1), IPR027409 (GroEL-like apical domain), IPR027413 (GroEL-like equatorial domain) | GO:0005524 (ATP binding), GO:0005737 (cytoplasm), GO:0006457 (protein folding), GO:0042026 (protein refolding), GO:0044267 (cellular protein metabolic process) |
| Niben101Scf14410g00001.1 | 229 | 335 | 0.684 | 0.028067933 | sp\|Q6C9R9\|SFH5_YARLI | Phosphatidylinositol transfer protein SFH5 | IPR001251 (CRAL-TRIO domain), IPR009038 (GOLD), IPR011074 (CRAL/TRIO, N-terminal domain) | GO:0006810 (transport), GO:0016021 (integral component of membrane) |
| Niben101Scf01101g04010.1 | 68 | 98 | 0.684 | 0.018801441 | ref\|WP_030619022.1\| | glycosidase [Streptomyces achromogenes] | IPR013783 (Immunoglobulin-like fold), IPR013784 (Carbohydrate-binding-like fold), IPR015902 (Glycoside hydrolase, family 13) | GO:0003824 (catalytic activity), GO:0030246 (carbohydrate binding), GO:2001070 (starch binding) |
| Niben101Scf05950g01001.1 | 337 | 487 | 0.683 | 0.002522179 | sp\|A8MQA2\|NLTPD_ARATH | Non-specific lipid-transfer protein 13 | IPR016140 (Bifunctional inhibitor/plant lipid transfer protein/seed storage helical domain) | NA |
| Niben101Scf00883g03018.1 | 81 | 117 | 0.683 | 0.023540024 | AT3G49780.1 | phytosulfokine 4 precursor LENGTH=79 | IPR009438 (Phytosulfokine) | GO:0005576 (extracellular region), GO:0008083 (growth factor activity), GO:0008283 (cell proliferation) |
| Niben101Scf05492g00001.1 | 330 | 470 | 0.682 | 0.010940991 | AT3G07700.3 | Protein kinase superfamily protein LENGTH=724 | IPR011009 (Protein kinase-like domain) | GO:0016772 (transferase activity, transferring phosphorus-containing groups) |
| Niben101Scf01427g00011.1 | 129 | 187 | 0.682 | 0.005241549 | sp\|Q3ZBR5\|TTC1_BOVIN | Tetratricopeptide repeat protein 1 | IPR000270 (Phox/Bem1p), IPR011990 (Tetratricopeptide-like helical domain) | GO:0005515 (protein binding) |
| Niben101Scf01587g02019.1 | 234 | 341 | 0.681 | 0.005837906 | ref\|WP_008310550.1\| | Protein of unknown function (DUF3067) [Leptolyngbya sp. PCC 6406] | IPR021420 (Protein of unknown function DUF3067) | NA |
| Niben101Scf10400g00005.1 | 827 | 1188 | 0.68 | 0.005837906 | sp\|A5GV53\|CH60_SYNR3 | 60 kDa chaperonin | IPR002423 (Chaperonin Cpn60/TCP-1), IPR027409 (GroEL-like apical domain), IPR027413 (GroEL-like equatorial domain) | GO:0005524 (ATP binding), GO:0005737 (cytoplasm), GO:0006457 (protein folding), GO:0042026 (protein refolding), GO:0044267 (cellular protein metabolic process) |
| Niben101Scf03717g01006.1 | 206 | 295 | 0.675 | 0.003741639 | AT3G26570.1 | phosphate transporter 2;1 LENGTH=613 | IPR001204 (Phosphate transporter) | GO:0005315 (inorganic phosphate transmembrane transporter activity), GO:0006817 (phosphate ion transport), GO:0016020 (membrane) |
| Niben101Scf07664g01009.1 | 53 | 77 | 0.675 | 0.019839073 | AT5G63870.2 | serine/threonine phosphatase 7 LENGTH=413 | IPR029052 (Metallo-dependent phosphatase-like) | GO:0016787 (hydrolase activity) |
| Niben101Scf14440g00003.1 | 99 | 143 | 0.674 | 0.047590729 | sp\|B0R3A3\|SYP_HALS3 | Proline--tRNA ligase | IPR002316 (Proline-tRNA ligase, class IIa), IPR017449 (Prolyl-tRNA synthetase, class II) | GO:0000166 (nucleotide binding), GO:0004812 (aminoacyl-tRNA ligase activity), GO:0004827 (proline-tRNA ligase activity), GO:0005524 (ATP binding), GO:0005737 (cytoplasm), GO:0006418 (tRNA aminoacylation for protein translation), GO:0006433 (prolyl-tRNA aminoacylation) |
| Niben101Scf05018g00001.1 | 77 | 111 | 0.673 | 0.012158774 | sp\|P41129\|RL132_BRANA | 60S ribosomal protein L13-2 | IPR001380 (Ribosomal protein L13e) | GO:0003735 (structural constituent of ribosome), GO:0005622 (intracellular), GO:0005840 (ribosome), GO:0006412 (translation) |
| Niben101Scf12360g00011.1 | 75 | 107 | 0.673 | 0.015270046 | gb\|KEH27963.1\| | Zn-dependent hydrolase of the beta-lactamase fold protein [Medicago truncatula] | IPR001279 (Beta-lactamase-like) | NA |
| Niben101Scf09678g00006.1 | 63 | 89 | 0.669 | 0.048340163 | sp\|Q76LC6\|RBM24_DANRE | RNA-binding protein 24 | IPR012677 (Nucleotide-binding alpha-beta plait domain) | GO:0000166 (nucleotide binding), GO:0003676 (nucleic acid binding) |
| Niben101Scf01980g10004.1 | 376 | 536 | 0.668 | 0.000792199 | AT1G22400.1 | UDP-Glycosyltransferase superfamily protein LENGTH=489 | IPR002213 (UDP-glucuronosyl/UDP-glucosyltransferase) | GO:0008152 (metabolic process), GO:0016758 (transferase activity, transferring hexosyl groups) |
| Niben101Scf11802g02006.1 | 315 | 448 | 0.667 | 0.042512295 | sp\|Q089P5\|RS17_SHEFN | 30S ribosomal protein S17 | IPR000266 (Ribosomal protein S17), IPR012340 (Nucleic acid-binding, OB-fold) | GO:0003735 (structural constituent of ribosome), GO:0005622 (intracellular), GO:0005840 (ribosome), GO:0006412 (translation) |
| Niben101Scf01232g03010.1 | 216 | 305 | 0.667 | 0.012579582 | AT5G52570.1 | beta-carotene hydroxylase 2 LENGTH=303 | IPR006694 (Fatty acid hydroxylase) | GO:0005506 (iron ion binding), GO:0006633 (fatty acid biosynthetic process), GO:0016491 (oxidoreductase activity), GO:0055114 (oxidation-reduction process) |
| Niben101Scf02289g02010.1 | 106 | 152 | 0.667 | 0.006117304 | sp\|Q9CAA4\|BIM2_ARATH | Transcription factor BIM2 | NA | NA |
| Niben101Scf05458g00003.1 | 104 | 148 | 0.667 | 0.006764772 | AT5G48930.1 | hydroxycinnamoyl-CoA shikimate/quinate hydroxycinnamoyl transferase LENGTH=433 | IPR003480 (Transferase), IPR023213 (Chloramphenicol acetyltransferase-like domain) | GO:0016747 (transferase activity, transferring acyl groups other than amino-acyl groups) |
| Niben101Scf07508g08003.1 | 99 | 140 | 0.666 | 0.020929164 | gb\|KEH33317.1\| | transmembrane protein, putative [Medicago truncatula] | NA | NA |
| Niben101Scf06896g00005.1 | 120 | 171 | 0.664 | 0.01966796 | AT4G34200.1 | D-3-phosphoglycerate dehydrogenase LENGTH=603 | IPR006236 (D-3-phosphoglycerate dehydrogenase), IPR015878 (S-adenosyl-L-homocysteine hydrolase, NAD binding domain), IPR016040 (NAD(P)-binding domain), IPR029009 (Allosteric substrate binding domain), IPR029752 (D-isomer specific 2-hydroxyacid dehydrogenase, NAD-binding domain conserved site 1), IPR029753 (D-isomer specific 2-hydroxyacid dehydrogenase, NAD-binding domain conserved site) | GO:0004617 (phosphoglycerate dehydrogenase activity), GO:0006564 (L-serine biosynthetic process), GO:0008152 (metabolic process), GO:0016597 (amino acid binding), GO:0016616 (oxidoreductase activity, acting on the CH-OH group of donors, NAD or NADP as acceptor), GO:0051287 (NAD binding), GO:0055114 (oxidation-reduction process) |
| Niben101Scf01422g00009.1 | 147 | 210 | 0.663 | 0.008114478 | ref\|WP_009457354.1\| | regulator [Fischerella sp. JSC-11] gb\|EHC13058.1\| response regulator receiver protein [Fischerella sp. JSC-11] | IPR022552 (Uncharacterised protein family Ycf55) | NA |
| Niben101Scf00123g06011.1 | 129 | 183 | 0.663 | 0.002255897 | sp\|Q9SSB8\|CX5B2_ARATH | Cytochrome c oxidase subunit 5b-2, mitochondrial | IPR002124 (Cytochrome c oxidase, subunit Vb) | GO:0004129 (cytochrome-c oxidase activity), GO:0005740 (mitochondrial envelope) |
| Niben101Scf00132g01003.1 | 176 | 253 | 0.662 | 0.024575321 | AT1G64500.1 | Glutaredoxin family protein LENGTH=368 | IPR012336 (Thioredoxin-like fold) | GO:0009055 (electron carrier activity), GO:0015035 (protein disulfide oxidoreductase activity), GO:0045454 (cell redox homeostasis) |
| Niben101Scf08774g00012.1 | 101 | 142 | 0.661 | 0.017135045 | sp\|Q7U4J2\|RL29_SYNPX | 50S ribosomal protein L29 | IPR001854 (Ribosomal protein L29) | GO:0003735 (structural constituent of ribosome), GO:0005622 (intracellular), GO:0005840 (ribosome), GO:0006412 (translation) |
| Niben101Scf28395g00001.1 | 237 | 332 | 0.66 | 0.014804059 | sp\|Q06721\|RCA_ANASC | Ribulose bisphosphate carboxylase/oxygenase activase | IPR003959 (ATPase, AAA-type, core), IPR027417 (P-loop containing nucleoside triphosphate hydrolase) | GO:0005524 (ATP binding) |
| Niben101Scf01521g16018.1 | 147 | 209 | 0.659 | 0.004420383 | ref\|WP_025946871.1\| | MULTISPECIES: ribonuclease P [Prochlorococcus] | IPR005182 (Domain of unknown function DUF304) | NA |
| Niben101Scf01513g01010.1 | 256 | 362 | 0.655 | 0.002105966 | sp\|P70718\|6PGD_AGGAC | 6-phosphogluconate dehydrogenase, decarboxylating | IPR008927 (6-phosphogluconate dehydrogenase, C-terminal-like), IPR015815 (Hydroxy monocarboxylic acid anion dehydrogenase, HIBADH-type), IPR016040 (NAD(P)-binding domain) | GO:0004616 (phosphogluconate dehydrogenase (decarboxylating) activity), GO:0006098 (pentose-phosphate shunt), GO:0016491 (oxidoreductase activity), GO:0016616 (oxidoreductase activity, acting on the CH-OH group of donors, NAD or NADP as acceptor), GO:0050662 (coenzyme binding), GO:0051287 (NAD binding), GO:0055114 (oxidation-reduction process) |
| Niben101Scf02613g03001.1 | 54 | 76 | 0.655 | 0.04552707 | AT2G26640.1 | 3-ketoacyl-CoA synthase 11 LENGTH=509 | IPR012392 (Very-long-chain 3-ketoacyl-CoA synthase), IPR016039 (Thiolase-like) | GO:0003824 (catalytic activity), GO:0006633 (fatty acid biosynthetic process), GO:0008152 (metabolic process), GO:0008610 (lipid biosynthetic process), GO:0016020 (membrane), GO:0016747 (transferase activity, transferring acyl groups other than amino-acyl groups) |
| Niben101Scf00774g01030.1 | 72 | 101 | 0.654 | 0.04881534 | sp\|Q3T0Z8\|RUXF_BOVIN | Small nuclear ribonucleoprotein F | IPR010920 (Like-Sm (LSM) domain) | NA |
| Niben101Scf01259g04002.1 | 78 | 111 | 0.653 | 0.031609236 | sp\|Q4R4P3\|CYBP_MACFA | Calcyclin-binding protein | IPR007699 (SGS), IPR008978 (HSP20-like chaperone), IPR015120 (Siah interacting protein, N-terminal) | NA |
| Niben101Scf00109g00017.1 | 133 | 188 | 0.651 | 0.020470188 | ref\|XP_003630569.1\| | Mitochondrial phosphate carrier protein [Medicago truncatula] | IPR018108 (Mitochondrial substrate/solute carrier), IPR023395 (Mitochondrial carrier domain) | NA |
| Niben101Scf02990g05004.1 | 105 | 149 | 0.647 | 0.015084334 | sp\|Q8LDW9\|XTH9_ARATH | Xyloglucan endotransglucosylase/hydrolase protein 9 | IPR013320 (Concanavalin A-like lectin/glucanase domain), IPR016455 (Xyloglucan endotransglucosylase/hydrolase) | GO:0004553 (hydrolase activity, hydrolyzing O-glycosyl compounds), GO:0005618 (cell wall), GO:0005975 (carbohydrate metabolic process), GO:0006073 (cellular glucan metabolic process), GO:0016762 (xyloglucan:xyloglucosyl transferase activity), GO:0048046 (apoplast) |
| Niben101Scf04738g02001.1 | 851 | 1201 | 0.646 | 0.030875757 | sp\|Q8LB81\|GDL79_ARATH | GDSL esterase/lipase | IPR013830 (SGNH hydrolase-type esterase domain) | GO:0006629 (lipid metabolic process), GO:0016788 (hydrolase activity, acting on ester bonds) |
| Niben101Scf02182g18008.1 | 285 | 401 | 0.645 | 0.014843984 | sp\|O82528\|RL15_PETHY | 60S ribosomal protein L15 | IPR000439 (Ribosomal protein L15e) | GO:0003735 (structural constituent of ribosome), GO:0005622 (intracellular), GO:0005840 (ribosome), GO:0006412 (translation) |
| Niben101Scf09436g05005.1 | 98 | 138 | 0.645 | 0.039751815 | sp\|Q6LXN1\|RS13_METMP | 30S ribosomal protein S13 | IPR001892 (Ribosomal protein S13), IPR010979 (Ribosomal protein S13-like, H2TH), IPR027437 (30s ribosomal protein S13, C-terminal) | GO:0003676 (nucleic acid binding), GO:0003723 (RNA binding), GO:0003735 (structural constituent of ribosome), GO:0005622 (intracellular), GO:0005840 (ribosome), GO:0006412 (translation) |
| Niben101Scf06504g01026.1 | 607 | 854 | 0.644 | 0.011205231 | AT1G64500.1 | Glutaredoxin family protein LENGTH=368 | IPR012336 (Thioredoxin-like fold) | GO:0009055 (electron carrier activity), GO:0015035 (protein disulfide oxidoreductase activity), GO:0045454 (cell redox homeostasis) |
| Niben101Scf22589g00001.1 | 269 | 379 | 0.644 | 0.007747848 | AT5G57800.1 | Fatty acid hydroxylase superfamily LENGTH=632 | IPR006694 (Fatty acid hydroxylase), IPR021940 (Uncharacterised domain Wax2, C-terminal) | GO:0005506 (iron ion binding), GO:0006633 (fatty acid biosynthetic process), GO:0016491 (oxidoreductase activity), GO:0055114 (oxidation-reduction process) |
| Niben101Scf11852g00012.1 | 2001 | 2807 | 0.639 | 0.004152953 | sp\|Q9SMB4\|PSBS_TOBAC | Photosystem II 22 kDa protein, chloroplastic | IPR022796 (Chlorophyll A-B binding protein), IPR023329 (Chlorophyll a/b binding protein domain) | NA |
| Niben101Scf01689g04015.1 | 336 | 470 | 0.639 | 0.021846818 | sp\|Q9SW09\|RS101_ARATH | 40S ribosomal protein S10-1 | IPR005326 (Plectin/S10, N-terminal) | NA |
| Niben101Scf31398g00007.1 | 141 | 197 | 0.636 | 0.016245873 | sp\|Q9FJ40\|GDL86_ARATH | GDSL esterase/lipase | IPR013830 (SGNH hydrolase-type esterase domain) | GO:0006629 (lipid metabolic process), GO:0016788 (hydrolase activity, acting on ester bonds) |
| Niben101Scf05250g02013.1 | 368 | 515 | 0.635 | 0.027040915 | sp\|Q67A25\|NCS_THLFG | S-norcoclaurine synthase | IPR000916 (Bet v I domain), IPR023393 (START-like domain) | GO:0006952 (defense response), GO:0009607 (response to biotic stimulus) |
| Niben101Scf13748g00016.1 | 64 | 89 | 0.635 | 0.048394988 | sp\|Q6GGC1\|DNAJ_STAAR | Chaperone protein DnaJ | IPR001623 (DnaJ domain) | NA |
| Niben101Scf09552g01001.1 | 230 | 324 | 0.631 | 0.033734963 | sp\|B4S6P7\|DNAK_PROA2 | Chaperone protein DnaK | IPR013126 (Heat shock protein 70 family), IPR029047 (Heat shock protein 70kD, peptide-binding domain), IPR029048 (Heat shock protein 70kD, C-terminal domain) | GO:0005524 (ATP binding), GO:0006457 (protein folding), GO:0051082 (unfolded protein binding) |
| Niben101Scf12154g01009.1 | 730 | 1019 | 0.629 | 0.02229234 | sp\|Q4R888\|HS71L_MACFA | Heat shock 70 kDa protein 1-like | IPR013126 (Heat shock protein 70 family), IPR029047 (Heat shock protein 70kD, peptide-binding domain), IPR029048 (Heat shock protein 70kD, C-terminal domain) | NA |
| Niben101Scf04323g04020.1 | 120 | 167 | 0.628 | 0.019829365 | sp\|Q84JP1\|NFYA7_ARATH | Nuclear transcription factor Y subunit A-7 | IPR001289 (CCAAT-binding transcription factor, subunit B) | GO:0003677 (DNA binding), GO:0003700 (sequence-specific DNA binding transcription factor activity), GO:0006355 (regulation of transcription, DNA-templated), GO:0016602 (CCAAT-binding factor complex) |
| Niben101Scf14325g00002.1 | 117 | 163 | 0.626 | 0.009705938 | sp\|A0M527\|MURG_GRAFK | UDP-N-acetylglucosamine--N-acetylmuramyl-(pentapeptide) pyrophosphoryl-undecaprenol N-acetylglucosamine transferase | IPR007235 (Glycosyl transferase, family 28, C-terminal), IPR009695 (Diacylglycerol glucosyltransferase, N-terminal) | GO:0005975 (carbohydrate metabolic process), GO:0009247 (glycolipid biosynthetic process), GO:0016758 (transferase activity, transferring hexosyl groups), GO:0030246 (carbohydrate binding), GO:0030259 (lipid glycosylation) |
| Niben101Scf32851g00036.1 | 332 | 461 | 0.625 | 0.025726479 | sp\|P46300\|RS4_SOLTU | 40S ribosomal protein S4 | IPR000876 (Ribosomal protein S4e) | GO:0003723 (RNA binding), GO:0003735 (structural constituent of ribosome), GO:0005622 (intracellular), GO:0005840 (ribosome), GO:0006412 (translation) |
| Niben101Scf07123g00001.1 | 120 | 165 | 0.622 | 0.010069307 | sp\|Q56YA5\|SGAT_ARATH | Serine--glyoxylate aminotransferase | IPR015424 (Pyridoxal phosphate-dependent transferase), IPR024169 (Serine-pyruvate aminotransferase/2-aminoethylphosphonate-pyruvate transaminase) | GO:0003824 (catalytic activity), GO:0008152 (metabolic process), GO:0030170 (pyridoxal phosphate binding) |
| Niben101Scf01764g00009.1 | 270 | 374 | 0.62 | 0.030098874 | sp\|O82204\|RL281_ARATH | 60S ribosomal protein L28-1 | IPR002672 (Ribosomal protein L28e), IPR029004 (Ribosomal L28e/Mak16) | GO:0003735 (structural constituent of ribosome), GO:0005622 (intracellular), GO:0005840 (ribosome), GO:0006412 (translation) |
| Niben101Scf02840g04002.1 | 316 | 436 | 0.619 | 0.016233329 | sp\|Q9C554\|EXPA1_ARATH | Expansin-A1 | IPR007118 (Expansin/Lol pI) | GO:0005576 (extracellular region), GO:0009664 (plant-type cell wall organization) |
| Niben101Scf05078g08029.1 | 487 | 672 | 0.618 | 0.017823263 | AT3G43720.1 | Bifunctional inhibitor/lipid-transfer protein/seed storage 2S albumin superfamily protein LENGTH=193 | IPR016140 (Bifunctional inhibitor/plant lipid transfer protein/seed storage helical domain) | NA |
| Niben101Ctg16098g00002.1 | 458 | 627 | 0.618 | 0.029205286 | AT2G37180.1 | Aquaporin-like superfamily protein LENGTH=285 | IPR000425 (Major intrinsic protein), IPR023271 (Aquaporin-like) | GO:0005215 (transporter activity), GO:0006810 (transport), GO:0016020 (membrane) |
| Niben101Scf07576g01015.1 | 63 | 88 | 0.617 | 0.046596961 | sp\|P92949\|FRO2_ARATH | Ferric reduction oxidase 2 | IPR013121 (Ferric reductase, NAD binding), IPR013130 (Ferric reductase transmembrane component-like domain), IPR017938 (Riboflavin synthase-like beta-barrel) | GO:0016491 (oxidoreductase activity), GO:0055114 (oxidation-reduction process) |
| Niben101Scf03538g02003.1 | 64 | 87 | 0.615 | 0.041531077 | sp\|P43672\|UUP_ECOLI | ABC transporter ATP-binding protein uup | IPR003439 (ABC transporter-like), IPR027417 (P-loop containing nucleoside triphosphate hydrolase) | GO:0005524 (ATP binding), GO:0016887 (ATPase activity) |
| Niben101Scf03133g00003.1 | 1203 | 1655 | 0.613 | 0.010069307 | sp\|Q5HF86\|G3P2_STAAC | Glyceraldehyde-3-phosphate dehydrogenase 2 | IPR003823 (Domain of unknown function CP12), IPR020831 (Glyceraldehyde/Erythrose phosphate dehydrogenase family) | GO:0016620 (oxidoreductase activity, acting on the aldehyde or oxo group of donors, NAD or NADP as acceptor), GO:0055114 (oxidation-reduction process) |
| Niben101Scf00626g01002.1 | 434 | 596 | 0.611 | 0.033427574 | sp\|Q9M352\|RL362_ARATH | 60S ribosomal protein L36-2 | IPR000509 (Ribosomal protein L36e) | GO:0003735 (structural constituent of ribosome), GO:0005622 (intracellular), GO:0005840 (ribosome), GO:0006412 (translation) |
| Niben101Scf01585g01017.1 | 147 | 201 | 0.611 | 0.038563996 | sp\|Q9LHP1\|RL74_ARATH | 60S ribosomal protein L7-4 | IPR005998 (Ribosomal protein L7, eukaryotic) | NA |
| Niben101Scf00087g01004.1 | 106 | 146 | 0.61 | 0.028495819 | AT2G24395.1 | chaperone protein dnaJ-related LENGTH=132 | NA | NA |
| Niben101Scf10556g00024.1 | 390 | 531 | 0.607 | 0.006764772 | AT5G07020.1 | proline-rich family protein LENGTH=235 | NA | NA |
| Niben101Scf07648g01017.1 | 94 | 129 | 0.607 | 0.044152419 | gb\|ACG27246.1\| | tubulin alpha-6 chain [Zea mays] | NA | NA |
| Niben101Scf29588g00003.1 | 93 | 129 | 0.607 | 0.032902691 | sp\|Q8GYX0\|MOB1_ARATH | MOB kinase activator-like 1 | IPR005301 (Mob1/phocein) | NA |
| Niben101Scf15760g00012.1 | 270 | 372 | 0.605 | 0.036947323 | sp\|Q9SCU0\|SDR2A_ARATH | Short-chain dehydrogenase reductase 2a | IPR002347 (Glucose/ribitol dehydrogenase) | GO:0008152 (metabolic process), GO:0016491 (oxidoreductase activity) |
| Niben101Scf04252g01009.1 | 165 | 226 | 0.603 | 0.005944857 | sp\|Q8YHC2\|SSB_BRUME | Single-stranded DNA-binding protein | IPR000424 (Primosome PriB/single-strand DNA-binding) | GO:0003697 (single-stranded DNA binding), GO:0006260 (DNA replication) |
| Niben101Scf04019g02005.1 | 75 | 103 | 0.6 | 0.040469735 | sp\|O67623\|DNAJ1_AQUAE | Chaperone protein DnaJ 1 | IPR001623 (DnaJ domain) | NA |
| Niben101Scf01182g01006.1 | 260 | 354 | 0.598 | 0.042381889 | sp\|Q9XEG7\|RS3A_TORRU | 40S ribosomal protein S3a | IPR001593 (Ribosomal protein S3Ae) | NA |
| Niben101Scf05615g01004.1 | 170 | 229 | 0.597 | 0.046550874 | AT2G26710.1 | Cytochrome P450 superfamily protein LENGTH=520 | IPR001128 (Cytochrome P450) | GO:0005506 (iron ion binding), GO:0016705 (oxidoreductase activity, acting on paired donors, with incorporation or reduction of molecular oxygen), GO:0020037 (heme binding), GO:0055114 (oxidation-reduction process) |
| Niben101Scf04847g00009.1 | 117 | 160 | 0.597 | 0.040940074 | sp\|Q9FM65\|FLA1_ARATH | Fasciclin-like arabinogalactan protein 1 | IPR000782 (FAS1 domain) | NA |
| Niben101Scf03377g05007.1 | 196 | 266 | 0.595 | 0.005058165 | sp\|B0U3J7\|DNAJ_XYLFM | Chaperone protein DnaJ | IPR001623 (DnaJ domain), IPR002939 (Chaperone DnaJ, C-terminal) | GO:0006457 (protein folding), GO:0051082 (unfolded protein binding) |
| Niben101Scf01774g10020.1 | 236 | 318 | 0.589 | 0.006119011 | ref\|WP_032378363.1\| | membrane protein [Rhodococcus fascians] | IPR021414 (Protein of unknown function DUF3054) | NA |
| Niben101Scf05688g08010.1 | 4453 | 5988 | 0.587 | 0.000407745 | AT1G56190.1 | Phosphoglycerate kinase family protein LENGTH=478 | IPR001576 (Phosphoglycerate kinase) | GO:0004618 (phosphoglycerate kinase activity), GO:0006096 (glycolytic process) |
| Niben101Scf10157g05001.1 | 88 | 119 | 0.587 | 0.04468573 | sp\|P10691\|SUS1_SOLTU | Sucrose synthase | IPR012820 (Sucrose synthase, plant/cyanobacteria) | GO:0005985 (sucrose metabolic process), GO:0009058 (biosynthetic process), GO:0016157 (sucrose synthase activity) |
| Niben101Scf07244g03021.1 | 1139 | 1546 | 0.586 | 0.034295037 | gb\|AFW71223.1\| | meiosis 5 [Zea mays] | NA | NA |
| Niben101Scf04664g01003.1 | 5383 | 7197 | 0.585 | 0.001428352 | AT4G38970.1 | fructose-bisphosphate aldolase 2 LENGTH=398 | IPR000741 (Fructose-bisphosphate aldolase, class-I), IPR013785 (Aldolase-type TIM barrel), IPR029768 (Fructose-bisphosphate aldolase class-I active site) | GO:0003824 (catalytic activity), GO:0004332 (fructose-bisphosphate aldolase activity), GO:0006096 (glycolytic process) |
| Niben101Scf08690g02001.1 | 125 | 169 | 0.584 | 0.02583416 | AT3G61770.1 | Acid phosphatase/vanadium-dependent haloperoxidase-related protein LENGTH=284 | IPR003832 (Acid phosphatase/vanadium-dependent haloperoxidase-related) | NA |
| Niben101Scf06236g00008.1 | 2129 | 2879 | 0.583 | 0.014561325 | AT2G45290.1 | Transketolase LENGTH=741 | IPR005478 (Transketolase, bacterial-like), IPR009014 (Transketolase, C-terminal/Pyruvate-ferredoxin oxidoreductase, domain II), IPR029061 (Thiamin diphosphate-binding fold) | GO:0003824 (catalytic activity), GO:0004802 (transketolase activity), GO:0008152 (metabolic process) |
| Niben101Scf00797g18009.1 | 1390 | 1853 | 0.583 | 0.015261588 | sp\|Q56YA5\|SGAT_ARATH | Serine--glyoxylate aminotransferase | IPR015424 (Pyridoxal phosphate-dependent transferase), IPR024169 (Serine-pyruvate aminotransferase/2-aminoethylphosphonate-pyruvate transaminase) | GO:0003824 (catalytic activity), GO:0008152 (metabolic process), GO:0030170 (pyridoxal phosphate binding) |
| Niben101Scf01958g02007.1 | 2290 | 3089 | 0.58 | 0.037243984 | NA | Unknown protein | NA | NA |
| Niben101Scf04024g02026.1 | 108 | 145 | 0.58 | 0.020716644 | sp\|P54086\|TATC_SYNY3 | Sec-independent protein translocase protein TatC | IPR002033 (Sec-independent periplasmic protein translocase TatC) | GO:0016021 (integral component of membrane) |
| Niben101Scf21171g00004.1 | 616 | 826 | 0.579 | 0.026307017 | sp\|O82528\|RL15_PETHY | 60S ribosomal protein L15 | IPR000439 (Ribosomal protein L15e) | GO:0003735 (structural constituent of ribosome), GO:0005622 (intracellular), GO:0005840 (ribosome), GO:0006412 (translation) |
| Niben101Scf00123g06029.1 | 91 | 122 | 0.579 | 0.044802988 | sp\|Q9QYL8\|LYPA2_RAT | Acyl-protein thioesterase 2 | IPR029058 (Alpha/Beta hydrolase fold) | GO:0016787 (hydrolase activity) |
| Niben101Scf08091g00018.1 | 72 | 96 | 0.579 | 0.046676126 | sp\|P62194\|PRS8_BOVIN | 26S protease regulatory subunit 8 | IPR003959 (ATPase, AAA-type, core), IPR005937 (26S proteasome subunit P45), IPR027417 (P-loop containing nucleoside triphosphate hydrolase) | GO:0005524 (ATP binding), GO:0005737 (cytoplasm), GO:0016787 (hydrolase activity), GO:0030163 (protein catabolic process) |
| Niben101Scf00603g08015.1 | 2487 | 3315 | 0.574 | 0.01052324 | AT2G45290.1 | Transketolase LENGTH=741 | IPR005478 (Transketolase, bacterial-like), IPR009014 (Transketolase, C-terminal/Pyruvate-ferredoxin oxidoreductase, domain II), IPR029061 (Thiamin diphosphate-binding fold) | GO:0003824 (catalytic activity), GO:0004802 (transketolase activity), GO:0008152 (metabolic process) |
| Niben101Scf06795g00001.1 | 174 | 235 | 0.574 | 0.037425584 | AT1G26761.1 | Arabinanase/levansucrase/invertase LENGTH=444 | IPR007184 (Mannoside phosphorylase), IPR023296 (Glycosyl hydrolase, five-bladed beta-propellor domain) | NA |
| Niben101Scf01848g04005.1 | 156 | 209 | 0.568 | 0.045654906 | sp\|C3N5S0\|EF2_SULIA | Elongation factor 2 | IPR000640 (Translation elongation factor EFG, V domain), IPR009000 (Translation protein, beta-barrel domain), IPR009022 (Elongation factor G, III-V domain), IPR020568 (Ribosomal protein S5 domain 2-type fold) | GO:0005525 (GTP binding) |
| Niben101Scf05368g03015.1 | 16523 | 21982 | 0.565 | 0.000559185 | sp\|Q06721\|RCA_ANASC | Ribulose bisphosphate carboxylase/oxygenase activase | IPR003959 (ATPase, AAA-type, core), IPR027417 (P-loop containing nucleoside triphosphate hydrolase) | GO:0005524 (ATP binding) |
| Niben101Scf07826g04012.1 | 352 | 469 | 0.563 | 0.012515861 | sp\|Q9LUD4\|R18A3_ARATH | 60S ribosomal protein L18a-3 | IPR023573 (Ribosomal protein L18a/LX), IPR028877 (50S ribosomal protein L18Ae/60S ribosomal protein L20 and L18a) | GO:0003735 (structural constituent of ribosome), GO:0005840 (ribosome), GO:0006412 (translation) |
| Niben101Scf07004g01004.1 | 174 | 231 | 0.563 | 0.009714094 | AT1G47420.1 | succinate dehydrogenase 5 LENGTH=257 | IPR025397 (Protein of unknown function DUF4370) | NA |
| Niben101Scf08680g01009.1 | 266 | 355 | 0.562 | 0.024270484 | sp\|Q98G96\|CLPB_RHILO | Chaperone protein ClpB | IPR001270 (ClpA/B family), IPR023150 (Double Clp-N motif), IPR027417 (P-loop containing nucleoside triphosphate hydrolase), IPR028299 (ClpA/B, conserved site 2) | GO:0005524 (ATP binding), GO:0005737 (cytoplasm), GO:0009408 (response to heat), GO:0016485 (protein processing), GO:0019538 (protein metabolic process) |
| Niben101Scf15689g00030.1 | 237 | 315 | 0.562 | 0.014918712 | sp\|P92949\|FRO2_ARATH | Ferric reduction oxidase 2 | IPR013121 (Ferric reductase, NAD binding), IPR013130 (Ferric reductase transmembrane component-like domain), IPR017938 (Riboflavin synthase-like beta-barrel) | GO:0016491 (oxidoreductase activity), GO:0055114 (oxidation-reduction process) |
| Niben101Scf01849g03015.1 | 115 | 153 | 0.562 | 0.027723292 | sp\|Q9S7M9\|P24B2_ARATH | Transmembrane emp24 domain-containing protein p24beta2 | NA | NA |
| Niben101Scf08777g03003.1 | 101 | 134 | 0.561 | 0.031146351 | AT1G15120.2 | Ubiquinol-cytochrome C reductase hinge protein LENGTH=166 | IPR003422 (Cytochrome b-c1 complex, subunit 6), IPR023184 (Ubiquinol-cytochrome C reductase hinge domain) | GO:0006122 (mitochondrial electron transport, ubiquinol to cytochrome c), GO:0008121 (ubiquinol-cytochrome-c reductase activity) |
| Niben101Scf05017g06019.1 | 1916 | 2537 | 0.555 | 0.006478884 | sp\|Q4V0I4\|F16PA_XANC8 | Fructose-1,6-bisphosphatase class 1 | IPR000146 (Fructose-1,6-bisphosphatase class 1/Sedoheputulose-1,7-bisphosphatase) | GO:0005975 (carbohydrate metabolic process), GO:0042132 (fructose 1,6-bisphosphate 1-phosphatase activity), GO:0042578 (phosphoric ester hydrolase activity) |
| Niben101Scf16833g00004.1 | 267 | 354 | 0.554 | 0.025719435 | sp\|O23184\|CML19_ARATH | Calcium-binding protein CML19 | IPR011992 (EF-hand domain pair) | GO:0005509 (calcium ion binding) |
| Niben101Scf03446g12007.1 | 103 | 136 | 0.554 | 0.034283145 | sp\|B7M3W3\|YBAB_ECO8A | Nucleoid-associated protein YbaB | IPR004401 (Nucleoid-associated protein YbaB) | NA |
| Niben101Scf08151g00005.1 | 177 | 234 | 0.553 | 0.019464671 | sp\|Q51876\|LEP_PHOLA | Signal peptidase I | IPR000223 (Peptidase S26A, signal peptidase I), IPR015927 (Peptidase S24/S26A/S26B/S26C), IPR028360 (Peptidase S24/S26, beta-ribbon domain) | GO:0006508 (proteolysis), GO:0008236 (serine-type peptidase activity), GO:0016020 (membrane), GO:0016021 (integral component of membrane) |
| Niben101Scf02237g01007.1 | 163 | 215 | 0.553 | 0.034022357 | sp\|P56533\|BADH_GADMC | Betaine aldehyde dehydrogenase | IPR016161 (Aldehyde/histidinol dehydrogenase), IPR029510 (Aldehyde dehydrogenase, glutamic acid active site) | GO:0008152 (metabolic process), GO:0016491 (oxidoreductase activity), GO:0016620 (oxidoreductase activity, acting on the aldehyde or oxo group of donors, NAD or NADP as acceptor), GO:0055114 (oxidation-reduction process) |
| Niben101Scf04926g04006.1 | 164 | 214 | 0.553 | 0.0374643 | sp\|Q885L1\|PHS_PSESM | Pterin-4-alpha-carbinolamine dehydratase | IPR001533 (Transcriptional coactivator/pterin dehydratase) | GO:0006729 (tetrahydrobiopterin biosynthetic process), GO:0008124 (4-alpha-hydroxytetrahydrobiopterin dehydratase activity) |
| Niben101Scf02638g00005.1 | 96 | 127 | 0.552 | 0.045840285 | NA | Unknown protein | NA | NA |
| Niben101Scf00472g00003.1 | 175 | 229 | 0.546 | 0.048521773 | sp\|Q9XHS0\|RS12_HORVU | 40S ribosomal protein S12 | IPR000530 (Ribosomal protein S12e), IPR029064 (50S ribosomal protein L30e-like) | GO:0003735 (structural constituent of ribosome), GO:0005622 (intracellular), GO:0005840 (ribosome), GO:0006412 (translation) |
| Niben101Scf13103g01029.1 | 931 | 1217 | 0.545 | 0.021250116 | AT4G38970.1 | fructose-bisphosphate aldolase 2 LENGTH=398 | IPR000741 (Fructose-bisphosphate aldolase, class-I), IPR013785 (Aldolase-type TIM barrel), IPR029768 (Fructose-bisphosphate aldolase class-I active site) | GO:0003824 (catalytic activity), GO:0004332 (fructose-bisphosphate aldolase activity), GO:0006096 (glycolytic process) |
| Niben101Scf02455g01011.1 | 181 | 235 | 0.545 | 0.026903153 | sp\|P31414\|AVP1_ARATH | Pyrophosphate-energized vacuolar membrane proton pump 1 | IPR004131 (Pyrophosphate-energised proton pump) | GO:0004427 (inorganic diphosphatase activity), GO:0009678 (hydrogen-translocating pyrophosphatase activity), GO:0015992 (proton transport), GO:0016020 (membrane) |
| Niben101Scf04439g01023.1 | 122 | 160 | 0.545 | 0.032570615 | gb\|KEH40500.1\| | zinc finger CCCH domain protein, putative [Medicago truncatula] | NA | NA |
| Niben101Scf02558g00020.1 | 155 | 203 | 0.542 | 0.024642948 | sp\|P62980\|RS27A_SOLLC | Ubiquitin-40S ribosomal protein S27a | IPR011332 (Zinc-binding ribosomal protein) | GO:0003735 (structural constituent of ribosome), GO:0005622 (intracellular), GO:0005840 (ribosome), GO:0006412 (translation) |
| Niben101Scf01385g00004.1 | 83 | 108 | 0.542 | 0.049943648 | gb\|AES94740.2\| | transcription factor [Medicago truncatula] | IPR006869 (Domain of unknown function DUF547), IPR025757 (Ternary complex factor MIP1, leucine-zipper) | NA |
| Niben101Scf04174g05001.1 | 3253 | 4221 | 0.541 | 0.014480119 | sp\|A1AHE2\|LLDD_ECOK1 | L-lactate dehydrogenase | IPR012133 (Alpha-hydroxy acid dehydrogenase, FMN-dependent), IPR013785 (Aldolase-type TIM barrel) | GO:0003824 (catalytic activity), GO:0010181 (FMN binding), GO:0016491 (oxidoreductase activity), GO:0055114 (oxidation-reduction process) |
| Niben101Scf05417g03027.1 | 141 | 185 | 0.541 | 0.045925926 | AT4G23100.1 | glutamate-cysteine ligase LENGTH=522 | IPR006336 (Glutamate--cysteine ligase, GCS2) | GO:0004357 (glutamate-cysteine ligase activity), GO:0006750 (glutathione biosynthetic process), GO:0042398 (cellular modified amino acid biosynthetic process) |
| Niben101Scf09075g03002.1 | 3054 | 4025 | 0.539 | 0.028338156 | AT5G65730.1 | xyloglucan endotransglucosylase/hydrolase 6 LENGTH=292 | IPR013320 (Concanavalin A-like lectin/glucanase domain), IPR016455 (Xyloglucan endotransglucosylase/hydrolase) | GO:0004553 (hydrolase activity, hydrolyzing O-glycosyl compounds), GO:0005618 (cell wall), GO:0005975 (carbohydrate metabolic process), GO:0006073 (cellular glucan metabolic process), GO:0016762 (xyloglucan:xyloglucosyl transferase activity), GO:0048046 (apoplast) |
| Niben101Scf14882g01006.1 | 188 | 246 | 0.538 | 0.047049432 | sp\|Q18GF3\|RS19_HALWD | 30S ribosomal protein S19 | IPR002222 (Ribosomal protein S19/S15), IPR023575 (Ribosomal protein S19, superfamily) | GO:0003723 (RNA binding), GO:0003735 (structural constituent of ribosome), GO:0005840 (ribosome), GO:0006412 (translation), GO:0015935 (small ribosomal subunit) |
| Niben101Scf03437g02009.1 | 367 | 478 | 0.536 | 0.041541594 | emb\|CDX81747.1\| | BnaC08g38540D [Brassica napus] | IPR013328 (Dehydrogenase, multihelical) | GO:0016491 (oxidoreductase activity), GO:0016616 (oxidoreductase activity, acting on the CH-OH group of donors, NAD or NADP as acceptor), GO:0050662 (coenzyme binding), GO:0055114 (oxidation-reduction process) |
| Niben101Scf06091g07004.1 | 1089 | 1408 | 0.535 | 0.024016409 | sp\|Q9LND8\|ADSL2_ARATH | Delta-9 desaturase-like 2 protein | IPR015876 (Fatty acid desaturase, type 1, core) | GO:0006629 (lipid metabolic process), GO:0016717 (oxidoreductase activity, acting on paired donors, with oxidation of a pair of donors resulting in the reduction of molecular oxygen to two molecules of water), GO:0055114 (oxidation-reduction process) |
| Niben101Scf07123g00001.1 | 2085 | 2681 | 0.534 | 0.030797601 | sp\|Q56YA5\|SGAT_ARATH | Serine--glyoxylate aminotransferase | IPR015424 (Pyridoxal phosphate-dependent transferase), IPR024169 (Serine-pyruvate aminotransferase/2-aminoethylphosphonate-pyruvate transaminase) | GO:0003824 (catalytic activity), GO:0008152 (metabolic process), GO:0030170 (pyridoxal phosphate binding) |
| Niben101Scf07493g03007.1 | 315 | 409 | 0.534 | 0.044163837 | sp\|P0CB38\|PAB4L_HUMAN | Polyadenylate-binding protein 4-like | IPR006515 (Polyadenylate binding protein, human types 1, 2, 3, 4), IPR012677 (Nucleotide-binding alpha-beta plait domain) | GO:0000166 (nucleotide binding), GO:0003676 (nucleic acid binding), GO:0003723 (RNA binding) |
| Niben101Scf09348g00004.1 | 515 | 665 | 0.532 | 0.029855212 | emb\|CBY05740.1\| | ASR2 protein [Solanum chilense] | IPR003496 (ABA/WDS induced protein) | GO:0006950 (response to stress) |
| Niben101Scf00800g04004.1 | 266 | 347 | 0.53 | 0.047109814 | AT4G02930.1 | GTP binding Elongation factor Tu family protein LENGTH=454 | IPR005225 (Small GTP-binding protein domain), IPR006298 (GTP-binding protein TypA), IPR009000 (Translation protein, beta-barrel domain), IPR027417 (P-loop containing nucleoside triphosphate hydrolase) | GO:0003924 (GTPase activity), GO:0005525 (GTP binding) |
| Niben101Scf00414g06008.1 | 252 | 326 | 0.527 | 0.027459737 | sp\|Q1CZI7\|HTPG_MYXXD | Chaperone protein HtpG | IPR001404 (Heat shock protein Hsp90 family) | GO:0005524 (ATP binding), GO:0006457 (protein folding), GO:0006950 (response to stress), GO:0051082 (unfolded protein binding) |
| Niben101Scf16508g00001.1 | 615 | 801 | 0.526 | 0.037179774 | sp\|Q5HL01\|SAT_STAEQ | Sulfate adenylyltransferase | IPR002650 (Sulphate adenylyltransferase), IPR014729 (Rossmann-like alpha/beta/alpha sandwich fold), IPR015947 (PUA-like domain) | GO:0000103 (sulfate assimilation), GO:0004781 (sulfate adenylyltransferase (ATP) activity) |
| Niben101Scf07639g05011.1 | 263 | 341 | 0.526 | 0.047464534 | sp\|B0JUI1\|CH10_MICAN | 10 kDa chaperonin | IPR020818 (Chaperonin Cpn10) | GO:0005737 (cytoplasm), GO:0006457 (protein folding) |
| Niben101Scf10608g03010.1 | 258 | 334 | 0.525 | 0.035127249 | AT3G23820.1 | UDP-D-glucuronate 4-epimerase 6 LENGTH=460 | IPR001509 (NAD-dependent epimerase/dehydratase, N-terminal domain), IPR016040 (NAD(P)-binding domain) | GO:0003824 (catalytic activity), GO:0050662 (coenzyme binding) |
| Niben101Scf11906g00004.1 | 412 | 530 | 0.524 | 0.038297417 | sp\|A7RK30\|QORL2_NEMVE | Quinone oxidoreductase-like protein 2 homolog | IPR002085 (Alcohol dehydrogenase superfamily, zinc-type), IPR016040 (NAD(P)-binding domain), IPR020843 (Polyketide synthase, enoylreductase domain) | GO:0008270 (zinc ion binding), GO:0016491 (oxidoreductase activity), GO:0055114 (oxidation-reduction process) |
| Niben101Scf05368g03015.1 | 809 | 1040 | 0.523 | 0.02248233 | sp\|Q06721\|RCA_ANASC | Ribulose bisphosphate carboxylase/oxygenase activase | IPR003959 (ATPase, AAA-type, core), IPR027417 (P-loop containing nucleoside triphosphate hydrolase) | GO:0005524 (ATP binding) |
| Niben101Scf28250g00015.1 | 200 | 259 | 0.523 | 0.035416944 | AT1G47480.1 | alpha/beta-Hydrolases superfamily protein LENGTH=314 | IPR029058 (Alpha/Beta hydrolase fold) | GO:0008152 (metabolic process), GO:0016787 (hydrolase activity) |
| Niben101Scf31945g00009.1 | 292 | 377 | 0.52 | 0.041045315 | sp\|Q8RWD0\|COL16_ARATH | Zinc finger protein CONSTANS-LIKE 16 | IPR000315 (Zinc finger, B-box), IPR010402 (CCT domain) | GO:0005515 (protein binding), GO:0005622 (intracellular), GO:0008270 (zinc ion binding) |
| Niben101Scf00653g01009.1 | 707 | 913 | 0.519 | 0.020475969 | AT5G16400.1 | thioredoxin F2 LENGTH=185 | IPR005746 (Thioredoxin), IPR012336 (Thioredoxin-like fold) | GO:0006662 (glycerol ether metabolic process), GO:0015035 (protein disulfide oxidoreductase activity), GO:0045454 (cell redox homeostasis) |
| Niben101Scf03737g06007.1 | 10799 | 14034 | 0.516 | 0.04344976 | AT2G10940.1 | Bifunctional inhibitor/lipid-transfer protein/seed storage 2S albumin superfamily protein LENGTH=291 | IPR016140 (Bifunctional inhibitor/plant lipid transfer protein/seed storage helical domain) | NA |
| Niben101Scf02864g04008.1 | 2190 | 2800 | 0.515 | 0.016858947 | AT4G38970.1 | fructose-bisphosphate aldolase 2 LENGTH=398 | IPR000741 (Fructose-bisphosphate aldolase, class-I), IPR013785 (Aldolase-type TIM barrel), IPR029768 (Fructose-bisphosphate aldolase class-I active site) | GO:0003824 (catalytic activity), GO:0004332 (fructose-bisphosphate aldolase activity), GO:0006096 (glycolytic process) |
| Niben101Scf01365g06006.1 | 178 | 230 | 0.515 | 0.037713071 | sp\|Q4P6E2\|EIF3I_USTMA | Eukaryotic translation initiation factor 3 subunit I | IPR015943 (WD40/YVTN repeat-like-containing domain), IPR020472 (G-protein beta WD-40 repeat), IPR027525 (Eukaryotic translation initiation factor 3 subunit I) | GO:0003743 (translation initiation factor activity), GO:0005515 (protein binding), GO:0005737 (cytoplasm), GO:0005852 (eukaryotic translation initiation factor 3 complex) |
